# Molecular evidence for segmental duplication across chromosomes of soybean using transcription factor gene family

**DOI:** 10.1101/2021.12.20.473405

**Authors:** Manoj Kumar Srivastava, Gyanesh Kumar Satpute

## Abstract

Duplication of genome is an important genetic innovation. Large genome size (1.1 Gb) along with ancient and recent duplication events make the soybean genome more complex. Analyzing the distribution and duplication event in soybean transcription family genes, the segmental duplication within chromosomes was revealed. Our study provides a strong evidence that the large segmental duplication event in genome architecture and evolution of soybean genome using simple method of sequence and order analysis of TF genes. Finally, a scheme for interrelationship of different chromosomes has been proposed.

## Introduction

Duplication of genome is an important event leading to evolution of organism. Gene and genome duplication may contribute to evolution and domestication of crops. The role of genome duplication in present architecture/ topology of soybean genome have vital influence on agronomic traits, yield potential and adaptation of crop plants. The redundant copy of gene arising by the duplication accumulates the beneficial mutations resulting in new function of duplicated gene product. Thus gene duplication provides a source of plasticity to genome for adapting to changing environment. Large genome size (1.1 Gb) along with ancient and recent duplication events make the soybean genome more complex. Having complex genome structure makes it rather difficult to design and develop effective breeding strategies in soybean for desired traits. Several QTLs for various traits and linkage maps have been developed for 20 chromosomes of soybean genome. Translocation, inversion, deletion and duplication play important roles in creating small duplication events.

In flowering plants, two ancestral whole genome duplication (WGD) is reported. In soybean, two additional sequential WGDs are established: one had occurred 59 MYA in the common ancestor of legumes and other about 8-13 MYA in *Glycine* lineage (Schmutz et al., 2010). Due to multiple genome duplication events, the number of predicted coding genes in soybean is much higher than in *Arabidopsis* and grapes (Sterck et al., 2007; Cannon and Shoemaker, 2012). Several small blocks of homeologus retention and chromosomal arrangements are shown to exist in 20 chromosomes of soybean (Schmutz et al., 2010; Lestari et al., 2013). The segmental duplication in soybean has been reported to result in the evolution of several phenotypic traits such as disease resistance (Shin et al., 2008; Kim et al., 2009). Several QTLs associated with seed related traits, disease resistance and high content of carbohydrates, proteins and oil were reported to be conserved in the duplicated segment of the soybean genome. The major seed protein QTL is mapped on chromosome 20 (Brummer et al., 1997). This QTL have been studied using several different approaches (Wang et al., 2008; Qi et al., 2011).

Transcription factors are DNA binding regulatory proteins, which interact with other proteins and regulate the process of transcription. Most of the TF genes have several family members. In soybean, there are 57 families of TF genes reported with are distributed at 3747 locus in 20 chromosomes. In the present study the duplication and order of these TF genes were analyzed.

## Materials and methods

All the transcription factor genes of soybean were downloaded from Plant transcription factor database (http://planttfdb.gao-lab.org/index.php?sp=Gma). Duplication of TF genes across all chromosomes was studied by BLAST analysis of each gene against total transcript database (Wm82.a2.v1 Transcript Sequences). The top BLAST hit having E value of 0.0 was considered as transcript arising of duplicated gene. The E value having more than 0.0 was considered as non-duplicated gene product. The chromosome wise duplicated gene pair was recorded. The putative chromosome pair for duplicated segment was observed for similarity in order of TF genes.

## Results and discussion

Most of the high copy number TF genes are distributed among all the chromosomes, however, the relative number of TF varied for each chromosome (Table 1). Five out of 20 chromosomes (Chr 2, Chr 6, Chr 8, Chr 10 and Chr 13) contain more than 200 loci of TF genes. The chromosome 16 has the lowest number of TF genes while chr 13 has the maximum number of TF genes. No specific pattern of chromosomal location for different TF gene family was observed i.e. different TF gene family have different distribution pattern.

**Table 1:**
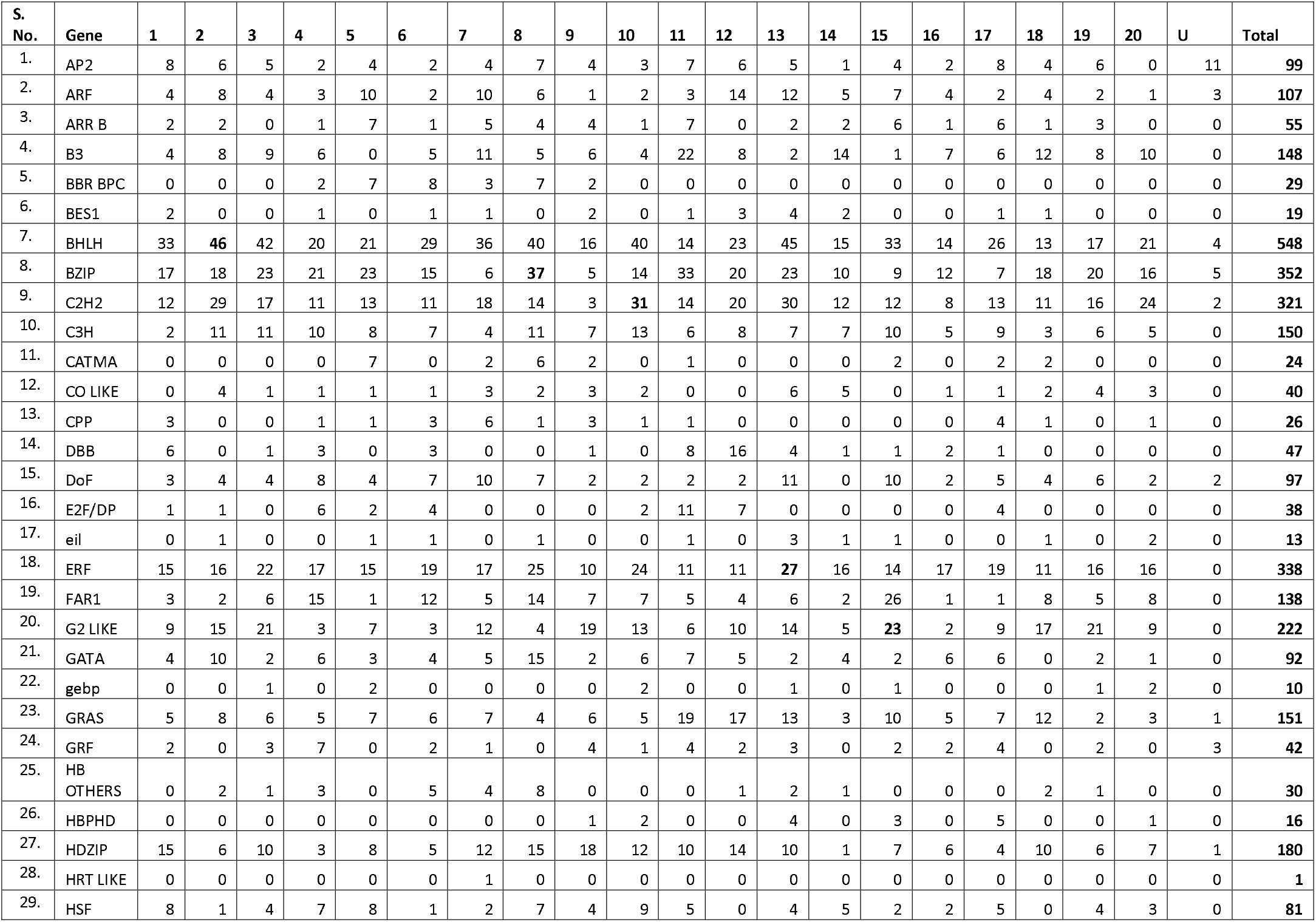
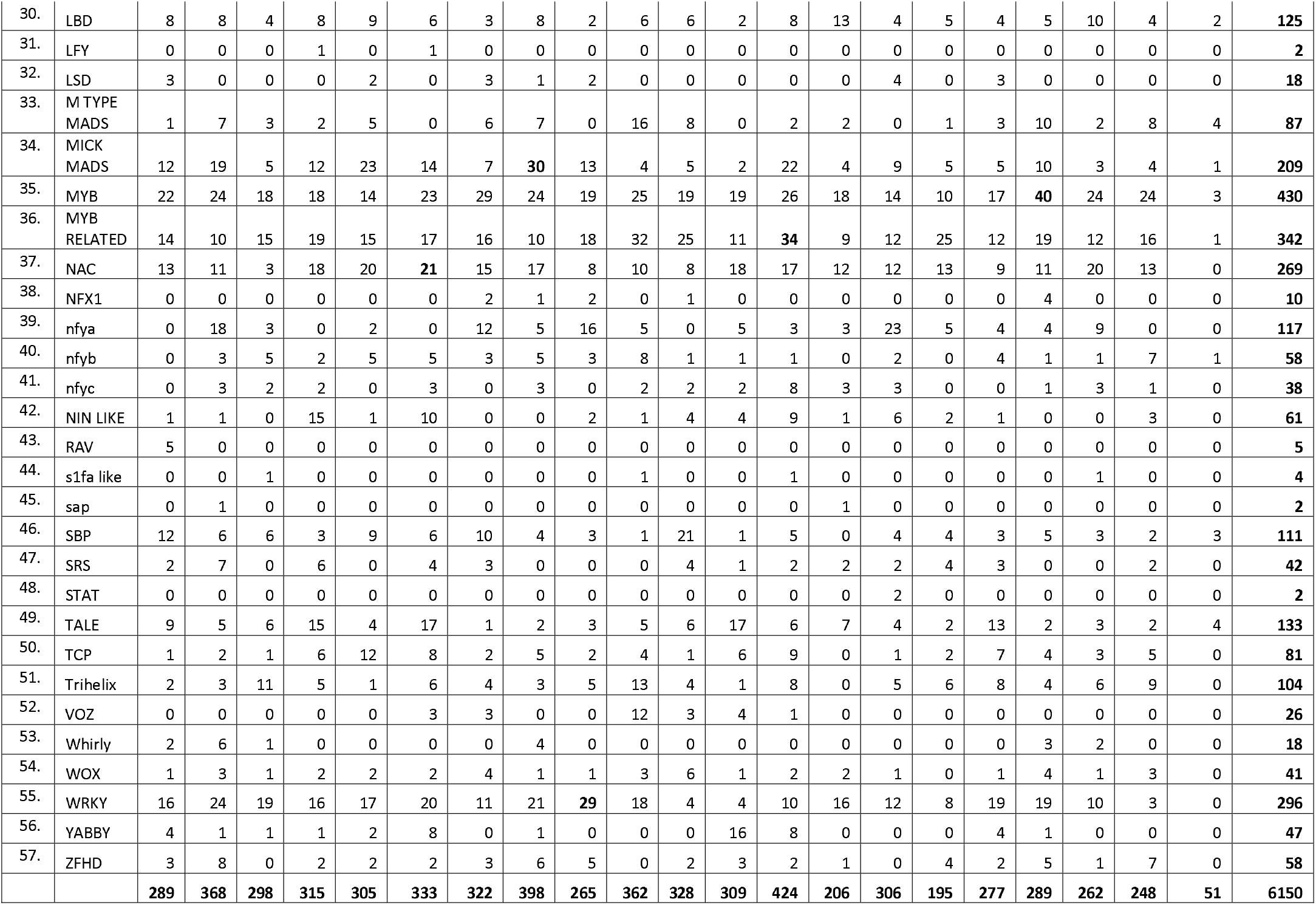
Distribution of TF gene products in soybean chromosomes

Locus frequency of less than 40% represents the TF genes which produce more copies of products from the same gene (Table 2 and S1). These include BBR-BCP (31.03%; 10/29), E2F/DP (36.84%; 14/38), VOZ (23.08%; 6/28), Whirly (38.89%; 7/18) and YABBY (34.04%; 17/47). Five out of 57 families have same number of gene products and gene locus (HRT-like, LFY, RAV, S1Fa-like and SAP). These are also low copy number transcription factor genes. The genes having locus to gene product ratio of more than 0.8 produce less gene product variants. These are BES1 (0.84), Dof (0.82), EIL (0.92), ERF (0.88), GeBP (0.9), M-type_MADS (0.95), WOX (0.80) and ZF-HD (0.89).

**Table 2:**
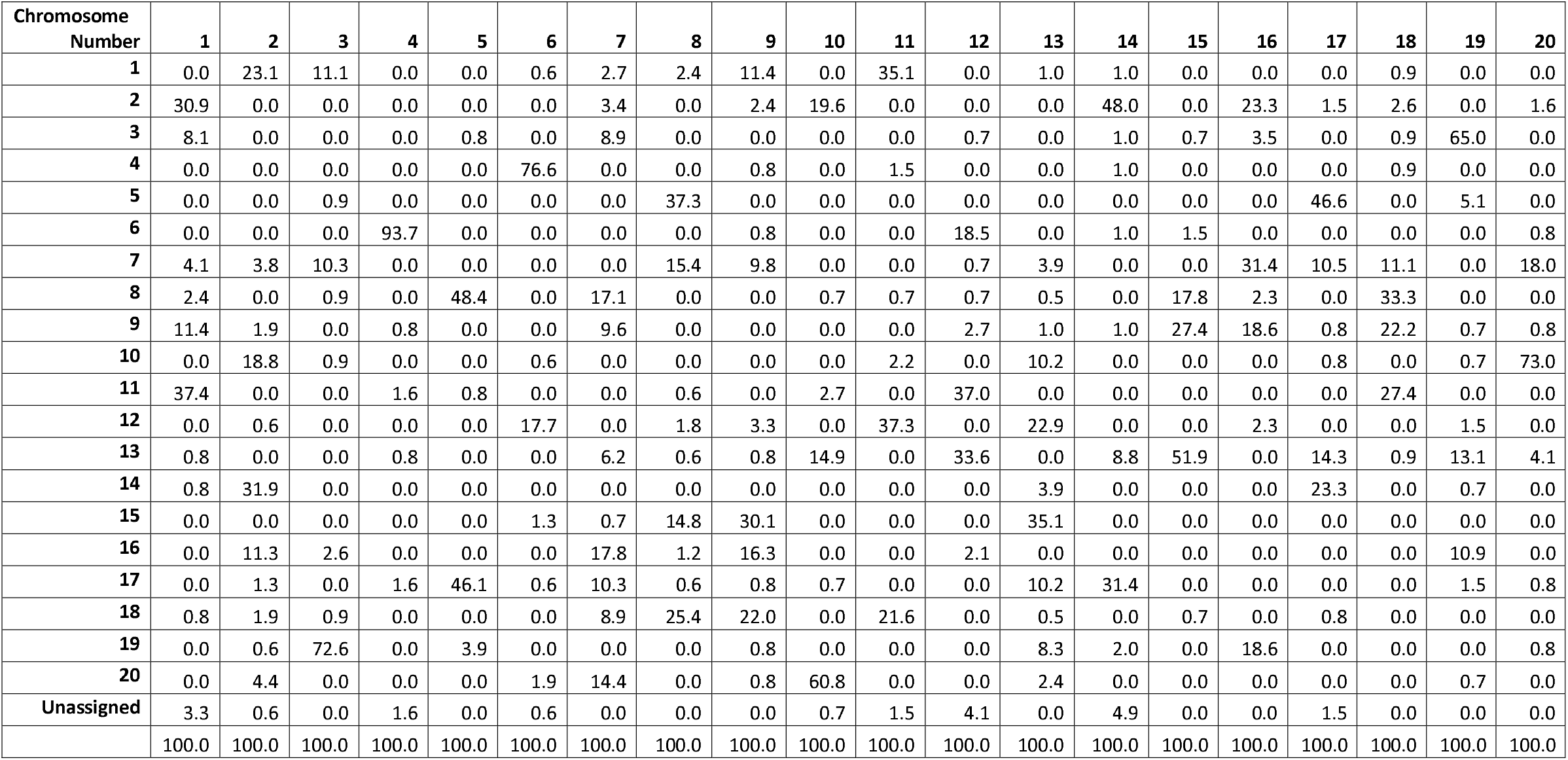
Distribution of duplicated genes (%) in soybean chromosome

This analysis for cDNA of all the 3747 locus genes revealed that at least 72.63% of the TF genes were duplicated paralog pair (Table 3 and S2). Also, there was a specific preference of duplication among various chromosomes, e.g. chromosome 1 have more duplication segments from chr 2, chr 9 and chr 11; chromosome 2 have more duplication fragments from chr 1, chr 10, chr 14 and chr 16; Chromosome 3 have major segment from chr 19 and minor contribution from chr 1 and chr 7; Chromosome 4 has almost entire (95%) TF gene duplication from chr 6. The detail distribution is given in Fig 1, Table 3 and supplementary Table S2.

**Fig 1:**
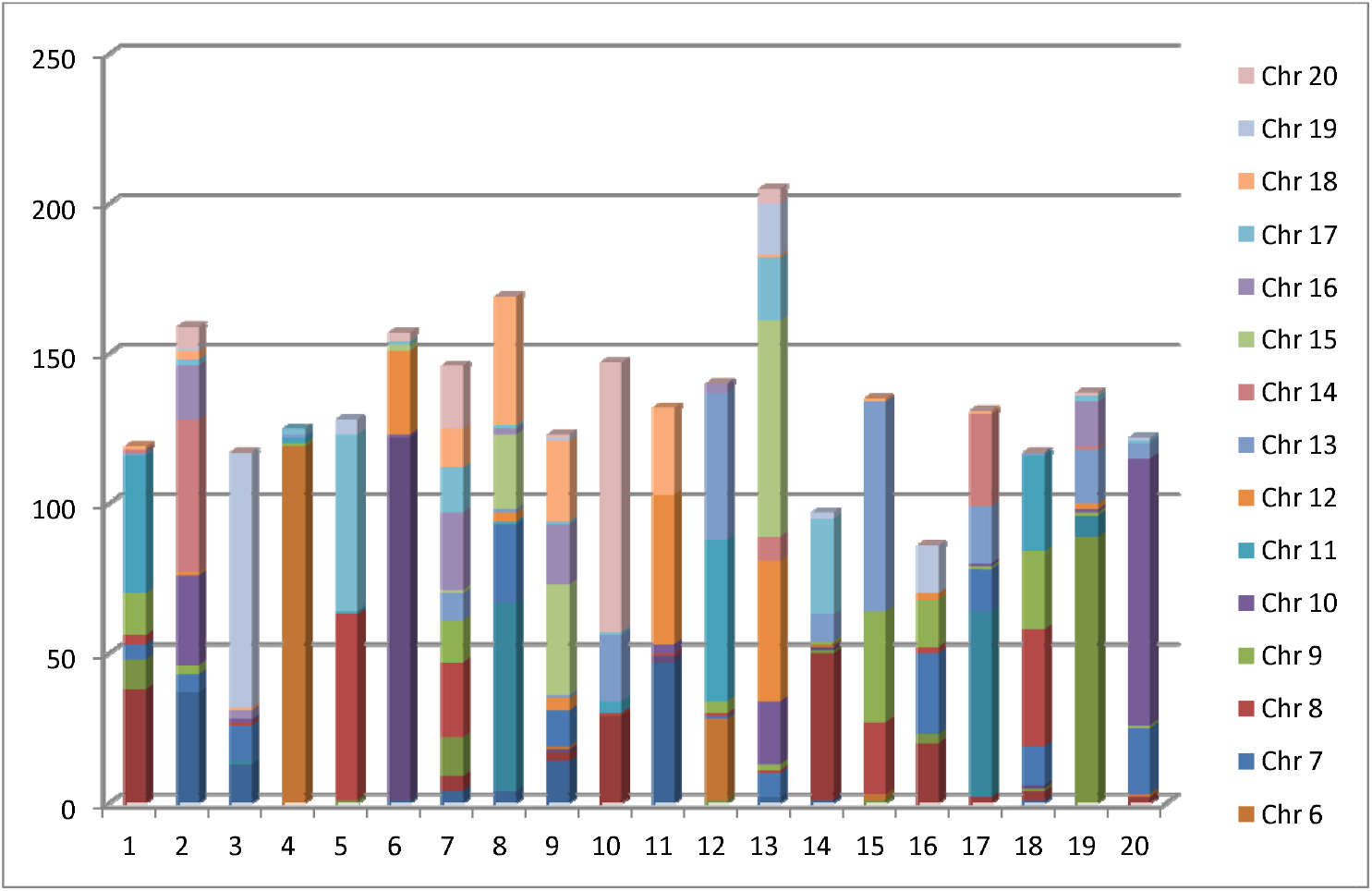
Distribution of duplicated transcription factor genes in different chromosomes of soybean

The gene order of different TFs was then compared and similarity was investigated among putative chromosome pairs (Table S3). The data supported the finding of preferential chromosome duplication. Furthermore, in the segment of similar TF genes the order of various TF genes was found to be collinear either in direct or reverse direction. This indicated that long segmental duplication event in soybean is major event in the evolution of soybean chromosome. The preferential segmental duplication is presented in Fig S1. This is in accordance with the distribution data of duplicated TF genes. However this may be noted that careful sequence analysis is required at gene to gene basis for the accumulated mutations in duplicated TF genes. This will lead to assigning new function to the duplicated TF gene.

It has been established that the recent genome duplication occurred on many soybean chromosomes (Cannon and Shoemaker, 2012). QTLs across duplicated regions of chr 4/ chr 6, chr 3/ chr 19 and chr 10/chr 20 were shown to be correlated (Lestani et al., 2013). Small rearrangements were found in duplicated homeologus regions of QTLs due to recent duplication event. Soybean gene duplication may also lead to gene regulation (Shoemaker et al., 1996). A large inversion with synteny in the corresponding regions of chr 10 and chr 20 has also been reported (Cannon et al., 2004).

Yang et al (2013) studied similarities between soybean and Arabidopsis genomes using dot-plot analysis and reported that whole genome duplication event occurred more than once during the evolution of soybean genome. About 70% of total genes in soybean genome have duplicated paralog pairs. The block of 2140 genes were found to be the largest pair of duplicated paralog gene in chromosome 3 and 19 of soybean (Yang et al., 2013). The data presented also indicated towards the ongoing duplication event in soybean genome. DNA rearrangement and codon mutations resulted in emergence of new gene sequences which may have new functions leading to better adaptation in new challenging environment.

Chromosome 1- contains 6 duplicated TF gene segments from chr 9, chr 2, chr 3 and chr 11. The largest segment was from chr 11 having 519 genes.

Chromosome 2- contains 8 distinct segments from chr 1, chr10, chr 14 and chr 16. There were four different fragments duplicated from chr 14, two in forward direction and two in reverse direction.

Chromosome 3- Block of 1355 genes is directly duplicated in chromosome 19 (1325 genes). All the TF genes are collinear. 160, 29 and 88 genes are duplicated in smaller segments and the order of TFs are collinear but in reverse and forward direction.

Chromosome 4- has 5 different segments from chr 6. The first two segments are in forward direction while last three are in reverse direction. Chromosome 6 has an additional segment from chromosome 12 in reverse direction.

Chromosome 5- has seven smaller segments, four from chromosome 17 and three from chromosome 8. Three segments of chr 17 were duplicated in reverse direction. All the three segments duplicated from chromosome 8 were in forward direction. Most of the TF gene order was conserved in duplicated segments.

The duplicated segments have almost similar number of transcription factor genes (Table 4). However, the number of embedded genes is quite variable in respective duplicated segments (Fig S1). There are 48 segmental pairs detected having 4-107 conserved TF genes and 29-1431 embedded genes. Only segment number 36 has identical 20 TF genes and 423 embedded genes on chr 9 and chr 16. Other segment pairs have variable number of genes. This may be due to ongoing translocation, mutations, and deletions of gene within the segments. Based on these segment pairs an inter-relation of various chromosomes of soybean has been proposed and shown as Fig 2.

**Figure 2:**
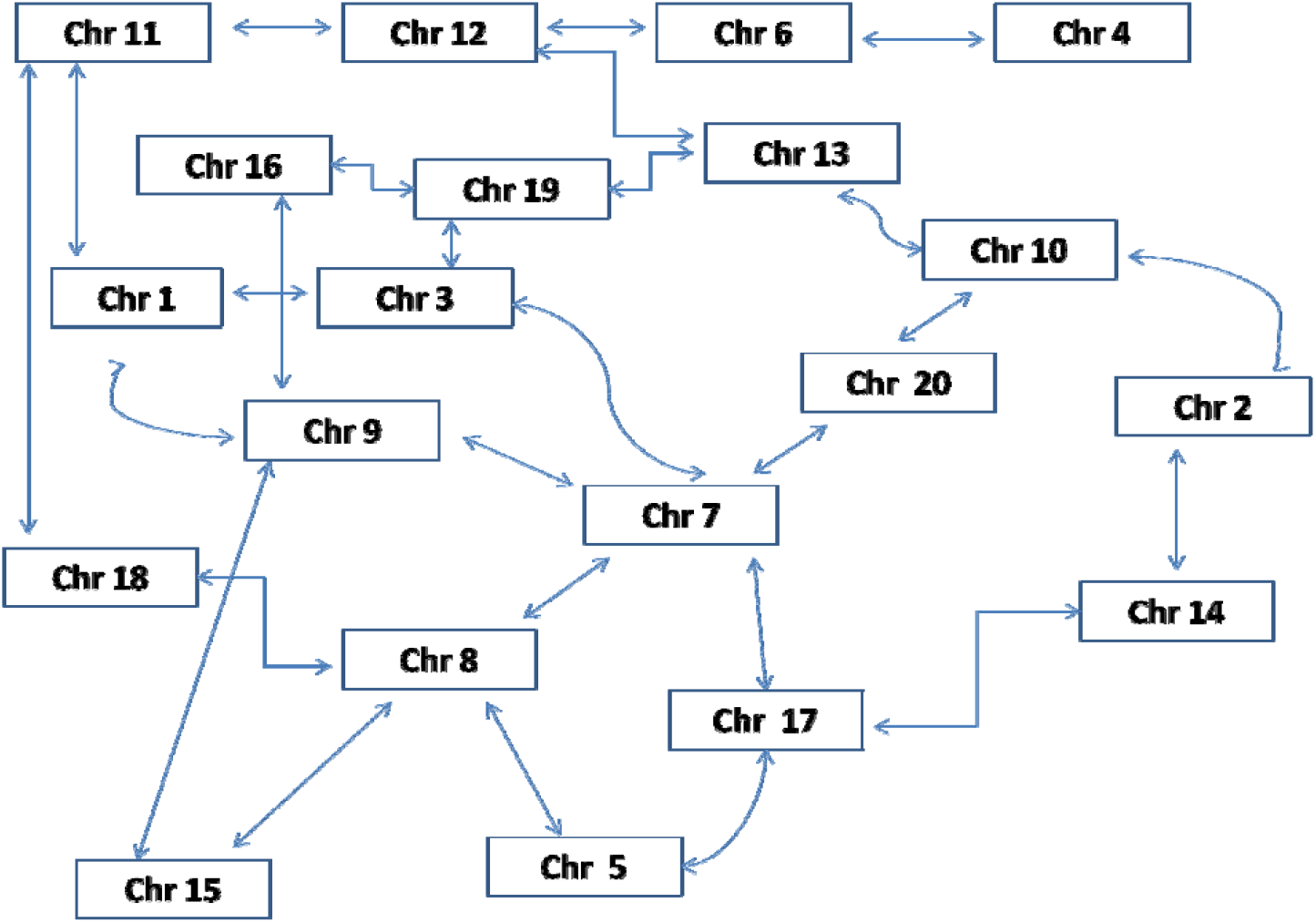
Inter relationship of different soybean chromosomes.

**Table 3:** Distribution of duplicated genes in soybean chromosome

| Chromosome Number | 1 | 2 | 3 | 4 | 5 | 6 | 7 | 8 | 9 | 10 | 11 | 12 | 13 | 14 | 15 | 16 | 17 | 18 | 19 | 20 | Unassigned | Not duplicated | TOTAL |
| --- | --- | --- | --- | --- | --- | --- | --- | --- | --- | --- | --- | --- | --- | --- | --- | --- | --- | --- | --- | --- | --- | --- | --- |
| **1** | 0 | 37 | 13 | 0 | 0 | 1 | 4 | 4 | 14 | 0 | 47 | 0 | 2 | 1 | 0 | 0 | 0 | 1 | 0 | 0 | 4 | 44 | 172 |
| **2** | 38 | 0 | 0 | 0 | 0 | 0 | 5 | 0 | 3 | 29 | 0 | 0 | 0 | 49 | 0 | 20 | 2 | 3 | 0 | 2 | 1 | 69 | 221 |
| **3** | 10 | 0 | 0 | 0 | 1 | 0 | 13 | 0 | 0 | 0 | 0 | 1 | 0 | 1 | 1 | 3 | 0 | 1 | 89 | 0 | 0 | 57 | 177 |
| **4** | 0 | 0 | 0 | 0 | 0 | 121 | 0 | 0 | 1 | 0 | 2 | 0 | 0 | 1 | 0 | 0 | 0 | 1 | 0 | 0 | 2 | 60 | 188 |
| **5** | 0 | 0 | 1 | 0 | 0 | 0 | 0 | 63 | 0 | 0 | 0 | 0 | 0 | 0 | 0 | 0 | 62 | 0 | 7 | 0 | 0 | 43 | 176 |
| **6** | 0 | 0 | 0 | 119 | 0 | 0 | 0 | 0 | 1 | 0 | 0 | 27 | 0 | 1 | 2 | 0 | 0 | 0 | 0 | 1 | 1 | 58 | 210 |
| **7** | 5 | 6 | 12 | 0 | 0 | 0 | 0 | 26 | 12 | 0 | 0 | 1 | 8 | 0 | 0 | 27 | 14 | 13 | 0 | 22 | 0 | 52 | 198 |
| **8** | 3 | 0 | 1 | 0 | 62 | 0 | 25 | 0 | 0 | 1 | 1 | 1 | 1 | 0 | 24 | 2 | 0 | 39 | 0 | 0 | 0 | 70 | 230 |
| **9** | 14 | 3 | 0 | 1 | 0 | 0 | 14 | 0 | 0 | 0 | 0 | 4 | 2 | 1 | 37 | 16 | 1 | 26 | 1 | 1 | 0 | 43 | 164 |
| **10** | 0 | 30 | 1 | 0 | 0 | 1 | 0 | 0 | 0 | 0 | 3 | 0 | 21 | 0 | 0 | 0 | 1 | 0 | 1 | 89 | 0 | 74 | 221 |
| **11** | 46 | 0 | 0 | 2 | 1 | 0 | 0 | 1 | 0 | 4 | 0 | 54 | 0 | 0 | 0 | 0 | 0 | 32 | 0 | 0 | 2 | 42 | 184 |
| **12** | 0 | 1 | 0 | 0 | 0 | 28 | 0 | 3 | 4 | 0 | 50 | 0 | 47 | 0 | 0 | 2 | 0 | 0 | 2 | 0 | 6 | 38 | 181 |
| **13** | 1 | 0 | 0 | 1 | 0 | 0 | 9 | 1 | 1 | 22 | 0 | 49 | 0 | 9 | 70 | 0 | 19 | 1 | 18 | 5 | 0 | 68 | 274 |
| **14** | 1 | 51 | 0 | 0 | 0 | 0 | 0 | 0 | 0 | 0 | 0 | 0 | 8 | 0 | 0 | 0 | 31 | 0 | 1 | 0 | 5 | 39 | 136 |
| **15** | 0 | 0 | 0 | 0 | 0 | 2 | 1 | 25 | 37 | 0 | 0 | 0 | 72 | 0 | 0 | 0 | 0 | 0 | 0 | 0 | 0 | 29 | 166 |
| **16** | 0 | 18 | 3 | 0 | 0 | 0 | 26 | 2 | 20 | 0 | 0 | 3 | 0 | 0 | 0 | 0 | 0 | 0 | 15 | 0 | 0 | 31 | 118 |
| **17** | 0 | 2 | 0 | 2 | 59 | 1 | 15 | 1 | 1 | 1 | 0 | 0 | 21 | 32 | 0 | 0 | 0 | 0 | 2 | 1 | 2 | 41 | 181 |
| **18** | 1 | 3 | 1 | 0 | 0 | 0 | 13 | 43 | 27 | 0 | 29 | 0 | 1 | 0 | 1 | 0 | 1 | 0 | 0 | 0 | 0 | 50 | 170 |
| **19** | 0 | 1 | 85 | 0 | 5 | 0 | 0 | 0 | 1 | 0 | 0 | 0 | 17 | 2 | 0 | 16 | 0 | 0 | 0 | 1 | 0 | 47 | 175 |
| **20** | 0 | 7 | 0 | 0 | 0 | 3 | 21 | 0 | 1 | 90 | 0 | 0 | 5 | 0 | 0 | 0 | 0 | 0 | 1 | 0 | 0 | 46 | 174 |
| **Unassigned** | 4 | 1 | 0 | 2 | 0 | 1 | 0 | 0 | 0 | 1 | 2 | 6 | 0 | 5 | 0 | 0 | 2 | 0 | 0 | 0 | 0 | 7 | 31 |
|  |  |  |  |  |  |  |  |  |  |  |  |  |  |  |  |  |  |  |  |  |  |  | 3747 |
|  | 123 | 160 | 117 | 127 | 128 | 158 | 146 | 169 | 123 | 148 | 134 | 146 | 205 | 102 | 135 | 86 | 133 | 117 | 137 | 122 | 23 | 1008 |  |

**Table 4:** Distribution and orientation of genes in duplicated segment pairs of soybean chromosomes.

| **Chromosome Number** | **Segment number** | **Number of TF genes in the segment** | **Total number of genes in the segment** | **Chromosome number of Duplicated segment** | **Number of TF genes in the segment** | **Total number of genes in the segment** | **Orientation** |
| --- | --- | --- | --- | --- | --- | --- | --- |
| Chr 1 | 1 | 11 | 130 | Chr 9 | 11 | 117 | REVERSE |
|  | 2 | 16 | 214 | Chr 2 | 16 | 227 | REVERSE |
|  | 3 | 22 | 239 | Chr 2 | 22 | 203 | DIRECT |
|  | 4 | 4 | 29 | Chr 3 | 4 | 40 | DIRECT |
|  | 5 | 8 | 110 | Chr 3 | 8 | 123 | REVERSE |
|  | 6 | 59 | 630 | Chr 11 | 54 | 619 | REVERSE |
| Chr 2 | 7 | 24 | 291 | Chr 16 | 24 | 345 | DIRECT |
|  | 8 | 9 | 114 | Chr 10 | 9 | 75 | REVERSE |
|  | 9 | 17 | 224 | Chr 14 | 20 | 253 | DIRECT |
|  | 10 | 5 | 33 | Chr 14 | 5 | 20 | DIRECT |
|  | 11 | 12 | 143 | Chr 14 | 13 | 168 | REVERSE |
|  | 12 | 10 | 170 | Chr 14 | 10 | 186 | REVERSE |
| Chr 3 | 13 | 19 | 166 | Chr 7 | 14 | 88 | REVERSE |
|  | 14 | 105 | 1356 | Chr 19 | 105 | 1326 | DIRECT |
| Chr 4 | 15 | 37 | 407 | Chr 6 | 35 | 418 | DIRECT |
|  | 16 | 41 | 586 | Chr 6 | 36 | 574 | DIRECT |
|  | 17 | 25 | 255 | Chr 6 | 25 | 247 | REVERSE |
|  | 18 | 28 | 350 | Chr 6 | 27 | 354 | REVERSE |
|  | 19 | 18 | 293 | Chr 6 | 19 | 319 | REVERSE |
| Chr 5 | 20 | 7 | 50 | Chr 17 | 7 | 53 | REVERSE |
|  | 21 | 15 | 149 | Chr 17 | 15 | 151 | REVERSE |
|  | 22 | 20 | 260 | Chr 17 | 22 | 261 | DIRECT |
|  | 23 | 14 | 99 | Chr 17 | 13 | 81 | REVERSE |
|  | 24 | 6 | 107 | Chr 8 | 6 | 104 | DIRECT |
|  | 25 | 35 | 476 | Chr 8 | 33 | 502 | DIRECT |
|  | 26 | 37 | 485 | Chr 8 | 38 | 497 | DIRECT |
| Chr 6 | 27 | 25 | 257 | Chr 12 | 25 | 269 | REVERSE |
| Chr 7 | 28 | 11 | 103 | Chr 8 | 11 | 93 | REVERSE |
|  | 29 | 28 | 243 | Chr 16 | 27 | 236 | DIRECT |
|  | 30 | 19 | 223 | Chr 9 | 17 | 215 | REVERSE |
|  | 31 | 22 | 262 | Chr 20 | 26 | 323 | DIRECT |
|  | 32 | 17 | 361 | Chr 17 | 19 | 354 | REVERSE |
| Chr 8 | 33 | 7 | 58 | Chr 15 | 6 | 44 | DIRECT |
|  | 34 | 11 | 168 | Chr 18 | 12 | 185 | DIRECT |
| Chr 9 | 35 | 46 | 942 | Chr 15 | 49 | 972 | DIRECT |
|  | 36 | 20 | 423 | Chr 16 | 20 | 423 | DIRECT |
|  | 37 | 38 | 465 | Chr 18 | 40 | 541 | REVERSE |
| Chr 10 | 38 | 35 | 342 | Chr 13 | 34 | 314 | DIRECT |
|  | 39 | 107 | 1431 | Chr 20 | 110 | 1361 | REVERSE |
| Chr 11 | 40 | 43 | 486 | Chr 12 | 37 | 460 | DIRECT |
|  | 41 | 46 | 676 | Chr 18 | 40 | 503 | REVERSE |
| Chr 12 | 42 | 40 | 441 | Chr 13 | 40 | 460 | REVERSE |
| Chr 13 | 43 | 19 | 186 | Chr 19 | 22 | 265 | REVERSE |
|  | 44 | 17 | 239 | Chr 17 | 18 | 244 | REVERSE |
|  | 45 | 40 | 488 | Chr 15 | 39 | 468 | REVERSE |
|  | 46 | 38 | 417 | Chr 15 | 42 | 420 | REVERSE |
| Chr 14 | 47 | 37 | 425 | Chr 17 | 36 | 392 | REVERSE |
| Chr 16 | 48 | 15 | 146 | Chr 19 | 15 | 192 | REVERSE |

Duplication creates genetic redundancy leading to evolutionary innovation. Over the passage of time the duplicated copy acquire a beneficial mutation resulting in retention of both copies. Alternatively the mutation in duplicated segment may make it non functional. The recognition of fact that a single protein can have a multiple catalytic or structural functions supports the contribution of gene duplication. In recent studies, the genome-wide analysis of duplication of individual transcription factors have been reported (Liu et al. 2020; Chen et al., 2019; Li et al., 2019; Ullah et al., 2019). Our results also validate these studies; however, there are some minor differences about the position of duplicated segments. Our study has provided a strong evidence that the large segmental duplication event in genome architecture and evolution of soybean genome using simple method of sequence and order analysis of TF genes. A detailed analysis of these genes using Bioinformatics tools may help in establishing the process of gene duplication in other species and genera.

## Conclusions

By analyzing the distribution and order of transcription factor genes the early mode of genome duplication was established. This method provides an easy and effective tool to study genome duplication in different species and genera. The functional analysis of duplicated genes is required for complete elucidation of the process of genome duplication.

## Data Availability

The data underlying this article are available in the article and in its online supplementary material. The raw data of transcription factor sequences can be found at http://planttfdb.gao-lab.org/index.php?sp=Gma.

## Acknowledgments

We thank Dr Sanjay Gupta, Dr Mehar H Asif and Dr. A. Radhakrishna for discussion during the preparation of the manuscript. We are thankful to Director, ICAR-Indian Institute of Soybean Research, Indore for encouragement and facility provided by for this work.

