## supplementary Fig 1 for "Molecular evidence for segmental duplication across chromosomes of soybean using transcription factor gene family"

### Slide 1
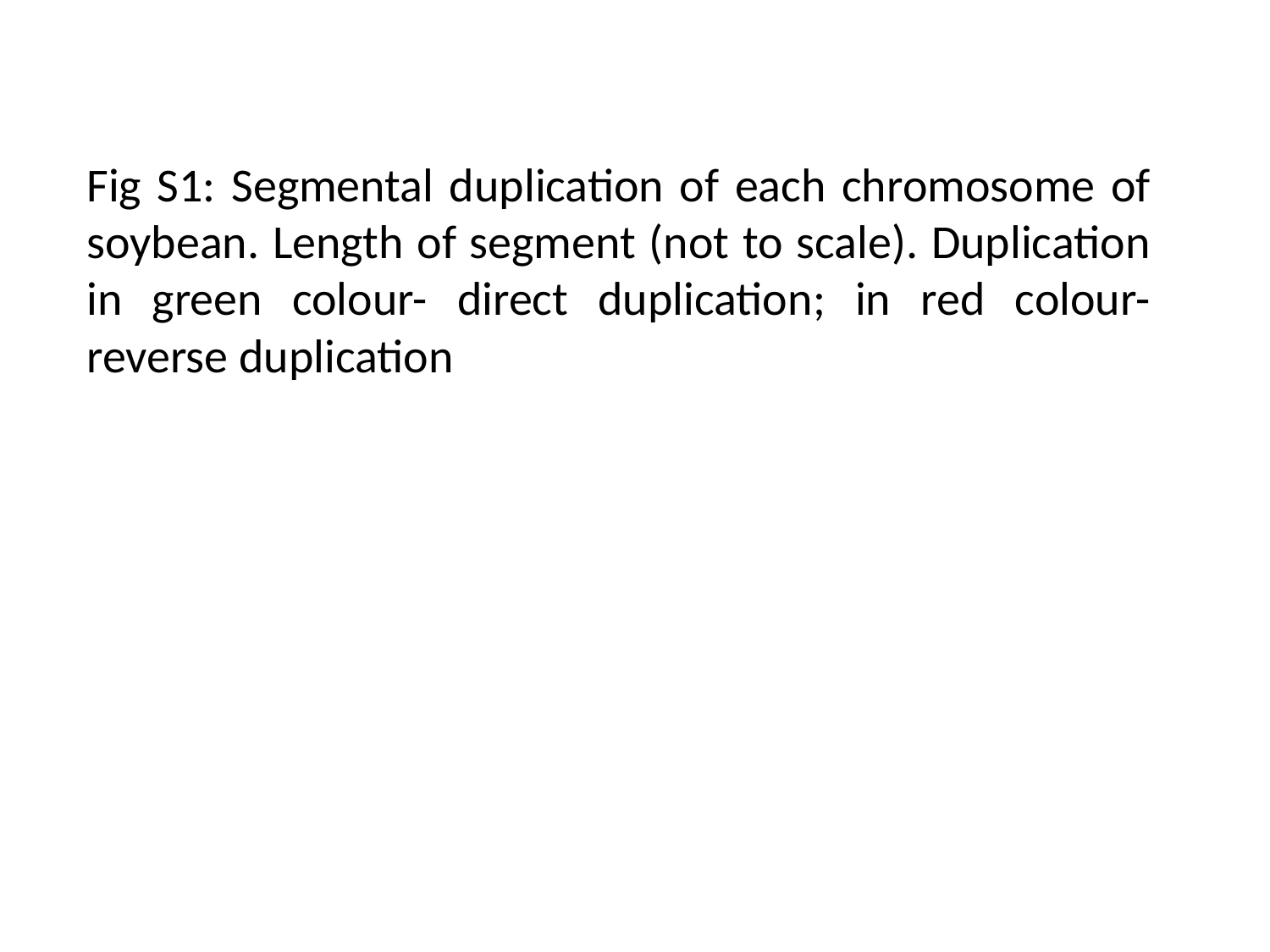

Fig S1: Segmental duplication of each chromosome of soybean. Length of segment (not to scale). Duplication in green colour- direct duplication; in red colour- reverse duplication

### Slide 2
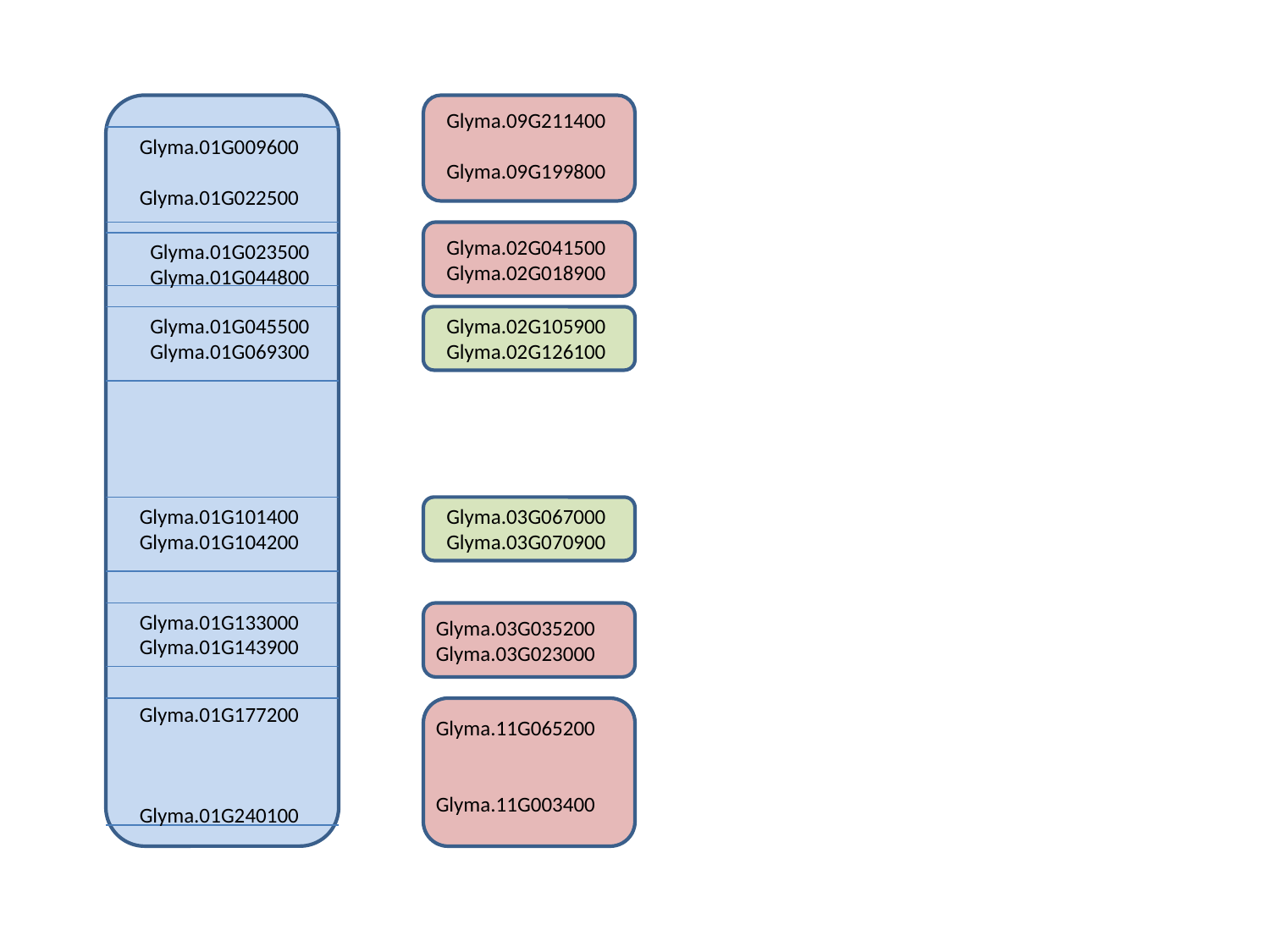

Glyma.09G211400
Glyma.09G199800
Glyma.01G009600
Glyma.01G022500
Glyma.02G041500
Glyma.02G018900
Glyma.01G023500
Glyma.01G044800
Glyma.01G045500
Glyma.01G069300
Glyma.02G105900
Glyma.02G126100
Glyma.01G101400
Glyma.01G104200
Glyma.03G067000
Glyma.03G070900
Glyma.01G133000
Glyma.01G143900
Glyma.03G035200
Glyma.03G023000
Glyma.01G177200
Glyma.01G240100
Glyma.11G065200
Glyma.11G003400

### Slide 3
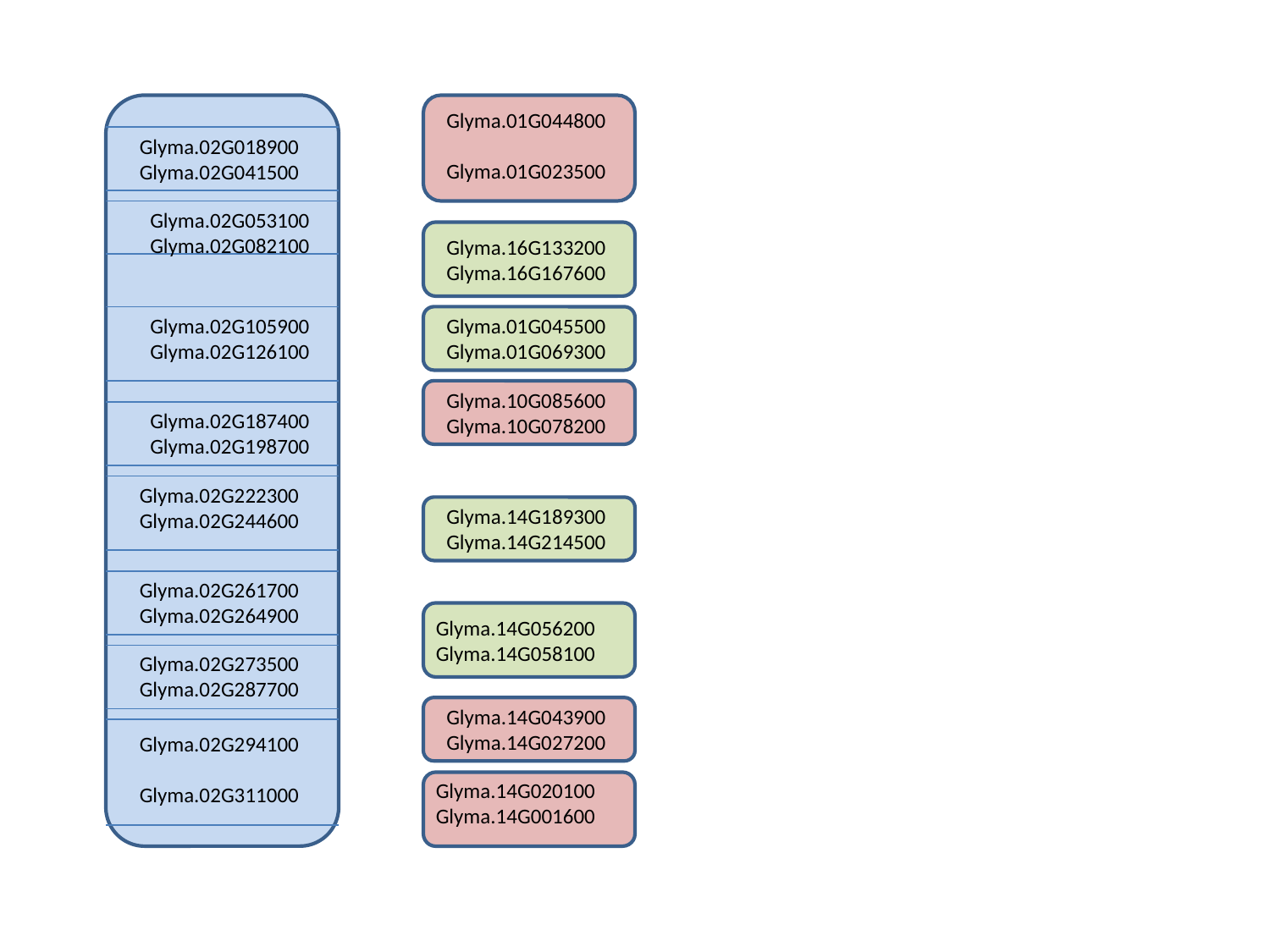

Glyma.01G044800
Glyma.01G023500
Glyma.02G018900
Glyma.02G041500
Glyma.02G053100
Glyma.02G082100
Glyma.16G133200
Glyma.16G167600
Glyma.02G105900
Glyma.02G126100
Glyma.01G045500
Glyma.01G069300
Glyma.10G085600
Glyma.10G078200
Glyma.02G187400
Glyma.02G198700
Glyma.02G222300
Glyma.02G244600
Glyma.14G189300
Glyma.14G214500
Glyma.02G261700
Glyma.02G264900
Glyma.14G056200
Glyma.14G058100
Glyma.02G273500
Glyma.02G287700
Glyma.14G043900
Glyma.14G027200
Glyma.02G294100
Glyma.02G311000
Glyma.14G020100
Glyma.14G001600

### Slide 4
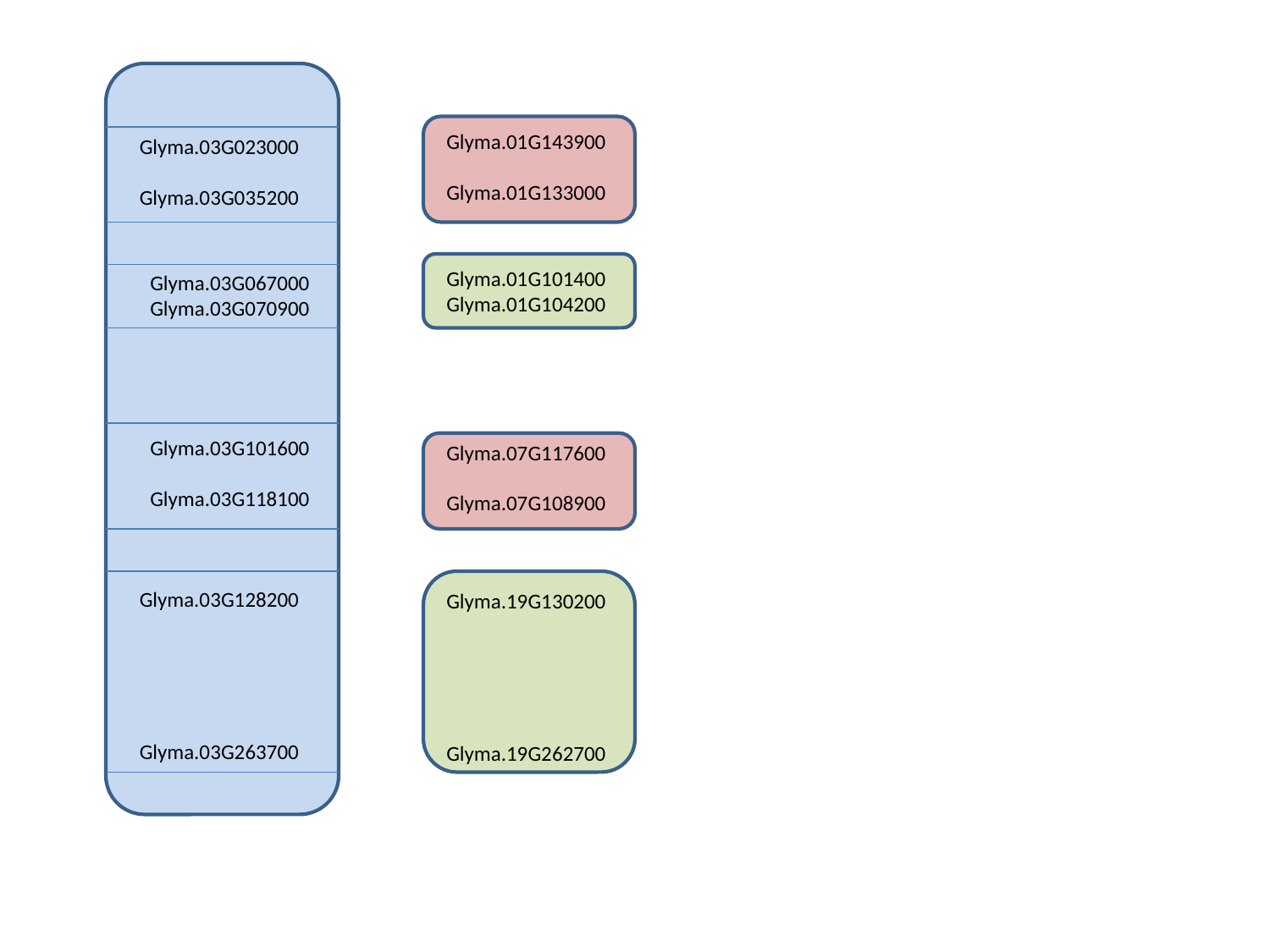

Glyma.01G143900
Glyma.01G133000
Glyma.03G023000
Glyma.03G035200
Glyma.01G101400
Glyma.01G104200
Glyma.03G067000
Glyma.03G070900
Glyma.03G101600
Glyma.03G118100
Glyma.07G117600
Glyma.07G108900
Glyma.03G128200
Glyma.03G263700
Glyma.19G130200
Glyma.19G262700

### Slide 5
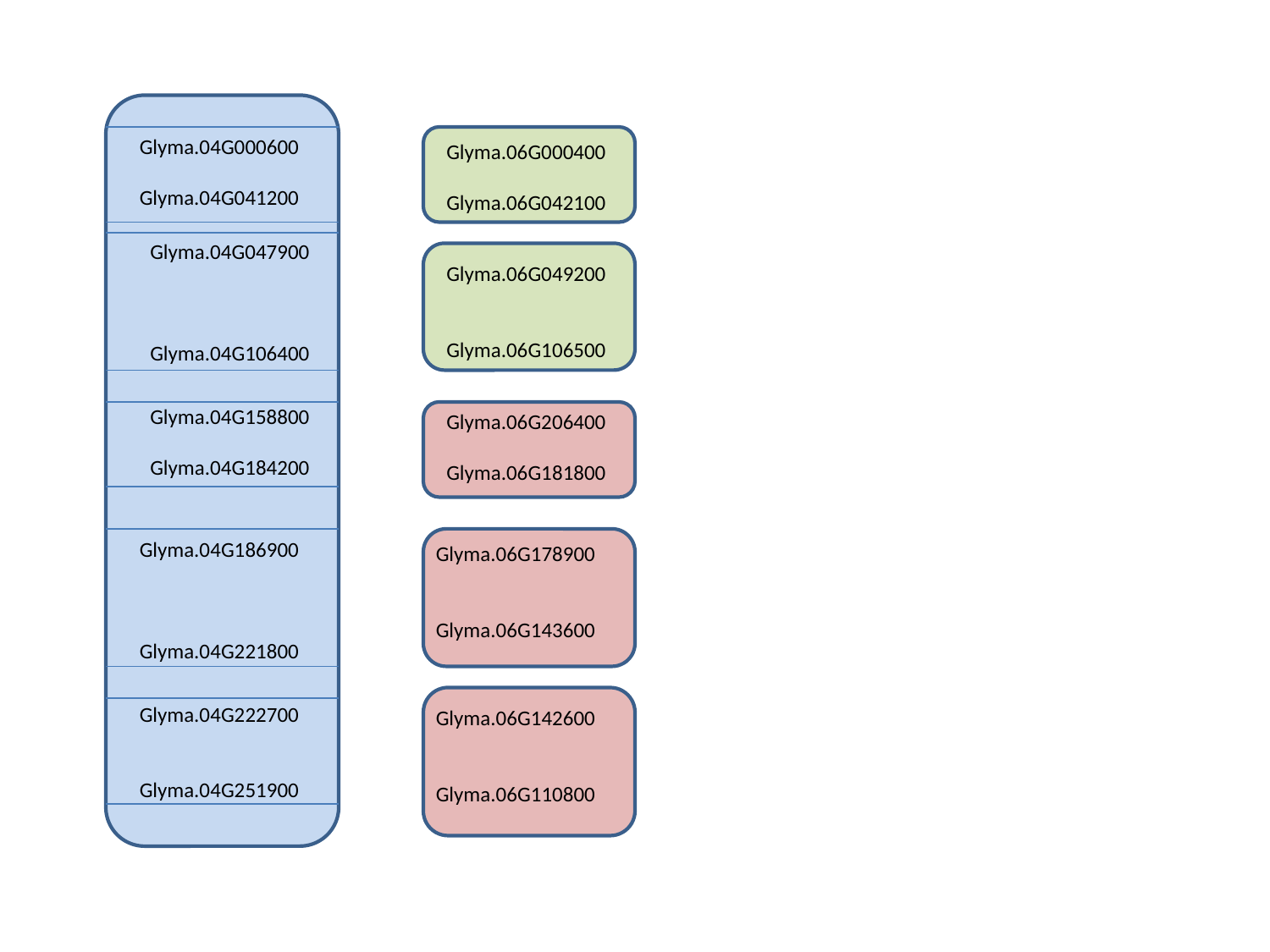

Glyma.04G000600
Glyma.04G041200
Glyma.06G000400
Glyma.06G042100
Glyma.04G047900
Glyma.04G106400
Glyma.06G049200
Glyma.06G106500
Glyma.04G158800
Glyma.04G184200
Glyma.06G206400
Glyma.06G181800
Glyma.04G186900
Glyma.04G221800
Glyma.06G178900
Glyma.06G143600
Glyma.04G222700
Glyma.04G251900
Glyma.06G142600
Glyma.06G110800

### Slide 6
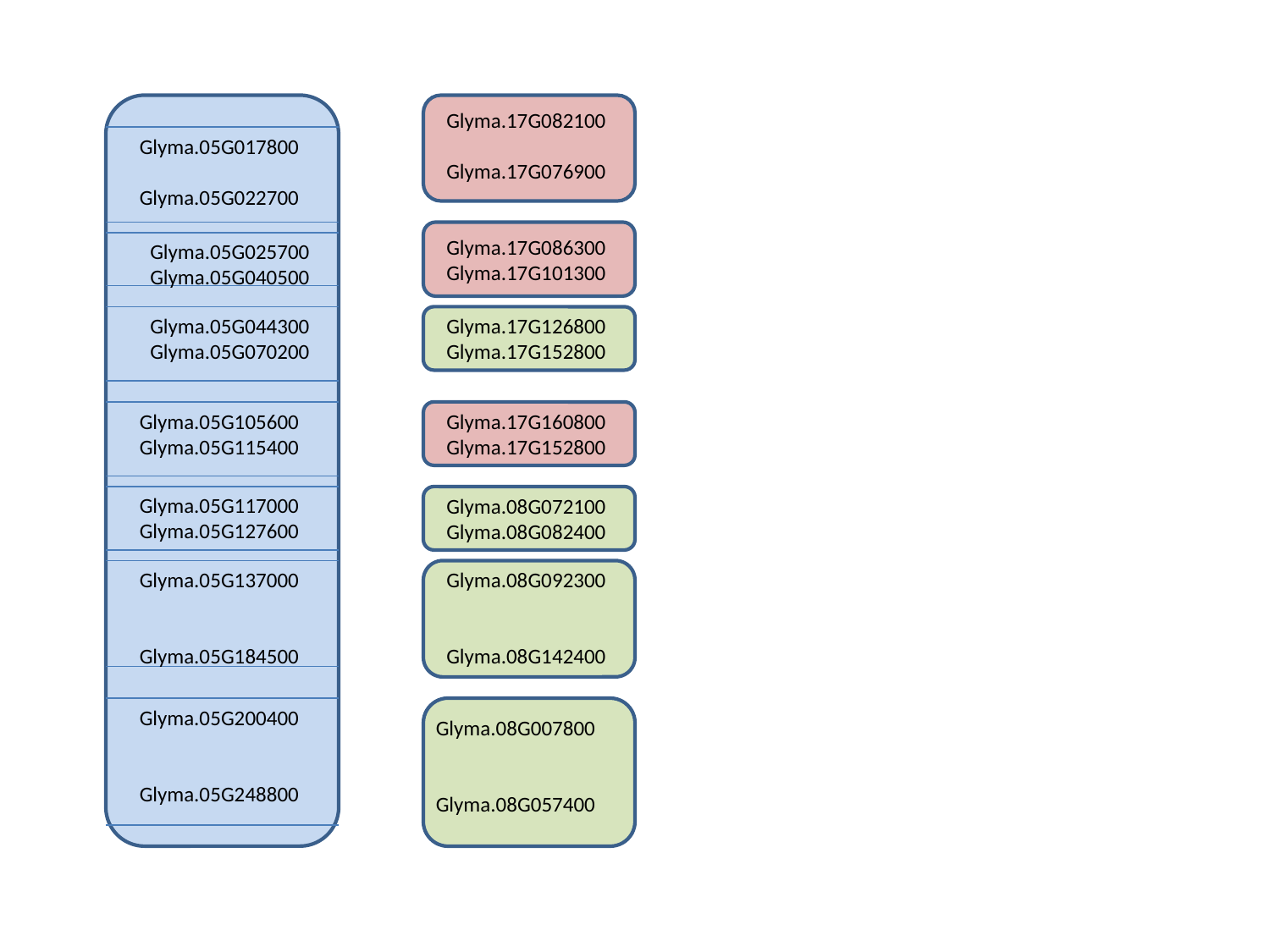

Glyma.17G082100
Glyma.17G076900
Glyma.05G017800
Glyma.05G022700
Glyma.17G086300
Glyma.17G101300
Glyma.05G025700
Glyma.05G040500
Glyma.05G044300
Glyma.05G070200
Glyma.17G126800
Glyma.17G152800
Glyma.05G105600
Glyma.05G115400
Glyma.17G160800
Glyma.17G152800
Glyma.05G117000
Glyma.05G127600
Glyma.08G072100
Glyma.08G082400
Glyma.05G137000
Glyma.05G184500
Glyma.08G092300
Glyma.08G142400
Glyma.05G200400
Glyma.05G248800
Glyma.08G007800
Glyma.08G057400

### Slide 7
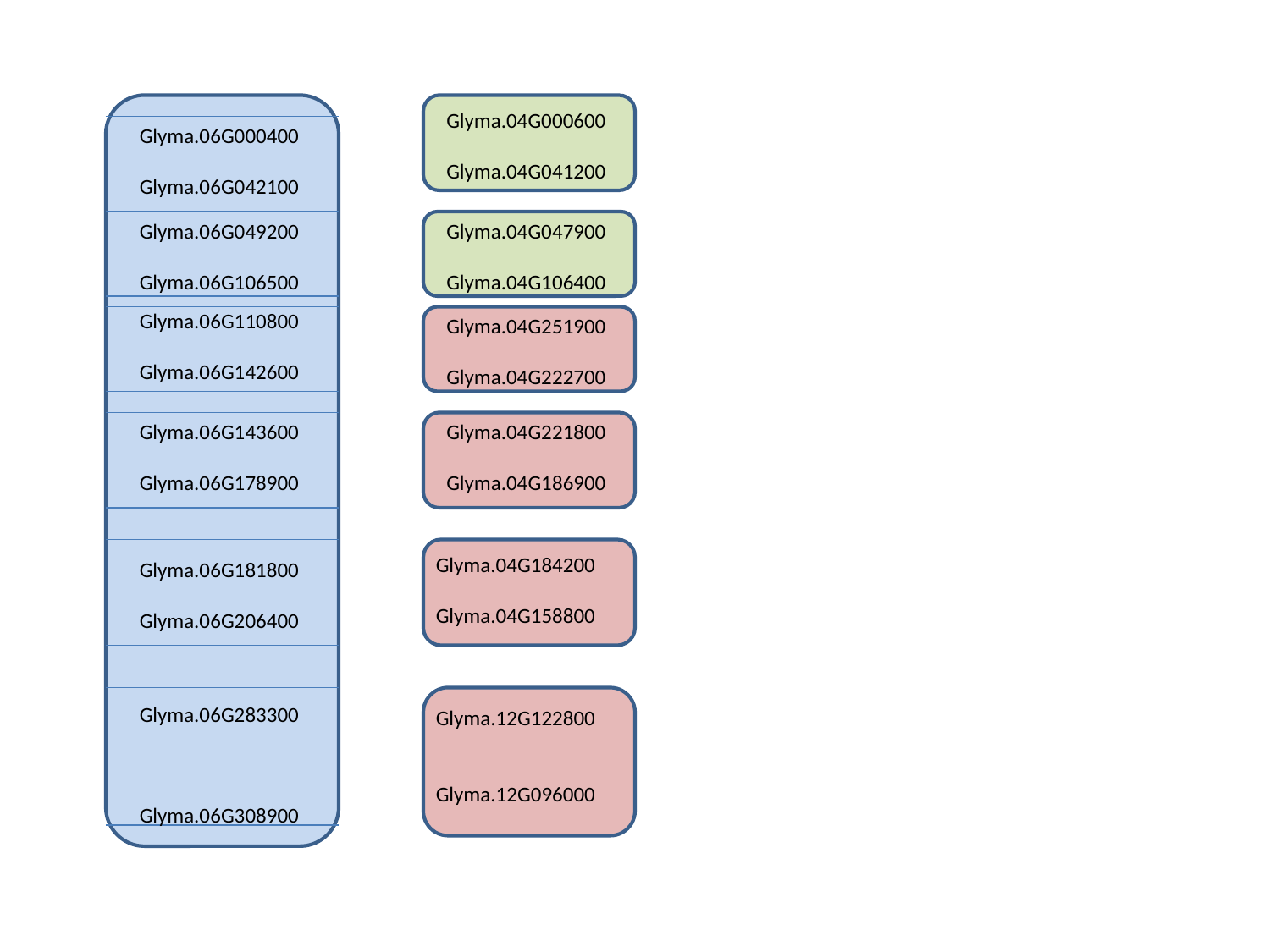

Glyma.04G000600
Glyma.04G041200
Glyma.06G000400
Glyma.06G042100
Glyma.06G049200
Glyma.06G106500
Glyma.04G047900
Glyma.04G106400
Glyma.06G110800
Glyma.06G142600
Glyma.04G251900
Glyma.04G222700
Glyma.06G143600
Glyma.06G178900
Glyma.04G221800
Glyma.04G186900
Glyma.04G184200
Glyma.04G158800
Glyma.06G181800
Glyma.06G206400
Glyma.06G283300
Glyma.06G308900
Glyma.12G122800
Glyma.12G096000

### Slide 8
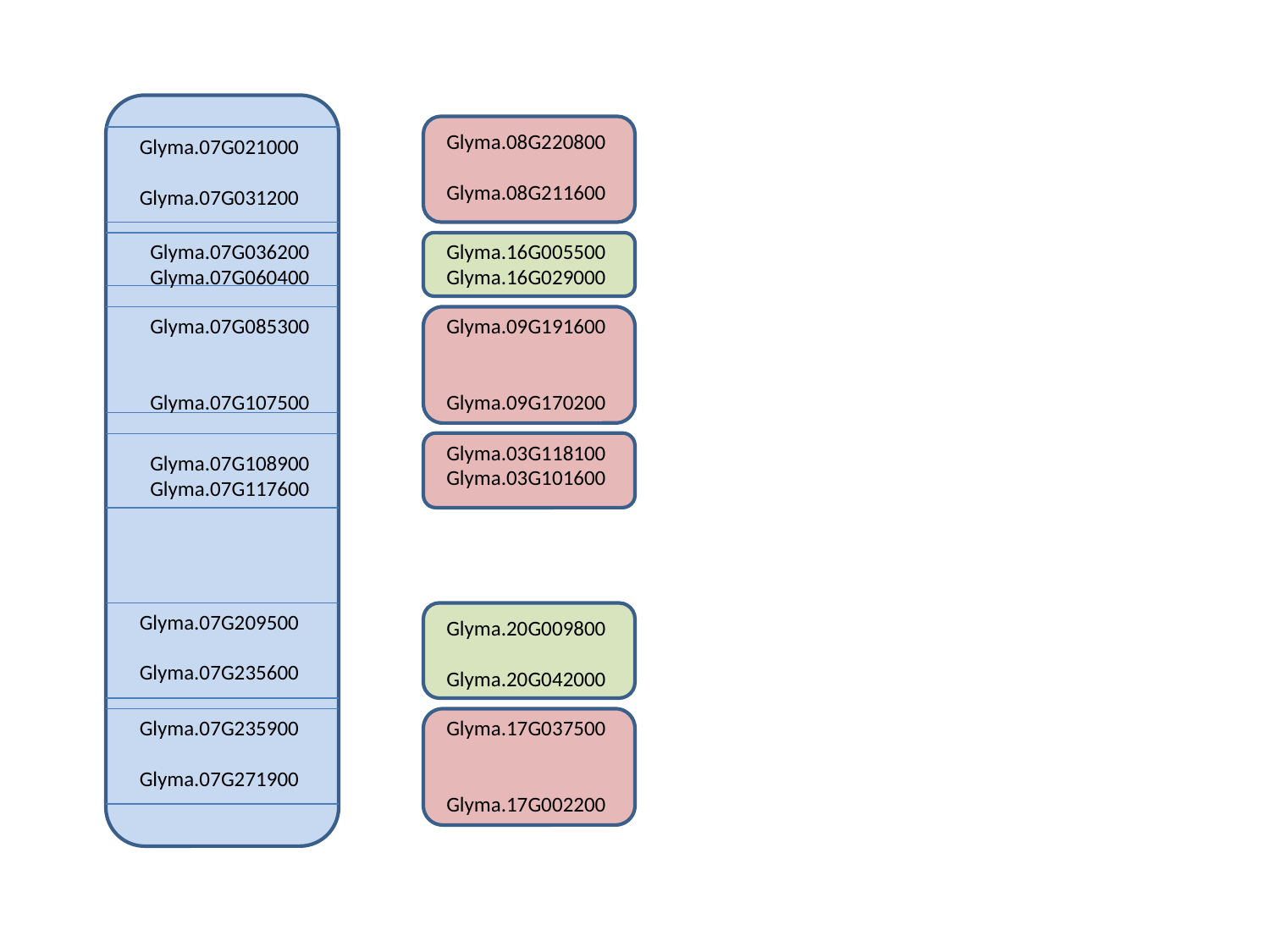

Glyma.08G220800
Glyma.08G211600
Glyma.07G021000
Glyma.07G031200
Glyma.07G036200
Glyma.07G060400
Glyma.16G005500
Glyma.16G029000
Glyma.07G085300
Glyma.07G107500
Glyma.09G191600
Glyma.09G170200
Glyma.03G118100
Glyma.03G101600
Glyma.07G108900
Glyma.07G117600
Glyma.07G209500
Glyma.07G235600
Glyma.20G009800
Glyma.20G042000
Glyma.07G235900
Glyma.07G271900
Glyma.17G037500
Glyma.17G002200

### Slide 9
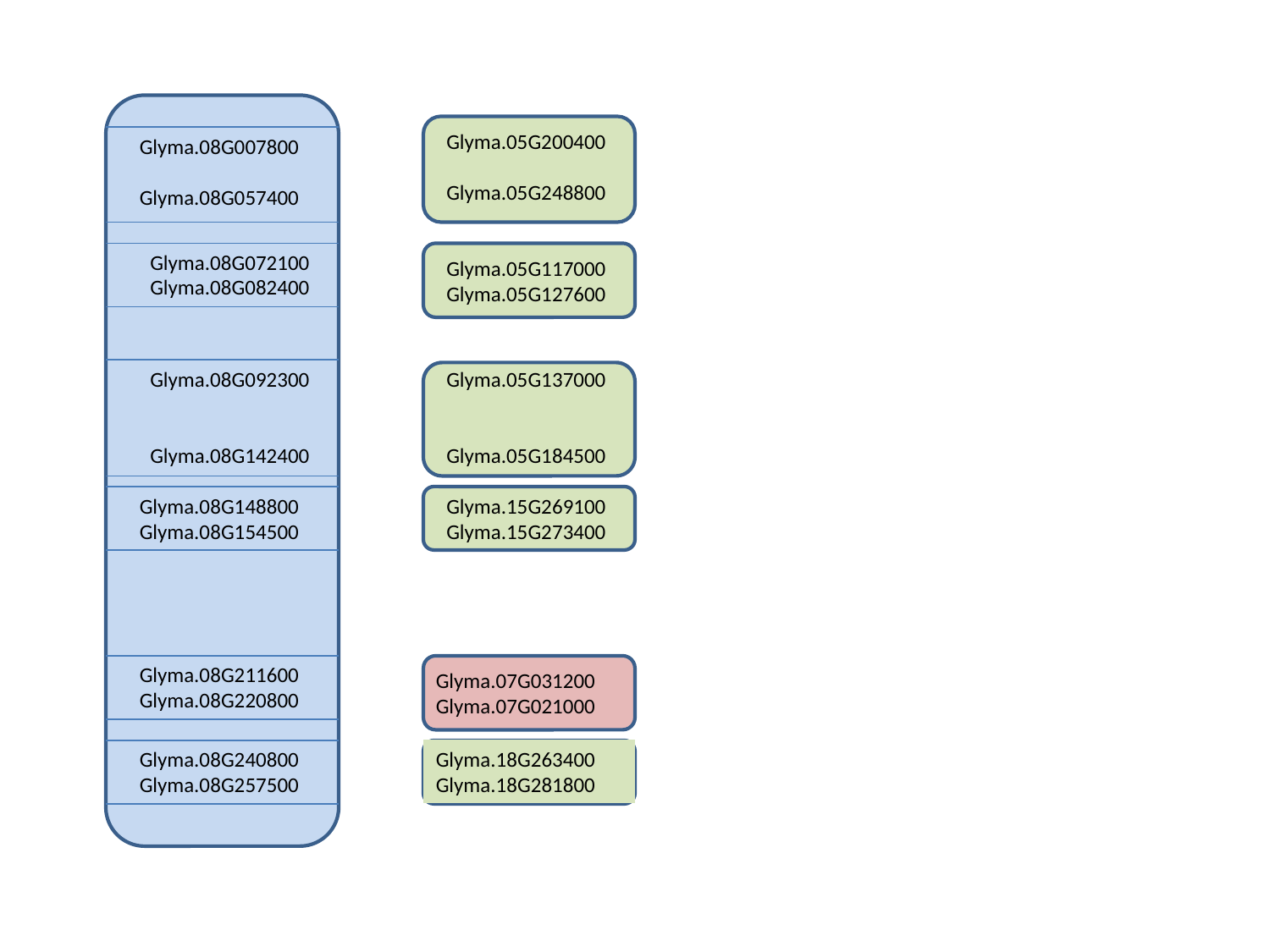

Glyma.05G200400
Glyma.05G248800
Glyma.08G007800
Glyma.08G057400
Glyma.08G072100
Glyma.08G082400
Glyma.05G117000
Glyma.05G127600
Glyma.08G092300
Glyma.08G142400
Glyma.05G137000
Glyma.05G184500
Glyma.08G148800
Glyma.08G154500
Glyma.15G269100
Glyma.15G273400
Glyma.08G211600
Glyma.08G220800
Glyma.07G031200
Glyma.07G021000
Glyma.08G240800
Glyma.08G257500
Glyma.18G263400
Glyma.18G281800

### Slide 10
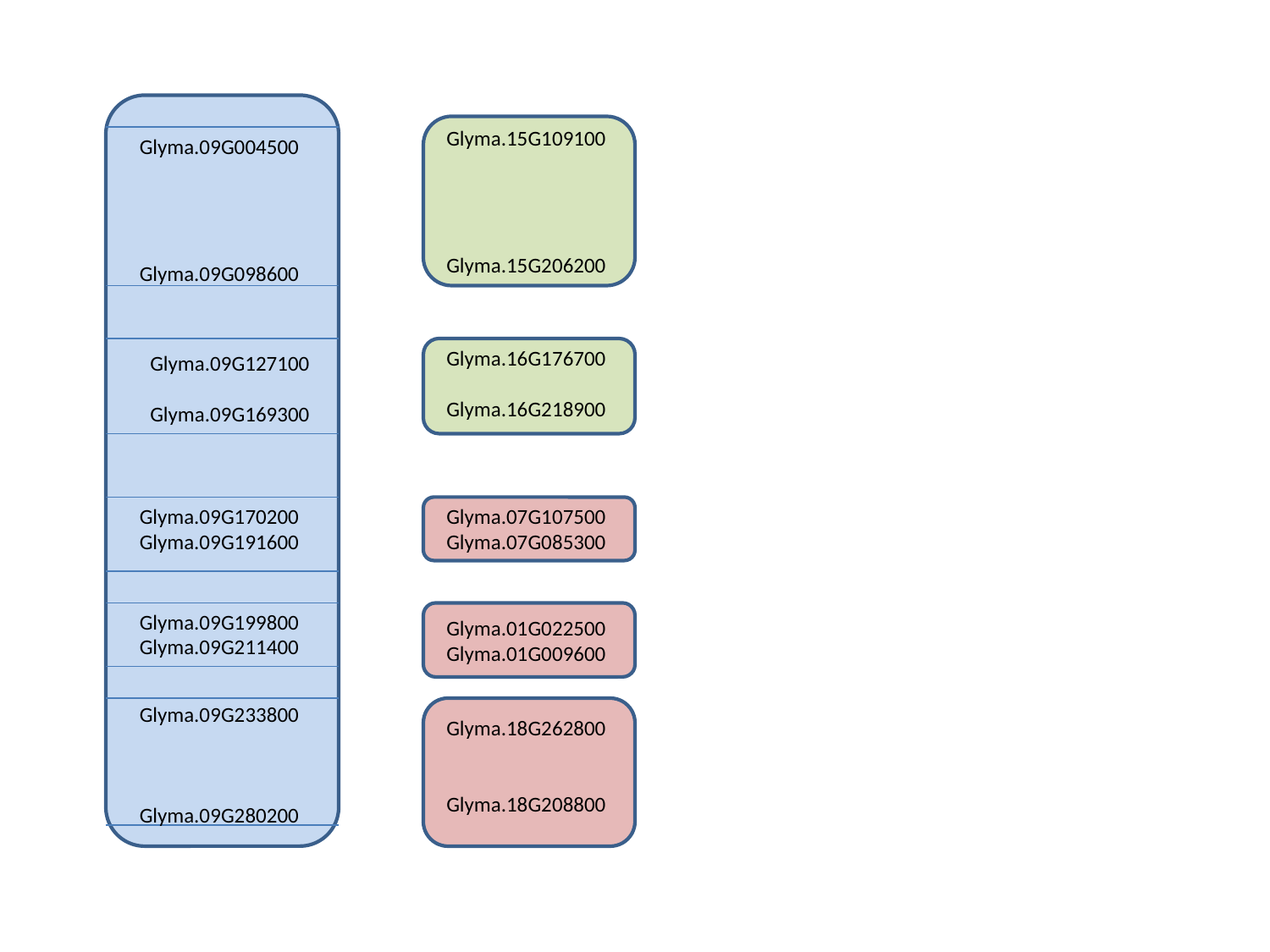

Glyma.15G109100
Glyma.15G206200
Glyma.09G004500
Glyma.09G098600
Glyma.16G176700
Glyma.16G218900
Glyma.09G127100
Glyma.09G169300
Glyma.09G170200
Glyma.09G191600
Glyma.07G107500
Glyma.07G085300
Glyma.09G199800
Glyma.09G211400
Glyma.01G022500
Glyma.01G009600
Glyma.09G233800
Glyma.09G280200
Glyma.18G262800
Glyma.18G208800

### Slide 11
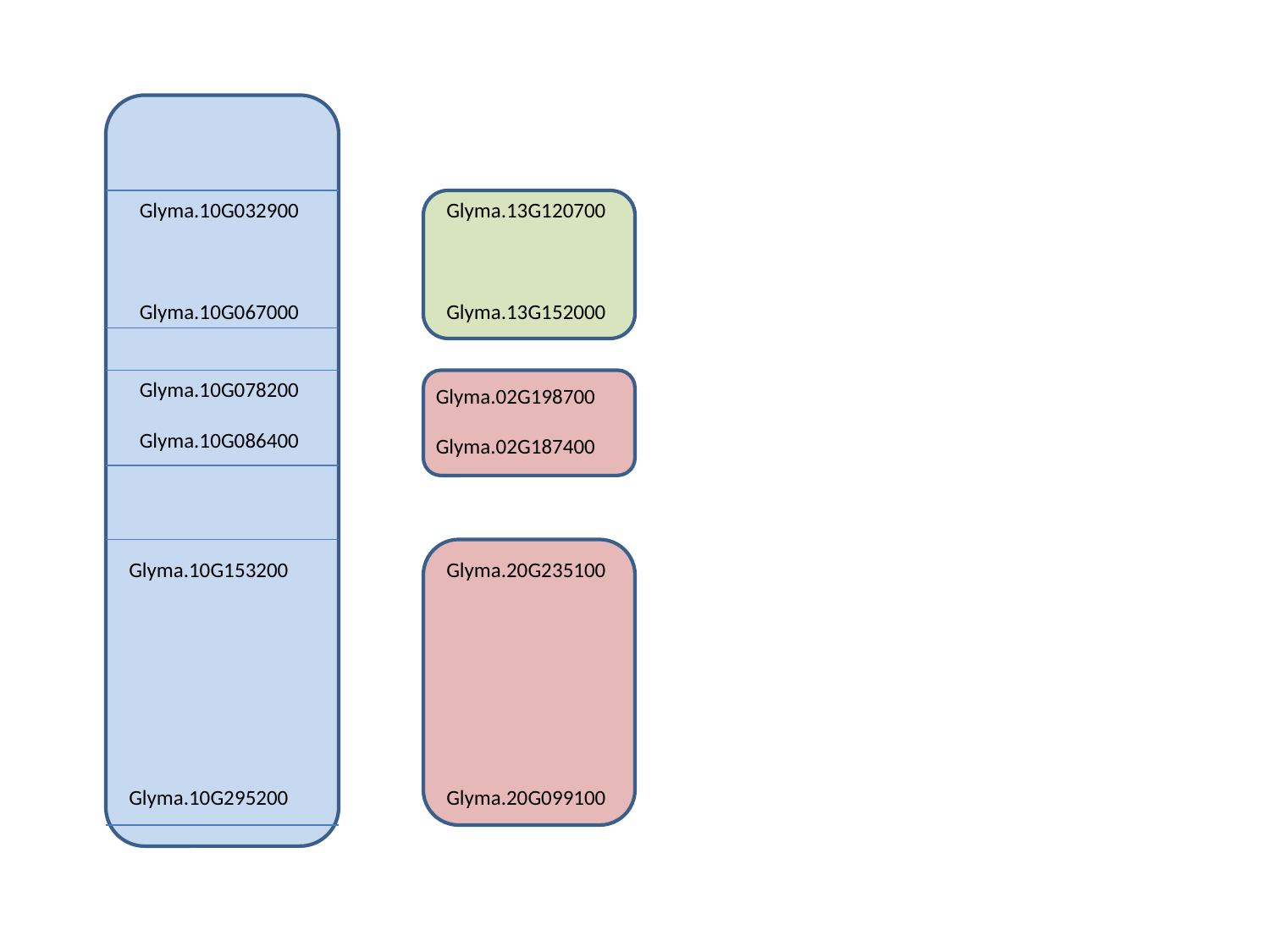

Glyma.10G032900
Glyma.10G067000
Glyma.13G120700
Glyma.13G152000
Glyma.10G078200
Glyma.10G086400
Glyma.02G198700
Glyma.02G187400
Glyma.10G153200
Glyma.10G295200
Glyma.20G235100
Glyma.20G099100

### Slide 12
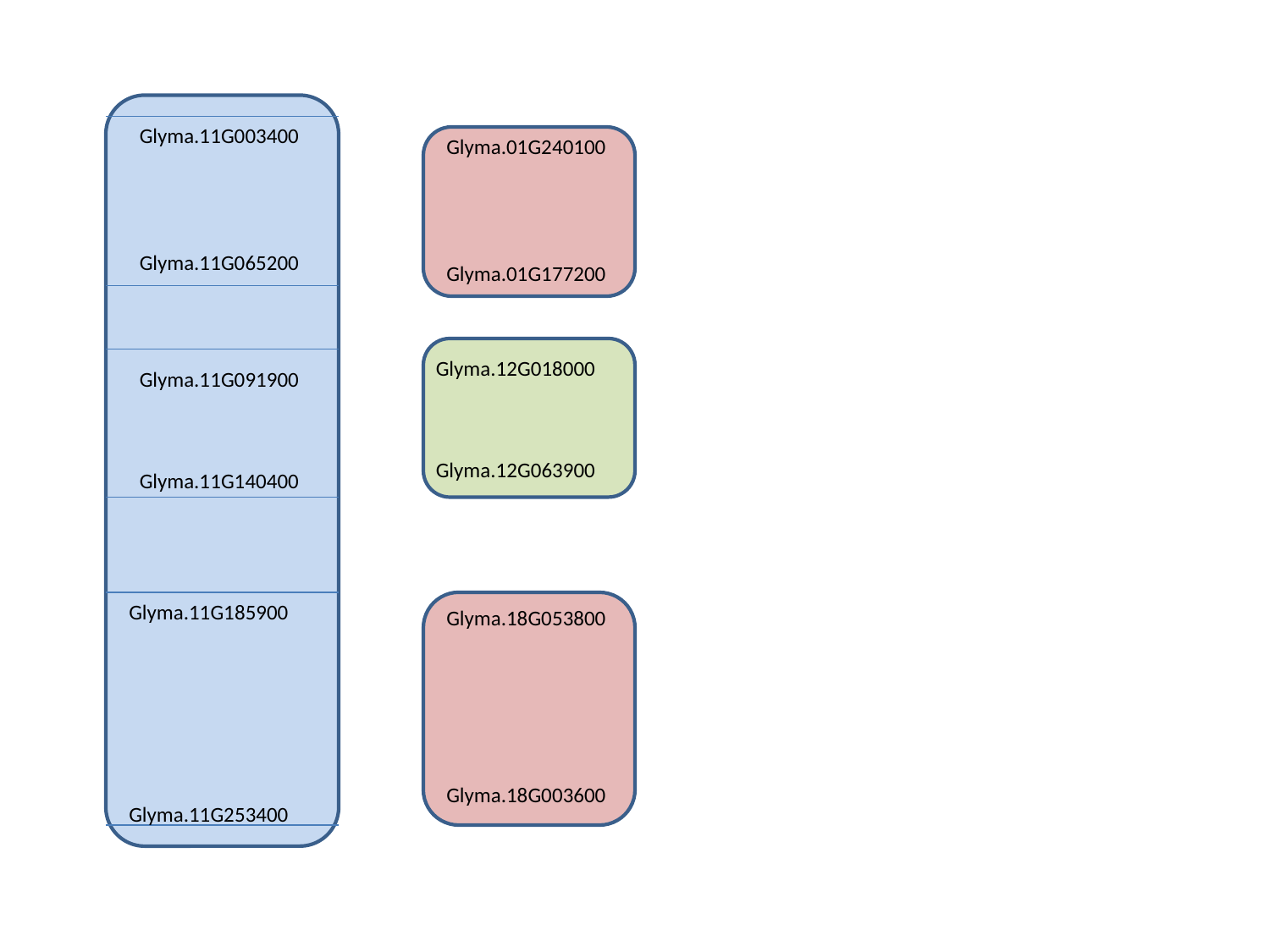

Glyma.11G003400
Glyma.11G065200
Glyma.01G240100
Glyma.01G177200
Glyma.12G018000
Glyma.12G063900
Glyma.11G091900
Glyma.11G140400
Glyma.11G185900
Glyma.11G253400
Glyma.18G053800
Glyma.18G003600

### Slide 13
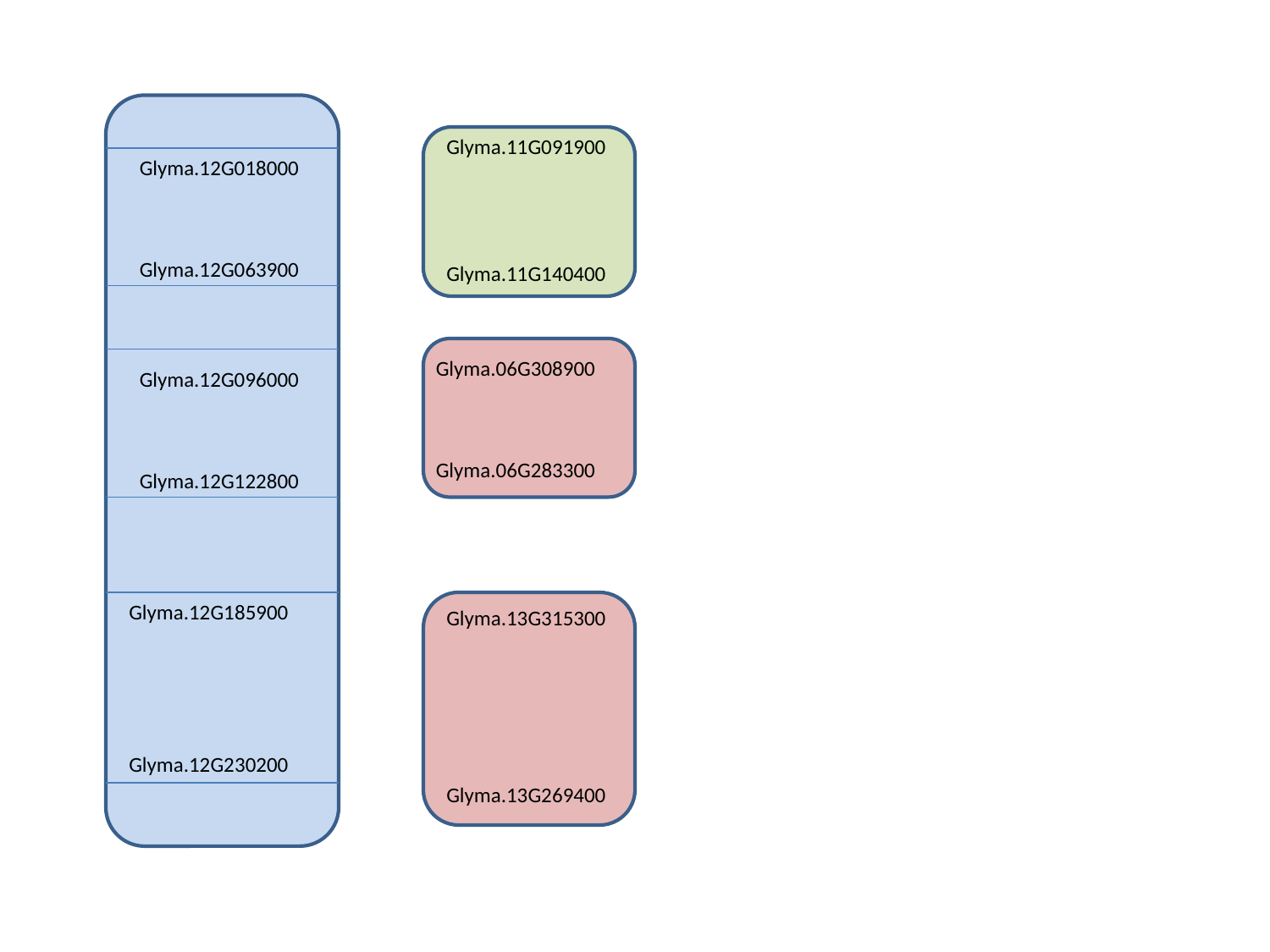

Glyma.11G091900
Glyma.11G140400
Glyma.12G018000
Glyma.12G063900
Glyma.06G308900
Glyma.06G283300
Glyma.12G096000
Glyma.12G122800
Glyma.12G185900
Glyma.12G230200
Glyma.13G315300
Glyma.13G269400

### Slide 14
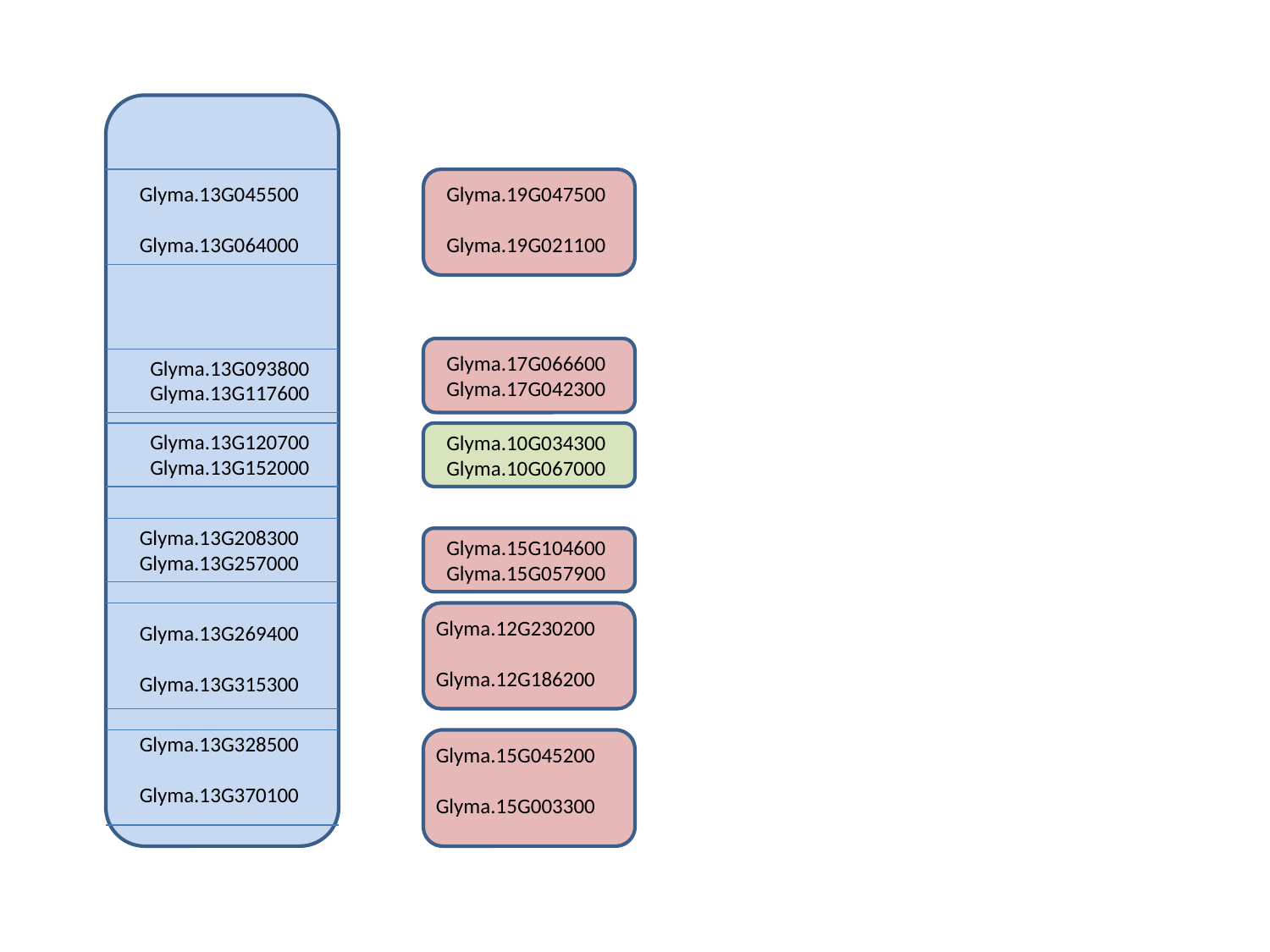

Glyma.13G045500
Glyma.13G064000
Glyma.19G047500
Glyma.19G021100
Glyma.17G066600
Glyma.17G042300
Glyma.13G093800
Glyma.13G117600
Glyma.13G120700
Glyma.13G152000
Glyma.10G034300
Glyma.10G067000
Glyma.13G208300
Glyma.13G257000
Glyma.15G104600
Glyma.15G057900
Glyma.12G230200
Glyma.12G186200
Glyma.13G269400
Glyma.13G315300
Glyma.13G328500
Glyma.13G370100
Glyma.15G045200
Glyma.15G003300

### Slide 15
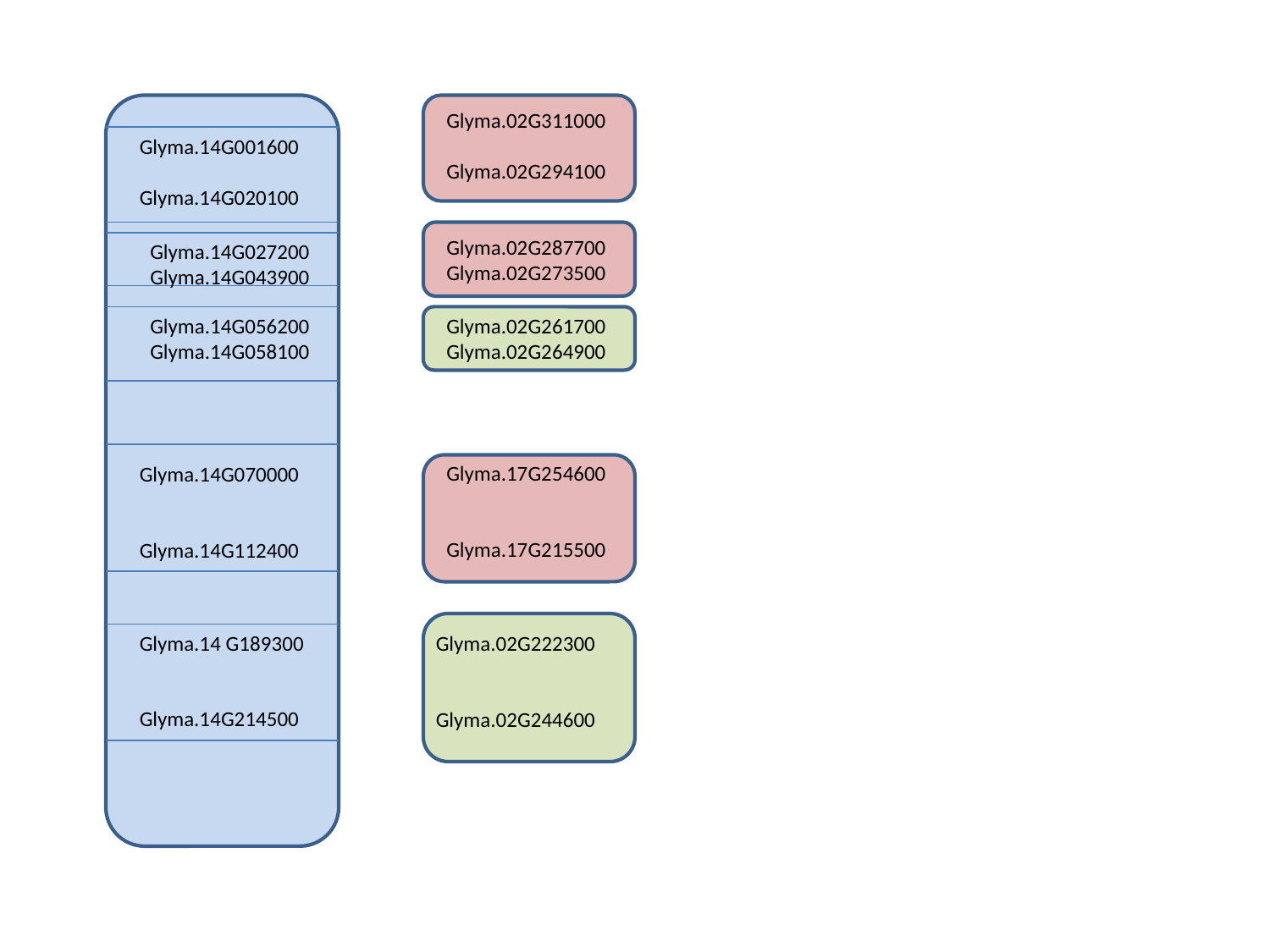

Glyma.02G311000
Glyma.02G294100
Glyma.14G001600
Glyma.14G020100
Glyma.02G287700
Glyma.02G273500
Glyma.14G027200
Glyma.14G043900
Glyma.14G056200
Glyma.14G058100
Glyma.02G261700
Glyma.02G264900
Glyma.17G254600
Glyma.17G215500
Glyma.14G070000
Glyma.14G112400
Glyma.14 G189300
Glyma.14G214500
Glyma.02G222300
Glyma.02G244600

### Slide 16
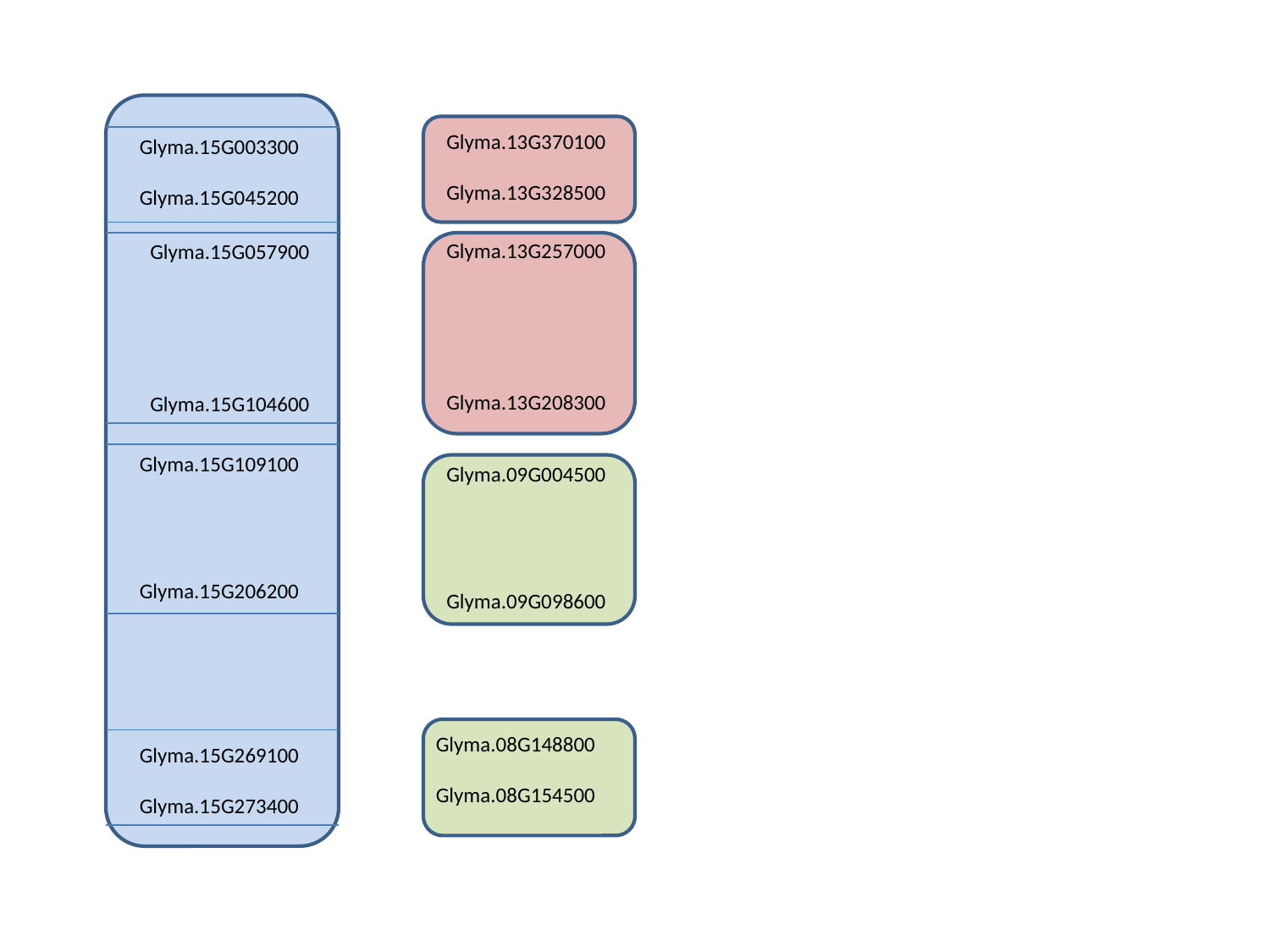

Glyma.13G370100
Glyma.13G328500
Glyma.15G003300
Glyma.15G045200
Glyma.13G257000
Glyma.13G208300
Glyma.15G057900
Glyma.15G104600
Glyma.15G109100
Glyma.15G206200
Glyma.09G004500
Glyma.09G098600
Glyma.08G148800
Glyma.08G154500
Glyma.15G269100
Glyma.15G273400

### Slide 17
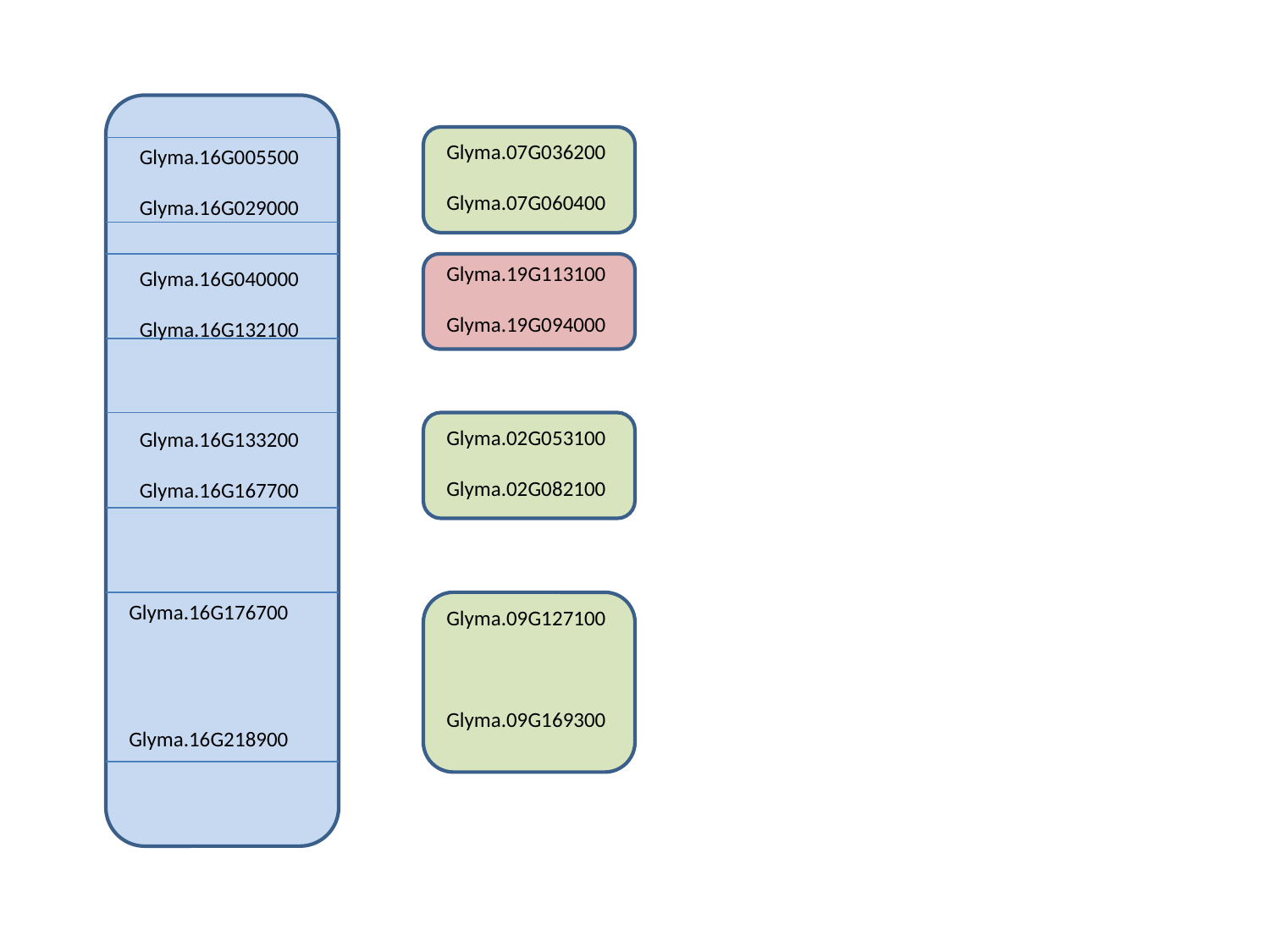

Glyma.07G036200
Glyma.07G060400
Glyma.16G005500
Glyma.16G029000
Glyma.19G113100
Glyma.19G094000
Glyma.16G040000
Glyma.16G132100
Glyma.02G053100
Glyma.02G082100
Glyma.16G133200
Glyma.16G167700
Glyma.16G176700
Glyma.16G218900
Glyma.09G127100
Glyma.09G169300

### Slide 18
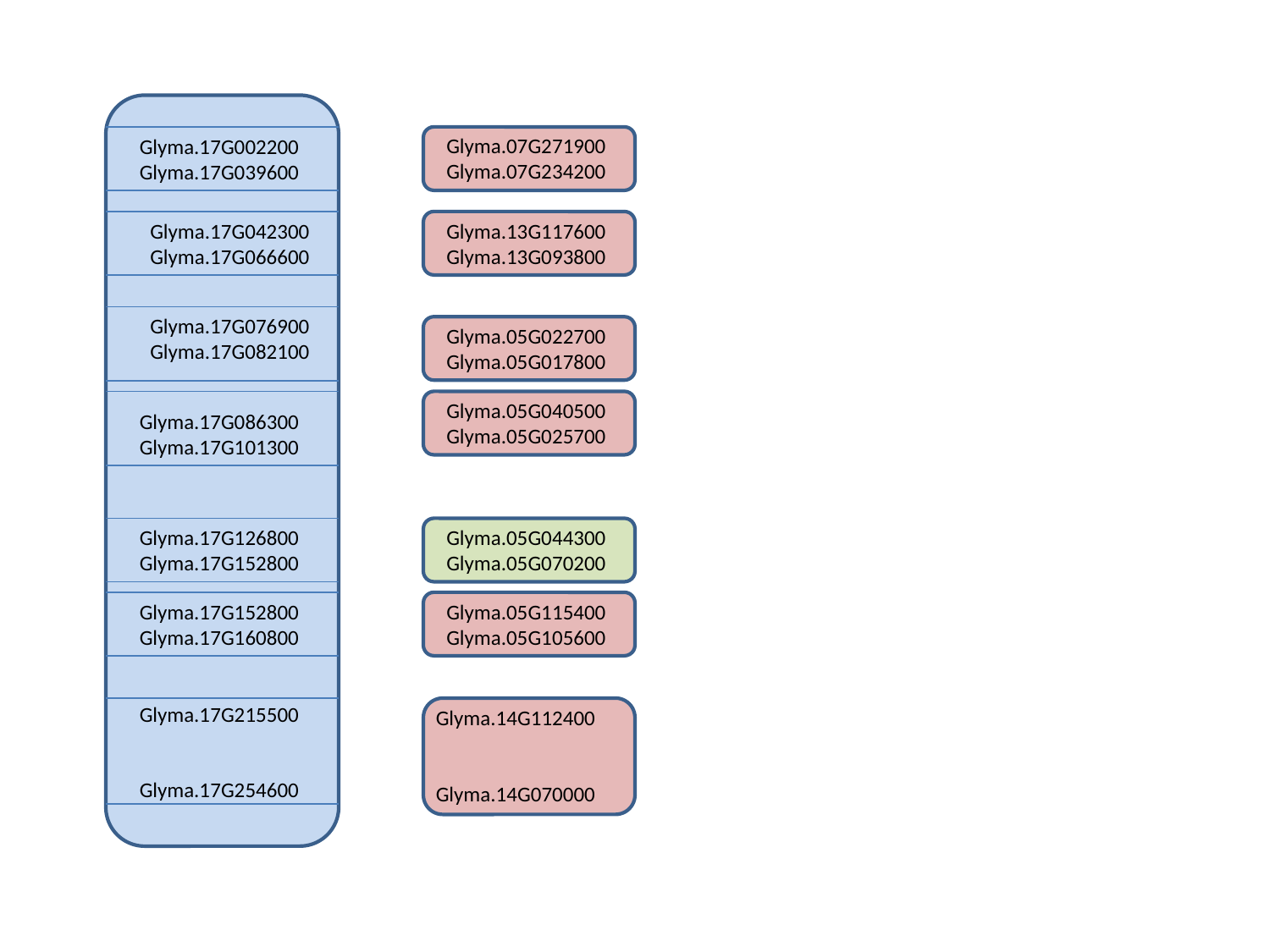

Glyma.07G271900
Glyma.07G234200
Glyma.17G002200
Glyma.17G039600
Glyma.17G042300
Glyma.17G066600
Glyma.13G117600
Glyma.13G093800
Glyma.17G076900
Glyma.17G082100
Glyma.05G022700
Glyma.05G017800
Glyma.05G040500
Glyma.05G025700
Glyma.17G086300
Glyma.17G101300
Glyma.17G126800
Glyma.17G152800
Glyma.05G044300
Glyma.05G070200
Glyma.17G152800
Glyma.17G160800
Glyma.05G115400
Glyma.05G105600
Glyma.17G215500
Glyma.17G254600
Glyma.14G112400
Glyma.14G070000

### Slide 19
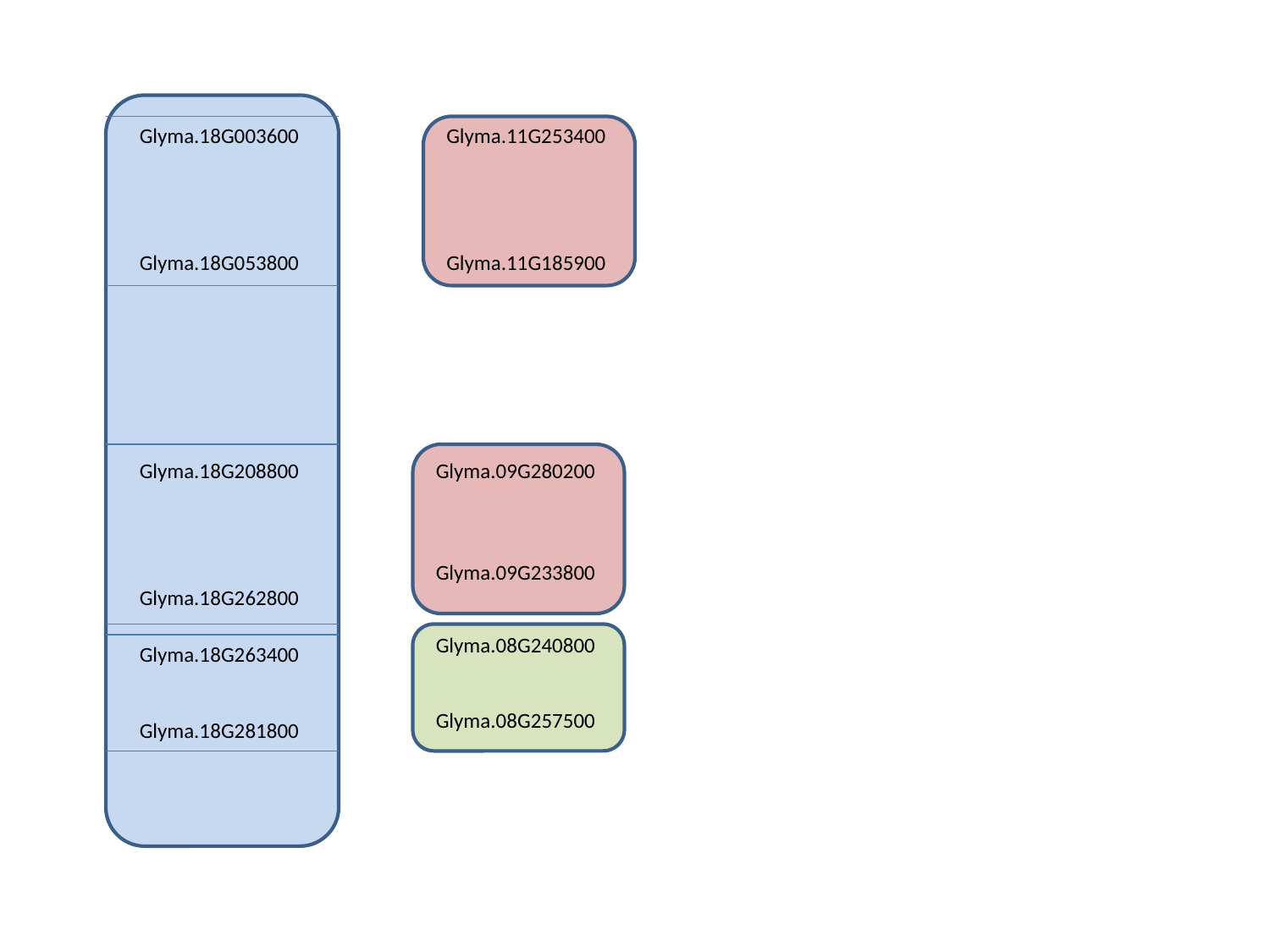

Glyma.18G003600
Glyma.18G053800
Glyma.11G253400
Glyma.11G185900
Glyma.18G208800
Glyma.18G262800
Glyma.09G280200
Glyma.09G233800
Glyma.08G240800
Glyma.08G257500
Glyma.18G263400
Glyma.18G281800

### Slide 20
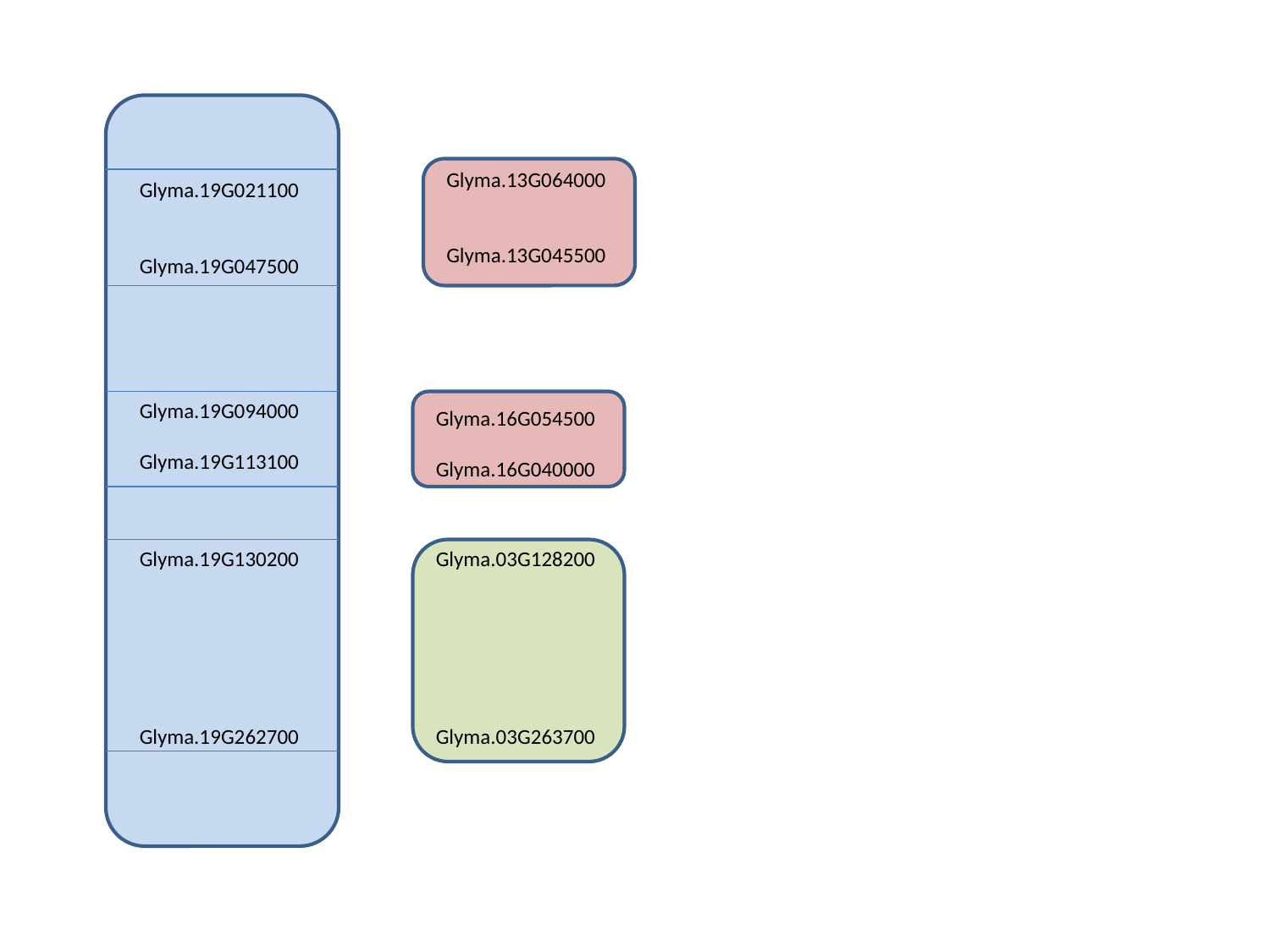

Glyma.13G064000
Glyma.13G045500
Glyma.19G021100
Glyma.19G047500
Glyma.19G094000
Glyma.19G113100
Glyma.16G054500
Glyma.16G040000
Glyma.19G130200
Glyma.19G262700
Glyma.03G128200
Glyma.03G263700

### Slide 21
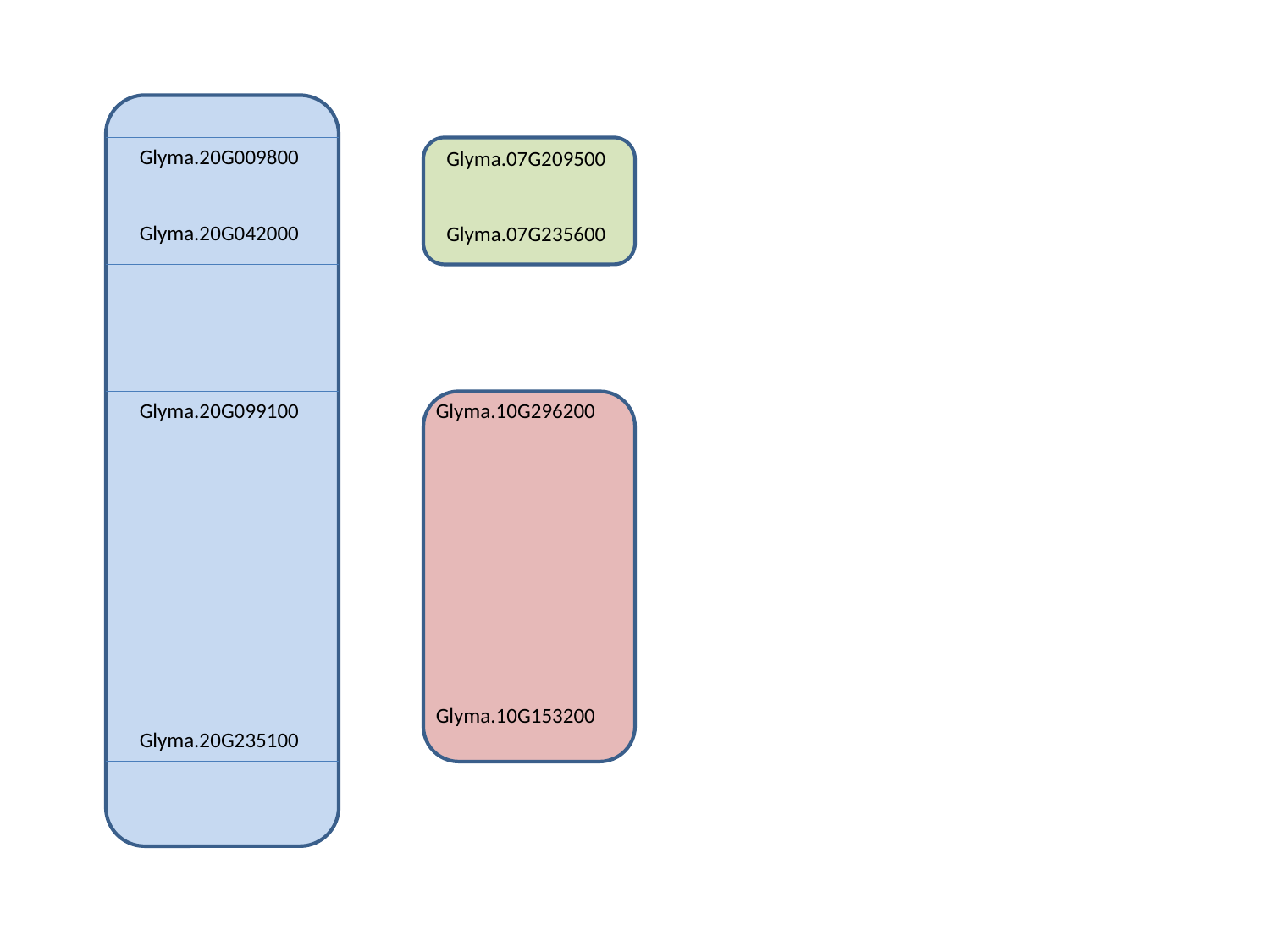

Glyma.20G009800
Glyma.20G042000
Glyma.07G209500
Glyma.07G235600
Glyma.20G099100
Glyma.20G235100
Glyma.10G296200
Glyma.10G153200
