## Supplementary Table 1 for "Molecular evidence for segmental duplication across chromosomes of soybean using transcription factor gene family"

Table S1: Distribution of TF family locus in different chromosome

| **S. No.** | **Gene** | **1** | **2** | **3** | **4** | **5** | **6** | **7** | **8** | **9** | **10** | **11** | **12** | **13** | **14** | **15** | **16** | **17** | **18** | **19** | **20** | **U** | **Total** |
| --- | --- | --- | --- | --- | --- | --- | --- | --- | --- | --- | --- | --- | --- | --- | --- | --- | --- | --- | --- | --- | --- | --- | --- |
| 1. | AP2 | 3 | 4 | 2 | 2 | 2 | 2 | 3 | 4 | 3 | 2 | 3 | 2 | 2 | 1 | 2 | 1 | 4 | 4 | 2 | 0 | 2 | **50** |
| 2. | ARF | 2 | 2 | 3 | 2 | 3 | 1 | 5 | 3 | 1 | 2 | 2 | 5 | 8 | 4 | 3 | 2 | 2 | 2 | 2 | 1 | 1 | **56** |
| 3. | ARR B | 1 | 1 | 0 | 1 | 4 | 1 | 3 | 3 | 2 | 1 | 2 | 0 | 1 | 1 | 2 | 1 | 4 | 1 | 2 | 0 | 0 | **31** |
| 4. | B3 | 2 | 3 | 4 | 4 | 0 | 2 | 4 | 3 | 5 | 4 | 6 | 4 | 2 | 4 | 1 | 3 | 3 | 4 | 4 | 8 | 0 | **70** |
| 5. | BBR- BPC | 0 | 0 | 0 | 1 | 2 | 1 | 2 | 2 | 2 | 0 | 0 | 0 | 0 | 0 | 0 | 0 | 0 | 0 | 0 | 0 | 0 | **10** |
| 6. | BES1 | 2 | 0 | 0 | 1 | 0 | 1 | 1 | 0 | 2 | 0 | 1 | 2 | 2 | 2 | 0 | 0 | 1 | 1 | 0 | 0 | 0 | **16** |
| 7. | bHLH | 20 | 29 | 19 | 15 | 14 | 21 | 23 | **30** | 11 | 20 | 9 | 16 | 25 | 12 | 20 | 10 | 17 | 12 | 14 | 14 | 4 | **355** |
| 8. | BZIP | 6 | 10 | 11 | 10 | 7 | 9 | 2 | 9 | 2 | 7 | **12** | 11 | 11 | 5 | 4 | 5 | 5 | 4 | 11 | 6 | 2 | **149** |
| 9. | C2H2 | 11 | 17 | 16 | 8 | 9 | 10 | 12 | 11 | 3 | **24** | 13 | 16 | 21 | 8 | 6 | 5 | 10 | 8 | 13 | 16 | 2 | **239** |
| 10. | C3H | 2 | 6 | 5 | 3 | 6 | 4 | 2 | 6 | 3 | 7 | 4 | 5 | 4 | 3 | 6 | 2 | 5 | 2 | 2 | 4 | 0 | **81** |
| 11. | CATMA | 0 | 0 | 0 | 0 | 3 | 0 | 1 | 4 | 1 | 0 | 1 | 0 | 0 | 0 | 2 | 0 | 2 | 1 | 0 | 0 | 0 | **15** |
| 12. | CO- like | 0 | 2 | 1 | 1 | 1 | 1 | 1 | 2 | 1 | 2 | 0 | 0 | 3 | 1 | 0 | 1 | 1 | 1 | 3 | 2 | 0 | **24** |
| 13. | CPP | 1 | 0 | 0 | 1 | 1 | 1 | 1 | 1 | 2 | 1 | 1 | 0 | 0 | 0 | 0 | 0 | 1 | 1 | 0 | 1 | 0 | **13** |
| 14. | DBB | 1 | 0 | 1 | 2 | 0 | 2 | 0 | 0 | 1 | 0 | 4 | 3 | 2 | 1 | 1 | 1 | 1 | 0 | 0 | 0 | 0 | **20** |
| 15. | Dof | 3 | 4 | 2 | 5 | 4 | 5 | 6 | 5 | 2 | 2 | 2 | 2 | 11 | 0 | 8 | 2 | 4 | 4 | 5 | 2 | 1 | **79** |
| 16. | E2F/DP | 1 | 1 | 0 | 2 | 1 | 2 | 0 | 0 | 0 | 1 | 3 | 2 | 0 | 0 | 0 | 0 | 1 | 0 | 0 | 0 | 0 | **14** |
| 17. | EIL | 0 | 1 | 0 | 0 | 1 | 1 | 0 | 1 | 0 | 0 | 1 | 0 | 3 | 1 | 1 | 0 | 0 | 1 | 0 | 1 | 0 | **12** |
| 18. | ERF | 14 | 15 | 21 | 14 | 14 | 16 | 15 | 16 | 10 | 23 | 11 | 10 | **27** | 14 | 14 | 11 | 17 | 10 | 15 | 15 | 0 | **302** |
| 19. | FAR1 | 2 | 1 | 3 | 6 | 1 | 5 | 5 | 3 | 5 | 5 | 4 | 3 | 5 | 2 | 10 | 1 | 1 | 7 | 2 | 5 | 0 | **76** |
| 20. | G2 LIKE | 7 | **9** | 8 | 2 | 5 | 3 | 6 | 2 | 9 | 5 | 4 | 7 | 5 | 4 | 6 | 1 | 4 | 4 | **9** | 6 | 0 | **106** |
| 21. | GATA | 3 | 5 | 2 | 5 | 3 | 4 | 4 | 6 | 2 | 2 | 5 | 3 | 2 | 3 | 2 | 4 | 5 | 0 | 2 | 1 | 0 | **63** |
| 22. | GeBP | 0 | 0 | 1 | 0 | 1 | 0 | 0 | 0 | 0 | 2 | 0 | 0 | 1 | 0 | 1 | 0 | 0 | 0 | 1 | 2 | 0 | **9** |
| 23. | GRAS | 5 | 5 | 4 | 4 | 5 | 4 | 5 | 4 | 6 | 5 | 16 | 11 | 11 | 3 | 9 | 5 | 5 | 5 | 2 | 3 | 1 | **118** |
| 24. | GRF | 2 | 0 | 1 | 1 | 0 | 1 | 1 | 0 | 2 | 1 | 3 | 1 | 1 | 0 | 1 | 1 | 3 | 0 | 1 | 0 | 2 | **22** |
| 25. | HB-other | 0 | 2 | 1 | 3 | 0 | 3 | 2 | 1 | 0 | 0 | 0 | 1 | 2 | 1 | 0 | 0 | 0 | 1 | 1 | 0 | 0 | **18** |
| 26. | HB-PHD | 0 | 0 | 0 | 0 | 0 | 0 | 0 | 0 | 1 | 1 | 0 | 0 | 1 | 0 | 1 | 0 | 1 | 0 | 0 | 1 | 0 | **6** |
| 27. | HD-ZIP | 7 | 4 | 4 | 3 | 6 | 5 | 7 | 10 | 11 | 3 | 6 | 4 | 6 | 1 | 5 | 3 | 4 | 6 | 4 | 3 | 1 | **103** |
| 28. | HRT- like | 0 | 0 | 0 | 0 | 0 | 0 | 1 | 0 | 0 | 0 | 0 | 0 | 0 | 0 | 0 | 0 | 0 | 0 | 0 | 0 | 0 | **1** |
| 29. | HSF | 5 | 1 | 3 | 2 | 4 | 1 | 2 | 3 | 3 | 5 | 3 | 0 | 4 | 3 | 1 | 2 | 4 | 0 | 3 | 3 | 0 | **52** |
| 30. | LBD | 5 | 6 | 4 | 4 | 5 | 4 | 3 | 7 | 2 | 5 | 4 | 2 | 6 | 6 | 4 | 2 | 4 | 5 | 7 | 4 | 2 | **91** |
| 31. | LFY | 0 | 0 | 0 | 1 | 0 | 1 | 0 | 0 | 0 | 0 | 0 | 0 | 0 | 0 | 0 | 0 | 0 | 0 | 0 | 0 | 0 | **2** |
| 32. | LSD | 1 | 0 | 0 | 0 | 2 | 0 | 1 | 1 | 1 | 0 | 0 | 0 | 0 | 0 | 1 | 0 | 1 | 0 | 0 | 0 | 0 | **8** |
| 33. | M TYPE MADS | 1 | 7 | 3 | 2 | 4 | 0 | 6 | 7 | 0 | 16 | 8 | 0 | 2 | 1 | 0 | 1 | 2 | 10 | 2 | 7 | 4 | **83** |
| 34. | MIKC MADS | 6 | 6 | 3 | 5 | 6 | 5 | 3 | **10** | 6 | 2 | 2 | 2 | 8 | 3 | 4 | 3 | 3 | 6 | 2 | 3 | 1 | **89** |
| 35. | MYB | 15 | 17 | 16 | 15 | 14 | 20 | 20 | 15 | 16 | 20 | 14 | 19 | **22** | 13 | 10 | 9 | 15 | 20 | **22** | 16 | 3 | **331** |
| 36. | MYB RELATED | 10 | 8 | 8 | 11 | 6 | 11 | 8 | 9 | 8 | 7 | 11 | 8 | **18** | 7 | 7 | 10 | 9 | 12 | 9 | 9 | 1 | **187** |
| 37. | NAC | 5 | 9 | 3 | 11 | 13 | 14 | 10 | 13 | 5 | 8 | 5 | **17** | 13 | 6 | 7 | 12 | 6 | 5 | 12 | 6 | 0 | **180** |
| 38. | NF-X1 | 0 | 0 | 0 | 0 | 0 | 0 | 1 | 1 | 1 | 0 | 1 | 0 | 0 | 0 | 0 | 0 | 0 | 1 | 0 | 0 | 0 | **5** |
| 39. | NF-YA | 0 | 2 | 1 | 0 | 1 | 0 | 1 | 2 | 2 | 1 | 0 | 1 | 2 | 1 | 3 | 1 | 1 | 1 | 1 | 0 | 0 | **21** |
| 40. | NF-FB | 0 | 3 | 3 | 2 | 4 | 1 | 3 | 3 | 3 | 4 | 1 | 1 | 1 | 0 | 2 | 0 | 3 | 1 | 1 | 3 | 1 | **40** |
| 41. | NF-YC | 0 | 2 | 1 | 1 | 0 | 3 | 0 | 2 | 0 | 1 | 2 | 2 | 5 | 1 | 2 | 0 | 0 | 1 | 1 | 1 | 0 | **25** |
| 42. | Nin-like | 1 | 1 | 0 | 5 | 1 | 7 | 0 | 0 | 1 | 1 | 2 | 1 | 1 | 1 | 1 | 1 | 1 | 0 | 0 | 3 | 0 | **28** |
| 43. | RAV | 1 | 1 | 0 | 0 | 0 | 0 | 0 | 0 | 0 | 1 | 0 | 0 | 0 | 0 | 0 | 0 | 0 | 0 | 0 | 2 | 0 | **5** |
| 44. | S1Fa like | 0 | 0 | 1 | 0 | 0 | 0 | 0 | 0 | 0 | 1 | 0 | 0 | 1 | 0 | 0 | 0 | 0 | 0 | 1 | 0 | 0 | **4** |
| 45. | SAP | 0 | 1 | 0 | 0 | 0 | 0 | 0 | 0 | 0 | 0 | 0 | 0 | 0 | 1 | 0 | 0 | 0 | 0 | 0 | 0 | 0 | **2** |
| 46. | SBP | 3 | 3 | 3 | 2 | 2 | 3 | 5 | 1 | 2 | 1 | 2 | 1 | 5 | 0 | 3 | 2 | 2 | 1 | 2 | 1 | 2 | **46** |
| 47. | SRS | 1 | 2 | 0 | 3 | 0 | 2 | 1 | 0 | 0 | 0 | 3 | 1 | 1 | 2 | 1 | 1 | 2 | 0 | 0 | 1 | 0 | **21** |
| 48. | STAT | 0 | 0 | 0 | 0 | 0 | 0 | 0 | 0 | 0 | 0 | 0 | 0 | 0 | 0 | 1 | 0 | 0 | 0 | 0 | 0 | 0 | **1** |
| 49. | TALE | 5 | 3 | 3 | 7 | 2 | 5 | 1 | 2 | 2 | 2 | 4 | 4 | 3 | 4 | 2 | 1 | 6 | 2 | 2 | 1 | 2 | **63** |
| 50. | TCP | 1 | 1 | 1 | 3 | 5 | 5 | 2 | 4 | 2 | 4 | 1 | 5 | 5 | 0 | 1 | 2 | 4 | 3 | 3 | 4 | 0 | **56** |
| 51. | Trihelix | 2 | 2 | 5 | 3 | 1 | 2 | 3 | 3 | 5 | 10 | 2 | 1 | 7 | 0 | 5 | 6 | 1 | 3 | 3 | 8 | 0 | **72** |
| 52. | VOZ | 0 | 0 | 0 | 0 | 0 | 1 | 1 | 0 | 0 | 1 | 1 | 1 | 1 | 0 | 0 | 0 | 0 | 0 | 0 | 0 | 0 | **6** |
| 53. | Whirly | 1 | 2 | 1 | 0 | 0 | 0 | 0 | 1 | 0 | 0 | 0 | 0 | 0 | 0 | 0 | 0 | 0 | 1 | 1 | 0 | 0 | **7** |
| 54. | WOX | 1 | 2 | 1 | 2 | 1 | 2 | 4 | 1 | 1 | 2 | 4 | 1 | 2 | 1 | 1 | 0 | 1 | 3 | 1 | 2 | 0 | **33** |
| 55. | WRKY | 7 | 12 | 9 | 11 | 12 | **15** | 8 | 13 | 14 | 9 | 3 | 3 | 9 | 13 | 6 | 7 | 12 | 12 | 7 | 3 | 0 | **185** |
| 56. | YABBY | 3 | 1 | 1 | 1 | 1 | 2 | 0 | 1 | 0 | 0 | 0 | 2 | 2 | 0 | 0 | 0 | 2 | 1 | 0 | 0 | 0 | **17** |
| 57. | ZFHD | 3 | 6 | 0 | 2 | 2 | 2 | 3 | 6 | 5 | 0 | 1 | 3 | 2 | 1 | 0 | 2 | 2 | 5 | 1 | 7 | 0 | **53** |
|  |  | **172** | **219** | **179** | **187** | **176** | **210** | **197** | **234** | **162** | **221** | **185** | **182** | **275** | **136** | **168** | **120** | **183** | **161** | **176** | **174** | **31** | **3771** |
