## Supplementary Table 2 for "Molecular evidence for segmental duplication across chromosomes of soybean using transcription factor gene family"

Table S2: Distribution of TF family locus in different chromosome

| **TF gene family** | **Total genes** | **Loci** | **Loci %** |
| --- | --- | --- | --- |
| AP2 | 99 | 50 | 50.51 |
| ARF | 107 | 56 | 52.34 |
| ARR B | 55 | 31 | 56.36 |
| B3 | 148 | 70 | 47.30 |
| BBR BPC | 29 | 10 | 34.48 |
| BES1 | 19 | 16 | 84.21 |
| BHLH | 548 | 355 | 64.78 |
| BZIP | 352 | 149 | 42.33 |
| C2H2 | 321 | 239 | 74.45 |
| C3H | 150 | 81 | 54.00 |
| CATMA | 24 | 15 | 62.50 |
| CO LIKE | 40 | 24 | 60.00 |
| CPP | 26 | 13 | 50.00 |
| DBB | 47 | 20 | 42.55 |
| DoF | 97 | 79 | 81.44 |
| E2F/DP | 38 | 14 | 36.84 |
| eil | 13 | 12 | 92.31 |
| ERF | 338 | 302 | 89.35 |
| FAR1 | 138 | 76 | 55.07 |
| G2 LIKE | 222 | 106 | 47.75 |
| GATA | 92 | 63 | 68.48 |
| gebp | 10 | 9 | 90.00 |
| GRAS | 151 | 118 | 78.15 |
| GRF | 42 | 22 | 52.38 |
| HB OTHERS | 30 | 18 | 60.00 |
| HBPHD | 16 | 6 | 37.50 |
| HDZIP | 180 | 103 | 57.22 |
| HRT LIKE | 1 | 1 | 100.00 |
| HSF | 81 | 52 | 64.20 |
| LBD | 125 | 91 | 72.80 |
| LFY | 2 | 2 | 100.00 |
| LSD | 18 | 8 | 44.44 |
| M TYPE MADS | 87 | 83 | 95.40 |
| MICK MADS | 209 | 89 | 42.58 |
| MYB | 430 | 331 | 76.98 |
| MYB RELATED | 342 | 187 | 54.68 |
| NAC | 269 | 180 | 66.91 |
| NFX1 | 10 | 5 | 50.00 |
| nfya | 117 | 21 | 17.95 |
| nfyb | 58 | 40 | 68.97 |
| nfyc | 38 | 25 | 65.79 |
| NIN LIKE | 61 | 28 | 45.90 |
| RAV | 5 | 5 | 100.00 |
| s1fa like | 4 | 4 | 100.00 |
| sap | 2 | 2 | 100.00 |
| SBP | 111 | 46 | 41.44 |
| SRS | 42 | 21 | 50.00 |
| STAT | 2 | 1 | 50.00 |
| TALE | 133 | 63 | 47.37 |
| TCP | 81 | 56 | 69.14 |
| Trihelix | 104 | 72 | 69.23 |
| VOZ | 26 | 6 | 23.08 |
| Whirly | 18 | 7 | 38.89 |
| WOX | 41 | 33 | 80.49 |
| WRKY | 296 | 185 | 62.50 |
| YABBY | 47 | 17 | 36.17 |
| ZFHD | 58 | 53 | 91.38 |
| **Total** | **6150** | **3771** | **61.32** |
