## Supplementary Table 3 for "Molecular evidence for segmental duplication across chromosomes of soybean using transcription factor gene family"

Table S3: Distribution and order of TF Locus in different chromosomes of soybean

| 1. | Glyma.01G000600.1.p | Glyma.01G000600 | FAR1 |
| --- | --- | --- | --- |
| 2. | Glyma.01G002100.1.p | Glyma.01G002100 | ARF |
| 3. | Glyma.01G003000.1.p | Glyma.01G003000 | MYB_related |
| 4. | Glyma.01G004900.1.p | Glyma.01G004900 | MYB |
| 5. | Glyma.01G005000.1.p | Glyma.01G005000 | C3H |
| 6. | Glyma.01G005500.1.p | Glyma.01G005500 | NAC |
| 7. | Glyma.01G009600.1.p | Glyma.01G009600 | G2-like |
| 8. | Glyma.01G013600.1.p | Glyma.01G013600 | bZIP |
| 9. | Glyma.01G014600.1.p | Glyma.01G014600 | HD-ZIP |
| 10. | Glyma.01G014800.1.p | Glyma.01G014800 | G2-like |
| 11. | Glyma.01G015900.1.p | Glyma.01G015900 | HSF |
| 12. | Glyma.01G016600.1.p | Glyma.01G016600 | MYB |
| 13. | Glyma.01G018400.1.p | Glyma.01G018400 | bHLH |
| 14. | Glyma.01G019700.1.p | Glyma.01G019700 | bHLH |
| 15. | Glyma.01G020500.1.p | Glyma.01G020500 | MIKC_MADS |
| 16. | Glyma.01G021100.1.p | Glyma.01G021100 | Dof |
| 17. | Glyma.01G022500.1.p | Glyma.01G022500 | AP2 |
| 18. | Glyma.01G023500.1.p | Glyma.01G023500 | MIKC_MADS |
| 19. | Glyma.01G023800.1.p | Glyma.01G023800 | bHLH |
| 20. | Glyma.01G025000.1.p | Glyma.01G025000 | ZF-HD |
| 21. | Glyma.01G025400.1.p | Glyma.01G025400 | ERF |
| 22. | Glyma.01G028300.1.p | Glyma.01G028300 | TALE |
| 23. | Glyma.01G029300.1.p | Glyma.01G029300 | YABBY |
| 24. | Glyma.01G034500.1.p | Glyma.01G034500 | C2H2 |
| 25. | Glyma.01G036600.1.p | Glyma.01G036600 | C2H2 |
| 26. | Glyma.01G038600.1.p | Glyma.01G038600 | MYB_related |
| 27. | Glyma.01G039200.1.p | Glyma.01G039200 | bHLH |
| 28. | Glyma.01G041700.1.p | Glyma.01G041700 | HD-ZIP |
| 29. | Glyma.01G043000.1.p | Glyma.01G043000 | Whirly |
| 30. | Glyma.01G043300.1.p | Glyma.01G043300 | WRKY |
| 31. | Glyma.01G044300.1.p | Glyma.01G044300 | MYB |
| 32. | Glyma.01G044600.1.p | Glyma.01G044600 | HD-ZIP |
| 33. | Glyma.01G044800.1.p | Glyma.01G044800 | bHLH |
| 34. | Glyma.01G045500.1.p | Glyma.01G045500 | TCP |
| 35. | Glyma.01G046800.1.p | Glyma.01G046800 | NAC |
| 36. | Glyma.01G047900.1.p | Glyma.01G047900 | ZF-HD |
| 37. | Glyma.01G049100.1.p | Glyma.01G049100 | G2-like |
| 38. | Glyma.01G049400.1.p | Glyma.01G049400 | Dof |
| 39. | Glyma.01G049600.1.p | Glyma.01G049600 | MYB |
| 40. | Glyma.01G050000.1.p | Glyma.01G050000 | MIKC_MADS |
| 41. | Glyma.01G051300.1.p | Glyma.01G051300 | NAC |
| 42. | Glyma.01G051600.1.p | Glyma.01G051600 | MYB_related |
| 43. | Glyma.01G051700.1.p | Glyma.01G051700 | MYB |
| 44. | Glyma.01G053800.1.p | Glyma.01G053800 | WRKY |
| 45. | Glyma.01G056800.1.p | Glyma.01G056800 | WRKY |
| 46. | Glyma.01G056900.1.p | Glyma.01G056900 | C2H2 |
| 47. | Glyma.01G059300.1.p | Glyma.01G059300 | MYB_related |
| 48. | Glyma.01G063500.1.p | Glyma.01G063500 | YABBY |
| 49. | Glyma.01G063700.1.p | Glyma.01G063700 | SBP |
| 50. | Glyma.01G064100.1.p | Glyma.01G064100 | MIKC_MADS |
| 51. | Glyma.01G064200.1.p | Glyma.01G064200 | MIKC_MADS |
| 52. | Glyma.01G066600.1.p | Glyma.01G066600 | bHLH |
| 53. | Glyma.01G067800.1.p | Glyma.01G067800 | MYB |
| 54. | Glyma.01G068600.1.p | Glyma.01G068600 | bHLH |
| 55. | Glyma.01G069300.1.p | Glyma.01G069300 | bZIP |
| 56. | Glyma.01G073100.1.p | Glyma.01G073100 | GATA |
| 57. | Glyma.01G074200.1.p | Glyma.01G074200 | ERF |
| 58. | Glyma.01G075800.1.p | Glyma.01G075800 | SBP |
| 59. | Glyma.01G075900.1.p | Glyma.01G075900 | SBP |
| 60. | Glyma.01G076900.1.p | Glyma.01G076900 | bHLH |
| 61. | Glyma.01G079400.1.p | Glyma.01G079400 | M-type_MADS |
| 62. | Glyma.01G079500.1.p | Glyma.01G079500 | GRAS |
| 63. | Glyma.01G079900.1.p | Glyma.01G079900 | C2H2 |
| 64. | Glyma.01G081100.1.p | Glyma.01G081100 | ERF |
| 65. | Glyma.01G084200.1.p | Glyma.01G084200 | bZIP |
| 66. | Glyma.01G086700.1.p | Glyma.01G086700 | G2-like |
| 67. | Glyma.01G086800.1.p | Glyma.01G086800 | C2H2 |
| 68. | Glyma.01G087500.1.p | Glyma.01G087500 | RAV |
| 69. | Glyma.01G088200.1.p | Glyma.01G088200 | NAC |
| 70. | Glyma.01G088300.1.p | Glyma.01G088300 | bHLH |
| 71. | Glyma.01G093400.1.p | Glyma.01G093400 | bHLH |
| 72. | Glyma.01G095100.1.p | Glyma.01G095100 | C3H |
| 73. | Glyma.01G095500.1.p | Glyma.01G095500 | bHLH |
| 74. | Glyma.01G096600.1.p | Glyma.01G096600 | bHLH |
| 75. | Glyma.01G101400.1.p | Glyma.01G101400 | MYB_related |
| 76. | Glyma.01G101800.1.p | Glyma.01G101800 | C2H2 |
| 77. | Glyma.01G103500.1.p | Glyma.01G103500 | ARF |
| 78. | Glyma.01G104200.1.p | Glyma.01G104200 | TALE |
| 79. | Glyma.01G107500.1.p | Glyma.01G107500 | MYB |
| 80. | Glyma.01G112700.1.p | Glyma.01G112700 | MYB |
| 81. | Glyma.01G113300.1.p | Glyma.01G113300 | C2H2 |
| 82. | Glyma.01G121000.1.p | Glyma.01G121000 | Trihelix |
| 83. | Glyma.01G123000.1.p | Glyma.01G123000 | bHLH |
| 84. | Glyma.01G123600.1.p | Glyma.01G123600 | G2-like |
| 85. | Glyma.01G128100.1.p | Glyma.01G128100 | WRKY |
| 86. | Glyma.01G129700.1.p | Glyma.01G129700 | bHLH |
| 87. | Glyma.01G131000.1.p | Glyma.01G131000 | LBD |
| 88. | Glyma.01G133000.1.p | Glyma.01G133000 | B3 |
| 89. | Glyma.01G133500.1.p | Glyma.01G133500 | bHLH |
| 90. | Glyma.01G134200.1.p | Glyma.01G134200 | C2H2 |
| 91. | Glyma.01G136300.1.p | Glyma.01G136300 | GRAS |
| 92. | Glyma.01G137200.1.p | Glyma.01G137200 | YABBY |
| 93. | Glyma.01G143500.1.p | Glyma.01G143500 | HSF |
| 94. | Glyma.01G143800.1.p | Glyma.01G143800 | LBD |
| 95. | Glyma.01G143900.1.p | Glyma.01G143900 | LBD |
| 96. | Glyma.01G147600.1.p | Glyma.01G147600 | ERF |
| 97. | Glyma.01G148600.1.p | Glyma.01G148600 | GRF |
| 98. | Glyma.01G150500.1.p | Glyma.01G150500 | Trihelix |
| 99. | Glyma.01G151800.1.p | Glyma.01G151800 | BES1 |
| 100. | Glyma.01G159200.1.p | Glyma.01G159200 | Nin-like |
| 101. | Glyma.01G162600.1.p | Glyma.01G162600 | G2-like |
| 102. | Glyma.01G163500.1.p | Glyma.01G163500 | bZIP |
| 103. | Glyma.01G166800.1.p | Glyma.01G166800 | WOX |
| 104. | Glyma.01G167900.1.p | Glyma.01G167900 | NAC |
| 105. | Glyma.01G168500.1.p | Glyma.01G168500 | DBB |
| 106. | Glyma.01G169400.1.p | Glyma.01G169400 | GATA |
| 107. | Glyma.01G169600.1.p | Glyma.01G169600 | MIKC_MADS |
| 108. | Glyma.01G170700.1.p | Glyma.01G170700 | SRS |
| 109. | Glyma.01G173900.1.p | Glyma.01G173900 | MYB_related |
| 110. | Glyma.01G174200.1.p | Glyma.01G174200 | ZF-HD |
| 111. | Glyma.01G176100.1.p | Glyma.01G176100 | MYB_related |
| 112. | Glyma.01G176600.1.p | Glyma.01G176600 | C2H2 |
| 113. | Glyma.01G177200.1.p | Glyma.01G177200 | GRAS |
| 114. | Glyma.01G177400.1.p | Glyma.01G177400 | bZIP |
| 115. | Glyma.01G177500.1.p | Glyma.01G177500 | HD-ZIP |
| 116. | Glyma.01G178000.1.p | Glyma.01G178000 | BES1 |
| 117. | Glyma.01G179900.1.p | Glyma.01G179900 | TALE |
| 118. | Glyma.01G183000.1.p | Glyma.01G183000 | Dof |
| 119. | Glyma.01G183700.1.p | Glyma.01G183700 | G2-like |
| 120. | Glyma.01G183900.1.p | Glyma.01G183900 | LBD |
| 121. | Glyma.01G185800.1.p | Glyma.01G185800 | HSF |
| 122. | Glyma.01G186700.1.p | Glyma.01G186700 | bHLH |
| 123. | Glyma.01G187600.1.p | Glyma.01G187600 | bHLH |
| 124. | Glyma.01G188400.1.p | Glyma.01G188400 | AP2 |
| 125. | Glyma.01G188600.1.p | Glyma.01G188600 | ERF |
| 126. | Glyma.01G189100.1.p | Glyma.01G189100 | WRKY |
| 127. | Glyma.01G190100.1.p | Glyma.01G190100 | MYB |
| 128. | Glyma.01G192600.1.p | Glyma.01G192600 | LBD |
| 129. | Glyma.01G194200.1.p | Glyma.01G194200 | GRAS |
| 130. | Glyma.01G194600.1.p | Glyma.01G194600 | MYB_related |
| 131. | Glyma.01G195800.1.p | Glyma.01G195800 | bZIP |
| 132. | Glyma.01G195900.1.p | Glyma.01G195900 | AP2 |
| 133. | Glyma.01G196200.1.p | Glyma.01G196200 | MYB |
| 134. | Glyma.01G196600.1.p | Glyma.01G196600 | HD-ZIP |
| 135. | Glyma.01G197900.1.p | Glyma.01G197900 | bHLH |
| 136. | Glyma.01G198000.1.p | Glyma.01G198000 | bHLH |
| 137. | Glyma.01G198100.1.p | Glyma.01G198100 | bHLH |
| 138. | Glyma.01G198700.1.p | Glyma.01G198700 | C2H2 |
| 139. | Glyma.01G199200.1.p | Glyma.01G199200 | LSD |
| 140. | Glyma.01G200800.1.p | Glyma.01G200800 | ARR-B |
| 141. | Glyma.01G202900.1.p | Glyma.01G202900 | FAR1 |
| 142. | Glyma.01G205100.1.p | Glyma.01G205100 | GATA |
| 143. | Glyma.01G206600.1.p | Glyma.01G206600 | ERF |
| 144. | Glyma.01G206700.1.p | Glyma.01G206700 | ERF |
| 145. | Glyma.01G207300.1.p | Glyma.01G207300 | HD-ZIP |
| 146. | Glyma.01G207600.1.p | Glyma.01G207600 | MYB |
| 147. | Glyma.01G209100.1.p | Glyma.01G209100 | C2H2 |
| 148. | Glyma.01G211500.1.p | Glyma.01G211500 | MYB |
| 149. | Glyma.01G214800.1.p | Glyma.01G214800 | TALE |
| 150. | Glyma.01G216000.1.p | Glyma.01G216000 | ERF |
| 151. | Glyma.01G216200.1.p | Glyma.01G216200 | ERF |
| 152. | Glyma.01G217400.1.p | Glyma.01G217400 | HSF |
| 153. | Glyma.01G217500.1.p | Glyma.01G217500 | MYB |
| 154. | Glyma.01G221500.1.p | Glyma.01G221500 | TALE |
| 155. | Glyma.01G222200.1.p | Glyma.01G222200 | MYB |
| 156. | Glyma.01G222300.1.p | Glyma.01G222300 | WRKY |
| 157. | Glyma.01G224100.1.p | Glyma.01G224100 | ERF |
| 158. | Glyma.01G224800.1.p | Glyma.01G224800 | WRKY |
| 159. | Glyma.01G224900.1.p | Glyma.01G224900 | MYB_related |
| 160. | Glyma.01G225000.1.p | Glyma.01G225000 | ERF |
| 161. | Glyma.01G226600.1.p | Glyma.01G226600 | GRAS |
| 162. | Glyma.01G228100.1.p | Glyma.01G228100 | E2F/DP |
| 163. | Glyma.01G228200.1.p | Glyma.01G228200 | MYB_related |
| 164. | Glyma.01G231000.1.p | Glyma.01G231000 | ERF |
| 165. | Glyma.01G231200.1.p | Glyma.01G231200 | ERF |
| 166. | Glyma.01G232000.1.p | Glyma.01G232000 | ERF |
| 167. | Glyma.01G233000.1.p | Glyma.01G233000 | HSF |
| 168. | Glyma.01G233400.1.p | Glyma.01G233400 | MYB |
| 169. | Glyma.01G234400.1.p | Glyma.01G234400 | GRF |
| 170. | Glyma.01G236300.1.p | Glyma.01G236300 | CPP |
| 171. | Glyma.01G240100.1.p | Glyma.01G240100 | HD-ZIP |

| 172. | Glyma.01G244800.1.p | Glyma.01G244800 | B3 |
| --- | --- | --- | --- |
| 173. | Glyma.02G000800.1.p | Glyma.02G000800 | bHLH |
| 174. | Glyma.02G004300.1.p | Glyma.02G004300 | C2H2 |
| 175. | Glyma.02G005600.1.p | Glyma.02G005600 | MYB |
| 176. | Glyma.02G006200.1.p | Glyma.02G006200 | ERF |
| 177. | Glyma.02G006300.1.p | Glyma.02G006300 | ERF |
| 178. | Glyma.02G006800.1.p | Glyma.02G006800 | MYB |
| 179. | Glyma.02G007000.1.p | Glyma.02G007000 | bHLH |
| 180. | Glyma.02G007500.1.p | Glyma.02G007500 | WRKY |
| 181. | Glyma.02G008600.1.p | Glyma.02G008600 | SBP |
| 182. | Glyma.02G009800.1.p | Glyma.02G009800 | MYB |
| 183. | Glyma.02G010900.1.p | Glyma.02G010900 | WRKY |
| 184. | Glyma.02G012000.1.p | Glyma.02G012000 | GRAS |
| 185. | Glyma.02G012700.1.p | Glyma.02G012700 | bZIP |
| 186. | Glyma.02G013900.1.p | Glyma.02G013900 | MYB |
| 187. | Glyma.02G016100.1.p | Glyma.02G016100 | ERF |
| 188. | Glyma.02G018900.1.p | Glyma.02G018900 | bHLH |
| 189. | Glyma.02G019100.1.p | Glyma.02G019100 | HD-ZIP |
| 190. | Glyma.02G019300.1.p | Glyma.02G019300 | MYB |
| 191. | Glyma.02G019900.1.p | Glyma.02G019900 | Whirly |
| 192. | Glyma.02G020300.1.p | Glyma.02G020300 | WRKY |
| 193. | Glyma.02G020400.1.p | Glyma.02G020400 | Whirly |
| 194. | Glyma.02G022300.1.p | Glyma.02G022300 | HD-ZIP |
| 195. | Glyma.02G025400.1.p | Glyma.02G025400 | bHLH |
| 196. | Glyma.02G025500.1.p | Glyma.02G025500 | bHLH |
| 197. | Glyma.02G026300.1.p | Glyma.02G026300 | MYB_related |
| 198. | Glyma.02G029400.1.p | Glyma.02G029400 | C2H2 |
| 199. | Glyma.02G036900.1.p | Glyma.02G036900 | TALE |
| 200. | Glyma.02G039300.1.p | Glyma.02G039300 | ERF |
| 201. | Glyma.02G040100.1.p | Glyma.02G040100 | ZF-HD |
| 202. | Glyma.02G041000.1.p | Glyma.02G041000 | bHLH |
| 203. | Glyma.02G041500.1.p | Glyma.02G041500 | MIKC_MADS |
| 204. | Glyma.02G045500.1.p | Glyma.02G045500 | bZIP |
| 205. | Glyma.02G048100.1.p | Glyma.02G048100 | MYB_related |
| 206. | Glyma.02G050100.1.p | Glyma.02G050100 | NAC |
| 207. | Glyma.02G051100.1.p | Glyma.02G051100 | GATA |
| 208. | Glyma.02G051900.1.p | Glyma.02G051900 | SRS |
| 209. | Glyma.02G053100.1.p | Glyma.02G053100 | bHLH |
| 210. | Glyma.02G053100.2.p | Glyma.02G053100 | bHLH |
| 211. | Glyma.02G053100.3.p | Glyma.02G053100 | bHLH |
| 212. | Glyma.02G055800.1.p | Glyma.02G055800 | MYB_related |
| 213. | Glyma.02G055900.1.p | Glyma.02G055900 | MYB_related |
| 214. | Glyma.02G056100.1.p | Glyma.02G056100 | ZF-HD |
| 215. | Glyma.02G056600.1.p | Glyma.02G056600 | GATA |
| 216. | Glyma.02G058400.1.p | Glyma.02G058400 | C2H2 |
| 217. | Glyma.02G058500.1.p | Glyma.02G058500 | C2H2 |
| 218. | Glyma.02G058700.1.p | Glyma.02G058700 | GRAS |
| 219. | Glyma.02G058800.1.p | Glyma.02G058800 | bZIP |
| 220. | Glyma.02G058900.1.p | Glyma.02G058900 | HD-ZIP |
| 221. | Glyma.02G060000.1.p | Glyma.02G060000 | TALE |
| 222. | Glyma.02G062700.1.p | Glyma.02G062700 | Dof |
| 223. | Glyma.02G066200.1.p | Glyma.02G066200 | ERF |
| 224. | Glyma.02G067600.1.p | Glyma.02G067600 | ERF |
| 225. | Glyma.02G070000.1.p | Glyma.02G070000 | NAC |
| 226. | Glyma.02G070600.1.p | Glyma.02G070600 | NAC |
| 227. | Glyma.02G070900.1.p | Glyma.02G070900 | G2-like |
| 228. | Glyma.02G071500.1.p | Glyma.02G071500 | GATA |
| 229. | Glyma.02G072800.1.p | Glyma.02G072800 | ERF |
| 230. | Glyma.02G073900.1.p | Glyma.02G073900 | GATA |
| 231. | Glyma.02G074800.1.p | Glyma.02G074800 | GRAS |
| 232. | Glyma.02G080200.1.p | Glyma.02G080200 | ERF |
| 233. | Glyma.02G082000.1.p | Glyma.02G082000 | Trihelix |
| 234. | Glyma.02G082100.1.p | Glyma.02G082100 | Trihelix |
| 235. | Glyma.02G082800.1.p | Glyma.02G082800 | bZIP |
| 236. | Glyma.02G085900.1.p | Glyma.02G085900 | ARR-B |
| 237. | Glyma.02G087200.1.p | Glyma.02G087200 | NF-YB |
| 238. | Glyma.02G087400.1.p | Glyma.02G087400 | AP2 |
| 239. | Glyma.02G089600.1.p | Glyma.02G089600 | NF-YC |
| 240. | Glyma.02G092700.1.p | Glyma.02G092700 | Dof |
| 241. | Glyma.02G093900.1.p | Glyma.02G093900 | WOX |
| 242. | Glyma.02G094500.1.p | Glyma.02G094500 | C2H2 |
| 243. | Glyma.02G097900.1.p | Glyma.02G097900 | bZIP |
| 244. | Glyma.02G098800.1.p | Glyma.02G098800 | G2-like |
| 245. | Glyma.02G099100.1.p | Glyma.02G099100 | C2H2 |
| 246. | Glyma.02G099500.1.p | Glyma.02G099500 | RAV |
| 247. | Glyma.02G100200.1.p | Glyma.02G100200 | NAC |
| 248. | Glyma.02G100700.1.p | Glyma.02G100700 | bHLH |
| 249. | Glyma.02G103100.1.p | Glyma.02G103100 | bHLH |
| 250. | Glyma.02G103200.1.p | Glyma.02G103200 | bHLH |
| 251. | Glyma.02G105900.1.p | Glyma.02G105900 | TCP |
| 252. | Glyma.02G107000.1.p | Glyma.02G107000 | NAC |
| 253. | Glyma.02G107300.1.p | Glyma.02G107300 | ZF-HD |
| 254. | Glyma.02G108500.1.p | Glyma.02G108500 | G2-like |
| 255. | Glyma.02G108600.1.p | Glyma.02G108600 | Dof |
| 256. | Glyma.02G108800.1.p | Glyma.02G108800 | MYB |
| 257. | Glyma.02G109000.1.p | Glyma.02G109000 | M-type_MADS |
| 258. | Glyma.02G109800.1.p | Glyma.02G109800 | NAC |
| 259. | Glyma.02G110000.1.p | Glyma.02G110000 | MYB |
| 260. | Glyma.02G110100.1.p | Glyma.02G110100 | MYB |
| 261. | Glyma.02G110200.1.p | Glyma.02G110200 | MYB |
| 262. | Glyma.02G112100.1.p | Glyma.02G112100 | WRKY |
| 263. | Glyma.02G115200.1.p | Glyma.02G115200 | WRKY |
| 264. | Glyma.02G115300.1.p | Glyma.02G115300 | C2H2 |
| 265. | Glyma.02G121100.1.p | Glyma.02G121100 | YABBY |
| 266. | Glyma.02G121300.1.p | Glyma.02G121300 | SBP |
| 267. | Glyma.02G121500.1.p | Glyma.02G121500 | MIKC_MADS |
| 268. | Glyma.02G121600.1.p | Glyma.02G121600 | MIKC_MADS |
| 269. | Glyma.02G123400.1.p | Glyma.02G123400 | bHLH |
| 270. | Glyma.02G124300.1.p | Glyma.02G124300 | MYB |
| 271. | Glyma.02G125200.1.p | Glyma.02G125200 | bHLH |
| 272. | Glyma.02G126100.1.p | Glyma.02G126100 | bZIP |
| 273. | Glyma.02G129100.1.p | Glyma.02G129100 | bHLH |
| 274. | Glyma.02G131700.1.p | Glyma.02G131700 | bZIP |
| 275. | Glyma.02G132500.1.p | Glyma.02G132500 | ERF |
| 276. | Glyma.02G135400.1.p | Glyma.02G135400 | G2-like |
| 277. | Glyma.02G137400.1.p | Glyma.02G137400 | bHLH |
| 278. | Glyma.02G139700.1.p | Glyma.02G139700 | bHLH |
| 279. | Glyma.02G141000.1.p | Glyma.02G141000 | WRKY |
| 280. | Glyma.02G142800.1.p | Glyma.02G142800 | bHLH |
| 281. | Glyma.02G144400.1.p | Glyma.02G144400 | C2H2 |
| 282. | Glyma.02G147800.1.p | Glyma.02G147800 | bHLH |
| 283. | Glyma.02G148500.1.p | Glyma.02G148500 | bHLH |
| 284. | Glyma.02G152900.1.p | Glyma.02G152900 | CO-like |
| 285. | Glyma.02G153600.1.p | Glyma.02G153600 | C3H |
| 286. | Glyma.02G153700.1.p | Glyma.02G153700 | C3H |
| 287. | Glyma.02G153900.1.p | Glyma.02G153900 | C2H2 |
| 288. | Glyma.02G154000.1.p | Glyma.02G154000 | NF-YB |
| 289. | Glyma.02G157300.1.p | Glyma.02G157300 | MYB_related |
| 290. | Glyma.02G160200.1.p | Glyma.02G160200 | bHLH |
| 291. | Glyma.02G161100.1.p | Glyma.02G161100 | bZIP |
| 292. | Glyma.02G162200.1.p | Glyma.02G162200 | C2H2 |
| 293. | Glyma.02G162600.1.p | Glyma.02G162600 | G2-like |
| 294. | Glyma.02G166400.1.p | Glyma.02G166400 | LBD |
| 295. | Glyma.02G171300.1.p | Glyma.02G171300 | HD-ZIP |
| 296. | Glyma.02G173900.1.p | Glyma.02G173900 | C3H |
| 297. | Glyma.02G174800.1.p | Glyma.02G174800 | bHLH |
| 298. | Glyma.02G175700.1.p | Glyma.02G175700 | bHLH |
| 299. | Glyma.02G176800.1.p | Glyma.02G176800 | bZIP |
| 300. | Glyma.02G177500.1.p | Glyma.02G177500 | SBP |
| 301. | Glyma.02G177800.1.p | Glyma.02G177800 | G2-like |
| 302. | Glyma.02G178100.1.p | Glyma.02G178100 | G2-like |
| 303. | Glyma.02G179300.1.p | Glyma.02G179300 | M-type_MADS |
| 304. | Glyma.02G179500.1.p | Glyma.02G179500 | AP2 |
| 305. | Glyma.02G180800.1.p | Glyma.02G180800 | ZF-HD |
| 306. | Glyma.02G180900.1.p | Glyma.02G180900 | ZF-HD |
| 307. | Glyma.02G181100.1.p | Glyma.02G181100 | C2H2 |
| 308. | Glyma.02G181600.1.p | Glyma.02G181600 | ERF |
| 309. | Glyma.02G183100.1.p | Glyma.02G183100 | C2H2 |
| 310. | Glyma.02G185200.1.p | Glyma.02G185200 | AP2 |
| 311. | Glyma.02G185500.1.p | Glyma.02G185500 | MIKC_MADS |
| 312. | Glyma.02G187400.1.p | Glyma.02G187400 | M-type_MADS |
| 313. | Glyma.02G188900.1.p | Glyma.02G188900 | M-type_MADS |
| 314. | Glyma.02G189500.1.p | Glyma.02G189500 | M-type_MADS |
| 315. | Glyma.02G194000.1.p | Glyma.02G194000 | M-type_MADS |
| 316. | Glyma.02G194300.1.p | Glyma.02G194300 | M-type_MADS |
| 317. | Glyma.02G195000.1.p | Glyma.02G195000 | NF-YA |
| 318. | Glyma.02G195700.1.p | Glyma.02G195700 | Dof |
| 319. | Glyma.02G196500.1.p | Glyma.02G196500 | TALE |
| 320. | Glyma.02G198700.1.p | Glyma.02G198700 | E2F/DP |
| 321. | Glyma.02G200500.1.p | Glyma.02G200500 | B3 |
| 322. | Glyma.02G202600.1.p | Glyma.02G202600 | bHLH |
| 323. | Glyma.02G203800.1.p | Glyma.02G203800 | WRKY |
| 324. | Glyma.02G207100.1.p | Glyma.02G207100 | AP2 |
| 325. | Glyma.02G210500.1.p | Glyma.02G210500 | bHLH |
| 326. | Glyma.02G211900.1.p | Glyma.02G211900 | ZF-HD |
| 327. | Glyma.02G213800.1.p | Glyma.02G213800 | bHLH |
| 328. | Glyma.02G214600.1.p | Glyma.02G214600 | MYB_related |
| 329. | Glyma.02G215600.1.p | Glyma.02G215600 | GATA |
| 330. | Glyma.02G216600.1.p | Glyma.02G216600 | MIKC_MADS |
| 331. | Glyma.02G217800.1.p | Glyma.02G217800 | bHLH |
| 332. | Glyma.02G222300.1.p | Glyma.02G222300 | NAC |
| 333. | Glyma.02G223700.1.p | Glyma.02G223700 | CO-like |
| 334. | Glyma.02G224900.1.p | Glyma.02G224900 | MYB |
| 335. | Glyma.02G225700.1.p | Glyma.02G225700 | MYB |
| 336. | Glyma.02G227000.1.p | Glyma.02G227000 | C3H |
| 337. | Glyma.02G232600.1.p | Glyma.02G232600 | WRKY |
| 338. | Glyma.02G234600.1.p | Glyma.02G234600 | LBD |
| 339. | Glyma.02G236200.1.p | Glyma.02G236200 | bZIP |
| 340. | Glyma.02G236400.1.p | Glyma.02G236400 | B3 |
| 341. | Glyma.02G236800.1.p | Glyma.02G236800 | ERF |
| 342. | Glyma.02G237500.1.p | Glyma.02G237500 | B3 |
| 343. | Glyma.02G239600.1.p | Glyma.02G239600 | ARF |
| 344. | Glyma.02G240500.1.p | Glyma.02G240500 | NAC |
| 345. | Glyma.02G241000.1.p | Glyma.02G241000 | MYB_related |
| 346. | Glyma.02G242100.1.p | Glyma.02G242100 | G2-like |
| 347. | Glyma.02G242400.1.p | Glyma.02G242400 | C2H2 |
| 348. | Glyma.02G244600.1.p | Glyma.02G244600 | MYB |
| 349. | Glyma.02G246400.1.p | Glyma.02G246400 | bHLH |
| 350. | Glyma.02G247100.1.p | Glyma.02G247100 | MYB |
| 351. | Glyma.02G249400.1.p | Glyma.02G249400 | MYB_related |
| 352. | Glyma.02G249700.1.p | Glyma.02G249700 | LBD |
| 353. | Glyma.02G250100.1.p | Glyma.02G250100 | C2H2 |
| 354. | Glyma.02G252400.1.p | Glyma.02G252400 | bHLH |
| 355. | Glyma.02G253100.1.p | Glyma.02G253100 | MYB |
| 356. | Glyma.02G254200.1.p | Glyma.02G254200 | MYB |
| 357. | Glyma.02G254800.1.p | Glyma.02G254800 | WOX |
| 358. | Glyma.02G258300.1.p | Glyma.02G258300 | bHLH |
| 359. | Glyma.02G261700.1.p | Glyma.02G261700 | ERF |
| 360. | Glyma.02G262400.1.p | Glyma.02G262400 | LBD |
| 361. | Glyma.02G264500.1.p | Glyma.02G264500 | LBD |
| 362. | Glyma.02G264700.1.p | Glyma.02G264700 | ERF |
| 363. | Glyma.02G264900.1.p | Glyma.02G264900 | MYB |
| 364. | Glyma.02G267400.1.p | Glyma.02G267400 | ERF |
| 365. | Glyma.02G267500.1.p | Glyma.02G267500 | C3H |
| 366. | Glyma.02G269900.1.p | Glyma.02G269900 | HB-other |
| 367. | Glyma.02G273500.1.p | Glyma.02G273500 | FAR1 |
| 368. | Glyma.02G274600.1.p | Glyma.02G274600 | EIL |
| 369. | Glyma.02G277000.1.p | Glyma.02G277000 | NF-YC |
| 370. | Glyma.02G278400.1.p | Glyma.02G278400 | HSF |
| 371. | Glyma.02G280000.1.p | Glyma.02G280000 | SRS |
| 372. | Glyma.02G280300.1.p | Glyma.02G280300 | HB-other |
| 373. | Glyma.02G281700.1.p | Glyma.02G281700 | ARF |
| 374. | Glyma.02G282100.1.p | Glyma.02G282100 | bHLH |
| 375. | Glyma.02G282900.1.p | Glyma.02G282900 | G2-like |
| 376. | Glyma.02G284300.1.p | Glyma.02G284300 | NAC |
| 377. | Glyma.02G285900.1.p | Glyma.02G285900 | WRKY |
| 378. | Glyma.02G287700.1.p | Glyma.02G287700 | MIKC_MADS |
| 379. | Glyma.02G292000.1.p | Glyma.02G292000 | LBD |
| 380. | Glyma.02G292900.1.p | Glyma.02G292900 | C2H2 |
| 381. | Glyma.02G293300.1.p | Glyma.02G293300 | C2H2 |
| 382. | Glyma.02G293400.1.p | Glyma.02G293400 | WRKY |
| 383. | Glyma.02G294100.1.p | Glyma.02G294100 | ERF |
| 384. | Glyma.02G296600.1.p | Glyma.02G296600 | C3H |
| 385. | Glyma.02G297400.1.p | Glyma.02G297400 | WRKY |
| 386. | Glyma.02G297700.1.p | Glyma.02G297700 | GRAS |
| 387. | Glyma.02G300200.1.p | Glyma.02G300200 | NF-YB |
| 388. | Glyma.02G303800.1.p | Glyma.02G303800 | NF-YA |
| 389. | Glyma.02G304600.1.p | Glyma.02G304600 | SAP |
| 390. | Glyma.02G306200.1.p | Glyma.02G306200 | GRAS |
| 391. | Glyma.02G306300.1.p | Glyma.02G306300 | WRKY |
| 392. | Glyma.02G311000.1.p | Glyma.02G311000 | Nin-like |
| 393. | Glyma.02G311100.1.p | Glyma.02G311100 | C2H2 |

| 394. | Glyma.03G002300.1.p | Glyma.03G002300 | WRKY |
| --- | --- | --- | --- |
| 395. | Glyma.03G003400.1.p | Glyma.03G003400 | bZIP |
| 396. | Glyma.03G003500.1.p | Glyma.03G003500 | G2-like |
| 397. | Glyma.03G006600.1.p | Glyma.03G006600 | MYB |
| 398. | Glyma.03G007500.1.p | Glyma.03G007500 | MYB |
| 399. | Glyma.03G007600.1.p | Glyma.03G007600 | WOX |
| 400. | Glyma.03G007900.1.p | Glyma.03G007900 | Dof |
| 401. | Glyma.03G013300.1.p | Glyma.03G013300 | MYB |
| 402. | Glyma.03G016600.1.p | Glyma.03G016600 | HD-ZIP |
| 403. | Glyma.03G017300.1.p | Glyma.03G017300 | ERF |
| 404. | Glyma.03G018000.1.p | Glyma.03G018000 | C3H |
| 405. | Glyma.03G018800.1.p | Glyma.03G018800 | TCP |
| 406. | Glyma.03G019300.1.p | Glyma.03G019300 | MIKC_MADS |
| 407. | Glyma.03G019400.1.p | Glyma.03G019400 | MIKC_MADS |
| 408. | Glyma.03G023000.1.p | Glyma.03G023000 | LBD |
| 409. | Glyma.03G023100.1.p | Glyma.03G023100 | LBD |
| 410. | Glyma.03G025400.1.p | Glyma.03G025400 | C3H |
| 411. | Glyma.03G029800.1.p | Glyma.03G029800 | YABBY |
| 412. | Glyma.03G031800.1.p | Glyma.03G031800 | GRAS |
| 413. | Glyma.03G033600.1.p | Glyma.03G033600 | C2H2 |
| 414. | Glyma.03G034000.1.p | Glyma.03G034000 | bHLH |
| 415. | Glyma.03G035200.1.p | Glyma.03G035200 | B3 |
| 416. | Glyma.03G042700.1.p | Glyma.03G042700 | WRKY |
| 417. | Glyma.03G047600.1.p | Glyma.03G047600 | MYB_related |
| 418. | Glyma.03G050800.1.p | Glyma.03G050800 | GRAS |
| 419. | Glyma.03G051400.1.p | Glyma.03G051400 | G2-like |
| 420. | Glyma.03G052300.1.p | Glyma.03G052300 | bHLH |
| 421. | Glyma.03G056900.1.p | Glyma.03G056900 | Trihelix |
| 422. | Glyma.03G065700.1.p | Glyma.03G065700 | GRAS |
| 423. | Glyma.03G067000.1.p | Glyma.03G067000 | MYB_related |
| 424. | Glyma.03G067700.1.p | Glyma.03G067700 | C2H2 |
| 425. | Glyma.03G070500.1.p | Glyma.03G070500 | ARF |
| 426. | Glyma.03G070900.1.p | Glyma.03G070900 | TALE |
| 427. | Glyma.03G073900.1.p | Glyma.03G073900 | bZIP |
| 428. | Glyma.03G078000.1.p | Glyma.03G078000 | MYB |
| 429. | Glyma.03G080000.1.p | Glyma.03G080000 | M-type_MADS |
| 430. | Glyma.03G080700.1.p | Glyma.03G080700 | NF-YB |
| 431. | Glyma.03G081500.1.p | Glyma.03G081500 | Trihelix |
| 432. | Glyma.03G081700.1.p | Glyma.03G081700 | bZIP |
| 433. | Glyma.03G081900.1.p | Glyma.03G081900 | MYB |
| 434. | Glyma.03G082400.1.p | Glyma.03G082400 | MYB |
| 435. | Glyma.03G083700.1.p | Glyma.03G083700 | M-type_MADS |
| 436. | Glyma.03G086800.1.p | Glyma.03G086800 | bHLH |
| 437. | Glyma.03G090800.1.p | Glyma.03G090800 | MYB_related |
| 438. | Glyma.03G091500.1.p | Glyma.03G091500 | FAR1 |
| 439. | Glyma.03G091600.1.p | Glyma.03G091600 | G2-like |
| 440. | Glyma.03G091700.1.p | Glyma.03G091700 | NF-YB |
| 441. | Glyma.03G094700.1.p | Glyma.03G094700 | ERF |
| 442. | Glyma.03G101600.1.p | Glyma.03G101600 | bHLH |
| 443. | Glyma.03G105000.1.p | Glyma.03G105000 | bHLH |
| 444. | Glyma.03G105700.1.p | Glyma.03G105700 | bHLH |
| 445. | Glyma.03G107400.1.p | Glyma.03G107400 | FAR1 |
| 446. | Glyma.03G109100.1.p | Glyma.03G109100 | WRKY |
| 447. | Glyma.03G111500.1.p | Glyma.03G111500 | MIKC_MADS |
| 448. | Glyma.03G111700.1.p | Glyma.03G111700 | ERF |
| 449. | Glyma.03G112000.1.p | Glyma.03G112000 | ERF |
| 450. | Glyma.03G112100.1.p | Glyma.03G112100 | ERF |
| 451. | Glyma.03G112400.1.p | Glyma.03G112400 | ERF |
| 452. | Glyma.03G112600.1.p | Glyma.03G112600 | ERF |
| 453. | Glyma.03G112700.1.p | Glyma.03G112700 | ERF |
| 454. | Glyma.03G112800.1.p | Glyma.03G112800 | ERF |
| 455. | Glyma.03G114300.1.p | Glyma.03G114300 | HD-ZIP |
| 456. | Glyma.03G115400.1.p | Glyma.03G115400 | MYB |
| 457. | Glyma.03G116700.1.p | Glyma.03G116700 | ERF |
| 458. | Glyma.03G117500.1.p | Glyma.03G117500 | SBP |
| 459. | Glyma.03G117600.1.p | Glyma.03G117600 | SBP |
| 460. | Glyma.03G118100.1.p | Glyma.03G118100 | GATA |
| 461. | Glyma.03G123200.1.p | Glyma.03G123200 | bZIP |
| 462. | Glyma.03G123400.1.p | Glyma.03G123400 | G2-like |
| 463. | Glyma.03G125000.1.p | Glyma.03G125000 | MYB_related |
| 464. | Glyma.03G126100.1.p | Glyma.03G126100 | bHLH |
| 465. | Glyma.03G127600.1.p | Glyma.03G127600 | bZIP |
| 466. | Glyma.03G128200.1.p | Glyma.03G128200 | bZIP |
| 467. | Glyma.03G130400.1.p | Glyma.03G130400 | bHLH |
| 468. | Glyma.03G130600.1.p | Glyma.03G130600 | bHLH |
| 469. | Glyma.03G135800.1.p | Glyma.03G135800 | HSF |
| 470. | Glyma.03G136100.1.p | Glyma.03G136100 | AP2 |
| 471. | Glyma.03G136800.1.p | Glyma.03G136800 | FAR1 |
| 472. | Glyma.03G137500.1.p | Glyma.03G137500 | C2H2 |
| 473. | Glyma.03G138600.1.p | Glyma.03G138600 | C3H |
| 474. | Glyma.03G138900.1.p | Glyma.03G138900 | ERF |
| 475. | Glyma.03G139800.1.p | Glyma.03G139800 | C2H2 |
| 476. | Glyma.03G140800.1.p | Glyma.03G140800 | ERF |
| 477. | Glyma.03G141200.1.p | Glyma.03G141200 | bHLH |
| 478. | Glyma.03G141700.1.p | Glyma.03G141700 | bHLH |
| 479. | Glyma.03G142400.1.p | Glyma.03G142400 | bZIP |
| 480. | Glyma.03G143100.1.p | Glyma.03G143100 | SBP |
| 481. | Glyma.03G143500.1.p | Glyma.03G143500 | G2-like |
| 482. | Glyma.03G143600.1.p | Glyma.03G143600 | G2-like |
| 483. | Glyma.03G144200.1.p | Glyma.03G144200 | C3H |
| 484. | Glyma.03G144300.1.p | Glyma.03G144300 | C3H |
| 485. | Glyma.03G145800.1.p | Glyma.03G145800 | HD-ZIP |
| 486. | Glyma.03G152800.1.p | Glyma.03G152800 | bHLH |
| 487. | Glyma.03G157300.1.p | Glyma.03G157300 | HSF |
| 488. | Glyma.03G157400.1.p | Glyma.03G157400 | C2H2 |
| 489. | Glyma.03G158500.1.p | Glyma.03G158500 | bHLH |
| 490. | Glyma.03G159700.1.p | Glyma.03G159700 | WRKY |
| 491. | Glyma.03G159800.1.p | Glyma.03G159800 | ERF |
| 492. | Glyma.03G160700.1.p | Glyma.03G160700 | C2H2 |
| 493. | Glyma.03G161400.1.p | Glyma.03G161400 | LBD |
| 494. | Glyma.03G161500.1.p | Glyma.03G161500 | LBD |
| 495. | Glyma.03G161800.1.p | Glyma.03G161800 | bHLH |
| 496. | Glyma.03G162400.1.p | Glyma.03G162400 | ERF |
| 497. | Glyma.03G162500.1.p | Glyma.03G162500 | ERF |
| 498. | Glyma.03G162600.1.p | Glyma.03G162600 | ERF |
| 499. | Glyma.03G162700.1.p | Glyma.03G162700 | ERF |
| 500. | Glyma.03G163100.1.p | Glyma.03G163100 | MYB |
| 501. | Glyma.03G164200.1.p | Glyma.03G164200 | NAC |
| 502. | Glyma.03G166400.1.p | Glyma.03G166400 | G2-like |
| 503. | Glyma.03G170300.1.p | Glyma.03G170300 | bHLH |
| 504. | Glyma.03G173100.1.p | Glyma.03G173100 | C2H2 |
| 505. | Glyma.03G173200.1.p | Glyma.03G173200 | C2H2 |
| 506. | Glyma.03G173300.1.p | Glyma.03G173300 | C2H2 |
| 507. | Glyma.03G176600.1.p | Glyma.03G176600 | WRKY |
| 508. | Glyma.03G177300.1.p | Glyma.03G177300 | MYB_related |
| 509. | Glyma.03G177500.1.p | Glyma.03G177500 | AP2 |
| 510. | Glyma.03G177700.1.p | Glyma.03G177700 | NF-YB |
| 511. | Glyma.03G179000.1.p | Glyma.03G179000 | S1Fa-like |
| 512. | Glyma.03G179600.1.p | Glyma.03G179600 | NAC |
| 513. | Glyma.03G179700.1.p | Glyma.03G179700 | C2H2 |
| 514. | Glyma.03G183900.1.p | Glyma.03G183900 | MYB |
| 515. | Glyma.03G186100.1.p | Glyma.03G186100 | C2H2 |
| 516. | Glyma.03G189400.1.p | Glyma.03G189400 | HD-ZIP |
| 517. | Glyma.03G189600.1.p | Glyma.03G189600 | Trihelix |
| 518. | Glyma.03G190700.1.p | Glyma.03G190700 | C2H2 |
| 519. | Glyma.03G191100.1.p | Glyma.03G191100 | HSF |
| 520. | Glyma.03G191700.1.p | Glyma.03G191700 | Trihelix |
| 521. | Glyma.03G191800.1.p | Glyma.03G191800 | ERF |
| 522. | Glyma.03G192200.1.p | Glyma.03G192200 | GRF |
| 523. | Glyma.03G193300.1.p | Glyma.03G193300 | bZIP |
| 524. | Glyma.03G197900.1.p | Glyma.03G197900 | NAC |
| 525. | Glyma.03G199000.1.p | Glyma.03G199000 | B3 |
| 526. | Glyma.03G202300.1.p | Glyma.03G202300 | TALE |
| 527. | Glyma.03G203000.1.p | Glyma.03G203000 | NF-YA |
| 528. | Glyma.03G203300.1.p | Glyma.03G203300 | M-type_MADS |
| 529. | Glyma.03G208800.1.p | Glyma.03G208800 | ARF |
| 530. | Glyma.03G209800.1.p | Glyma.03G209800 | CO-like |
| 531. | Glyma.03G211500.1.p | Glyma.03G211500 | HB-other |
| 532. | Glyma.03G211700.1.p | Glyma.03G211700 | C2H2 |
| 533. | Glyma.03G217900.1.p | Glyma.03G217900 | C2H2 |
| 534. | Glyma.03G218200.1.p | Glyma.03G218200 | MYB |
| 535. | Glyma.03G219300.1.p | Glyma.03G219300 | bZIP |
| 536. | Glyma.03G219900.1.p | Glyma.03G219900 | GRAS |
| 537. | Glyma.03G220100.1.p | Glyma.03G220100 | WRKY |
| 538. | Glyma.03G220800.1.p | Glyma.03G220800 | WRKY |
| 539. | Glyma.03G221500.1.p | Glyma.03G221500 | MYB_related |
| 540. | Glyma.03G221700.1.p | Glyma.03G221700 | MYB |
| 541. | Glyma.03G221900.1.p | Glyma.03G221900 | MYB |
| 542. | Glyma.03G224700.1.p | Glyma.03G224700 | WRKY |
| 543. | Glyma.03G225000.1.p | Glyma.03G225000 | bHLH |
| 544. | Glyma.03G225200.1.p | Glyma.03G225200 | MYB |
| 545. | Glyma.03G227700.1.p | Glyma.03G227700 | MYB |
| 546. | Glyma.03G227800.1.p | Glyma.03G227800 | bHLH |
| 547. | Glyma.03G228100.1.p | Glyma.03G228100 | C2H2 |
| 548. | Glyma.03G231100.1.p | Glyma.03G231100 | TALE |
| 549. | Glyma.03G232900.1.p | Glyma.03G232900 | GATA |
| 550. | Glyma.03G236600.1.p | Glyma.03G236600 | C2H2 |
| 551. | Glyma.03G239400.1.p | Glyma.03G239400 | NF-YC |
| 552. | Glyma.03G240000.1.p | Glyma.03G240000 | bHLH |
| 553. | Glyma.03G245200.1.p | Glyma.03G245200 | GeBP |
| 554. | Glyma.03G246000.1.p | Glyma.03G246000 | Trihelix |
| 555. | Glyma.03G246400.1.p | Glyma.03G246400 | B3 |
| 556. | Glyma.03G247100.1.p | Glyma.03G247100 | bZIP |
| 557. | Glyma.03G250000.1.p | Glyma.03G250000 | G2-like |
| 558. | Glyma.03G250600.1.p | Glyma.03G250600 | MYB |
| 559. | Glyma.03G252100.1.p | Glyma.03G252100 | Whirly |
| 560. | Glyma.03G255000.1.p | Glyma.03G255000 | bZIP |
| 561. | Glyma.03G255500.1.p | Glyma.03G255500 | ERF |
| 562. | Glyma.03G256700.1.p | Glyma.03G256700 | WRKY |
| 563. | Glyma.03G258100.1.p | Glyma.03G258100 | bHLH |
| 564. | Glyma.03G258200.1.p | Glyma.03G258200 | ERF |
| 565. | Glyma.03G258300.1.p | Glyma.03G258300 | ARF |
| 566. | Glyma.03G258700.1.p | Glyma.03G258700 | MYB |
| 567. | Glyma.03G258800.1.p | Glyma.03G258800 | Dof |
| 568. | Glyma.03G261800.1.p | Glyma.03G261800 | MYB_related |
| 569. | Glyma.03G262200.1.p | Glyma.03G262200 | B3 |
| 570. | Glyma.03G263700.1.p | Glyma.03G263700 | ERF |

| 571. | Glyma.04G000600.1.p | Glyma.04G000600 | Nin-like |
| --- | --- | --- | --- |
| 572. | Glyma.04G004100.1.p | Glyma.04G004100 | MYB |
| 573. | Glyma.04G004700.1.p | Glyma.04G004700 | MYB_related |
| 574. | Glyma.04G008900.1.p | Glyma.04G008900 | GATA |
| 575. | Glyma.04G009200.1.p | Glyma.04G009200 | DBB |
| 576. | Glyma.04G009300.1.p | Glyma.04G009300 | SRS |
| 577. | Glyma.04G009600.1.p | Glyma.04G009600 | TALE |
| 578. | Glyma.04G010300.1.p | Glyma.04G010300 | bZIP |
| 579. | Glyma.04G012500.1.p | Glyma.04G012500 | bHLH |
| 580. | Glyma.04G012600.1.p | Glyma.04G012600 | MYB_related |
| 581. | Glyma.04G014900.1.p | Glyma.04G014900 | NAC |
| 582. | Glyma.04G016700.1.p | Glyma.04G016700 | WOX |
| 583. | Glyma.04G017400.1.p | Glyma.04G017400 | Nin-like |
| 584. | Glyma.04G022100.1.p | Glyma.04G022100 | bZIP |
| 585. | Glyma.04G024800.1.p | Glyma.04G024800 | E2F/DP |
| 586. | Glyma.04G026200.1.p | Glyma.04G026200 | bZIP |
| 587. | Glyma.04G026300.1.p | Glyma.04G026300 | BBR-BPC |
| 588. | Glyma.04G027000.1.p | Glyma.04G027000 | DBB |
| 589. | Glyma.04G027200.1.p | Glyma.04G027200 | MIKC_MADS |
| 590. | Glyma.04G027400.1.p | Glyma.04G027400 | SRS |
| 591. | Glyma.04G028100.1.p | Glyma.04G028100 | ERF |
| 592. | Glyma.04G029100.1.p | Glyma.04G029100 | TALE |
| 593. | Glyma.04G029200.1.p | Glyma.04G029200 | TALE |
| 594. | Glyma.04G029600.1.p | Glyma.04G029600 | bZIP |
| 595. | Glyma.04G031300.1.p | Glyma.04G031300 | MYB_related |
| 596. | Glyma.04G031700.1.p | Glyma.04G031700 | MYB_related |
| 597. | Glyma.04G033300.1.p | Glyma.04G033300 | C2H2 |
| 598. | Glyma.04G033800.1.p | Glyma.04G033800 | BES1 |
| 599. | Glyma.04G035700.1.p | Glyma.04G035700 | G2-like |
| 600. | Glyma.04G036700.1.p | Glyma.04G036700 | MYB |
| 601. | Glyma.04G037800.1.p | Glyma.04G037800 | B3 |
| 602. | Glyma.04G037900.1.p | Glyma.04G037900 | B3 |
| 603. | Glyma.04G039300.1.p | Glyma.04G039300 | bZIP |
| 604. | Glyma.04G039500.1.p | Glyma.04G039500 | bHLH |
| 605. | Glyma.04G039600.1.p | Glyma.04G039600 | HSF |
| 606. | Glyma.04G040900.1.p | Glyma.04G040900 | WOX |
| 607. | Glyma.04G041200.1.p | Glyma.04G041200 | ERF |
| 608. | Glyma.04G042300.1.p | Glyma.04G042300 | MYB |
| 609. | Glyma.04G043800.1.p | Glyma.04G043800 | M-type_MADS |
| 610. | Glyma.04G044800.1.p | Glyma.04G044800 | LBD |
| 611. | Glyma.04G044900.1.p | Glyma.04G044900 | C2H2 |
| 612. | Glyma.04G045300.1.p | Glyma.04G045300 | bHLH |
| 613. | Glyma.04G047900.1.p | Glyma.04G047900 | AP2 |
| 614. | Glyma.04G048000.1.p | Glyma.04G048000 | bHLH |
| 615. | Glyma.04G048900.1.p | Glyma.04G048900 | MYB |
| 616. | Glyma.04G049300.1.p | Glyma.04G049300 | HD-ZIP |
| 617. | Glyma.04G049500.1.p | Glyma.04G049500 | TALE |
| 618. | Glyma.04G050300.1.p | Glyma.04G050300 | C3H |
| 619. | Glyma.04G050800.1.p | Glyma.04G050800 | TALE |
| 620. | Glyma.04G051000.1.p | Glyma.04G051000 | MYB_related |
| 621. | Glyma.04G051300.1.p | Glyma.04G051300 | GATA |
| 622. | Glyma.04G052000.1.p | Glyma.04G052000 | HSF |
| 623. | Glyma.04G054200.1.p | Glyma.04G054200 | WRKY |
| 624. | Glyma.04G054800.1.p | Glyma.04G054800 | Nin-like |
| 625. | Glyma.04G057700.1.p | Glyma.04G057700 | ERF |
| 626. | Glyma.04G058900.1.p | Glyma.04G058900 | CO-like |
| 627. | Glyma.04G061300.1.p | Glyma.04G061300 | WRKY |
| 628. | Glyma.04G061400.1.p | Glyma.04G061400 | WRKY |
| 629. | Glyma.04G062500.1.p | Glyma.04G062500 | ARR-B |
| 630. | Glyma.04G062600.1.p | Glyma.04G062600 | C2H2 |
| 631. | Glyma.04G062900.1.p | Glyma.04G062900 | ERF |
| 632. | Glyma.04G064100.1.p | Glyma.04G064100 | TALE |
| 633. | Glyma.04G067200.1.p | Glyma.04G067200 | ERF |
| 634. | Glyma.04G068000.1.p | Glyma.04G068000 | HB-other |
| 635. | Glyma.04G076200.1.p | Glyma.04G076200 | WRKY |
| 636. | Glyma.04G078300.1.p | Glyma.04G078300 | bZIP |
| 637. | Glyma.04G078600.1.p | Glyma.04G078600 | NAC |
| 638. | Glyma.04G080600.1.p | Glyma.04G080600 | MYB |
| 639. | Glyma.04G081700.1.p | Glyma.04G081700 | CPP |
| 640. | Glyma.04G084000.1.p | Glyma.04G084000 | ERF |
| 641. | Glyma.04G084900.1.p | Glyma.04G084900 | GATA |
| 642. | Glyma.04G085000.1.p | Glyma.04G085000 | HD-ZIP |
| 643. | Glyma.04G090100.1.p | Glyma.04G090100 | bHLH |
| 644. | Glyma.04G093300.1.p | Glyma.04G093300 | ZF-HD |
| 645. | Glyma.04G093900.1.p | Glyma.04G093900 | M-type_MADS |
| 646. | Glyma.04G094800.1.p | Glyma.04G094800 | YABBY |
| 647. | Glyma.04G095300.1.p | Glyma.04G095300 | MYB_related |
| 648. | Glyma.04G096200.1.p | Glyma.04G096200 | GATA |
| 649. | Glyma.04G096300.1.p | Glyma.04G096300 | GATA |
| 650. | Glyma.04G098400.1.p | Glyma.04G098400 | bHLH |
| 651. | Glyma.04G101900.1.p | Glyma.04G101900 | MYB |
| 652. | Glyma.04G103900.1.p | Glyma.04G103900 | ERF |
| 653. | Glyma.04G106400.1.p | Glyma.04G106400 | LBD |
| 654. | Glyma.04G110100.1.p | Glyma.04G110100 | MYB |
| 655. | Glyma.04G111200.1.p | Glyma.04G111200 | FAR1 |
| 656. | Glyma.04G115500.1.p | Glyma.04G115500 | WRKY |
| 657. | Glyma.04G119500.1.p | Glyma.04G119500 | NAC |
| 658. | Glyma.04G124200.1.p | Glyma.04G124200 | bZIP |
| 659. | Glyma.04G124300.1.p | Glyma.04G124300 | FAR1 |
| 660. | Glyma.04G125700.1.p | Glyma.04G125700 | MYB |
| 661. | Glyma.04G126200.1.p | Glyma.04G126200 | HB-other |
| 662. | Glyma.04G127600.1.p | Glyma.04G127600 | C2H2 |
| 663. | Glyma.04G130300.1.p | Glyma.04G130300 | C2H2 |
| 664. | Glyma.04G132200.1.p | Glyma.04G132200 | MYB_related |
| 665. | Glyma.04G135400.1.p | Glyma.04G135400 | Trihelix |
| 666. | Glyma.04G136700.1.p | Glyma.04G136700 | SRS |
| 667. | Glyma.04G138900.1.p | Glyma.04G138900 | GRAS |
| 668. | Glyma.04G141700.1.p | Glyma.04G141700 | MYB |
| 669. | Glyma.04G142500.1.p | Glyma.04G142500 | MIKC_MADS |
| 670. | Glyma.04G144700.1.p | Glyma.04G144700 | FAR1 |
| 671. | Glyma.04G145000.1.p | Glyma.04G145000 | NF-YB |
| 672. | Glyma.04G147500.1.p | Glyma.04G147500 | ERF |
| 673. | Glyma.04G148800.1.p | Glyma.04G148800 | MYB_related |
| 674. | Glyma.04G150500.1.p | Glyma.04G150500 | GRAS |
| 675. | Glyma.04G150800.1.p | Glyma.04G150800 | FAR1 |
| 676. | Glyma.04G151000.1.p | Glyma.04G151000 | G2-like |
| 677. | Glyma.04G151100.1.p | Glyma.04G151100 | ERF |
| 678. | Glyma.04G152400.1.p | Glyma.04G152400 | TCP |
| 679. | Glyma.04G154900.1.p | Glyma.04G154900 | AP2 |
| 680. | Glyma.04G158300.1.p | Glyma.04G158300 | bZIP |
| 681. | Glyma.04G158800.1.p | Glyma.04G158800 | Dof |
| 682. | Glyma.04G159300.1.p | Glyma.04G159300 | MIKC_MADS |
| 683. | Glyma.04G159600.1.p | Glyma.04G159600 | SBP |
| 684. | Glyma.04G161400.1.p | Glyma.04G161400 | TCP |
| 685. | Glyma.04G166900.1.p | Glyma.04G166900 | MYB |
| 686. | Glyma.04G167200.1.p | Glyma.04G167200 | NAC |
| 687. | Glyma.04G168300.1.p | Glyma.04G168300 | Dof |
| 688. | Glyma.04G169000.1.p | Glyma.04G169000 | Nin-like |
| 689. | Glyma.04G169900.1.p | Glyma.04G169900 | ZF-HD |
| 690. | Glyma.04G170100.1.p | Glyma.04G170100 | MYB |
| 691. | Glyma.04G170600.1.p | Glyma.04G170600 | TCP |
| 692. | Glyma.04G171600.1.p | Glyma.04G171600 | bHLH |
| 693. | Glyma.04G172500.1.p | Glyma.04G172500 | bHLH |
| 694. | Glyma.04G173500.1.p | Glyma.04G173500 | WRKY |
| 695. | Glyma.04G174500.1.p | Glyma.04G174500 | HB-other |
| 696. | Glyma.04G175800.1.p | Glyma.04G175800 | NAC |
| 697. | Glyma.04G176700.1.p | Glyma.04G176700 | MYB |
| 698. | Glyma.04G176900.1.p | Glyma.04G176900 | bHLH |
| 699. | Glyma.04G177300.1.p | Glyma.04G177300 | MYB_related |
| 700. | Glyma.04G177400.1.p | Glyma.04G177400 | FAR1 |
| 701. | Glyma.04G178300.1.p | Glyma.04G178300 | E2F/DP |
| 702. | Glyma.04G178700.1.p | Glyma.04G178700 | bHLH |
| 703. | Glyma.04G182600.1.p | Glyma.04G182600 | bHLH |
| 704. | Glyma.04G183700.1.p | Glyma.04G183700 | Dof |
| 705. | Glyma.04G184200.1.p | Glyma.04G184200 | MYB_related |
| 706. | Glyma.04G185100.1.p | Glyma.04G185100 | TALE |
| 707. | Glyma.04G186100.1.p | Glyma.04G186100 | FAR1 |
| 708. | Glyma.04G186900.1.p | Glyma.04G186900 | LBD |
| 709. | Glyma.04G187300.1.p | Glyma.04G187300 | MYB |
| 710. | Glyma.04G191900.1.p | Glyma.04G191900 | C2H2 |
| 711. | Glyma.04G192000.1.p | Glyma.04G192000 | C2H2 |
| 712. | Glyma.04G192100.1.p | Glyma.04G192100 | C3H |
| 713. | Glyma.04G192400.1.p | Glyma.04G192400 | B3 |
| 714. | Glyma.04G194600.1.p | Glyma.04G194600 | Trihelix |
| 715. | Glyma.04G196200.1.p | Glyma.04G196200 | NF-YC |
| 716. | Glyma.04G197100.1.p | Glyma.04G197100 | SBP |
| 717. | Glyma.04G199000.1.p | Glyma.04G199000 | NAC |
| 718. | Glyma.04G199900.1.p | Glyma.04G199900 | bHLH |
| 719. | Glyma.04G200500.1.p | Glyma.04G200500 | bHLH |
| 720. | Glyma.04G201700.1.p | Glyma.04G201700 | ERF |
| 721. | Glyma.04G201900.1.p | Glyma.04G201900 | ERF |
| 722. | Glyma.04G202000.1.p | Glyma.04G202000 | LFY |
| 723. | Glyma.04G205100.1.p | Glyma.04G205100 | MYB |
| 724. | Glyma.04G208300.1.p | Glyma.04G208300 | NAC |
| 725. | Glyma.04G211200.1.p | Glyma.04G211200 | NF-YB |
| 726. | Glyma.04G212000.1.p | Glyma.04G212000 | NAC |
| 727. | Glyma.04G213300.1.p | Glyma.04G213300 | NAC |
| 728. | Glyma.04G214100.1.p | Glyma.04G214100 | bHLH |
| 729. | Glyma.04G214200.1.p | Glyma.04G214200 | C3H |
| 730. | Glyma.04G216100.1.p | Glyma.04G216100 | Trihelix |
| 731. | Glyma.04G217400.1.p | Glyma.04G217400 | ERF |
| 732. | Glyma.04G218400.1.p | Glyma.04G218400 | WRKY |
| 733. | Glyma.04G218700.1.p | Glyma.04G218700 | WRKY |
| 734. | Glyma.04G220500.1.p | Glyma.04G220500 | LBD |
| 735. | Glyma.04G221800.1.p | Glyma.04G221800 | MYB_related |
| 736. | Glyma.04G222200.1.p | Glyma.04G222200 | bZIP |
| 737. | Glyma.04G222700.1.p | Glyma.04G222700 | C2H2 |
| 738. | Glyma.04G223200.1.p | Glyma.04G223200 | WRKY |
| 739. | Glyma.04G223300.1.p | Glyma.04G223300 | WRKY |
| 740. | Glyma.04G226700.1.p | Glyma.04G226700 | NAC |
| 741. | Glyma.04G230600.1.p | Glyma.04G230600 | GRF |
| 742. | Glyma.04G231400.1.p | Glyma.04G231400 | HD-ZIP |
| 743. | Glyma.04G233300.1.p | Glyma.04G233300 | Dof |
| 744. | Glyma.04G234600.1.p | Glyma.04G234600 | Nin-like |
| 745. | Glyma.04G238300.1.p | Glyma.04G238300 | WRKY |
| 746. | Glyma.04G238400.1.p | Glyma.04G238400 | bHLH |
| 747. | Glyma.04G238700.1.p | Glyma.04G238700 | ERF |
| 748. | Glyma.04G239500.1.p | Glyma.04G239500 | Dof |
| 749. | Glyma.04G242100.1.p | Glyma.04G242100 | GRAS |
| 750. | Glyma.04G242200.1.p | Glyma.04G242200 | MYB |
| 751. | Glyma.04G245500.1.p | Glyma.04G245500 | MIKC_MADS |
| 752. | Glyma.04G249000.1.p | Glyma.04G249000 | NAC |
| 753. | Glyma.04G251400.1.p | Glyma.04G251400 | ERF |
| 754. | Glyma.04G251900.1.p | Glyma.04G251900 | GRAS |
| 755. | Glyma.04G254200.1.p | Glyma.04G254200 | ARF |
| 756. | Glyma.04G254800.1.p | Glyma.04G254800 | bZIP |
| 757. | Glyma.04G257000.1.p | Glyma.04G257000 | B3 |
| 758. | Glyma.04G257100.1.p | Glyma.04G257100 | MIKC_MADS |

| 759. | Glyma.05G002700.1.p | Glyma.05G002700 | NAC |
| --- | --- | --- | --- |
| 760. | Glyma.05G004400.1.p | Glyma.05G004400 | LBD |
| 761. | Glyma.05G005400.1.p | Glyma.05G005400 | MYB |
| 762. | Glyma.05G006100.1.p | Glyma.05G006100 | MYB |
| 763. | Glyma.05G008000.1.p | Glyma.05G008000 | C3H |
| 764. | Glyma.05G008400.1.p | Glyma.05G008400 | C3H |
| 765. | Glyma.05G011500.1.p | Glyma.05G011500 | G2-like |
| 766. | Glyma.05G013000.1.p | Glyma.05G013000 | MYB |
| 767. | Glyma.05G013300.1.p | Glyma.05G013300 | TCP |
| 768. | Glyma.05G015100.1.p | Glyma.05G015100 | NF-YB |
| 769. | Glyma.05G015900.1.p | Glyma.05G015900 | ERF |
| 770. | Glyma.05G017400.1.p | Glyma.05G017400 | bZIP |
| 771. | Glyma.05G017800.1.p | Glyma.05G017800 | bHLH |
| 772. | Glyma.05G018100.1.p | Glyma.05G018100 | Dof |
| 773. | Glyma.05G018800.1.p | Glyma.05G018800 | MIKC_MADS |
| 774. | Glyma.05G018900.1.p | Glyma.05G018900 | MIKC_MADS |
| 775. | Glyma.05G019000.1.p | Glyma.05G019000 | SBP |
| 776. | Glyma.05G019900.1.p | Glyma.05G019900 | TCP |
| 777. | Glyma.05G022700.1.p | Glyma.05G022700 | C2H2 |
| 778. | Glyma.05G025500.1.p | Glyma.05G025500 | NAC |
| 779. | Glyma.05G025700.1.p | Glyma.05G025700 | M-type_MADS |
| 780. | Glyma.05G025900.1.p | Glyma.05G025900 | Dof |
| 781. | Glyma.05G026500.1.p | Glyma.05G026500 | Nin-like |
| 782. | Glyma.05G026900.1.p | Glyma.05G026900 | ZF-HD |
| 783. | Glyma.05G027000.1.p | Glyma.05G027000 | MYB |
| 784. | Glyma.05G027400.1.p | Glyma.05G027400 | TCP |
| 785. | Glyma.05G029000.1.p | Glyma.05G029000 | WRKY |
| 786. | Glyma.05G030000.1.p | Glyma.05G030000 | HD-ZIP |
| 787. | Glyma.05G031800.1.p | Glyma.05G031800 | bHLH |
| 788. | Glyma.05G032200.1.p | Glyma.05G032200 | MYB_related |
| 789. | Glyma.05G033400.1.p | Glyma.05G033400 | E2F/DP |
| 790. | Glyma.05G036800.1.p | Glyma.05G036800 | bHLH |
| 791. | Glyma.05G037800.1.p | Glyma.05G037800 | Dof |
| 792. | Glyma.05G038500.1.p | Glyma.05G038500 | MYB |
| 793. | Glyma.05G040500.1.p | Glyma.05G040500 | LBD |
| 794. | Glyma.05G040700.1.p | Glyma.05G040700 | MYB |
| 795. | Glyma.05G044300.1.p | Glyma.05G044300 | C3H |
| 796. | Glyma.05G045300.1.p | Glyma.05G045300 | GRAS |
| 797. | Glyma.05G049300.1.p | Glyma.05G049300 | GRAS |
| 798. | Glyma.05G049800.1.p | Glyma.05G049800 | ERF |
| 799. | Glyma.05G049900.1.p | Glyma.05G049900 | ERF |
| 800. | Glyma.05G050400.1.p | Glyma.05G050400 | TCP |
| 801. | Glyma.05G050600.1.p | Glyma.05G050600 | TALE |
| 802. | Glyma.05G050700.1.p | Glyma.05G050700 | MIKC_MADS |
| 803. | Glyma.05G051700.1.p | Glyma.05G051700 | MYB |
| 804. | Glyma.05G055900.1.p | Glyma.05G055900 | NAC |
| 805. | Glyma.05G056000.1.p | Glyma.05G056000 | YABBY |
| 806. | Glyma.05G059800.1.p | Glyma.05G059800 | C2H2 |
| 807. | Glyma.05G061900.1.p | Glyma.05G061900 | MYB |
| 808. | Glyma.05G062100.1.p | Glyma.05G062100 | ERF |
| 809. | Glyma.05G062300.1.p | Glyma.05G062300 | MYB_related |
| 810. | Glyma.05G062700.1.p | Glyma.05G062700 | HD-ZIP |
| 811. | Glyma.05G063500.1.p | Glyma.05G063500 | ERF |
| 812. | Glyma.05G063600.1.p | Glyma.05G063600 | ERF |
| 813. | Glyma.05G064800.1.p | Glyma.05G064800 | GATA |
| 814. | Glyma.05G070200.1.p | Glyma.05G070200 | ARR-B |
| 815. | Glyma.05G072100.1.p | Glyma.05G072100 | FAR1 |
| 816. | Glyma.05G072600.1.p | Glyma.05G072600 | MYB |
| 817. | Glyma.05G079800.1.p | Glyma.05G079800 | bZIP |
| 818. | Glyma.05G081400.1.p | Glyma.05G081400 | LBD |
| 819. | Glyma.05G081700.1.p | Glyma.05G081700 | G2-like |
| 820. | Glyma.05G085900.1.p | Glyma.05G085900 | NF-YB |
| 821. | Glyma.05G086000.1.p | Glyma.05G086000 | NAC |
| 822. | Glyma.05G088300.1.p | Glyma.05G088300 | GeBP |
| 823. | Glyma.05G091200.1.p | Glyma.05G091200 | AP2 |
| 824. | Glyma.05G092800.1.p | Glyma.05G092800 | ERF |
| 825. | Glyma.05G094900.1.p | Glyma.05G094900 | bHLH |
| 826. | Glyma.05G095900.1.p | Glyma.05G095900 | HSF |
| 827. | Glyma.05G096500.1.p | Glyma.05G096500 | WRKY |
| 828. | Glyma.05G098200.1.p | Glyma.05G098200 | MYB |
| 829. | Glyma.05G103300.1.p | Glyma.05G103300 | LBD |
| 830. | Glyma.05G103400.1.p | Glyma.05G103400 | GRAS |
| 831. | Glyma.05G105600.1.p | Glyma.05G105600 | GRAS |
| 832. | Glyma.05G106000.1.p | Glyma.05G106000 | MYB_related |
| 833. | Glyma.05G108200.1.p | Glyma.05G108200 | bZIP |
| 834. | Glyma.05G108600.1.p | Glyma.05G108600 | AP2 |
| 835. | Glyma.05G108700.1.p | Glyma.05G108700 | NAC |
| 836. | Glyma.05G109200.1.p | Glyma.05G109200 | MYB |
| 837. | Glyma.05G109500.1.p | Glyma.05G109500 | HD-ZIP |
| 838. | Glyma.05G110600.1.p | Glyma.05G110600 | bHLH |
| 839. | Glyma.05G110700.1.p | Glyma.05G110700 | bHLH |
| 840. | Glyma.05G110900.1.p | Glyma.05G110900 | bHLH |
| 841. | Glyma.05G111400.1.p | Glyma.05G111400 | C2H2 |
| 842. | Glyma.05G112200.1.p | Glyma.05G112200 | LSD |
| 843. | Glyma.05G113000.1.p | Glyma.05G113000 | NAC |
| 844. | Glyma.05G115400.1.p | Glyma.05G115400 | ARR-B |
| 845. | Glyma.05G117000.1.p | Glyma.05G117000 | CAMTA |
| 846. | Glyma.05G120500.1.p | Glyma.05G120500 | NAC |
| 847. | Glyma.05G122400.1.p | Glyma.05G122400 | bZIP |
| 848. | Glyma.05G123000.1.p | Glyma.05G123000 | WRKY |
| 849. | Glyma.05G123600.1.p | Glyma.05G123600 | WRKY |
| 850. | Glyma.05G127600.1.p | Glyma.05G127600 | WRKY |
| 851. | Glyma.05G134400.1.p | Glyma.05G134400 | bHLH |
| 852. | Glyma.05G137000.1.p | Glyma.05G137000 | C2H2 |
| 853. | Glyma.05G137100.1.p | Glyma.05G137100 | BBR-BPC |
| 854. | Glyma.05G139000.1.p | Glyma.05G139000 | C2H2 |
| 855. | Glyma.05G140000.1.p | Glyma.05G140000 | M-type_MADS |
| 856. | Glyma.05G140400.1.p | Glyma.05G140400 | GRAS |
| 857. | Glyma.05G142000.1.p | Glyma.05G142000 | TCP |
| 858. | Glyma.05G143800.1.p | Glyma.05G143800 | ARF |
| 859. | Glyma.05G144500.1.p | Glyma.05G144500 | ARR-B |
| 860. | Glyma.05G148300.1.p | Glyma.05G148300 | CAMTA |
| 861. | Glyma.05G148700.1.p | Glyma.05G148700 | MIKC_MADS |
| 862. | Glyma.05G148800.1.p | Glyma.05G148800 | MIKC_MADS |
| 863. | Glyma.05G151800.1.p | Glyma.05G151800 | HSF |
| 864. | Glyma.05G157000.1.p | Glyma.05G157000 | bZIP |
| 865. | Glyma.05G157400.1.p | Glyma.05G157400 | ERF |
| 866. | Glyma.05G158200.1.p | Glyma.05G158200 | Dof |
| 867. | Glyma.05G158900.1.p | Glyma.05G158900 | G2-like |
| 868. | Glyma.05G160800.1.p | Glyma.05G160800 | WRKY |
| 869. | Glyma.05G162300.1.p | Glyma.05G162300 | HSF |
| 870. | Glyma.05G163200.1.p | Glyma.05G163200 | MIKC_MADS |
| 871. | Glyma.05G165800.1.p | Glyma.05G165800 | WRKY |
| 872. | Glyma.05G166100.1.p | Glyma.05G166100 | NF-YA |
| 873. | Glyma.05G166400.1.p | Glyma.05G166400 | HD-ZIP |
| 874. | Glyma.05G167900.1.p | Glyma.05G167900 | C2H2 |
| 875. | Glyma.05G168100.1.p | Glyma.05G168100 | bZIP |
| 876. | Glyma.05G170000.1.p | Glyma.05G170000 | GATA |
| 877. | Glyma.05G170100.1.p | Glyma.05G170100 | GATA |
| 878. | Glyma.05G171000.1.p | Glyma.05G171000 | LSD |
| 879. | Glyma.05G175600.1.p | Glyma.05G175600 | HD-ZIP |
| 880. | Glyma.05G177700.1.p | Glyma.05G177700 | C2H2 |
| 881. | Glyma.05G178200.1.p | Glyma.05G178200 | CAMTA |
| 882. | Glyma.05G179900.1.p | Glyma.05G179900 | ERF |
| 883. | Glyma.05G180300.1.p | Glyma.05G180300 | EIL |
| 884. | Glyma.05G182500.1.p | Glyma.05G182500 | bZIP |
| 885. | Glyma.05G183200.1.p | Glyma.05G183200 | NF-YB |
| 886. | Glyma.05G184500.1.p | Glyma.05G184500 | WRKY |
| 887. | Glyma.05G185400.1.p | Glyma.05G185400 | WRKY |
| 888. | Glyma.05G186700.1.p | Glyma.05G186700 | ERF |
| 889. | Glyma.05G189200.1.p | Glyma.05G189200 | MYB_related |
| 890. | Glyma.05G189700.1.p | Glyma.05G189700 | LBD |
| 891. | Glyma.05G190500.1.p | Glyma.05G190500 | C3H |
| 892. | Glyma.05G190600.1.p | Glyma.05G190600 | bHLH |
| 893. | Glyma.05G191300.1.p | Glyma.05G191300 | NAC |
| 894. | Glyma.05G192500.1.p | Glyma.05G192500 | NAC |
| 895. | Glyma.05G193300.1.p | Glyma.05G193300 | NF-YB |
| 896. | Glyma.05G195000.1.p | Glyma.05G195000 | NAC |
| 897. | Glyma.05G200100.1.p | Glyma.05G200100 | ERF |
| 898. | Glyma.05G200400.1.p | Glyma.05G200400 | Trihelix |
| 899. | Glyma.05G200800.1.p | Glyma.05G200800 | ARF |
| 900. | Glyma.05G200900.1.p | Glyma.05G200900 | bHLH |
| 901. | Glyma.05G201700.1.p | Glyma.05G201700 | bHLH |
| 902. | Glyma.05G201900.1.p | Glyma.05G201900 | BBR-BPC |
| 903. | Glyma.05G202300.1.p | Glyma.05G202300 | NAC |
| 904. | Glyma.05G203900.1.p | Glyma.05G203900 | WRKY |
| 905. | Glyma.05G204100.1.p | Glyma.05G204100 | SBP |
| 906. | Glyma.05G208300.1.p | Glyma.05G208300 | bHLH |
| 907. | Glyma.05G210300.1.p | Glyma.05G210300 | TALE |
| 908. | Glyma.05G210600.1.p | Glyma.05G210600 | C3H |
| 909. | Glyma.05G211200.1.p | Glyma.05G211200 | MYB |
| 910. | Glyma.05G211900.1.p | Glyma.05G211900 | WRKY |
| 911. | Glyma.05G214400.1.p | Glyma.05G214400 | ERF |
| 912. | Glyma.05G215900.1.p | Glyma.05G215900 | WRKY |
| 913. | Glyma.05G221300.1.p | Glyma.05G221300 | ARF |
| 914. | Glyma.05G222600.1.p | Glyma.05G222600 | MYB_related |
| 915. | Glyma.05G224300.1.p | Glyma.05G224300 | MYB |
| 916. | Glyma.05G224400.1.p | Glyma.05G224400 | C3H |
| 917. | Glyma.05G225100.1.p | Glyma.05G225100 | NAC |
| 918. | Glyma.05G227200.1.p | Glyma.05G227200 | M-type_MADS |
| 919. | Glyma.05G227300.1.p | Glyma.05G227300 | M-type_MADS |
| 920. | Glyma.05G228100.1.p | Glyma.05G228100 | ERF |
| 921. | Glyma.05G229500.1.p | Glyma.05G229500 | bHLH |
| 922. | Glyma.05G229900.1.p | Glyma.05G229900 | CPP |
| 923. | Glyma.05G233700.1.p | Glyma.05G233700 | CO-like |
| 924. | Glyma.05G234200.1.p | Glyma.05G234200 | NAC |
| 925. | Glyma.05G234500.1.p | Glyma.05G234500 | bHLH |
| 926. | Glyma.05G234600.1.p | Glyma.05G234600 | MYB |
| 927. | Glyma.05G235400.1.p | Glyma.05G235400 | C2H2 |
| 928. | Glyma.05G239500.1.p | Glyma.05G239500 | G2-like |
| 929. | Glyma.05G239800.1.p | Glyma.05G239800 | ARR-B |
| 930. | Glyma.05G240500.1.p | Glyma.05G240500 | HSF |
| 931. | Glyma.05G245900.1.p | Glyma.05G245900 | WOX |
| 932. | Glyma.05G248200.1.p | Glyma.05G248200 | ZF-HD |
| 933. | Glyma.05G248300.1.p | Glyma.05G248300 | C2H2 |
| 934. | Glyma.05G248800.1.p | Glyma.05G248800 | HD-ZIP |

| 935. | Glyma.06G000400.1.p | Glyma.06G000400 | Nin-like |
| --- | --- | --- | --- |
| 936. | Glyma.06G002400.1.p | Glyma.06G002400 | FAR1 |
| 937. | Glyma.06G003800.1.p | Glyma.06G003800 | MYB |
| 938. | Glyma.06G008800.1.p | Glyma.06G008800 | GATA |
| 939. | Glyma.06G009100.1.p | Glyma.06G009100 | DBB |
| 940. | Glyma.06G009200.1.p | Glyma.06G009200 | SRS |
| 941. | Glyma.06G009500.1.p | Glyma.06G009500 | TALE |
| 942. | Glyma.06G010200.1.p | Glyma.06G010200 | bZIP |
| 943. | Glyma.06G012200.1.p | Glyma.06G012200 | bHLH |
| 944. | Glyma.06G014900.1.p | Glyma.06G014900 | NAC |
| 945. | Glyma.06G016800.1.p | Glyma.06G016800 | WOX |
| 946. | Glyma.06G017800.1.p | Glyma.06G017800 | Nin-like |
| 947. | Glyma.06G022300.1.p | Glyma.06G022300 | bZIP |
| 948. | Glyma.06G024900.1.p | Glyma.06G024900 | E2F/DP |
| 949. | Glyma.06G026100.1.p | Glyma.06G026100 | BBR-BPC |
| 950. | Glyma.06G027000.1.p | Glyma.06G027000 | DBB |
| 951. | Glyma.06G027200.1.p | Glyma.06G027200 | MIKC_MADS |
| 952. | Glyma.06G027500.1.p | Glyma.06G027500 | SRS |
| 953. | Glyma.06G028300.1.p | Glyma.06G028300 | ERF |
| 954. | Glyma.06G029100.1.p | Glyma.06G029100 | TALE |
| 955. | Glyma.06G029200.1.p | Glyma.06G029200 | TALE |
| 956. | Glyma.06G029600.1.p | Glyma.06G029600 | bZIP |
| 957. | Glyma.06G031400.1.p | Glyma.06G031400 | MYB_related |
| 958. | Glyma.06G031800.1.p | Glyma.06G031800 | MYB_related |
| 959. | Glyma.06G033300.1.p | Glyma.06G033300 | C2H2 |
| 960. | Glyma.06G034000.1.p | Glyma.06G034000 | BES1 |
| 961. | Glyma.06G035700.1.p | Glyma.06G035700 | G2-like |
| 962. | Glyma.06G036800.1.p | Glyma.06G036800 | MYB |
| 963. | Glyma.06G038200.1.p | Glyma.06G038200 | NF-YC |
| 964. | Glyma.06G038900.1.p | Glyma.06G038900 | B3 |
| 965. | Glyma.06G040400.1.p | Glyma.06G040400 | bZIP |
| 966. | Glyma.06G040800.1.p | Glyma.06G040800 | bHLH |
| 967. | Glyma.06G040900.1.p | Glyma.06G040900 | HSF |
| 968. | Glyma.06G041800.1.p | Glyma.06G041800 | WOX |
| 969. | Glyma.06G042100.1.p | Glyma.06G042100 | ERF |
| 970. | Glyma.06G045400.1.p | Glyma.06G045400 | C2H2 |
| 971. | Glyma.06G045800.1.p | Glyma.06G045800 | bHLH |
| 972. | Glyma.06G048500.1.p | Glyma.06G048500 | bZIP |
| 973. | Glyma.06G049200.1.p | Glyma.06G049200 | AP2 |
| 974. | Glyma.06G049300.1.p | Glyma.06G049300 | bHLH |
| 975. | Glyma.06G050000.1.p | Glyma.06G050000 | MYB |
| 976. | Glyma.06G050300.1.p | Glyma.06G050300 | C3H |
| 977. | Glyma.06G051600.1.p | Glyma.06G051600 | TALE |
| 978. | Glyma.06G051900.1.p | Glyma.06G051900 | MYB_related |
| 979. | Glyma.06G054500.1.p | Glyma.06G054500 | WRKY |
| 980. | Glyma.06G054900.1.p | Glyma.06G054900 | Nin-like |
| 981. | Glyma.06G058400.1.p | Glyma.06G058400 | ERF |
| 982. | Glyma.06G059600.1.p | Glyma.06G059600 | CO-like |
| 983. | Glyma.06G061900.1.p | Glyma.06G061900 | WRKY |
| 984. | Glyma.06G063500.1.p | Glyma.06G063500 | ARR-B |
| 985. | Glyma.06G063700.1.p | Glyma.06G063700 | C2H2 |
| 986. | Glyma.06G064000.1.p | Glyma.06G064000 | ERF |
| 987. | Glyma.06G065200.1.p | Glyma.06G065200 | TALE |
| 988. | Glyma.06G068800.1.p | Glyma.06G068800 | ERF |
| 989. | Glyma.06G069700.1.p | Glyma.06G069700 | HB-other |
| 990. | Glyma.06G077400.1.p | Glyma.06G077400 | WRKY |
| 991. | Glyma.06G079800.1.p | Glyma.06G079800 | bZIP |
| 992. | Glyma.06G080200.1.p | Glyma.06G080200 | NAC |
| 993. | Glyma.06G082300.1.p | Glyma.06G082300 | MYB |
| 994. | Glyma.06G083400.1.p | Glyma.06G083400 | CPP |
| 995. | Glyma.06G085700.1.p | Glyma.06G085700 | ERF |
| 996. | Glyma.06G086400.1.p | Glyma.06G086400 | GATA |
| 997. | Glyma.06G086600.1.p | Glyma.06G086600 | HD-ZIP |
| 998. | Glyma.06G092000.1.p | Glyma.06G092000 | bHLH |
| 999. | Glyma.06G095200.1.p | Glyma.06G095200 | ZF-HD |
| 1000. | Glyma.06G095700.1.p | Glyma.06G095700 | MIKC_MADS |
| 1001. | Glyma.06G096500.1.p | Glyma.06G096500 | YABBY |
| 1002. | Glyma.06G097100.1.p | Glyma.06G097100 | MYB_related |
| 1003. | Glyma.06G097900.1.p | Glyma.06G097900 | GATA |
| 1004. | Glyma.06G098000.1.p | Glyma.06G098000 | GATA |
| 1005. | Glyma.06G100000.1.p | Glyma.06G100000 | bHLH |
| 1006. | Glyma.06G103300.1.p | Glyma.06G103300 | MYB |
| 1007. | Glyma.06G105000.1.p | Glyma.06G105000 | ERF |
| 1008. | Glyma.06G106500.1.p | Glyma.06G106500 | LBD |
| 1009. | Glyma.06G107300.1.p | Glyma.06G107300 | bZIP |
| 1010. | Glyma.06G110800.1.p | Glyma.06G110800 | GRAS |
| 1011. | Glyma.06G111300.1.p | Glyma.06G111300 | ERF |
| 1012. | Glyma.06G114000.1.p | Glyma.06G114000 | NAC |
| 1013. | Glyma.06G117600.1.p | Glyma.06G117600 | MIKC_MADS |
| 1014. | Glyma.06G121200.1.p | Glyma.06G121200 | MYB |
| 1015. | Glyma.06G121300.1.p | Glyma.06G121300 | GRAS |
| 1016. | Glyma.06G124300.1.p | Glyma.06G124300 | Dof |
| 1017. | Glyma.06G125100.1.p | Glyma.06G125100 | ERF |
| 1018. | Glyma.06G125500.1.p | Glyma.06G125500 | bHLH |
| 1019. | Glyma.06G125600.1.p | Glyma.06G125600 | WRKY |
| 1020. | Glyma.06G129900.1.p | Glyma.06G129900 | Nin-like |
| 1021. | Glyma.06G130000.1.p | Glyma.06G130000 | Nin-like |
| 1022. | Glyma.06G131500.1.p | Glyma.06G131500 | Dof |
| 1023. | Glyma.06G133800.1.p | Glyma.06G133800 | HD-ZIP |
| 1024. | Glyma.06G134600.1.p | Glyma.06G134600 | GRF |
| 1025. | Glyma.06G138100.1.p | Glyma.06G138100 | NAC |
| 1026. | Glyma.06G142000.1.p | Glyma.06G142000 | WRKY |
| 1027. | Glyma.06G142100.1.p | Glyma.06G142100 | WRKY |
| 1028. | Glyma.06G142600.1.p | Glyma.06G142600 | C2H2 |
| 1029. | Glyma.06G143600.1.p | Glyma.06G143600 | MYB_related |
| 1030. | Glyma.06G145400.1.p | Glyma.06G145400 | LBD |
| 1031. | Glyma.06G147100.1.p | Glyma.06G147100 | WRKY |
| 1032. | Glyma.06G147500.1.p | Glyma.06G147500 | WRKY |
| 1033. | Glyma.06G148400.1.p | Glyma.06G148400 | ERF |
| 1034. | Glyma.06G149900.1.p | Glyma.06G149900 | Trihelix |
| 1035. | Glyma.06G152200.1.p | Glyma.06G152200 | C3H |
| 1036. | Glyma.06G152300.1.p | Glyma.06G152300 | bHLH |
| 1037. | Glyma.06G152900.1.p | Glyma.06G152900 | NAC |
| 1038. | Glyma.06G154400.1.p | Glyma.06G154400 | NAC |
| 1039. | Glyma.06G157400.1.p | Glyma.06G157400 | NAC |
| 1040. | Glyma.06G160500.1.p | Glyma.06G160500 | MYB |
| 1041. | Glyma.06G163600.1.p | Glyma.06G163600 | LFY |
| 1042. | Glyma.06G163700.1.p | Glyma.06G163700 | ERF |
| 1043. | Glyma.06G164900.1.p | Glyma.06G164900 | ARF |
| 1044. | Glyma.06G165000.1.p | Glyma.06G165000 | bHLH |
| 1045. | Glyma.06G165700.1.p | Glyma.06G165700 | bHLH |
| 1046. | Glyma.06G166500.1.p | Glyma.06G166500 | NAC |
| 1047. | Glyma.06G168400.1.p | Glyma.06G168400 | WRKY |
| 1048. | Glyma.06G168600.1.p | Glyma.06G168600 | SBP |
| 1049. | Glyma.06G169600.1.p | Glyma.06G169600 | NF-YC |
| 1050. | Glyma.06G171400.1.p | Glyma.06G171400 | Trihelix |
| 1051. | Glyma.06G173700.1.p | Glyma.06G173700 | C3H |
| 1052. | Glyma.06G173800.1.p | Glyma.06G173800 | C2H2 |
| 1053. | Glyma.06G174000.1.p | Glyma.06G174000 | C2H2 |
| 1054. | Glyma.06G178600.1.p | Glyma.06G178600 | MYB |
| 1055. | Glyma.06G178900.1.p | Glyma.06G178900 | LBD |
| 1056. | Glyma.06G181800.1.p | Glyma.06G181800 | MYB |
| 1057. | Glyma.06G182200.1.p | Glyma.06G182200 | Dof |
| 1058. | Glyma.06G186800.1.p | Glyma.06G186800 | E2F/DP |
| 1059. | Glyma.06G187600.1.p | Glyma.06G187600 | MYB_related |
| 1060. | Glyma.06G188100.1.p | Glyma.06G188100 | bHLH |
| 1061. | Glyma.06G188400.1.p | Glyma.06G188400 | MYB_related |
| 1062. | Glyma.06G190200.1.p | Glyma.06G190200 | HD-ZIP |
| 1063. | Glyma.06G190800.1.p | Glyma.06G190800 | WRKY |
| 1064. | Glyma.06G191800.1.p | Glyma.06G191800 | bHLH |
| 1065. | Glyma.06G192300.1.p | Glyma.06G192300 | bHLH |
| 1066. | Glyma.06G193000.1.p | Glyma.06G193000 | TCP |
| 1067. | Glyma.06G193600.1.p | Glyma.06G193600 | MYB |
| 1068. | Glyma.06G193700.1.p | Glyma.06G193700 | ZF-HD |
| 1069. | Glyma.06G194300.1.p | Glyma.06G194300 | Nin-like |
| 1070. | Glyma.06G194800.1.p | Glyma.06G194800 | Dof |
| 1071. | Glyma.06G195500.1.p | Glyma.06G195500 | NAC |
| 1072. | Glyma.06G195600.1.p | Glyma.06G195600 | MYB |
| 1073. | Glyma.06G197700.1.p | Glyma.06G197700 | C2H2 |
| 1074. | Glyma.06G198300.1.p | Glyma.06G198300 | Nin-like |
| 1075. | Glyma.06G199600.1.p | Glyma.06G199600 | bHLH |
| 1076. | Glyma.06G204300.1.p | Glyma.06G204300 | TCP |
| 1077. | Glyma.06G205700.1.p | Glyma.06G205700 | SBP |
| 1078. | Glyma.06G205800.1.p | Glyma.06G205800 | MIKC_MADS |
| 1079. | Glyma.06G206300.1.p | Glyma.06G206300 | bHLH |
| 1080. | Glyma.06G206400.1.p | Glyma.06G206400 | Dof |
| 1081. | Glyma.06G209000.1.p | Glyma.06G209000 | NF-YB |
| 1082. | Glyma.06G210600.1.p | Glyma.06G210600 | TCP |
| 1083. | Glyma.06G212900.1.p | Glyma.06G212900 | WRKY |
| 1084. | Glyma.06G213100.1.p | Glyma.06G213100 | GRAS |
| 1085. | Glyma.06G213400.1.p | Glyma.06G213400 | G2-like |
| 1086. | Glyma.06G216400.1.p | Glyma.06G216400 | MYB_related |
| 1087. | Glyma.06G219800.1.p | Glyma.06G219800 | WRKY |
| 1088. | Glyma.06G220000.1.p | Glyma.06G220000 | B3 |
| 1089. | Glyma.06G220100.1.p | Glyma.06G220100 | MYB_related |
| 1090. | Glyma.06G221200.1.p | Glyma.06G221200 | bHLH |
| 1091. | Glyma.06G221800.1.p | Glyma.06G221800 | ERF |
| 1092. | Glyma.06G222900.1.p | Glyma.06G222900 | FAR1 |
| 1093. | Glyma.06G225200.1.p | Glyma.06G225200 | AP2 |
| 1094. | Glyma.06G232300.1.p | Glyma.06G232300 | TCP |
| 1095. | Glyma.06G234700.1.p | Glyma.06G234700 | HD-ZIP |
| 1096. | Glyma.06G235200.1.p | Glyma.06G235200 | HB-other |
| 1097. | Glyma.06G236000.1.p | Glyma.06G236000 | NAC |
| 1098. | Glyma.06G236400.1.p | Glyma.06G236400 | ERF |
| 1099. | Glyma.06G238100.1.p | Glyma.06G238100 | SBP |
| 1100. | Glyma.06G242200.1.p | Glyma.06G242200 | WRKY |
| 1101. | Glyma.06G247000.1.p | Glyma.06G247000 | FAR1 |
| 1102. | Glyma.06G248200.1.p | Glyma.06G248200 | MYB |
| 1103. | Glyma.06G248900.1.p | Glyma.06G248900 | NAC |
| 1104. | Glyma.06G249100.1.p | Glyma.06G249100 | NAC |
| 1105. | Glyma.06G251200.1.p | Glyma.06G251200 | bHLH |
| 1106. | Glyma.06G260000.1.p | Glyma.06G260000 | bHLH |
| 1107. | Glyma.06G265500.1.p | Glyma.06G265500 | GRAS |
| 1108. | Glyma.06G266800.1.p | Glyma.06G266800 | bHLH |
| 1109. | Glyma.06G283300.1.p | Glyma.06G283300 | bHLH |
| 1110. | Glyma.06G284500.1.p | Glyma.06G284500 | TCP |
| 1111. | Glyma.06G284900.1.p | Glyma.06G284900 | bZIP |
| 1112. | Glyma.06G285100.1.p | Glyma.06G285100 | LBD |
| 1113. | Glyma.06G285800.1.p | Glyma.06G285800 | VOZ |
| 1114. | Glyma.06G287200.1.p | Glyma.06G287200 | C2H2 |
| 1115. | Glyma.06G288500.1.p | Glyma.06G288500 | NAC |
| 1116. | Glyma.06G288900.1.p | Glyma.06G288900 | HB-other |
| 1117. | Glyma.06G289300.1.p | Glyma.06G289300 | G2-like |
| 1118. | Glyma.06G290000.1.p | Glyma.06G290000 | ERF |
| 1119. | Glyma.06G290100.1.p | Glyma.06G290100 | C3H |
| 1120. | Glyma.06G291900.1.p | Glyma.06G291900 | bHLH |
| 1121. | Glyma.06G295300.1.p | Glyma.06G295300 | ERF |
| 1122. | Glyma.06G299300.1.p | Glyma.06G299300 | MYB |
| 1123. | Glyma.06G299900.1.p | Glyma.06G299900 | MYB |
| 1124. | Glyma.06G300000.1.p | Glyma.06G300000 | MYB |
| 1125. | Glyma.06G300100.1.p | Glyma.06G300100 | MYB |
| 1126. | Glyma.06G300200.1.p | Glyma.06G300200 | MYB |
| 1127. | Glyma.06G300300.1.p | Glyma.06G300300 | MYB |
| 1128. | Glyma.06G300400.1.p | Glyma.06G300400 | MYB |
| 1129. | Glyma.06G301300.1.p | Glyma.06G301300 | ERF |
| 1130. | Glyma.06G303100.1.p | Glyma.06G303100 | MYB_related |
| 1131. | Glyma.06G303700.1.p | Glyma.06G303700 | HD-ZIP |
| 1132. | Glyma.06G307700.1.p | Glyma.06G307700 | WRKY |
| 1133. | Glyma.06G308900.1.p | Glyma.06G308900 | YABBY |
| 1134. | Glyma.06G311400.1.p | Glyma.06G311400 | NF-YC |
| 1135. | Glyma.06G311500.1.p | Glyma.06G311500 | C2H2 |
| 1136. | Glyma.06G312900.1.p | Glyma.06G312900 | MYB |
| 1137. | Glyma.06G314000.1.p | Glyma.06G314000 | EIL |
| 1138. | Glyma.06G314300.1.p | Glyma.06G314300 | FAR1 |
| 1139. | Glyma.06G314400.1.p | Glyma.06G314400 | bZIP |
| 1140. | Glyma.06G316400.1.p | Glyma.06G316400 | C2H2 |
| 1141. | Glyma.06G318900.1.p | Glyma.06G318900 | NAC |
| 1142. | Glyma.06G320700.1.p | Glyma.06G320700 | WRKY |
| 1143. | Glyma.06G323400.1.p | Glyma.06G323400 | FAR1 |
| 1144. | Glyma.06G324400.1.p | Glyma.06G324400 | MIKC_MADS |

| 1145. | Glyma.07G007200.1.p | Glyma.07G007200 | MYB_related |
| --- | --- | --- | --- |
| 1146. | Glyma.07G008500.1.p | Glyma.07G008500 | MYB |
| 1147. | Glyma.07G009100.1.p | Glyma.07G009100 | C2H2 |
| 1148. | Glyma.07G009500.1.p | Glyma.07G009500 | LBD |
| 1149. | Glyma.07G012100.1.p | Glyma.07G012100 | Dof |
| 1150. | Glyma.07G012200.1.p | Glyma.07G012200 | C2H2 |
| 1151. | Glyma.07G013600.1.p | Glyma.07G013600 | bHLH |
| 1152. | Glyma.07G016000.1.p | Glyma.07G016000 | HB-other |
| 1153. | Glyma.07G016500.1.p | Glyma.07G016500 | HD-ZIP |
| 1154. | Glyma.07G016700.1.p | Glyma.07G016700 | HD-ZIP |
| 1155. | Glyma.07G016800.1.p | Glyma.07G016800 | GATA |
| 1156. | Glyma.07G017300.1.p | Glyma.07G017300 | ERF |
| 1157. | Glyma.07G018500.1.p | Glyma.07G018500 | bHLH |
| 1158. | Glyma.07G019500.1.p | Glyma.07G019500 | HD-ZIP |
| 1159. | Glyma.07G021000.1.p | Glyma.07G021000 | AP2 |
| 1160. | Glyma.07G023300.1.p | Glyma.07G023300 | WRKY |
| 1161. | Glyma.07G025300.1.p | Glyma.07G025300 | C2H2 |
| 1162. | Glyma.07G025800.1.p | Glyma.07G025800 | ERF |
| 1163. | Glyma.07G027000.1.p | Glyma.07G027000 | ERF |
| 1164. | Glyma.07G027100.1.p | Glyma.07G027100 | bHLH |
| 1165. | Glyma.07G027200.1.p | Glyma.07G027200 | bHLH |
| 1166. | Glyma.07G027300.1.p | Glyma.07G027300 | bHLH |
| 1167. | Glyma.07G027400.1.p | Glyma.07G027400 | bHLH |
| 1168. | Glyma.07G030200.1.p | Glyma.07G030200 | M-type_MADS |
| 1169. | Glyma.07G031200.1.p | Glyma.07G031200 | ERF |
| 1170. | Glyma.07G036200.1.p | Glyma.07G036200 | NF-YA |
| 1171. | Glyma.07G037100.1.p | Glyma.07G037100 | MYB_related |
| 1172. | Glyma.07G037700.1.p | Glyma.07G037700 | MYB |
| 1173. | Glyma.07G038000.1.p | Glyma.07G038000 | MYB |
| 1174. | Glyma.07G038200.1.p | Glyma.07G038200 | AP2 |
| 1175. | Glyma.07G038400.1.p | Glyma.07G038400 | GRF |
| 1176. | Glyma.07G039400.1.p | Glyma.07G039400 | GRAS |
| 1177. | Glyma.07G042800.1.p | Glyma.07G042800 | Trihelix |
| 1178. | Glyma.07G044300.1.p | Glyma.07G044300 | ERF |
| 1179. | Glyma.07G044800.1.p | Glyma.07G044800 | M-type_MADS |
| 1180. | Glyma.07G045000.1.p | Glyma.07G045000 | M-type_MADS |
| 1181. | Glyma.07G045200.1.p | Glyma.07G045200 | M-type_MADS |
| 1182. | Glyma.07G047900.1.p | Glyma.07G047900 | NAC |
| 1183. | Glyma.07G048000.1.p | Glyma.07G048000 | NAC |
| 1184. | Glyma.07G048100.1.p | Glyma.07G048100 | NAC |
| 1185. | Glyma.07G048200.1.p | Glyma.07G048200 | B3 |
| 1186. | Glyma.07G048500.1.p | Glyma.07G048500 | MYB_related |
| 1187. | Glyma.07G049100.1.p | Glyma.07G049100 | bHLH |
| 1188. | Glyma.07G050600.1.p | Glyma.07G050600 | NAC |
| 1189. | Glyma.07G051500.1.p | Glyma.07G051500 | bHLH |
| 1190. | Glyma.07G052100.1.p | Glyma.07G052100 | HD-ZIP |
| 1191. | Glyma.07G053900.1.p | Glyma.07G053900 | Dof |
| 1192. | Glyma.07G054000.1.p | Glyma.07G054000 | MYB |
| 1193. | Glyma.07G054800.1.p | Glyma.07G054800 | ARF |
| 1194. | Glyma.07G055000.1.p | Glyma.07G055000 | ERF |
| 1195. | Glyma.07G055200.1.p | Glyma.07G055200 | bHLH |
| 1196. | Glyma.07G057400.1.p | Glyma.07G057400 | WRKY |
| 1197. | Glyma.07G060400.1.p | Glyma.07G060400 | bZIP |
| 1198. | Glyma.07G066100.1.p | Glyma.07G066100 | MYB_related |
| 1199. | Glyma.07G073000.1.p | Glyma.07G073000 | MYB |
| 1200. | Glyma.07G074300.1.p | Glyma.07G074300 | MYB_related |
| 1201. | Glyma.07G074500.1.p | Glyma.07G074500 | MYB_related |
| 1202. | Glyma.07G076800.1.p | Glyma.07G076800 | HD-ZIP |
| 1203. | Glyma.07G078600.1.p | Glyma.07G078600 | ERF |
| 1204. | Glyma.07G079000.1.p | Glyma.07G079000 | ARR-B |
| 1205. | Glyma.07G080300.1.p | Glyma.07G080300 | TCP |
| 1206. | Glyma.07G080900.1.p | Glyma.07G080900 | MIKC_MADS |
| 1207. | Glyma.07G081300.1.p | Glyma.07G081300 | MIKC_MADS |
| 1208. | Glyma.07G083500.1.p | Glyma.07G083500 | bHLH |
| 1209. | Glyma.07G085300.1.p | Glyma.07G085300 | CPP |
| 1210. | Glyma.07G086300.1.p | Glyma.07G086300 | HSF |
| 1211. | Glyma.07G087900.1.p | Glyma.07G087900 | Trihelix |
| 1212. | Glyma.07G091100.1.p | Glyma.07G091100 | ERF |
| 1213. | Glyma.07G091400.1.p | Glyma.07G091400 | CO-like |
| 1214. | Glyma.07G091600.1.p | Glyma.07G091600 | LBD |
| 1215. | Glyma.07G092000.1.p | Glyma.07G092000 | NAC |
| 1216. | Glyma.07G092700.1.p | Glyma.07G092700 | bHLH |
| 1217. | Glyma.07G092800.1.p | Glyma.07G092800 | MYB |
| 1218. | Glyma.07G099100.1.p | Glyma.07G099100 | BES1 |
| 1219. | Glyma.07G100700.1.p | Glyma.07G100700 | MYB |
| 1220. | Glyma.07G101000.1.p | Glyma.07G101000 | WOX |
| 1221. | Glyma.07G104700.1.p | Glyma.07G104700 | NF-X1 |
| 1222. | Glyma.07G105100.1.p | Glyma.07G105100 | GRAS |
| 1223. | Glyma.07G106800.1.p | Glyma.07G106800 | G2-like |
| 1224. | Glyma.07G107200.1.p | Glyma.07G107200 | ZF-HD |
| 1225. | Glyma.07G107300.1.p | Glyma.07G107300 | BBR-BPC |
| 1226. | Glyma.07G107400.1.p | Glyma.07G107400 | BBR-BPC |
| 1227. | Glyma.07G107500.1.p | Glyma.07G107500 | C2H2 |
| 1228. | Glyma.07G108900.1.p | Glyma.07G108900 | GATA |
| 1229. | Glyma.07G109500.1.p | Glyma.07G109500 | SBP |
| 1230. | Glyma.07G109600.1.p | Glyma.07G109600 | SBP |
| 1231. | Glyma.07G110000.1.p | Glyma.07G110000 | ERF |
| 1232. | Glyma.07G110700.1.p | Glyma.07G110700 | MYB |
| 1233. | Glyma.07G111800.1.p | Glyma.07G111800 | M-type_MADS |
| 1234. | Glyma.07G112400.1.p | Glyma.07G112400 | HD-ZIP |
| 1235. | Glyma.07G113800.1.p | Glyma.07G113800 | ERF |
| 1236. | Glyma.07G114000.1.p | Glyma.07G114000 | ERF |
| 1237. | Glyma.07G114300.1.p | Glyma.07G114300 | ERF |
| 1238. | Glyma.07G116300.1.p | Glyma.07G116300 | WRKY |
| 1239. | Glyma.07G116700.1.p | Glyma.07G116700 | bHLH |
| 1240. | Glyma.07G117500.1.p | Glyma.07G117500 | bHLH |
| 1241. | Glyma.07G117600.1.p | Glyma.07G117600 | bHLH |
| 1242. | Glyma.07G126500.1.p | Glyma.07G126500 | NAC |
| 1243. | Glyma.07G126800.1.p | Glyma.07G126800 | C3H |
| 1244. | Glyma.07G126900.1.p | Glyma.07G126900 | MYB |
| 1245. | Glyma.07G128700.1.p | Glyma.07G128700 | HRT-like |
| 1246. | Glyma.07G130400.1.p | Glyma.07G130400 | ARF |
| 1247. | Glyma.07G131000.1.p | Glyma.07G131000 | WOX |
| 1248. | Glyma.07G132200.1.p | Glyma.07G132200 | MYB |
| 1249. | Glyma.07G132400.1.p | Glyma.07G132400 | MYB |
| 1250. | Glyma.07G133100.1.p | Glyma.07G133100 | GRAS |
| 1251. | Glyma.07G133700.1.p | Glyma.07G133700 | WRKY |
| 1252. | Glyma.07G134800.1.p | Glyma.07G134800 | ARF |
| 1253. | Glyma.07G135800.1.p | Glyma.07G135800 | C2H2 |
| 1254. | Glyma.07G135900.1.p | Glyma.07G135900 | C2H2 |
| 1255. | Glyma.07G141100.1.p | Glyma.07G141100 | MYB |
| 1256. | Glyma.07G146300.1.p | Glyma.07G146300 | MYB |
| 1257. | Glyma.07G151100.1.p | Glyma.07G151100 | Trihelix |
| 1258. | Glyma.07G153600.1.p | Glyma.07G153600 | bHLH |
| 1259. | Glyma.07G153900.1.p | Glyma.07G153900 | G2-like |
| 1260. | Glyma.07G154300.1.p | Glyma.07G154300 | GRAS |
| 1261. | Glyma.07G156200.1.p | Glyma.07G156200 | ERF |
| 1262. | Glyma.07G157300.1.p | Glyma.07G157300 | B3 |
| 1263. | Glyma.07G158000.1.p | Glyma.07G158000 | C2H2 |
| 1264. | Glyma.07G158200.1.p | Glyma.07G158200 | C2H2 |
| 1265. | Glyma.07G158500.1.p | Glyma.07G158500 | FAR1 |
| 1266. | Glyma.07G158600.1.p | Glyma.07G158600 | G2-like |
| 1267. | Glyma.07G159300.1.p | Glyma.07G159300 | SBP |
| 1268. | Glyma.07G161100.1.p | Glyma.07G161100 | WRKY |
| 1269. | Glyma.07G163700.1.p | Glyma.07G163700 | B3 |
| 1270. | Glyma.07G168800.1.p | Glyma.07G168800 | FAR1 |
| 1271. | Glyma.07G170500.1.p | Glyma.07G170500 | C2H2 |
| 1272. | Glyma.07G171200.1.p | Glyma.07G171200 | ARR-B |
| 1273. | Glyma.07G171600.1.p | Glyma.07G171600 | bHLH |
| 1274. | Glyma.07G174000.1.p | Glyma.07G174000 | C2H2 |
| 1275. | Glyma.07G175500.1.p | Glyma.07G175500 | MYB |
| 1276. | Glyma.07G177200.1.p | Glyma.07G177200 | TCP |
| 1277. | Glyma.07G178500.1.p | Glyma.07G178500 | G2-like |
| 1278. | Glyma.07G180200.1.p | Glyma.07G180200 | MYB_related |
| 1279. | Glyma.07G180300.1.p | Glyma.07G180300 | NF-YB |
| 1280. | Glyma.07G181600.1.p | Glyma.07G181600 | MIKC_MADS |
| 1281. | Glyma.07G182700.1.p | Glyma.07G182700 | GATA |
| 1282. | Glyma.07G185300.1.p | Glyma.07G185300 | bHLH |
| 1283. | Glyma.07G187900.1.p | Glyma.07G187900 | bHLH |
| 1284. | Glyma.07G188100.1.p | Glyma.07G188100 | bHLH |
| 1285. | Glyma.07G189300.1.p | Glyma.07G189300 | MYB |
| 1286. | Glyma.07G192000.1.p | Glyma.07G192000 | MYB_related |
| 1287. | Glyma.07G192900.1.p | Glyma.07G192900 | NAC |
| 1288. | Glyma.07G193900.1.p | Glyma.07G193900 | Dof |
| 1289. | Glyma.07G194600.1.p | Glyma.07G194600 | FAR1 |
| 1290. | Glyma.07G196200.1.p | Glyma.07G196200 | LSD |
| 1291. | Glyma.07G197300.1.p | Glyma.07G197300 | bHLH |
| 1292. | Glyma.07G198800.1.p | Glyma.07G198800 | Dof |
| 1293. | Glyma.07G198900.1.p | Glyma.07G198900 | Dof |
| 1294. | Glyma.07G199000.1.p | Glyma.07G199000 | SBP |
| 1295. | Glyma.07G199700.1.p | Glyma.07G199700 | AP2 |
| 1296. | Glyma.07G201200.1.p | Glyma.07G201200 | HB-other |
| 1297. | Glyma.07G201800.1.p | Glyma.07G201800 | NAC |
| 1298. | Glyma.07G202200.1.p | Glyma.07G202200 | ARF |
| 1299. | Glyma.07G202700.1.p | Glyma.07G202700 | VOZ |
| 1300. | Glyma.07G203000.1.p | Glyma.07G203000 | WOX |
| 1301. | Glyma.07G205800.1.p | Glyma.07G205800 | bHLH |
| 1302. | Glyma.07G208000.1.p | Glyma.07G208000 | bHLH |
| 1303. | Glyma.07G209500.1.p | Glyma.07G209500 | G2-like |
| 1304. | Glyma.07G212400.1.p | Glyma.07G212400 | ERF |
| 1305. | Glyma.07G213100.1.p | Glyma.07G213100 | bZIP |
| 1306. | Glyma.07G216000.1.p | Glyma.07G216000 | MYB |
| 1307. | Glyma.07G218000.1.p | Glyma.07G218000 | HD-ZIP |
| 1308. | Glyma.07G219400.1.p | Glyma.07G219400 | WOX |
| 1309. | Glyma.07G223600.1.p | Glyma.07G223600 | LBD |
| 1310. | Glyma.07G224100.1.p | Glyma.07G224100 | B3 |
| 1311. | Glyma.07G224700.1.p | Glyma.07G224700 | FAR1 |
| 1312. | Glyma.07G226800.1.p | Glyma.07G226800 | C2H2 |
| 1313. | Glyma.07G227000.1.p | Glyma.07G227000 | FAR1 |
| 1314. | Glyma.07G227200.1.p | Glyma.07G227200 | WRKY |
| 1315. | Glyma.07G228600.1.p | Glyma.07G228600 | MYB |
| 1316. | Glyma.07G228700.1.p | Glyma.07G228700 | MYB |
| 1317. | Glyma.07G228900.1.p | Glyma.07G228900 | M-type_MADS |
| 1318. | Glyma.07G229100.1.p | Glyma.07G229100 | NAC |
| 1319. | Glyma.07G229600.1.p | Glyma.07G229600 | Dof |
| 1320. | Glyma.07G229800.1.p | Glyma.07G229800 | G2-like |
| 1321. | Glyma.07G230200.1.p | Glyma.07G230200 | ZF-HD |
| 1322. | Glyma.07G230400.1.p | Glyma.07G230400 | SRS |
| 1323. | Glyma.07G234200.1.p | Glyma.07G234200 | SBP |
| 1324. | Glyma.07G235600.1.p | Glyma.07G235600 | HSF |
| 1325. | Glyma.07G235900.1.p | Glyma.07G235900 | MYB |
| 1326. | Glyma.07G238000.1.p | Glyma.07G238000 | WRKY |
| 1327. | Glyma.07G242000.1.p | Glyma.07G242000 | CAMTA |
| 1328. | Glyma.07G242600.1.p | Glyma.07G242600 | MYB |
| 1329. | Glyma.07G242800.1.p | Glyma.07G242800 | C2H2 |
| 1330. | Glyma.07G243000.1.p | Glyma.07G243000 | GATA |
| 1331. | Glyma.07G243300.1.p | Glyma.07G243300 | ARR-B |
| 1332. | Glyma.07G249000.1.p | Glyma.07G249000 | NF-YB |
| 1333. | Glyma.07G250000.1.p | Glyma.07G250000 | C3H |
| 1334. | Glyma.07G250100.1.p | Glyma.07G250100 | ERF |
| 1335. | Glyma.07G251100.1.p | Glyma.07G251100 | bHLH |
| 1336. | Glyma.07G262700.1.p | Glyma.07G262700 | WRKY |
| 1337. | Glyma.07G263600.1.p | Glyma.07G263600 | TALE |
| 1338. | Glyma.07G266500.1.p | Glyma.07G266500 | GRAS |
| 1339. | Glyma.07G268100.1.p | Glyma.07G268100 | NF-YB |
| 1340. | Glyma.07G271100.1.p | Glyma.07G271100 | NAC |
| 1341. | Glyma.07G271900.1.p | Glyma.07G271900 | ZF-HD |
| 1342. | Glyma.07G272800.1.p | Glyma.07G272800 | ARF |

| 1343. | Glyma.08G001300.1.p | Glyma.08G001300 | NF-YB |
| --- | --- | --- | --- |
| 1344. | Glyma.08G005400.1.p | Glyma.08G005400 | MYB |
| 1345. | Glyma.08G007800.1.p | Glyma.08G007800 | Trihelix |
| 1346. | Glyma.08G008100.1.p | Glyma.08G008100 | ARF |
| 1347. | Glyma.08G008200.1.p | Glyma.08G008200 | bHLH |
| 1348. | Glyma.08G009100.1.p | Glyma.08G009100 | bHLH |
| 1349. | Glyma.08G009300.1.p | Glyma.08G009300 | BBR-BPC |
| 1350. | Glyma.08G009700.1.p | Glyma.08G009700 | NAC |
| 1351. | Glyma.08G011300.1.p | Glyma.08G011300 | WRKY |
| 1352. | Glyma.08G011500.1.p | Glyma.08G011500 | SBP |
| 1353. | Glyma.08G014900.1.p | Glyma.08G014900 | bHLH |
| 1354. | Glyma.08G016900.1.p | Glyma.08G016900 | TALE |
| 1355. | Glyma.08G017600.1.p | Glyma.08G017600 | MYB |
| 1356. | Glyma.08G018300.1.p | Glyma.08G018300 | WRKY |
| 1357. | Glyma.08G020900.1.p | Glyma.08G020900 | ERF |
| 1358. | Glyma.08G021900.1.p | Glyma.08G021900 | WRKY |
| 1359. | Glyma.08G027800.1.p | Glyma.08G027800 | ARF |
| 1360. | Glyma.08G029400.1.p | Glyma.08G029400 | MYB_related |
| 1361. | Glyma.08G031400.1.p | Glyma.08G031400 | C3H |
| 1362. | Glyma.08G031900.1.p | Glyma.08G031900 | NAC |
| 1363. | Glyma.08G033900.1.p | Glyma.08G033900 | M-type_MADS |
| 1364. | Glyma.08G034100.1.p | Glyma.08G034100 | M-type_MADS |
| 1365. | Glyma.08G035000.1.p | Glyma.08G035000 | ERF |
| 1366. | Glyma.08G036900.1.p | Glyma.08G036900 | bHLH |
| 1367. | Glyma.08G037400.1.p | Glyma.08G037400 | CPP |
| 1368. | Glyma.08G040900.1.p | Glyma.08G040900 | ERF |
| 1369. | Glyma.08G041100.1.p | Glyma.08G041100 | CO-like |
| 1370. | Glyma.08G041500.1.p | Glyma.08G041500 | NAC |
| 1371. | Glyma.08G042000.1.p | Glyma.08G042000 | bHLH |
| 1372. | Glyma.08G042100.1.p | Glyma.08G042100 | MYB |
| 1373. | Glyma.08G042900.1.p | Glyma.08G042900 | C2H2 |
| 1374. | Glyma.08G046700.1.p | Glyma.08G046700 | ARR-B |
| 1375. | Glyma.08G046800.1.p | Glyma.08G046800 | ARR-B |
| 1376. | Glyma.08G047400.1.p | Glyma.08G047400 | HSF |
| 1377. | Glyma.08G051800.1.p | Glyma.08G051800 | Trihelix |
| 1378. | Glyma.08G053700.1.p | Glyma.08G053700 | WOX |
| 1379. | Glyma.08G055900.1.p | Glyma.08G055900 | NF-X1 |
| 1380. | Glyma.08G056700.1.p | Glyma.08G056700 | ZF-HD |
| 1381. | Glyma.08G056800.1.p | Glyma.08G056800 | C2H2 |
| 1382. | Glyma.08G057400.1.p | Glyma.08G057400 | HD-ZIP |
| 1383. | Glyma.08G059900.1.p | Glyma.08G059900 | MYB |
| 1384. | Glyma.08G061300.1.p | Glyma.08G061300 | bHLH |
| 1385. | Glyma.08G063900.1.p | Glyma.08G063900 | bHLH |
| 1386. | Glyma.08G065300.1.p | Glyma.08G065300 | MIKC_MADS |
| 1387. | Glyma.08G065400.1.p | Glyma.08G065400 | MIKC_MADS |
| 1388. | Glyma.08G067200.1.p | Glyma.08G067200 | GATA |
| 1389. | Glyma.08G068200.1.p | Glyma.08G068200 | MIKC_MADS |
| 1390. | Glyma.08G072100.1.p | Glyma.08G072100 | CAMTA |
| 1391. | Glyma.08G075300.1.p | Glyma.08G075300 | NAC |
| 1392. | Glyma.08G077400.1.p | Glyma.08G077400 | bZIP |
| 1393. | Glyma.08G078100.1.p | Glyma.08G078100 | WRKY |
| 1394. | Glyma.08G078700.1.p | Glyma.08G078700 | WRKY |
| 1395. | Glyma.08G082400.1.p | Glyma.08G082400 | WRKY |
| 1396. | Glyma.08G083900.1.p | Glyma.08G083900 | M-type_MADS |
| 1397. | Glyma.08G089100.1.p | Glyma.08G089100 | bHLH |
| 1398. | Glyma.08G089200.1.p | Glyma.08G089200 | HD-ZIP |
| 1399. | Glyma.08G091000.1.p | Glyma.08G091000 | C3H |
| 1400. | Glyma.08G092300.1.p | Glyma.08G092300 | C2H2 |
| 1401. | Glyma.08G092400.1.p | Glyma.08G092400 | BBR-BPC |
| 1402. | Glyma.08G094200.1.p | Glyma.08G094200 | C2H2 |
| 1403. | Glyma.08G095300.1.p | Glyma.08G095300 | M-type_MADS |
| 1404. | Glyma.08G095500.1.p | Glyma.08G095500 | M-type_MADS |
| 1405. | Glyma.08G095800.1.p | Glyma.08G095800 | GRAS |
| 1406. | Glyma.08G097900.1.p | Glyma.08G097900 | TCP |
| 1407. | Glyma.08G100100.1.p | Glyma.08G100100 | ARF |
| 1408. | Glyma.08G100900.1.p | Glyma.08G100900 | ARR-B |
| 1409. | Glyma.08G105200.1.p | Glyma.08G105200 | CAMTA |
| 1410. | Glyma.08G105400.1.p | Glyma.08G105400 | MIKC_MADS |
| 1411. | Glyma.08G105500.1.p | Glyma.08G105500 | MIKC_MADS |
| 1412. | Glyma.08G108600.1.p | Glyma.08G108600 | HSF |
| 1413. | Glyma.08G110200.1.p | Glyma.08G110200 | MYB_related |
| 1414. | Glyma.08G115300.1.p | Glyma.08G115300 | bZIP |
| 1415. | Glyma.08G115900.1.p | Glyma.08G115900 | Dof |
| 1416. | Glyma.08G116700.1.p | Glyma.08G116700 | G2-like |
| 1417. | Glyma.08G118200.1.p | Glyma.08G118200 | WRKY |
| 1418. | Glyma.08G119900.1.p | Glyma.08G119900 | HSF |
| 1419. | Glyma.08G120600.1.p | Glyma.08G120600 | MIKC_MADS |
| 1420. | Glyma.08G124200.1.p | Glyma.08G124200 | NF-YA |
| 1421. | Glyma.08G124400.1.p | Glyma.08G124400 | HD-ZIP |
| 1422. | Glyma.08G126300.1.p | Glyma.08G126300 | C2H2 |
| 1423. | Glyma.08G126500.1.p | Glyma.08G126500 | bZIP |
| 1424. | Glyma.08G129400.1.p | Glyma.08G129400 | LSD |
| 1425. | Glyma.08G132800.1.p | Glyma.08G132800 | HD-ZIP |
| 1426. | Glyma.08G134700.1.p | Glyma.08G134700 | C2H2 |
| 1427. | Glyma.08G135200.1.p | Glyma.08G135200 | CAMTA |
| 1428. | Glyma.08G137600.1.p | Glyma.08G137600 | ERF |
| 1429. | Glyma.08G137800.1.p | Glyma.08G137800 | EIL |
| 1430. | Glyma.08G140100.1.p | Glyma.08G140100 | bZIP |
| 1431. | Glyma.08G141000.1.p | Glyma.08G141000 | NF-YB |
| 1432. | Glyma.08G142400.1.p | Glyma.08G142400 | WRKY |
| 1433. | Glyma.08G142500.1.p | Glyma.08G142500 | GATA |
| 1434. | Glyma.08G143400.1.p | Glyma.08G143400 | WRKY |
| 1435. | Glyma.08G145300.1.p | Glyma.08G145300 | ERF |
| 1436. | Glyma.08G146900.1.p | Glyma.08G146900 | GRAS |
| 1437. | Glyma.08G147300.1.p | Glyma.08G147300 | LBD |
| 1438. | Glyma.08G148200.1.p | Glyma.08G148200 | NF-YC |
| 1439. | Glyma.08G148500.1.p | Glyma.08G148500 | bHLH |
| 1440. | Glyma.08G148800.1.p | Glyma.08G148800 | HD-ZIP |
| 1441. | Glyma.08G149200.1.p | Glyma.08G149200 | ERF |
| 1442. | Glyma.08G149600.1.p | Glyma.08G149600 | MYB_related |
| 1443. | Glyma.08G152500.1.p | Glyma.08G152500 | bHLH |
| 1444. | Glyma.08G153800.1.p | Glyma.08G153800 | C3H |
| 1445. | Glyma.08G154100.1.p | Glyma.08G154100 | C3H |
| 1446. | Glyma.08G154500.1.p | Glyma.08G154500 | C2H2 |
| 1447. | Glyma.08G156000.1.p | Glyma.08G156000 | bHLH |
| 1448. | Glyma.08G156500.1.p | Glyma.08G156500 | NAC |
| 1449. | Glyma.08G161300.1.p | Glyma.08G161300 | NAC |
| 1450. | Glyma.08G163100.1.p | Glyma.08G163100 | NAC |
| 1451. | Glyma.08G163200.1.p | Glyma.08G163200 | MYB |
| 1452. | Glyma.08G163500.1.p | Glyma.08G163500 | G2-like |
| 1453. | Glyma.08G165700.1.p | Glyma.08G165700 | NF-YC |
| 1454. | Glyma.08G166900.1.p | Glyma.08G166900 | bHLH |
| 1455. | Glyma.08G167000.1.p | Glyma.08G167000 | bHLH |
| 1456. | Glyma.08G168100.1.p | Glyma.08G168100 | MYB |
| 1457. | Glyma.08G169000.1.p | Glyma.08G169000 | LBD |
| 1458. | Glyma.08G169400.1.p | Glyma.08G169400 | NAC |
| 1459. | Glyma.08G172600.1.p | Glyma.08G172600 | FAR1 |
| 1460. | Glyma.08G173400.1.p | Glyma.08G173400 | NAC |
| 1461. | Glyma.08G178900.1.p | Glyma.08G178900 | CAMTA |
| 1462. | Glyma.08G181100.1.p | Glyma.08G181100 | NAC |
| 1463. | Glyma.08G183600.1.p | Glyma.08G183600 | bZIP |
| 1464. | Glyma.08G184500.1.p | Glyma.08G184500 | GATA |
| 1465. | Glyma.08G188900.1.p | Glyma.08G188900 | MYB |
| 1466. | Glyma.08G189000.1.p | Glyma.08G189000 | ZF-HD |
| 1467. | Glyma.08G189900.1.p | Glyma.08G189900 | MYB_related |
| 1468. | Glyma.08G191500.1.p | Glyma.08G191500 | MYB |
| 1469. | Glyma.08G192300.1.p | Glyma.08G192300 | C2H2 |
| 1470. | Glyma.08G192600.1.p | Glyma.08G192600 | LBD |
| 1471. | Glyma.08G195300.1.p | Glyma.08G195300 | Dof |
| 1472. | Glyma.08G195700.1.p | Glyma.08G195700 | C2H2 |
| 1473. | Glyma.08G195800.1.p | Glyma.08G195800 | C2H2 |
| 1474. | Glyma.08G197600.1.p | Glyma.08G197600 | bHLH |
| 1475. | Glyma.08G201900.1.p | Glyma.08G201900 | HD-ZIP |
| 1476. | Glyma.08G202000.1.p | Glyma.08G202000 | HD-ZIP |
| 1477. | Glyma.08G202100.1.p | Glyma.08G202100 | GATA |
| 1478. | Glyma.08G202300.1.p | Glyma.08G202300 | ERF |
| 1479. | Glyma.08G203600.1.p | Glyma.08G203600 | bHLH |
| 1480. | Glyma.08G204700.1.p | Glyma.08G204700 | HD-ZIP |
| 1481. | Glyma.08G211600.1.p | Glyma.08G211600 | ERF |
| 1482. | Glyma.08G212700.1.p | Glyma.08G212700 | M-type_MADS |
| 1483. | Glyma.08G215300.1.p | Glyma.08G215300 | bHLH |
| 1484. | Glyma.08G215400.1.p | Glyma.08G215400 | bHLH |
| 1485. | Glyma.08G215500.1.p | Glyma.08G215500 | bHLH |
| 1486. | Glyma.08G215600.1.p | Glyma.08G215600 | bHLH |
| 1487. | Glyma.08G215700.1.p | Glyma.08G215700 | ERF |
| 1488. | Glyma.08G216600.1.p | Glyma.08G216600 | ERF |
| 1489. | Glyma.08G217100.1.p | Glyma.08G217100 | C2H2 |
| 1490. | Glyma.08G218600.1.p | Glyma.08G218600 | WRKY |
| 1491. | Glyma.08G220800.1.p | Glyma.08G220800 | AP2 |
| 1492. | Glyma.08G221800.1.p | Glyma.08G221800 | GATA |
| 1493. | Glyma.08G227000.1.p | Glyma.08G227000 | bZIP |
| 1494. | Glyma.08G227500.1.p | Glyma.08G227500 | FAR1 |
| 1495. | Glyma.08G227700.1.p | Glyma.08G227700 | AP2 |
| 1496. | Glyma.08G228900.1.p | Glyma.08G228900 | Dof |
| 1497. | Glyma.08G232700.1.p | Glyma.08G232700 | C3H |
| 1498. | Glyma.08G235200.1.p | Glyma.08G235200 | LBD |
| 1499. | Glyma.08G237300.1.p | Glyma.08G237300 | GRAS |
| 1500. | Glyma.08G239500.1.p | Glyma.08G239500 | bHLH |
| 1501. | Glyma.08G240800.1.p | Glyma.08G240800 | WRKY |
| 1502. | Glyma.08G247300.1.p | Glyma.08G247300 | TCP |
| 1503. | Glyma.08G250600.1.p | Glyma.08G250600 | MYB |
| 1504. | Glyma.08G250700.1.p | Glyma.08G250700 | MIKC_MADS |
| 1505. | Glyma.08G250800.1.p | Glyma.08G250800 | MIKC_MADS |
| 1506. | Glyma.08G252800.1.p | Glyma.08G252800 | bHLH |
| 1507. | Glyma.08G254400.1.p | Glyma.08G254400 | bZIP |
| 1508. | Glyma.08G255200.1.p | Glyma.08G255200 | CO-like |
| 1509. | Glyma.08G256400.1.p | Glyma.08G256400 | TCP |
| 1510. | Glyma.08G257300.1.p | Glyma.08G257300 | ERF |
| 1511. | Glyma.08G257500.1.p | Glyma.08G257500 | Trihelix |
| 1512. | Glyma.08G261500.1.p | Glyma.08G261500 | ERF |
| 1513. | Glyma.08G261900.1.p | Glyma.08G261900 | LBD |
| 1514. | Glyma.08G265600.1.p | Glyma.08G265600 | bZIP |
| 1515. | Glyma.08G269800.1.p | Glyma.08G269800 | MIKC_MADS |
| 1516. | Glyma.08G271000.1.p | Glyma.08G271000 | bHLH |
| 1517. | Glyma.08G271900.1.p | Glyma.08G271900 | bHLH |
| 1518. | Glyma.08G274200.1.p | Glyma.08G274200 | bHLH |
| 1519. | Glyma.08G276300.1.p | Glyma.08G276300 | Dof |
| 1520. | Glyma.08G278800.1.p | Glyma.08G278800 | ERF |
| 1521. | Glyma.08G279000.1.p | Glyma.08G279000 | AP2 |
| 1522. | Glyma.08G280900.1.p | Glyma.08G280900 | ZF-HD |
| 1523. | Glyma.08G281000.1.p | Glyma.08G281000 | ZF-HD |
| 1524. | Glyma.08G281100.1.p | Glyma.08G281100 | ZF-HD |
| 1525. | Glyma.08G281200.1.p | Glyma.08G281200 | ZF-HD |
| 1526. | Glyma.08G281900.1.p | Glyma.08G281900 | ERF |
| 1527. | Glyma.08G282500.1.p | Glyma.08G282500 | M-type_MADS |
| 1528. | Glyma.08G284300.1.p | Glyma.08G284300 | TALE |
| 1529. | Glyma.08G285200.1.p | Glyma.08G285200 | YABBY |
| 1530. | Glyma.08G286000.1.p | Glyma.08G286000 | C3H |
| 1531. | Glyma.08G286500.1.p | Glyma.08G286500 | bHLH |
| 1532. | Glyma.08G290100.1.p | Glyma.08G290100 | MYB |
| 1533. | Glyma.08G293300.1.p | Glyma.08G293300 | MYB_related |
| 1534. | Glyma.08G294000.1.p | Glyma.08G294000 | bHLH |
| 1535. | Glyma.08G295800.1.p | Glyma.08G295800 | HD-ZIP |
| 1536. | Glyma.08G297000.1.p | Glyma.08G297000 | AP2 |
| 1537. | Glyma.08G297200.1.p | Glyma.08G297200 | Whirly |
| 1538. | Glyma.08G298200.1.p | Glyma.08G298200 | MYB |
| 1539. | Glyma.08G298400.1.p | Glyma.08G298400 | HD-ZIP |
| 1540. | Glyma.08G298700.1.p | Glyma.08G298700 | bHLH |
| 1541. | Glyma.08G299400.1.p | Glyma.08G299400 | TCP |
| 1542. | Glyma.08G301100.1.p | Glyma.08G301100 | NAC |
| 1543. | Glyma.08G301700.1.p | Glyma.08G301700 | HB-other |
| 1544. | Glyma.08G302500.1.p | Glyma.08G302500 | bZIP |
| 1545. | Glyma.08G303900.1.p | Glyma.08G303900 | bHLH |
| 1546. | Glyma.08G307100.1.p | Glyma.08G307100 | NAC |
| 1547. | Glyma.08G310100.1.p | Glyma.08G310100 | MIKC_MADS |
| 1548. | Glyma.08G317300.1.p | Glyma.08G317300 | MYB |
| 1549. | Glyma.08G317600.1.p | Glyma.08G317600 | MYB |
| 1550. | Glyma.08G317800.1.p | Glyma.08G317800 | MYB |
| 1551. | Glyma.08G320200.1.p | Glyma.08G320200 | WRKY |
| 1552. | Glyma.08G320700.1.p | Glyma.08G320700 | ERF |
| 1553. | Glyma.08G325800.1.p | Glyma.08G325800 | WRKY |
| 1554. | Glyma.08G325900.1.p | Glyma.08G325900 | GRAS |
| 1555. | Glyma.08G329300.1.p | Glyma.08G329300 | NF-YB |
| 1556. | Glyma.08G331300.1.p | Glyma.08G331300 | MYB_related |
| 1557. | Glyma.08G333500.1.p | Glyma.08G333500 | MYB_related |
| 1558. | Glyma.08G333600.1.p | Glyma.08G333600 | MYB_related |
| 1559. | Glyma.08G334200.1.p | Glyma.08G334200 | LBD |
| 1560. | Glyma.08G335900.1.p | Glyma.08G335900 | NF-YA |
| 1561. | Glyma.08G336500.1.p | Glyma.08G336500 | MYB |
| 1562. | Glyma.08G339100.1.p | Glyma.08G339100 | B3 |
| 1563. | Glyma.08G339200.1.p | Glyma.08G339200 | B3 |
| 1564. | Glyma.08G342800.1.p | Glyma.08G342800 | FAR1 |
| 1565. | Glyma.08G344500.1.p | Glyma.08G344500 | GATA |
| 1566. | Glyma.08G346300.1.p | Glyma.08G346300 | bHLH |
| 1567. | Glyma.08G348300.1.p | Glyma.08G348300 | ERF |
| 1568. | Glyma.08G356900.1.p | Glyma.08G356900 | bHLH |
| 1569. | Glyma.08G357600.1.p | Glyma.08G357600 | B3 |
| 1570. | Glyma.08G358100.1.p | Glyma.08G358100 | Dof |
| 1571. | Glyma.08G360200.1.p | Glyma.08G360200 | NAC |
| 1572. | Glyma.08G365100.1.p | Glyma.08G365100 | LBD |

| 1573. | Glyma.09G001300.1.p | Glyma.09G001300 | FAR1 |
| --- | --- | --- | --- |
| 1574. | Glyma.09G001500.1.p | Glyma.09G001500 | MYB |
| 1575. | Glyma.09G004500.1.p | Glyma.09G004500 | G2-like |
| 1576. | Glyma.09G005700.1.p | Glyma.09G005700 | WRKY |
| 1577. | Glyma.09G007500.1.p | Glyma.09G007500 | TALE |
| 1578. | Glyma.09G011800.1.p | Glyma.09G011800 | GRAS |
| 1579. | Glyma.09G012900.1.p | Glyma.09G012900 | FAR1 |
| 1580. | Glyma.09G014100.1.p | Glyma.09G014100 | NF-YB |
| 1581. | Glyma.09G014300.1.p | Glyma.09G014300 | Trihelix |
| 1582. | Glyma.09G017300.1.p | Glyma.09G017300 | G2-like |
| 1583. | Glyma.09G017400.1.p | Glyma.09G017400 | G2-like |
| 1584. | Glyma.09G019300.1.p | Glyma.09G019300 | FAR1 |
| 1585. | Glyma.09G021600.1.p | Glyma.09G021600 | LBD |
| 1586. | Glyma.09G023600.1.p | Glyma.09G023600 | HD-ZIP |
| 1587. | Glyma.09G023800.1.p | Glyma.09G023800 | NF-YA |
| 1588. | Glyma.09G025800.1.p | Glyma.09G025800 | HD-ZIP |
| 1589. | Glyma.09G025900.1.p | Glyma.09G025900 | HD-ZIP |
| 1590. | Glyma.09G029300.1.p | Glyma.09G029300 | MYB_related |
| 1591. | Glyma.09G029800.1.p | Glyma.09G029800 | WRKY |
| 1592. | Glyma.09G032100.1.p | Glyma.09G032100 | MYB |
| 1593. | Glyma.09G034100.1.p | Glyma.09G034100 | C2H2 |
| 1594. | Glyma.09G034300.1.p | Glyma.09G034300 | WRKY |
| 1595. | Glyma.09G036500.1.p | Glyma.09G036500 | GRAS |
| 1596. | Glyma.09G038300.1.p | Glyma.09G038300 | CAMTA |
| 1597. | Glyma.09G038900.1.p | Glyma.09G038900 | MYB |
| 1598. | Glyma.09G039300.1.p | Glyma.09G039300 | FAR1 |
| 1599. | Glyma.09G040000.1.p | Glyma.09G040000 | ARR-B |
| 1600. | Glyma.09G041500.1.p | Glyma.09G041500 | ERF |
| 1601. | Glyma.09G046200.1.p | Glyma.09G046200 | NF-YB |
| 1602. | Glyma.09G052800.1.p | Glyma.09G052800 | ERF |
| 1603. | Glyma.09G052900.1.p | Glyma.09G052900 | ERF |
| 1604. | Glyma.09G053000.1.p | Glyma.09G053000 | ERF |
| 1605. | Glyma.09G059700.1.p | Glyma.09G059700 | HB-PHD |
| 1606. | Glyma.09G060200.1.p | Glyma.09G060200 | bHLH |
| 1607. | Glyma.09G061900.1.p | Glyma.09G061900 | WRKY |
| 1608. | Glyma.09G062400.1.p | Glyma.09G062400 | HD-ZIP |
| 1609. | Glyma.09G062800.1.p | Glyma.09G062800 | GATA |
| 1610. | Glyma.09G064200.1.p | Glyma.09G064200 | bHLH |
| 1611. | Glyma.09G068400.1.p | Glyma.09G068400 | NF-YA |
| 1612. | Glyma.09G068700.1.p | Glyma.09G068700 | GRF |
| 1613. | Glyma.09G072000.1.p | Glyma.09G072000 | ERF |
| 1614. | Glyma.09G072200.1.p | Glyma.09G072200 | ARF |
| 1615. | Glyma.09G076800.1.p | Glyma.09G076800 | GATA |
| 1616. | Glyma.09G080000.1.p | Glyma.09G080000 | WRKY |
| 1617. | Glyma.09G082000.1.p | Glyma.09G082000 | SBP |
| 1618. | Glyma.09G083500.1.p | Glyma.09G083500 | LSD |
| 1619. | Glyma.09G092500.1.p | Glyma.09G092500 | MYB |
| 1620. | Glyma.09G092900.1.p | Glyma.09G092900 | MYB |
| 1621. | Glyma.09G095700.1.p | Glyma.09G095700 | TALE |
| 1622. | Glyma.09G098300.1.p | Glyma.09G098300 | bHLH |
| 1623. | Glyma.09G098600.1.p | Glyma.09G098600 | ARR-B |
| 1624. | Glyma.09G099000.1.p | Glyma.09G099000 | DBB |
| 1625. | Glyma.09G102700.1.p | Glyma.09G102700 | B3 |
| 1626. | Glyma.09G102900.1.p | Glyma.09G102900 | C3H |
| 1627. | Glyma.09G109700.1.p | Glyma.09G109700 | HD-ZIP |
| 1628. | Glyma.09G113000.1.p | Glyma.09G113000 | G2-like |
| 1629. | Glyma.09G113100.1.p | Glyma.09G113100 | G2-like |
| 1630. | Glyma.09G113800.1.p | Glyma.09G113800 | SBP |
| 1631. | Glyma.09G114900.1.p | Glyma.09G114900 | B3 |
| 1632. | Glyma.09G116400.1.p | Glyma.09G116400 | Trihelix |
| 1633. | Glyma.09G117200.1.p | Glyma.09G117200 | B3 |
| 1634. | Glyma.09G117500.1.p | Glyma.09G117500 | B3 |
| 1635. | Glyma.09G121900.1.p | Glyma.09G121900 | FAR1 |
| 1636. | Glyma.09G123300.1.p | Glyma.09G123300 | GRAS |
| 1637. | Glyma.09G127100.1.p | Glyma.09G127100 | WRKY |
| 1638. | Glyma.09G129100.1.p | Glyma.09G129100 | WRKY |
| 1639. | Glyma.09G131400.1.p | Glyma.09G131400 | MYB_related |
| 1640. | Glyma.09G133600.1.p | Glyma.09G133600 | GRAS |
| 1641. | Glyma.09G137000.1.p | Glyma.09G137000 | Nin-like |
| 1642. | Glyma.09G139000.1.p | Glyma.09G139000 | MYB |
| 1643. | Glyma.09G143200.1.p | Glyma.09G143200 | HSF |
| 1644. | Glyma.09G143700.1.p | Glyma.09G143700 | HD-ZIP |
| 1645. | Glyma.09G147100.1.p | Glyma.09G147100 | G2-like |
| 1646. | Glyma.09G147200.1.p | Glyma.09G147200 | ERF |
| 1647. | Glyma.09G148000.1.p | Glyma.09G148000 | C3H |
| 1648. | Glyma.09G149000.1.p | Glyma.09G149000 | MIKC_MADS |
| 1649. | Glyma.09G149700.1.p | Glyma.09G149700 | Trihelix |
| 1650. | Glyma.09G149900.1.p | Glyma.09G149900 | bHLH |
| 1651. | Glyma.09G150000.1.p | Glyma.09G150000 | bHLH |
| 1652. | Glyma.09G158400.1.p | Glyma.09G158400 | NF-YB |
| 1653. | Glyma.09G167400.1.p | Glyma.09G167400 | NAC |
| 1654. | Glyma.09G167900.1.p | Glyma.09G167900 | MYB_related |
| 1655. | Glyma.09G168000.1.p | Glyma.09G168000 | HD-ZIP |
| 1656. | Glyma.09G169300.1.p | Glyma.09G169300 | MYB |
| 1657. | Glyma.09G170200.1.p | Glyma.09G170200 | C2H2 |
| 1658. | Glyma.09G170300.1.p | Glyma.09G170300 | BBR-BPC |
| 1659. | Glyma.09G170400.1.p | Glyma.09G170400 | BBR-BPC |
| 1660. | Glyma.09G170500.1.p | Glyma.09G170500 | ZF-HD |
| 1661. | Glyma.09G171100.1.p | Glyma.09G171100 | G2-like |
| 1662. | Glyma.09G173000.1.p | Glyma.09G173000 | NF-X1 |
| 1663. | Glyma.09G177800.1.p | Glyma.09G177800 | WOX |
| 1664. | Glyma.09G178000.1.p | Glyma.09G178000 | MYB |
| 1665. | Glyma.09G183400.1.p | Glyma.09G183400 | MYB |
| 1666. | Glyma.09G183500.1.p | Glyma.09G183500 | bHLH |
| 1667. | Glyma.09G184100.1.p | Glyma.09G184100 | NAC |
| 1668. | Glyma.09G184300.1.p | Glyma.09G184300 | bZIP |
| 1669. | Glyma.09G184600.1.p | Glyma.09G184600 | CO-like |
| 1670. | Glyma.09G184800.1.p | Glyma.09G184800 | ERF |
| 1671. | Glyma.09G188900.1.p | Glyma.09G188900 | Trihelix |
| 1672. | Glyma.09G190600.1.p | Glyma.09G190600 | HSF |
| 1673. | Glyma.09G191600.1.p | Glyma.09G191600 | CPP |
| 1674. | Glyma.09G194300.1.p | Glyma.09G194300 | B3 |
| 1675. | Glyma.09G194800.1.p | Glyma.09G194800 | ERF |
| 1676. | Glyma.09G196500.1.p | Glyma.09G196500 | MYB_related |
| 1677. | Glyma.09G199800.1.p | Glyma.09G199800 | AP2 |
| 1678. | Glyma.09G201000.1.p | Glyma.09G201000 | Dof |
| 1679. | Glyma.09G201700.1.p | Glyma.09G201700 | MIKC_MADS |
| 1680. | Glyma.09G203000.1.p | Glyma.09G203000 | bHLH |
| 1681. | Glyma.09G204500.1.p | Glyma.09G204500 | bHLH |
| 1682. | Glyma.09G206200.1.p | Glyma.09G206200 | MYB |
| 1683. | Glyma.09G206600.1.p | Glyma.09G206600 | HSF |
| 1684. | Glyma.09G207300.1.p | Glyma.09G207300 | G2-like |
| 1685. | Glyma.09G207500.1.p | Glyma.09G207500 | HD-ZIP |
| 1686. | Glyma.09G208500.1.p | Glyma.09G208500 | bZIP |
| 1687. | Glyma.09G211400.1.p | Glyma.09G211400 | G2-like |
| 1688. | Glyma.09G212500.1.p | Glyma.09G212500 | GRF |
| 1689. | Glyma.09G217000.1.p | Glyma.09G217000 | BES1 |
| 1690. | Glyma.09G223500.1.p | Glyma.09G223500 | CPP |
| 1691. | Glyma.09G224900.1.p | Glyma.09G224900 | GRAS |
| 1692. | Glyma.09G230200.1.p | Glyma.09G230200 | C2H2 |
| 1693. | Glyma.09G231600.1.p | Glyma.09G231600 | MIKC_MADS |
| 1694. | Glyma.09G231700.1.p | Glyma.09G231700 | NAC |
| 1695. | Glyma.09G233600.1.p | Glyma.09G233600 | NAC |
| 1696. | Glyma.09G233800.1.p | Glyma.09G233800 | ERF |
| 1697. | Glyma.09G234900.1.p | Glyma.09G234900 | MYB |
| 1698. | Glyma.09G235100.1.p | Glyma.09G235100 | MYB |
| 1699. | Glyma.09G235300.1.p | Glyma.09G235300 | MYB |
| 1700. | Glyma.09G235600.1.p | Glyma.09G235600 | MYB |
| 1701. | Glyma.09G235700.1.p | Glyma.09G235700 | NAC |
| 1702. | Glyma.09G237000.1.p | Glyma.09G237000 | Dof |
| 1703. | Glyma.09G238400.1.p | Glyma.09G238400 | ZF-HD |
| 1704. | Glyma.09G238500.1.p | Glyma.09G238500 | ZF-HD |
| 1705. | Glyma.09G238600.1.p | Glyma.09G238600 | ZF-HD |
| 1706. | Glyma.09G238700.1.p | Glyma.09G238700 | ZF-HD |
| 1707. | Glyma.09G238800.1.p | Glyma.09G238800 | MYB |
| 1708. | Glyma.09G239100.1.p | Glyma.09G239100 | bHLH |
| 1709. | Glyma.09G239400.1.p | Glyma.09G239400 | HD-ZIP |
| 1710. | Glyma.09G240000.1.p | Glyma.09G240000 | WRKY |
| 1711. | Glyma.09G240400.1.p | Glyma.09G240400 | AP2 |
| 1712. | Glyma.09G241800.1.p | Glyma.09G241800 | HD-ZIP |
| 1713. | Glyma.09G242600.1.p | Glyma.09G242600 | ERF |
| 1714. | Glyma.09G244000.1.p | Glyma.09G244000 | WRKY |
| 1715. | Glyma.09G245300.1.p | Glyma.09G245300 | Trihelix |
| 1716. | Glyma.09G246800.1.p | Glyma.09G246800 | bHLH |
| 1717. | Glyma.09G248200.1.p | Glyma.09G248200 | AP2 |
| 1718. | Glyma.09G250500.1.p | Glyma.09G250500 | WRKY |
| 1719. | Glyma.09G251200.1.p | Glyma.09G251200 | C3H |
| 1720. | Glyma.09G254400.1.p | Glyma.09G254400 | WRKY |
| 1721. | Glyma.09G254800.1.p | Glyma.09G254800 | WRKY |
| 1722. | Glyma.09G255300.1.p | Glyma.09G255300 | MYB_related |
| 1723. | Glyma.09G257700.1.p | Glyma.09G257700 | MYB_related |
| 1724. | Glyma.09G260100.1.p | Glyma.09G260100 | BES1 |
| 1725. | Glyma.09G261600.1.p | Glyma.09G261600 | MYB |
| 1726. | Glyma.09G265200.1.p | Glyma.09G265200 | HD-ZIP |
| 1727. | Glyma.09G266200.1.p | Glyma.09G266200 | MIKC_MADS |
| 1728. | Glyma.09G266400.1.p | Glyma.09G266400 | MIKC_MADS |
| 1729. | Glyma.09G268700.1.p | Glyma.09G268700 | LBD |
| 1730. | Glyma.09G270000.1.p | Glyma.09G270000 | GRAS |
| 1731. | Glyma.09G274000.1.p | Glyma.09G274000 | WRKY |
| 1732. | Glyma.09G278100.1.p | Glyma.09G278100 | bHLH |
| 1733. | Glyma.09G280200.1.p | Glyma.09G280200 | WRKY |
| 1734. | Glyma.09G283800.1.p | Glyma.09G283800 | MIKC_MADS |
| 1735. | Glyma.09G284300.1.p | Glyma.09G284300 | TCP |
| 1736. | Glyma.09G284500.1.p | Glyma.09G284500 | TCP |

| 1737. | Glyma.10G001400.1.p | Glyma.10G001400 | FAR1 |
| --- | --- | --- | --- |
| 1738. | Glyma.10G003100.1.p | Glyma.10G003100 | HSF |
| 1739. | Glyma.10G004000.1.p | Glyma.10G004000 | C2H2 |
| 1740. | Glyma.10G006600.1.p | Glyma.10G006600 | MYB |
| 1741. | Glyma.10G007000.1.p | Glyma.10G007000 | ERF |
| 1742. | Glyma.10G007100.1.p | Glyma.10G007100 | ERF |
| 1743. | Glyma.10G007300.1.p | Glyma.10G007300 | bHLH |
| 1744. | Glyma.10G009200.1.p | Glyma.10G009200 | SBP |
| 1745. | Glyma.10G010300.1.p | Glyma.10G010300 | MYB |
| 1746. | Glyma.10G010400.1.p | Glyma.10G010400 | MYB |
| 1747. | Glyma.10G011300.1.p | Glyma.10G011300 | WRKY |
| 1748. | Glyma.10G012500.1.p | Glyma.10G012500 | GRAS |
| 1749. | Glyma.10G013300.1.p | Glyma.10G013300 | bZIP |
| 1750. | Glyma.10G014600.1.p | Glyma.10G014600 | MYB |
| 1751. | Glyma.10G016500.1.p | Glyma.10G016500 | ERF |
| 1752. | Glyma.10G020100.1.p | Glyma.10G020100 | NF-YB |
| 1753. | Glyma.10G020200.1.p | Glyma.10G020200 | C2H2 |
| 1754. | Glyma.10G020600.1.p | Glyma.10G020600 | C3H |
| 1755. | Glyma.10G020700.1.p | Glyma.10G020700 | C3H |
| 1756. | Glyma.10G021400.1.p | Glyma.10G021400 | CO-like |
| 1757. | Glyma.10G026000.1.p | Glyma.10G026000 | bHLH |
| 1758. | Glyma.10G029600.1.p | Glyma.10G029600 | HSF |
| 1759. | Glyma.10G029700.1.p | Glyma.10G029700 | C2H2 |
| 1760. | Glyma.10G031600.1.p | Glyma.10G031600 | bHLH |
| 1761. | Glyma.10G032900.1.p | Glyma.10G032900 | WRKY |
| 1762. | Glyma.10G034300.1.p | Glyma.10G034300 | bHLH |
| 1763. | Glyma.10G034600.1.p | Glyma.10G034600 | C2H2 |
| 1764. | Glyma.10G035000.1.p | Glyma.10G035000 | LBD |
| 1765. | Glyma.10G035100.1.p | Glyma.10G035100 | LBD |
| 1766. | Glyma.10G036200.1.p | Glyma.10G036200 | ERF |
| 1767. | Glyma.10G036300.1.p | Glyma.10G036300 | ERF |
| 1768. | Glyma.10G036600.1.p | Glyma.10G036600 | ERF |
| 1769. | Glyma.10G036700.1.p | Glyma.10G036700 | ERF |
| 1770. | Glyma.10G037000.1.p | Glyma.10G037000 | MYB |
| 1771. | Glyma.10G037700.1.p | Glyma.10G037700 | NAC |
| 1772. | Glyma.10G038500.1.p | Glyma.10G038500 | GRAS |
| 1773. | Glyma.10G039700.1.p | Glyma.10G039700 | G2-like |
| 1774. | Glyma.10G042800.1.p | Glyma.10G042800 | bHLH |
| 1775. | Glyma.10G045200.1.p | Glyma.10G045200 | C2H2 |
| 1776. | Glyma.10G045300.1.p | Glyma.10G045300 | C2H2 |
| 1777. | Glyma.10G045400.1.p | Glyma.10G045400 | C2H2 |
| 1778. | Glyma.10G048500.1.p | Glyma.10G048500 | MYB_related |
| 1779. | Glyma.10G048900.1.p | Glyma.10G048900 | NF-YB |
| 1780. | Glyma.10G050700.1.p | Glyma.10G050700 | S1Fa-like |
| 1781. | Glyma.10G051500.1.p | Glyma.10G051500 | C2H2 |
| 1782. | Glyma.10G053500.1.p | Glyma.10G053500 | ARF |
| 1783. | Glyma.10G054700.1.p | Glyma.10G054700 | MYB |
| 1784. | Glyma.10G057400.1.p | Glyma.10G057400 | TCP |
| 1785. | Glyma.10G057800.1.p | Glyma.10G057800 | MYB_related |
| 1786. | Glyma.10G059000.1.p | Glyma.10G059000 | MYB |
| 1787. | Glyma.10G059700.1.p | Glyma.10G059700 | C2H2 |
| 1788. | Glyma.10G061400.1.p | Glyma.10G061400 | ERF |
| 1789. | Glyma.10G064700.1.p | Glyma.10G064700 | HD-ZIP |
| 1790. | Glyma.10G064900.1.p | Glyma.10G064900 | Trihelix |
| 1791. | Glyma.10G065100.1.p | Glyma.10G065100 | Trihelix |
| 1792. | Glyma.10G066100.1.p | Glyma.10G066100 | HSF |
| 1793. | Glyma.10G066800.1.p | Glyma.10G066800 | Trihelix |
| 1794. | Glyma.10G066900.1.p | Glyma.10G066900 | ERF |
| 1795. | Glyma.10G067000.1.p | Glyma.10G067000 | ERF |
| 1796. | Glyma.10G067200.1.p | Glyma.10G067200 | GRF |
| 1797. | Glyma.10G068900.1.p | Glyma.10G068900 | VOZ |
| 1798. | Glyma.10G069700.1.p | Glyma.10G069700 | WOX |
| 1799. | Glyma.10G071700.1.p | Glyma.10G071700 | bZIP |
| 1800. | Glyma.10G076100.1.p | Glyma.10G076100 | B3 |
| 1801. | Glyma.10G077000.1.p | Glyma.10G077000 | NAC |
| 1802. | Glyma.10G077400.1.p | Glyma.10G077400 | NAC |
| 1803. | Glyma.10G078200.1.p | Glyma.10G078200 | E2F/DP |
| 1804. | Glyma.10G081700.1.p | Glyma.10G081700 | TALE |
| 1805. | Glyma.10G082000.1.p | Glyma.10G082000 | Dof |
| 1806. | Glyma.10G082800.1.p | Glyma.10G082800 | NF-YA |
| 1807. | Glyma.10G083200.1.p | Glyma.10G083200 | M-type_MADS |
| 1808. | Glyma.10G084700.1.p | Glyma.10G084700 | M-type_MADS |
| 1809. | Glyma.10G084800.1.p | Glyma.10G084800 | M-type_MADS |
| 1810. | Glyma.10G085100.1.p | Glyma.10G085100 | M-type_MADS |
| 1811. | Glyma.10G085600.1.p | Glyma.10G085600 | M-type_MADS |
| 1812. | Glyma.10G085700.1.p | Glyma.10G085700 | M-type_MADS |
| 1813. | Glyma.10G085900.1.p | Glyma.10G085900 | M-type_MADS |
| 1814. | Glyma.10G086200.1.p | Glyma.10G086200 | M-type_MADS |
| 1815. | Glyma.10G086300.1.p | Glyma.10G086300 | M-type_MADS |
| 1816. | Glyma.10G086400.1.p | Glyma.10G086400 | M-type_MADS |
| 1817. | Glyma.10G087200.1.p | Glyma.10G087200 | LBD |
| 1818. | Glyma.10G088600.1.p | Glyma.10G088600 | M-type_MADS |
| 1819. | Glyma.10G092100.1.p | Glyma.10G092100 | bZIP |
| 1820. | Glyma.10G093100.1.p | Glyma.10G093100 | bHLH |
| 1821. | Glyma.10G093600.1.p | Glyma.10G093600 | bHLH |
| 1822. | Glyma.10G094200.1.p | Glyma.10G094200 | M-type_MADS |
| 1823. | Glyma.10G095100.1.p | Glyma.10G095100 | C2H2 |
| 1824. | Glyma.10G099800.1.p | Glyma.10G099800 | ERF |
| 1825. | Glyma.10G100400.1.p | Glyma.10G100400 | C3H |
| 1826. | Glyma.10G101000.1.p | Glyma.10G101000 | ARR-B |
| 1827. | Glyma.10G101900.1.p | Glyma.10G101900 | C2H2 |
| 1828. | Glyma.10G103400.1.p | Glyma.10G103400 | C2H2 |
| 1829. | Glyma.10G111400.1.p | Glyma.10G111400 | WRKY |
| 1830. | Glyma.10G113800.1.p | Glyma.10G113800 | WRKY |
| 1831. | Glyma.10G116600.1.p | Glyma.10G116600 | AP2 |
| 1832. | Glyma.10G117500.1.p | Glyma.10G117500 | MYB_related |
| 1833. | Glyma.10G118400.1.p | Glyma.10G118400 | MYB_related |
| 1834. | Glyma.10G118900.1.p | Glyma.10G118900 | ERF |
| 1835. | Glyma.10G119100.1.p | Glyma.10G119100 | ERF |
| 1836. | Glyma.10G121700.1.p | Glyma.10G121700 | FAR1 |
| 1837. | Glyma.10G124900.1.p | Glyma.10G124900 | FAR1 |
| 1838. | Glyma.10G126900.1.p | Glyma.10G126900 | GATA |
| 1839. | Glyma.10G132200.1.p | Glyma.10G132200 | MYB |
| 1840. | Glyma.10G138300.1.p | Glyma.10G138300 | WRKY |
| 1841. | Glyma.10G138800.1.p | Glyma.10G138800 | bHLH |
| 1842. | Glyma.10G139000.1.p | Glyma.10G139000 | MYB |
| 1843. | Glyma.10G142200.1.p | Glyma.10G142200 | MYB |
| 1844. | Glyma.10G142600.1.p | Glyma.10G142600 | bHLH |
| 1845. | Glyma.10G142900.1.p | Glyma.10G142900 | M-type_MADS |
| 1846. | Glyma.10G144700.1.p | Glyma.10G144700 | C3H |
| 1847. | Glyma.10G147000.1.p | Glyma.10G147000 | TALE |
| 1848. | Glyma.10G153200.1.p | Glyma.10G153200 | C2H2 |
| 1849. | Glyma.10G153500.1.p | Glyma.10G153500 | NF-YB |
| 1850. | Glyma.10G153800.1.p | Glyma.10G153800 | MYB_related |
| 1851. | Glyma.10G155900.1.p | Glyma.10G155900 | NF-YC |
| 1852. | Glyma.10G156600.1.p | Glyma.10G156600 | bHLH |
| 1853. | Glyma.10G160000.1.p | Glyma.10G160000 | GeBP |
| 1854. | Glyma.10G160100.1.p | Glyma.10G160100 | GeBP |
| 1855. | Glyma.10G161200.1.p | Glyma.10G161200 | Trihelix |
| 1856. | Glyma.10G162100.1.p | Glyma.10G162100 | bZIP |
| 1857. | Glyma.10G162300.1.p | Glyma.10G162300 | bHLH |
| 1858. | Glyma.10G165800.1.p | Glyma.10G165800 | MYB |
| 1859. | Glyma.10G165900.1.p | Glyma.10G165900 | MYB |
| 1860. | Glyma.10G170800.1.p | Glyma.10G170800 | C3H |
| 1861. | Glyma.10G171000.1.p | Glyma.10G171000 | WRKY |
| 1862. | Glyma.10G171100.1.p | Glyma.10G171100 | WRKY |
| 1863. | Glyma.10G171200.1.p | Glyma.10G171200 | WRKY |
| 1864. | Glyma.10G171400.1.p | Glyma.10G171400 | AP2 |
| 1865. | Glyma.10G173500.1.p | Glyma.10G173500 | bHLH |
| 1866. | Glyma.10G173700.1.p | Glyma.10G173700 | Dof |
| 1867. | Glyma.10G180800.1.p | Glyma.10G180800 | MYB |
| 1868. | Glyma.10G182100.1.p | Glyma.10G182100 | C2H2 |
| 1869. | Glyma.10G184500.1.p | Glyma.10G184500 | C2H2 |
| 1870. | Glyma.10G186800.1.p | Glyma.10G186800 | ERF |
| 1871. | Glyma.10G186900.1.p | Glyma.10G186900 | ERF |
| 1872. | Glyma.10G187000.1.p | Glyma.10G187000 | ERF |
| 1873. | Glyma.10G190200.1.p | Glyma.10G190200 | GRAS |
| 1874. | Glyma.10G191000.1.p | Glyma.10G191000 | MYB |
| 1875. | Glyma.10G192000.1.p | Glyma.10G192000 | NF-YB |
| 1876. | Glyma.10G193400.1.p | Glyma.10G193400 | ERF |
| 1877. | Glyma.10G194200.1.p | Glyma.10G194200 | ERF |
| 1878. | Glyma.10G196600.1.p | Glyma.10G196600 | G2-like |
| 1879. | Glyma.10G197200.1.p | Glyma.10G197200 | C2H2 |
| 1880. | Glyma.10G197500.1.p | Glyma.10G197500 | NAC |
| 1881. | Glyma.10G197600.1.p | Glyma.10G197600 | NAC |
| 1882. | Glyma.10G201400.1.p | Glyma.10G201400 | Trihelix |
| 1883. | Glyma.10G202300.1.p | Glyma.10G202300 | Trihelix |
| 1884. | Glyma.10G204200.1.p | Glyma.10G204200 | G2-like |
| 1885. | Glyma.10G204300.1.p | Glyma.10G204300 | C2H2 |
| 1886. | Glyma.10G204400.1.p | Glyma.10G204400 | RAV |
| 1887. | Glyma.10G204700.1.p | Glyma.10G204700 | NAC |
| 1888. | Glyma.10G205000.1.p | Glyma.10G205000 | bHLH |
| 1889. | Glyma.10G206300.1.p | Glyma.10G206300 | bHLH |
| 1890. | Glyma.10G206400.1.p | Glyma.10G206400 | MYB |
| 1891. | Glyma.10G206500.1.p | Glyma.10G206500 | MYB |
| 1892. | Glyma.10G206600.1.p | Glyma.10G206600 | C2H2 |
| 1893. | Glyma.10G209600.1.p | Glyma.10G209600 | C2H2 |
| 1894. | Glyma.10G210500.1.p | Glyma.10G210500 | GATA |
| 1895. | Glyma.10G210600.1.p | Glyma.10G210600 | ARF |
| 1896. | Glyma.10G212600.1.p | Glyma.10G212600 | MYB |
| 1897. | Glyma.10G213500.1.p | Glyma.10G213500 | LBD |
| 1898. | Glyma.10G215000.1.p | Glyma.10G215000 | GRAS |
| 1899. | Glyma.10G215200.1.p | Glyma.10G215200 | C2H2 |
| 1900. | Glyma.10G216400.1.p | Glyma.10G216400 | NAC |
| 1901. | Glyma.10G219000.1.p | Glyma.10G219000 | ERF |
| 1902. | Glyma.10G219600.1.p | Glyma.10G219600 | NAC |
| 1903. | Glyma.10G223200.1.p | Glyma.10G223200 | ERF |
| 1904. | Glyma.10G223800.1.p | Glyma.10G223800 | bZIP |
| 1905. | Glyma.10G225100.1.p | Glyma.10G225100 | Trihelix |
| 1906. | Glyma.10G225200.1.p | Glyma.10G225200 | Trihelix |
| 1907. | Glyma.10G225300.1.p | Glyma.10G225300 | Trihelix |
| 1908. | Glyma.10G230200.1.p | Glyma.10G230200 | WRKY |
| 1909. | Glyma.10G230700.1.p | Glyma.10G230700 | MYB_related |
| 1910. | Glyma.10G232000.1.p | Glyma.10G232000 | GRAS |
| 1911. | Glyma.10G233400.1.p | Glyma.10G233400 | C2H2 |
| 1912. | Glyma.10G234100.1.p | Glyma.10G234100 | Nin-like |
| 1913. | Glyma.10G236400.1.p | Glyma.10G236400 | MYB |
| 1914. | Glyma.10G236600.1.p | Glyma.10G236600 | MYB |
| 1915. | Glyma.10G237800.1.p | Glyma.10G237800 | HSF |
| 1916. | Glyma.10G238100.1.p | Glyma.10G238100 | HD-ZIP |
| 1917. | Glyma.10G238500.1.p | Glyma.10G238500 | FAR1 |
| 1918. | Glyma.10G239300.1.p | Glyma.10G239300 | ERF |
| 1919. | Glyma.10G239400.1.p | Glyma.10G239400 | ERF |
| 1920. | Glyma.10G240200.1.p | Glyma.10G240200 | TCP |
| 1921. | Glyma.10G240500.1.p | Glyma.10G240500 | MIKC_MADS |
| 1922. | Glyma.10G240900.1.p | Glyma.10G240900 | MIKC_MADS |
| 1923. | Glyma.10G241300.1.p | Glyma.10G241300 | bHLH |
| 1924. | Glyma.10G241400.1.p | Glyma.10G241400 | bHLH |
| 1925. | Glyma.10G244000.1.p | Glyma.10G244000 | HSF |
| 1926. | Glyma.10G245500.1.p | Glyma.10G245500 | CPP |
| 1927. | Glyma.10G246200.1.p | Glyma.10G246200 | TCP |
| 1928. | Glyma.10G251300.1.p | Glyma.10G251300 | HD-ZIP |
| 1929. | Glyma.10G254600.1.p | Glyma.10G254600 | M-type_MADS |
| 1930. | Glyma.10G254700.1.p | Glyma.10G254700 | M-type_MADS |
| 1931. | Glyma.10G254800.1.p | Glyma.10G254800 | M-type_MADS |
| 1932. | Glyma.10G257400.1.p | Glyma.10G257400 | bHLH |
| 1933. | Glyma.10G257500.1.p | Glyma.10G257500 | bHLH |
| 1934. | Glyma.10G257900.1.p | Glyma.10G257900 | C2H2 |
| 1935. | Glyma.10G258100.1.p | Glyma.10G258100 | LBD |
| 1936. | Glyma.10G259000.1.p | Glyma.10G259000 | FAR1 |
| 1937. | Glyma.10G260400.1.p | Glyma.10G260400 | HB-PHD |
| 1938. | Glyma.10G261100.1.p | Glyma.10G261100 | bHLH |
| 1939. | Glyma.10G269300.1.p | Glyma.10G269300 | C3H |
| 1940. | Glyma.10G269600.1.p | Glyma.10G269600 | G2-like |
| 1941. | Glyma.10G273000.1.p | Glyma.10G273000 | MYB |
| 1942. | Glyma.10G274300.1.p | Glyma.10G274300 | CO-like |
| 1943. | Glyma.10G274600.1.p | Glyma.10G274600 | ERF |
| 1944. | Glyma.10G276100.1.p | Glyma.10G276100 | bZIP |
| 1945. | Glyma.10G277800.1.p | Glyma.10G277800 | MYB_related |
| 1946. | Glyma.10G280000.1.p | Glyma.10G280000 | C2H2 |
| 1947. | Glyma.10G281000.1.p | Glyma.10G281000 | G2-like |
| 1948. | Glyma.10G281100.1.p | Glyma.10G281100 | B3 |
| 1949. | Glyma.10G281200.1.p | Glyma.10G281200 | B3 |
| 1950. | Glyma.10G281300.1.p | Glyma.10G281300 | B3 |
| 1951. | Glyma.10G281800.1.p | Glyma.10G281800 | bHLH |
| 1952. | Glyma.10G285900.1.p | Glyma.10G285900 | TCP |
| 1953. | Glyma.10G289900.1.p | Glyma.10G289900 | WOX |
| 1954. | Glyma.10G295200.1.p | Glyma.10G295200 | C2H2 |
| 1955. | Glyma.10G296200.1.p | Glyma.10G296200 | bZIP |
| 1956. | Glyma.10G296700.1.p | Glyma.10G296700 | C3H |
| 1957. | Glyma.10G298700.1.p | Glyma.10G298700 | Trihelix |

| 1958. | Glyma.11G003400.1.p | Glyma.11G003400 | HD-ZIP |
| --- | --- | --- | --- |
| 1959. | Glyma.11G006900.1.p | Glyma.11G006900 | CPP |
| 1960. | Glyma.11G008500.1.p | Glyma.11G008500 | GRF |
| 1961. | Glyma.11G009400.1.p | Glyma.11G009400 | MYB |
| 1962. | Glyma.11G009800.1.p | Glyma.11G009800 | HSF |
| 1963. | Glyma.11G010800.1.p | Glyma.11G010800 | E2F/DP |
| 1964. | Glyma.11G010900.1.p | Glyma.11G010900 | MYB_related |
| 1965. | Glyma.11G014200.1.p | Glyma.11G014200 | ERF |
| 1966. | Glyma.11G014800.1.p | Glyma.11G014800 | ERF |
| 1967. | Glyma.11G016100.1.p | Glyma.11G016100 | GRAS |
| 1968. | Glyma.11G018100.1.p | Glyma.11G018100 | ERF |
| 1969. | Glyma.11G018200.1.p | Glyma.11G018200 | MYB_related |
| 1970. | Glyma.11G019000.1.p | Glyma.11G019000 | ERF |
| 1971. | Glyma.11G021200.1.p | Glyma.11G021200 | WRKY |
| 1972. | Glyma.11G021600.1.p | Glyma.11G021600 | MYB |
| 1973. | Glyma.11G022200.1.p | Glyma.11G022200 | TALE |
| 1974. | Glyma.11G025700.1.p | Glyma.11G025700 | HSF |
| 1975. | Glyma.11G027100.1.p | Glyma.11G027100 | TALE |
| 1976. | Glyma.11G029900.1.p | Glyma.11G029900 | M-type_MADS |
| 1977. | Glyma.11G030200.1.p | Glyma.11G030200 | MYB |
| 1978. | Glyma.11G030600.1.p | Glyma.11G030600 | NAC |
| 1979. | Glyma.11G033100.1.p | Glyma.11G033100 | C2H2 |
| 1980. | Glyma.11G034900.1.p | Glyma.11G034900 | MYB |
| 1981. | Glyma.11G035100.1.p | Glyma.11G035100 | ERF |
| 1982. | Glyma.11G035900.1.p | Glyma.11G035900 | HD-ZIP |
| 1983. | Glyma.11G036400.1.p | Glyma.11G036400 | ERF |
| 1984. | Glyma.11G036500.1.p | Glyma.11G036500 | ERF |
| 1985. | Glyma.11G037900.1.p | Glyma.11G037900 | GATA |
| 1986. | Glyma.11G041300.1.p | Glyma.11G041300 | ARR-B |
| 1987. | Glyma.11G043200.1.p | Glyma.11G043200 | C2H2 |
| 1988. | Glyma.11G043600.1.p | Glyma.11G043600 | bHLH |
| 1989. | Glyma.11G043700.1.p | Glyma.11G043700 | bHLH |
| 1990. | Glyma.11G045100.1.p | Glyma.11G045100 | HD-ZIP |
| 1991. | Glyma.11G045400.1.p | Glyma.11G045400 | MYB |
| 1992. | Glyma.11G045800.1.p | Glyma.11G045800 | AP2 |
| 1993. | Glyma.11G045900.1.p | Glyma.11G045900 | bZIP |
| 1994. | Glyma.11G047300.1.p | Glyma.11G047300 | MYB_related |
| 1995. | Glyma.11G047700.1.p | Glyma.11G047700 | GRAS |
| 1996. | Glyma.11G049300.1.p | Glyma.11G049300 | LBD |
| 1997. | Glyma.11G052100.1.p | Glyma.11G052100 | MYB |
| 1998. | Glyma.11G053100.1.p | Glyma.11G053100 | WRKY |
| 1999. | Glyma.11G053600.1.p | Glyma.11G053600 | ERF |
| 2000. | Glyma.11G053800.1.p | Glyma.11G053800 | AP2 |
| 2001. | Glyma.11G054600.1.p | Glyma.11G054600 | bHLH |
| 2002. | Glyma.11G055300.1.p | Glyma.11G055300 | bHLH |
| 2003. | Glyma.11G056200.1.p | Glyma.11G056200 | HSF |
| 2004. | Glyma.11G058400.1.p | Glyma.11G058400 | LBD |
| 2005. | Glyma.11G058600.1.p | Glyma.11G058600 | G2-like |
| 2006. | Glyma.11G059300.1.p | Glyma.11G059300 | Dof |
| 2007. | Glyma.11G062300.1.p | Glyma.11G062300 | TALE |
| 2008. | Glyma.11G064300.1.p | Glyma.11G064300 | BES1 |
| 2009. | Glyma.11G064800.1.p | Glyma.11G064800 | HD-ZIP |
| 2010. | Glyma.11G065000.1.p | Glyma.11G065000 | bZIP |
| 2011. | Glyma.11G065200.1.p | Glyma.11G065200 | GRAS |
| 2012. | Glyma.11G068400.1.p | Glyma.11G068400 | MYB_related |
| 2013. | Glyma.11G068700.1.p | Glyma.11G068700 | GATA |
| 2014. | Glyma.11G068800.1.p | Glyma.11G068800 | ZF-HD |
| 2015. | Glyma.11G069100.1.p | Glyma.11G069100 | MYB_related |
| 2016. | Glyma.11G072500.1.p | Glyma.11G072500 | SRS |
| 2017. | Glyma.11G073700.1.p | Glyma.11G073700 | MIKC_MADS |
| 2018. | Glyma.11G074900.1.p | Glyma.11G074900 | DBB |
| 2019. | Glyma.11G075400.1.p | Glyma.11G075400 | NAC |
| 2020. | Glyma.11G075900.1.p | Glyma.11G075900 | C3H |
| 2021. | Glyma.11G076500.1.p | Glyma.11G076500 | WOX |
| 2022. | Glyma.11G083900.1.p | Glyma.11G083900 | SBP |
| 2023. | Glyma.11G085500.1.p | Glyma.11G085500 | Nin-like |
| 2024. | Glyma.11G088700.1.p | Glyma.11G088700 | FAR1 |
| 2025. | Glyma.11G091900.1.p | Glyma.11G091900 | E2F/DP |
| 2026. | Glyma.11G092000.1.p | Glyma.11G092000 | GRAS |
| 2027. | Glyma.11G096000.1.p | Glyma.11G096000 | GRAS |
| 2028. | Glyma.11G096500.1.p | Glyma.11G096500 | GRAS |
| 2029. | Glyma.11G096600.1.p | Glyma.11G096600 | NAC |
| 2030. | Glyma.11G107100.1.p | Glyma.11G107100 | MYB |
| 2031. | Glyma.11G108100.1.p | Glyma.11G108100 | bZIP |
| 2032. | Glyma.11G108400.1.p | Glyma.11G108400 | MYB |
| 2033. | Glyma.11G109000.1.p | Glyma.11G109000 | C3H |
| 2034. | Glyma.11G110400.1.p | Glyma.11G110400 | bZIP |
| 2035. | Glyma.11G110700.1.p | Glyma.11G110700 | GRF |
| 2036. | Glyma.11G110900.1.p | Glyma.11G110900 | DBB |
| 2037. | Glyma.11G111700.1.p | Glyma.11G111700 | GATA |
| 2038. | Glyma.11G112800.1.p | Glyma.11G112800 | DBB |
| 2039. | Glyma.11G113200.1.p | Glyma.11G113200 | SRS |
| 2040. | Glyma.11G114800.1.p | Glyma.11G114800 | bZIP |
| 2041. | Glyma.11G117100.1.p | Glyma.11G117100 | bHLH |
| 2042. | Glyma.11G117200.1.p | Glyma.11G117200 | MYB_related |
| 2043. | Glyma.11G124100.1.p | Glyma.11G124100 | B3 |
| 2044. | Glyma.11G124200.1.p | Glyma.11G124200 | B3 |
| 2045. | Glyma.11G124300.1.p | Glyma.11G124300 | B3 |
| 2046. | Glyma.11G125200.1.p | Glyma.11G125200 | B3 |
| 2047. | Glyma.11G125500.1.p | Glyma.11G125500 | Nin-like |
| 2048. | Glyma.11G126600.1.p | Glyma.11G126600 | bZIP |
| 2049. | Glyma.11G126700.1.p | Glyma.11G126700 | FAR1 |
| 2050. | Glyma.11G127100.1.p | Glyma.11G127100 | DBB |
| 2051. | Glyma.11G127400.1.p | Glyma.11G127400 | FAR1 |
| 2052. | Glyma.11G131200.1.p | Glyma.11G131200 | bHLH |
| 2053. | Glyma.11G131900.1.p | Glyma.11G131900 | AP2 |
| 2054. | Glyma.11G132600.1.p | Glyma.11G132600 | C2H2 |
| 2055. | Glyma.11G133700.1.p | Glyma.11G133700 | MYB |
| 2056. | Glyma.11G135800.1.p | Glyma.11G135800 | C2H2 |
| 2057. | Glyma.11G136600.1.p | Glyma.11G136600 | G2-like |
| 2058. | Glyma.11G138000.1.p | Glyma.11G138000 | GRAS |
| 2059. | Glyma.11G138100.1.p | Glyma.11G138100 | GRAS |
| 2060. | Glyma.11G138200.1.p | Glyma.11G138200 | GRAS |
| 2061. | Glyma.11G138300.1.p | Glyma.11G138300 | GRAS |
| 2062. | Glyma.11G138400.1.p | Glyma.11G138400 | GRAS |
| 2063. | Glyma.11G138500.1.p | Glyma.11G138500 | GRAS |
| 2064. | Glyma.11G138600.1.p | Glyma.11G138600 | GRAS |
| 2065. | Glyma.11G138700.1.p | Glyma.11G138700 | MYB_related |
| 2066. | Glyma.11G140200.1.p | Glyma.11G140200 | Dof |
| 2067. | Glyma.11G140400.1.p | Glyma.11G140400 | WOX |
| 2068. | Glyma.11G142500.1.p | Glyma.11G142500 | C2H2 |
| 2069. | Glyma.11G142900.1.p | Glyma.11G142900 | MYB |
| 2070. | Glyma.11G145400.1.p | Glyma.11G145400 | GATA |
| 2071. | Glyma.11G145500.1.p | Glyma.11G145500 | ARF |
| 2072. | Glyma.11G145800.1.p | Glyma.11G145800 | HD-ZIP |
| 2073. | Glyma.11G147200.1.p | Glyma.11G147200 | C3H |
| 2074. | Glyma.11G148000.1.p | Glyma.11G148000 | NF-YC |
| 2075. | Glyma.11G150200.1.p | Glyma.11G150200 | GRAS |
| 2076. | Glyma.11G155400.1.p | Glyma.11G155400 | SRS |
| 2077. | Glyma.11G156200.1.p | Glyma.11G156200 | GATA |
| 2078. | Glyma.11G156700.1.p | Glyma.11G156700 | Trihelix |
| 2079. | Glyma.11G159700.1.p | Glyma.11G159700 | M-type_MADS |
| 2080. | Glyma.11G160200.1.p | Glyma.11G160200 | M-type_MADS |
| 2081. | Glyma.11G163300.1.p | Glyma.11G163300 | WRKY |
| 2082. | Glyma.11G164800.1.p | Glyma.11G164800 | LBD |
| 2083. | Glyma.11G165800.1.p | Glyma.11G165800 | FAR1 |
| 2084. | Glyma.11G167400.1.p | Glyma.11G167400 | bZIP |
| 2085. | Glyma.11G167600.1.p | Glyma.11G167600 | bZIP |
| 2086. | Glyma.11G167900.1.p | Glyma.11G167900 | bZIP |
| 2087. | Glyma.11G174400.1.p | Glyma.11G174400 | C3H |
| 2088. | Glyma.11G176500.1.p | Glyma.11G176500 | MYB |
| 2089. | Glyma.11G177100.1.p | Glyma.11G177100 | GRAS |
| 2090. | Glyma.11G180100.1.p | Glyma.11G180100 | E2F/DP |
| 2091. | Glyma.11G182000.1.p | Glyma.11G182000 | NAC |
| 2092. | Glyma.11G183100.1.p | Glyma.11G183100 | NF-YB |
| 2093. | Glyma.11G183400.1.p | Glyma.11G183400 | G2-like |
| 2094. | Glyma.11G183700.1.p | Glyma.11G183700 | bZIP |
| 2095. | Glyma.11G184000.1.p | Glyma.11G184000 | C2H2 |
| 2096. | Glyma.11G185900.1.p | Glyma.11G185900 | M-type_MADS |
| 2097. | Glyma.11G186000.1.p | Glyma.11G186000 | M-type_MADS |
| 2098. | Glyma.11G186100.1.p | Glyma.11G186100 | M-type_MADS |
| 2099. | Glyma.11G186400.1.p | Glyma.11G186400 | M-type_MADS |
| 2100. | Glyma.11G186600.1.p | Glyma.11G186600 | MYB_related |
| 2101. | Glyma.11G186700.1.p | Glyma.11G186700 | bZIP |
| 2102. | Glyma.11G188200.1.p | Glyma.11G188200 | ERF |
| 2103. | Glyma.11G189500.1.p | Glyma.11G189500 | C2H2 |
| 2104. | Glyma.11G192400.1.p | Glyma.11G192400 | C2H2 |
| 2105. | Glyma.11G192600.1.p | Glyma.11G192600 | C2H2 |
| 2106. | Glyma.11G192800.1.p | Glyma.11G192800 | bHLH |
| 2107. | Glyma.11G194100.1.p | Glyma.11G194100 | MYB |
| 2108. | Glyma.11G196000.1.p | Glyma.11G196000 | TCP |
| 2109. | Glyma.11G196300.1.p | Glyma.11G196300 | TALE |
| 2110. | Glyma.11G197900.1.p | Glyma.11G197900 | B3 |
| 2111. | Glyma.11G198800.1.p | Glyma.11G198800 | B3 |
| 2112. | Glyma.11G199300.1.p | Glyma.11G199300 | ERF |
| 2113. | Glyma.11G204200.1.p | Glyma.11G204200 | ARF |
| 2114. | Glyma.11G208800.1.p | Glyma.11G208800 | GRF |
| 2115. | Glyma.11G210800.1.p | Glyma.11G210800 | WOX |
| 2116. | Glyma.11G211900.1.p | Glyma.11G211900 | VOZ |
| 2117. | Glyma.11G212400.1.p | Glyma.11G212400 | NAC |
| 2118. | Glyma.11G213700.1.p | Glyma.11G213700 | G2-like |
| 2119. | Glyma.11G214300.1.p | Glyma.11G214300 | C2H2 |
| 2120. | Glyma.11G214600.1.p | Glyma.11G214600 | M-type_MADS |
| 2121. | Glyma.11G215800.1.p | Glyma.11G215800 | MYB |
| 2122. | Glyma.11G216500.1.p | Glyma.11G216500 | GRAS |
| 2123. | Glyma.11G217700.1.p | Glyma.11G217700 | bHLH |
| 2124. | Glyma.11G221500.1.p | Glyma.11G221500 | C2H2 |
| 2125. | Glyma.11G227200.1.p | Glyma.11G227200 | MYB |
| 2126. | Glyma.11G227800.1.p | Glyma.11G227800 | WOX |
| 2127. | Glyma.11G230200.1.p | Glyma.11G230200 | NF-X1 |
| 2128. | Glyma.11G231300.1.p | Glyma.11G231300 | bHLH |
| 2129. | Glyma.11G231500.1.p | Glyma.11G231500 | LBD |
| 2130. | Glyma.11G236300.1.p | Glyma.11G236300 | bZIP |
| 2131. | Glyma.11G239000.1.p | Glyma.11G239000 | EIL |
| 2132. | Glyma.11G239200.1.p | Glyma.11G239200 | ERF |
| 2133. | Glyma.11G240600.1.p | Glyma.11G240600 | C2H2 |
| 2134. | Glyma.11G242200.1.p | Glyma.11G242200 | HD-ZIP |
| 2135. | Glyma.11G246400.1.p | Glyma.11G246400 | ARR-B |
| 2136. | Glyma.11G247300.1.p | Glyma.11G247300 | Trihelix |
| 2137. | Glyma.11G250000.1.p | Glyma.11G250000 | NF-YC |
| 2138. | Glyma.11G251500.1.p | Glyma.11G251500 | SBP |
| 2139. | Glyma.11G251900.1.p | Glyma.11G251900 | CAMTA |
| 2140. | Glyma.11G252300.1.p | Glyma.11G252300 | MIKC_MADS |
| 2141. | Glyma.11G253400.1.p | Glyma.11G253400 | C2H2 |

| 2142. | Glyma.12G003200.1.p | Glyma.12G003200 | NAC |
| --- | --- | --- | --- |
| 2143. | Glyma.12G004900.1.p | Glyma.12G004900 | NAC |
| 2144. | Glyma.12G005000.1.p | Glyma.12G005000 | MIKC_MADS |
| 2145. | Glyma.12G011500.1.p | Glyma.12G011500 | FAR1 |
| 2146. | Glyma.12G012100.1.p | Glyma.12G012100 | GRAS |
| 2147. | Glyma.12G014700.1.p | Glyma.12G014700 | GRF |
| 2148. | Glyma.12G016400.1.p | Glyma.12G016400 | C3H |
| 2149. | Glyma.12G017000.1.p | Glyma.12G017000 | MYB |
| 2150. | Glyma.12G018000.1.p | Glyma.12G018000 | E2F/DP |
| 2151. | Glyma.12G018100.1.p | Glyma.12G018100 | GRAS |
| 2152. | Glyma.12G022100.1.p | Glyma.12G022100 | GRAS |
| 2153. | Glyma.12G022600.1.p | Glyma.12G022600 | GRAS |
| 2154. | Glyma.12G022700.1.p | Glyma.12G022700 | NAC |
| 2155. | Glyma.12G024400.1.p | Glyma.12G024400 | bHLH |
| 2156. | Glyma.12G032200.1.p | Glyma.12G032200 | MYB |
| 2157. | Glyma.12G033000.1.p | Glyma.12G033000 | bZIP |
| 2158. | Glyma.12G036400.1.p | Glyma.12G036400 | bZIP |
| 2159. | Glyma.12G037200.1.p | Glyma.12G037200 | DBB |
| 2160. | Glyma.12G037700.1.p | Glyma.12G037700 | GATA |
| 2161. | Glyma.12G039100.1.p | Glyma.12G039100 | SRS |
| 2162. | Glyma.12G040600.1.p | Glyma.12G040600 | bZIP |
| 2163. | Glyma.12G042800.1.p | Glyma.12G042800 | bHLH |
| 2164. | Glyma.12G042900.1.p | Glyma.12G042900 | MYB_related |
| 2165. | Glyma.12G045100.1.p | Glyma.12G045100 | bZIP |
| 2166. | Glyma.12G048500.1.p | Glyma.12G048500 | B3 |
| 2167. | Glyma.12G048600.1.p | Glyma.12G048600 | B3 |
| 2168. | Glyma.12G048700.1.p | Glyma.12G048700 | B3 |
| 2169. | Glyma.12G049900.1.p | Glyma.12G049900 | B3 |
| 2170. | Glyma.12G050100.1.p | Glyma.12G050100 | Nin-like |
| 2171. | Glyma.12G051200.1.p | Glyma.12G051200 | bZIP |
| 2172. | Glyma.12G051700.1.p | Glyma.12G051700 | DBB |
| 2173. | Glyma.12G052000.1.p | Glyma.12G052000 | FAR1 |
| 2174. | Glyma.12G055400.1.p | Glyma.12G055400 | bHLH |
| 2175. | Glyma.12G056300.1.p | Glyma.12G056300 | AP2 |
| 2176. | Glyma.12G057000.1.p | Glyma.12G057000 | C2H2 |
| 2177. | Glyma.12G057900.1.p | Glyma.12G057900 | MYB |
| 2178. | Glyma.12G059500.1.p | Glyma.12G059500 | C2H2 |
| 2179. | Glyma.12G060200.1.p | Glyma.12G060200 | G2-like |
| 2180. | Glyma.12G061900.1.p | Glyma.12G061900 | GRAS |
| 2181. | Glyma.12G062000.1.p | Glyma.12G062000 | GRAS |
| 2182. | Glyma.12G062100.1.p | Glyma.12G062100 | GRAS |
| 2183. | Glyma.12G062200.1.p | Glyma.12G062200 | GRAS |
| 2184. | Glyma.12G062300.1.p | Glyma.12G062300 | MYB_related |
| 2185. | Glyma.12G063800.1.p | Glyma.12G063800 | Dof |
| 2186. | Glyma.12G063900.1.p | Glyma.12G063900 | WOX |
| 2187. | Glyma.12G066000.1.p | Glyma.12G066000 | MYB |
| 2188. | Glyma.12G068100.1.p | Glyma.12G068100 | C3H |
| 2189. | Glyma.12G069100.2.p | Glyma.12G069100 | NF-YC |
| 2190. | Glyma.12G070300.1.p | Glyma.12G070300 | C2H2 |
| 2191. | Glyma.12G071000.1.p | Glyma.12G071000 | ARF |
| 2192. | Glyma.12G072000.1.p | Glyma.12G072000 | LBD |
| 2193. | Glyma.12G072400.1.p | Glyma.12G072400 | Dof |
| 2194. | Glyma.12G073300.1.p | Glyma.12G073300 | AP2 |
| 2195. | Glyma.12G075800.1.p | Glyma.12G075800 | HD-ZIP |
| 2196. | Glyma.12G076200.1.p | Glyma.12G076200 | ARF |
| 2197. | Glyma.12G076500.1.p | Glyma.12G076500 | GATA |
| 2198. | Glyma.12G077800.1.p | Glyma.12G077800 | TALE |
| 2199. | Glyma.12G079800.1.p | Glyma.12G079800 | MYB |
| 2200. | Glyma.12G081200.1.p | Glyma.12G081200 | bHLH |
| 2201. | Glyma.12G081400.1.p | Glyma.12G081400 | C2H2 |
| 2202. | Glyma.12G081700.1.p | Glyma.12G081700 | C2H2 |
| 2203. | Glyma.12G084700.1.p | Glyma.12G084700 | C2H2 |
| 2204. | Glyma.12G086200.1.p | Glyma.12G086200 | ERF |
| 2205. | Glyma.12G088400.1.p | Glyma.12G088400 | C2H2 |
| 2206. | Glyma.12G088700.1.p | Glyma.12G088700 | bZIP |
| 2207. | Glyma.12G089100.1.p | Glyma.12G089100 | G2-like |
| 2208. | Glyma.12G089400.1.p | Glyma.12G089400 | NF-YB |
| 2209. | Glyma.12G091200.1.p | Glyma.12G091200 | NAC |
| 2210. | Glyma.12G093400.1.p | Glyma.12G093400 | E2F/DP |
| 2211. | Glyma.12G094800.1.p | Glyma.12G094800 | TALE |
| 2212. | Glyma.12G096000.1.p | Glyma.12G096000 | YABBY |
| 2213. | Glyma.12G097100.1.p | Glyma.12G097100 | WRKY |
| 2214. | Glyma.12G100100.1.p | Glyma.12G100100 | HD-ZIP |
| 2215. | Glyma.12G100600.1.p | Glyma.12G100600 | MYB_related |
| 2216. | Glyma.12G103100.1.p | Glyma.12G103100 | ERF |
| 2217. | Glyma.12G104500.1.p | Glyma.12G104500 | MYB |
| 2218. | Glyma.12G104600.1.p | Glyma.12G104600 | MYB |
| 2219. | Glyma.12G104800.1.p | Glyma.12G104800 | MYB |
| 2220. | Glyma.12G105400.1.p | Glyma.12G105400 | MYB |
| 2221. | Glyma.12G106400.1.p | Glyma.12G106400 | MYB |
| 2222. | Glyma.12G110400.1.p | Glyma.12G110400 | ERF |
| 2223. | Glyma.12G114300.1.p | Glyma.12G114300 | bHLH |
| 2224. | Glyma.12G114800.1.p | Glyma.12G114800 | MYB_related |
| 2225. | Glyma.12G116900.1.p | Glyma.12G116900 | C3H |
| 2226. | Glyma.12G117000.1.p | Glyma.12G117000 | ERF |
| 2227. | Glyma.12G117700.1.p | Glyma.12G117700 | G2-like |
| 2228. | Glyma.12G118000.1.p | Glyma.12G118000 | G2-like |
| 2229. | Glyma.12G118100.1.p | Glyma.12G118100 | MIKC_MADS |
| 2230. | Glyma.12G118700.1.p | Glyma.12G118700 | NAC |
| 2231. | Glyma.12G118800.1.p | Glyma.12G118800 | C2H2 |
| 2232. | Glyma.12G120100.1.p | Glyma.12G120100 | VOZ |
| 2233. | Glyma.12G120700.1.p | Glyma.12G120700 | LBD |
| 2234. | Glyma.12G121000.1.p | Glyma.12G121000 | bZIP |
| 2235. | Glyma.12G121500.1.p | Glyma.12G121500 | TCP |
| 2236. | Glyma.12G122800.1.p | Glyma.12G122800 | bHLH |
| 2237. | Glyma.12G136300.1.p | Glyma.12G136300 | bHLH |
| 2238. | Glyma.12G137700.1.p | Glyma.12G137700 | GRAS |
| 2239. | Glyma.12G141700.1.p | Glyma.12G141700 | bHLH |
| 2240. | Glyma.12G145100.1.p | Glyma.12G145100 | NAC |
| 2241. | Glyma.12G148900.1.p | Glyma.12G148900 | NAC |
| 2242. | Glyma.12G149100.1.p | Glyma.12G149100 | NAC |
| 2243. | Glyma.12G152600.1.p | Glyma.12G152600 | WRKY |
| 2244. | Glyma.12G158000.1.p | Glyma.12G158000 | HB-other |
| 2245. | Glyma.12G158200.1.p | Glyma.12G158200 | FAR1 |
| 2246. | Glyma.12G158400.1.p | Glyma.12G158400 | HD-ZIP |
| 2247. | Glyma.12G158900.1.p | Glyma.12G158900 | TCP |
| 2248. | Glyma.12G160100.1.p | Glyma.12G160100 | NAC |
| 2249. | Glyma.12G161700.1.p | Glyma.12G161700 | NAC |
| 2250. | Glyma.12G162700.1.p | Glyma.12G162700 | ERF |
| 2251. | Glyma.12G164100.1.p | Glyma.12G164100 | ARF |
| 2252. | Glyma.12G165800.1.p | Glyma.12G165800 | ZF-HD |
| 2253. | Glyma.12G168300.1.p | Glyma.12G168300 | TCP |
| 2254. | Glyma.12G168600.1.p | Glyma.12G168600 | ZF-HD |
| 2255. | Glyma.12G168700.1.p | Glyma.12G168700 | ZF-HD |
| 2256. | Glyma.12G168800.1.p | Glyma.12G168800 | MYB |
| 2257. | Glyma.12G171000.1.p | Glyma.12G171000 | ARF |
| 2258. | Glyma.12G171600.1.p | Glyma.12G171600 | NAC |
| 2259. | Glyma.12G171700.1.p | Glyma.12G171700 | C2H2 |
| 2260. | Glyma.12G174100.1.p | Glyma.12G174100 | ARF |
| 2261. | Glyma.12G174200.1.p | Glyma.12G174200 | GATA |
| 2262. | Glyma.12G176400.1.p | Glyma.12G176400 | TALE |
| 2263. | Glyma.12G177700.1.p | Glyma.12G177700 | MYB |
| 2264. | Glyma.12G178500.1.p | Glyma.12G178500 | bHLH |
| 2265. | Glyma.12G178900.1.p | Glyma.12G178900 | C2H2 |
| 2266. | Glyma.12G179000.1.p | Glyma.12G179000 | C2H2 |
| 2267. | Glyma.12G181400.1.p | Glyma.12G181400 | C2H2 |
| 2268. | Glyma.12G182200.1.p | Glyma.12G182200 | ERF |
| 2269. | Glyma.12G182400.1.p | Glyma.12G182400 | ERF |
| 2270. | Glyma.12G183900.1.p | Glyma.12G183900 | C2H2 |
| 2271. | Glyma.12G184400.1.p | Glyma.12G184400 | bZIP |
| 2272. | Glyma.12G184500.1.p | Glyma.12G184500 | bZIP |
| 2273. | Glyma.12G184700.1.p | Glyma.12G184700 | G2-like |
| 2274. | Glyma.12G186200.1.p | Glyma.12G186200 | NAC |
| 2275. | Glyma.12G186900.1.p | Glyma.12G186900 | NAC |
| 2276. | Glyma.12G188800.1.p | Glyma.12G188800 | TALE |
| 2277. | Glyma.12G190500.1.p | Glyma.12G190500 | YABBY |
| 2278. | Glyma.12G193300.1.p | Glyma.12G193300 | MYB |
| 2279. | Glyma.12G194400.1.p | Glyma.12G194400 | HD-ZIP |
| 2280. | Glyma.12G195200.1.p | Glyma.12G195200 | MYB_related |
| 2281. | Glyma.12G196700.1.p | Glyma.12G196700 | bHLH |
| 2282. | Glyma.12G196900.1.p | Glyma.12G196900 | bHLH |
| 2283. | Glyma.12G197300.1.p | Glyma.12G197300 | GRAS |
| 2284. | Glyma.12G197700.1.p | Glyma.12G197700 | ERF |
| 2285. | Glyma.12G199100.1.p | Glyma.12G199100 | MYB |
| 2286. | Glyma.12G199200.1.p | Glyma.12G199200 | MYB |
| 2287. | Glyma.12G199600.1.p | Glyma.12G199600 | MYB |
| 2288. | Glyma.12G203100.1.p | Glyma.12G203100 | ERF |
| 2289. | Glyma.12G205200.1.p | Glyma.12G205200 | bHLH |
| 2290. | Glyma.12G205700.1.p | Glyma.12G205700 | C3H |
| 2291. | Glyma.12G206600.1.p | Glyma.12G206600 | G2-like |
| 2292. | Glyma.12G206900.1.p | Glyma.12G206900 | NAC |
| 2293. | Glyma.12G207200.1.p | Glyma.12G207200 | C2H2 |
| 2294. | Glyma.12G208400.1.p | Glyma.12G208400 | bZIP |
| 2295. | Glyma.12G208800.1.p | Glyma.12G208800 | TCP |
| 2296. | Glyma.12G209900.1.p | Glyma.12G209900 | bHLH |
| 2297. | Glyma.12G210900.1.p | Glyma.12G210900 | Trihelix |
| 2298. | Glyma.12G211600.1.p | Glyma.12G211600 | G2-like |
| 2299. | Glyma.12G212300.1.p | Glyma.12G212300 | WRKY |
| 2300. | Glyma.12G213900.1.p | Glyma.12G213900 | MYB_related |
| 2301. | Glyma.12G214900.1.p | Glyma.12G214900 | bHLH |
| 2302. | Glyma.12G216100.1.p | Glyma.12G216100 | GRAS |
| 2303. | Glyma.12G216800.1.p | Glyma.12G216800 | bHLH |
| 2304. | Glyma.12G217200.1.p | Glyma.12G217200 | NF-YC |
| 2305. | Glyma.12G218200.1.p | Glyma.12G218200 | MYB |
| 2306. | Glyma.12G221400.1.p | Glyma.12G221400 | NAC |
| 2307. | Glyma.12G221500.1.p | Glyma.12G221500 | NAC |
| 2308. | Glyma.12G226000.1.p | Glyma.12G226000 | SBP |
| 2309. | Glyma.12G226500.1.p | Glyma.12G226500 | NAC |
| 2310. | Glyma.12G226600.1.p | Glyma.12G226600 | ERF |
| 2311. | Glyma.12G227700.1.p | Glyma.12G227700 | bZIP |
| 2312. | Glyma.12G228300.1.p | Glyma.12G228300 | TCP |
| 2313. | Glyma.12G230200.1.p | Glyma.12G230200 | C2H2 |
| 2314. | Glyma.12G231400.1.p | Glyma.12G231400 | BES1 |
| 2315. | Glyma.12G231500.1.p | Glyma.12G231500 | BES1 |
| 2316. | Glyma.12G233900.1.p | Glyma.12G233900 | DBB |
| 2317. | Glyma.12G236800.1.p | Glyma.12G236800 | NF-YA |
| 2318. | Glyma.12G237300.1.p | Glyma.12G237300 | C3H |
| 2319. | Glyma.12G237500.1.p | Glyma.12G237500 | MYB |
| 2320. | Glyma.12G237800.1.p | Glyma.12G237800 | C2H2 |
| 2321. | Glyma.12G238700.1.p | Glyma.12G238700 | bHLH |
| 2322. | Glyma.12G241400.1.p | Glyma.12G241400 | MYB |

| 2323. | Glyma.13G000700.1.p | Glyma.13G000700 | FAR1 |
| --- | --- | --- | --- |
| 2324. | Glyma.13G003600.1.p | Glyma.13G003600 | NF-YB |
| 2325. | Glyma.13G005900.1.p | Glyma.13G005900 | MYB_related |
| 2326. | Glyma.13G006100.1.p | Glyma.13G006100 | MYB_related |
| 2327. | Glyma.13G009300.1.p | Glyma.13G009300 | CO-like |
| 2328. | Glyma.13G030000.1.p | Glyma.13G030000 | FAR1 |
| 2329. | Glyma.13G030900.1.p | Glyma.13G030900 | NAC |
| 2330. | Glyma.13G032200.1.p | Glyma.13G032200 | MYB |
| 2331. | Glyma.13G034100.1.p | Glyma.13G034100 | MIKC_MADS |
| 2332. | Glyma.13G037100.1.p | Glyma.13G037100 | GRAS |
| 2333. | Glyma.13G038100.1.p | Glyma.13G038100 | MYB_related |
| 2334. | Glyma.13G038200.1.p | Glyma.13G038200 | MYB_related |
| 2335. | Glyma.13G038500.1.p | Glyma.13G038500 | MYB |
| 2336. | Glyma.13G040100.1.p | Glyma.13G040100 | bHLH |
| 2337. | Glyma.13G040400.1.p | Glyma.13G040400 | ERF |
| 2338. | Glyma.13G045500.1.p | Glyma.13G045500 | M-type_MADS |
| 2339. | Glyma.13G046300.1.p | Glyma.13G046300 | LBD |
| 2340. | Glyma.13G047400.1.p | Glyma.13G047400 | TCP |
| 2341. | Glyma.13G050300.1.p | Glyma.13G050300 | CO-like |
| 2342. | Glyma.13G050400.1.p | Glyma.13G050400 | MYB |
| 2343. | Glyma.13G050700.1.p | Glyma.13G050700 | bZIP |
| 2344. | Glyma.13G052100.1.p | Glyma.13G052100 | MIKC_MADS |
| 2345. | Glyma.13G052700.1.p | Glyma.13G052700 | MIKC_MADS |
| 2346. | Glyma.13G058500.1.p | Glyma.13G058500 | B3 |
| 2347. | Glyma.13G060600.1.p | Glyma.13G060600 | ERF |
| 2348. | Glyma.13G061900.1.p | Glyma.13G061900 | MYB |
| 2349. | Glyma.13G062000.1.p | Glyma.13G062000 | NAC |
| 2350. | Glyma.13G062500.1.p | Glyma.13G062500 | Dof |
| 2351. | Glyma.13G062900.1.p | Glyma.13G062900 | ZF-HD |
| 2352. | Glyma.13G063100.1.p | Glyma.13G063100 | ZF-HD |
| 2353. | Glyma.13G063200.1.p | Glyma.13G063200 | MYB |
| 2354. | Glyma.13G063300.1.p | Glyma.13G063300 | NAC |
| 2355. | Glyma.13G063800.1.p | Glyma.13G063800 | bHLH |
| 2356. | Glyma.13G064000.1.p | Glyma.13G064000 | HD-ZIP |
| 2357. | Glyma.13G066600.1.p | Glyma.13G066600 | MYB |
| 2358. | Glyma.13G071700.1.p | Glyma.13G071700 | MYB |
| 2359. | Glyma.13G073300.1.p | Glyma.13G073300 | C2H2 |
| 2360. | Glyma.13G073400.1.p | Glyma.13G073400 | MYB |
| 2361. | Glyma.13G075000.1.p | Glyma.13G075000 | bZIP |
| 2362. | Glyma.13G076700.1.p | Glyma.13G076700 | EIL |
| 2363. | Glyma.13G076800.1.p | Glyma.13G076800 | EIL |
| 2364. | Glyma.13G081600.1.p | Glyma.13G081600 | ERF |
| 2365. | Glyma.13G081700.1.p | Glyma.13G081700 | GRAS |
| 2366. | Glyma.13G084700.1.p | Glyma.13G084700 | ARF |
| 2367. | Glyma.13G085100.1.p | Glyma.13G085100 | bZIP |
| 2368. | Glyma.13G086400.1.p | Glyma.13G086400 | MIKC_MADS |
| 2369. | Glyma.13G088100.1.p | Glyma.13G088100 | ERF |
| 2370. | Glyma.13G090000.1.p | Glyma.13G090000 | C2H2 |
| 2371. | Glyma.13G093800.1.p | Glyma.13G093800 | CO-like |
| 2372. | Glyma.13G094400.1.p | Glyma.13G094400 | MYB |
| 2373. | Glyma.13G095500.1.p | Glyma.13G095500 | C2H2 |
| 2374. | Glyma.13G096900.1.p | Glyma.13G096900 | AP2 |
| 2375. | Glyma.13G100500.1.p | Glyma.13G100500 | HB-PHD |
| 2376. | Glyma.13G101100.1.p | Glyma.13G101100 | bHLH |
| 2377. | Glyma.13G102000.1.p | Glyma.13G102000 | WRKY |
| 2378. | Glyma.13G102800.1.p | Glyma.13G102800 | HD-ZIP |
| 2379. | Glyma.13G103900.1.p | Glyma.13G103900 | GATA |
| 2380. | Glyma.13G105400.1.p | Glyma.13G105400 | bHLH |
| 2381. | Glyma.13G105700.1.p | Glyma.13G105700 | HSF |
| 2382. | Glyma.13G107900.1.p | Glyma.13G107900 | NF-YA |
| 2383. | Glyma.13G109100.1.p | Glyma.13G109100 | MYB |
| 2384. | Glyma.13G109500.1.p | Glyma.13G109500 | GRF |
| 2385. | Glyma.13G112400.1.p | Glyma.13G112400 | ERF |
| 2386. | Glyma.13G112600.1.p | Glyma.13G112600 | ARF |
| 2387. | Glyma.13G117600.1.p | Glyma.13G117600 | WRKY |
| 2388. | Glyma.13G120700.1.p | Glyma.13G120700 | bHLH |
| 2389. | Glyma.13G120900.1.p | Glyma.13G120900 | C2H2 |
| 2390. | Glyma.13G121000.1.p | Glyma.13G121000 | C2H2 |
| 2391. | Glyma.13G121300.1.p | Glyma.13G121300 | LBD |
| 2392. | Glyma.13G121400.1.p | Glyma.13G121400 | LBD |
| 2393. | Glyma.13G122500.1.p | Glyma.13G122500 | ERF |
| 2394. | Glyma.13G122600.1.p | Glyma.13G122600 | ERF |
| 2395. | Glyma.13G122700.1.p | Glyma.13G122700 | ERF |
| 2396. | Glyma.13G122800.1.p | Glyma.13G122800 | ERF |
| 2397. | Glyma.13G122900.1.p | Glyma.13G122900 | ERF |
| 2398. | Glyma.13G123000.1.p | Glyma.13G123000 | ERF |
| 2399. | Glyma.13G123100.1.p | Glyma.13G123100 | ERF |
| 2400. | Glyma.13G124900.1.p | Glyma.13G124900 | GRAS |
| 2401. | Glyma.13G126200.1.p | Glyma.13G126200 | G2-like |
| 2402. | Glyma.13G128500.1.p | Glyma.13G128500 | MYB_related |
| 2403. | Glyma.13G130100.1.p | Glyma.13G130100 | bHLH |
| 2404. | Glyma.13G132800.1.p | Glyma.13G132800 | C2H2 |
| 2405. | Glyma.13G133000.1.p | Glyma.13G133000 | C2H2 |
| 2406. | Glyma.13G133100.1.p | Glyma.13G133100 | C2H2 |
| 2407. | Glyma.13G136300.1.p | Glyma.13G136300 | MYB_related |
| 2408. | Glyma.13G138200.1.p | Glyma.13G138200 | S1Fa-like |
| 2409. | Glyma.13G139000.1.p | Glyma.13G139000 | C2H2 |
| 2410. | Glyma.13G140600.1.p | Glyma.13G140600 | ARF |
| 2411. | Glyma.13G141900.1.p | Glyma.13G141900 | MYB |
| 2412. | Glyma.13G144100.1.p | Glyma.13G144100 | TCP |
| 2413. | Glyma.13G144600.1.p | Glyma.13G144600 | MYB_related |
| 2414. | Glyma.13G145800.1.p | Glyma.13G145800 | MYB |
| 2415. | Glyma.13G146400.1.p | Glyma.13G146400 | C2H2 |
| 2416. | Glyma.13G149500.1.p | Glyma.13G149500 | HD-ZIP |
| 2417. | Glyma.13G149700.1.p | Glyma.13G149700 | Trihelix |
| 2418. | Glyma.13G149900.1.p | Glyma.13G149900 | Trihelix |
| 2419. | Glyma.13G151200.1.p | Glyma.13G151200 | HSF |
| 2420. | Glyma.13G151900.1.p | Glyma.13G151900 | ERF |
| 2421. | Glyma.13G152000.1.p | Glyma.13G152000 | ERF |
| 2422. | Glyma.13G152300.1.p | Glyma.13G152300 | MYB_related |
| 2423. | Glyma.13G153200.1.p | Glyma.13G153200 | bZIP |
| 2424. | Glyma.13G155400.1.p | Glyma.13G155400 | ARR-B |
| 2425. | Glyma.13G157300.1.p | Glyma.13G157300 | TALE |
| 2426. | Glyma.13G157800.1.p | Glyma.13G157800 | YABBY |
| 2427. | Glyma.13G166700.1.p | Glyma.13G166700 | ERF |
| 2428. | Glyma.13G169900.1.p | Glyma.13G169900 | HD-ZIP |
| 2429. | Glyma.13G172700.1.p | Glyma.13G172700 | WOX |
| 2430. | Glyma.13G172900.1.p | Glyma.13G172900 | VOZ |
| 2431. | Glyma.13G174000.1.p | Glyma.13G174000 | ARF |
| 2432. | Glyma.13G174700.1.p | Glyma.13G174700 | NAC |
| 2433. | Glyma.13G175200.1.p | Glyma.13G175200 | HB-other |
| 2434. | Glyma.13G177400.1.p | Glyma.13G177400 | SBP |
| 2435. | Glyma.13G177500.1.p | Glyma.13G177500 | Dof |
| 2436. | Glyma.13G177600.1.p | Glyma.13G177600 | Dof |
| 2437. | Glyma.13G178800.1.p | Glyma.13G178800 | bHLH |
| 2438. | Glyma.13G180200.1.p | Glyma.13G180200 | HSF |
| 2439. | Glyma.13G182700.1.p | Glyma.13G182700 | Dof |
| 2440. | Glyma.13G184500.1.p | Glyma.13G184500 | MYB_related |
| 2441. | Glyma.13G187500.1.p | Glyma.13G187500 | MYB |
| 2442. | Glyma.13G189400.1.p | Glyma.13G189400 | NF-YC |
| 2443. | Glyma.13G191000.1.p | Glyma.13G191000 | LBD |
| 2444. | Glyma.13G193700.1.p | Glyma.13G193700 | bZIP |
| 2445. | Glyma.13G195800.1.p | Glyma.13G195800 | Trihelix |
| 2446. | Glyma.13G197300.1.p | Glyma.13G197300 | SRS |
| 2447. | Glyma.13G197800.1.p | Glyma.13G197800 | MYB_related |
| 2448. | Glyma.13G202300.1.p | Glyma.13G202300 | NF-YA |
| 2449. | Glyma.13G202800.1.p | Glyma.13G202800 | C3H |
| 2450. | Glyma.13G203100.1.p | Glyma.13G203100 | MYB |
| 2451. | Glyma.13G203700.1.p | Glyma.13G203700 | C2H2 |
| 2452. | Glyma.13G204400.1.p | Glyma.13G204400 | bHLH |
| 2453. | Glyma.13G207500.1.p | Glyma.13G207500 | NF-YC |
| 2454. | Glyma.13G207600.1.p | Glyma.13G207600 | NF-YC |
| 2455. | Glyma.13G207700.1.p | Glyma.13G207700 | NF-YC |
| 2456. | Glyma.13G208300.1.p | Glyma.13G208300 | bHLH |
| 2457. | Glyma.13G211200.1.p | Glyma.13G211200 | FAR1 |
| 2458. | Glyma.13G213400.1.p | Glyma.13G213400 | SBP |
| 2459. | Glyma.13G216900.1.p | Glyma.13G216900 | ERF |
| 2460. | Glyma.13G217000.1.p | Glyma.13G217000 | C3H |
| 2461. | Glyma.13G218900.1.p | Glyma.13G218900 | C2H2 |
| 2462. | Glyma.13G219900.1.p | Glyma.13G219900 | TCP |
| 2463. | Glyma.13G221400.1.p | Glyma.13G221400 | ARF |
| 2464. | Glyma.13G223300.1.p | Glyma.13G223300 | MIKC_MADS |
| 2465. | Glyma.13G225700.1.p | Glyma.13G225700 | HSF |
| 2466. | Glyma.13G226800.1.p | Glyma.13G226800 | MYB_related |
| 2467. | Glyma.13G227100.1.p | Glyma.13G227100 | ERF |
| 2468. | Glyma.13G228900.1.p | Glyma.13G228900 | G2-like |
| 2469. | Glyma.13G229600.1.p | Glyma.13G229600 | Trihelix |
| 2470. | Glyma.13G230200.1.p | Glyma.13G230200 | Dof |
| 2471. | Glyma.13G233800.1.p | Glyma.13G233800 | ERF |
| 2472. | Glyma.13G233900.1.p | Glyma.13G233900 | ERF |
| 2473. | Glyma.13G234200.1.p | Glyma.13G234200 | ARF |
| 2474. | Glyma.13G234700.1.p | Glyma.13G234700 | NAC |
| 2475. | Glyma.13G235300.1.p | Glyma.13G235300 | HB-other |
| 2476. | Glyma.13G236500.1.p | Glyma.13G236500 | ERF |
| 2477. | Glyma.13G236600.1.p | Glyma.13G236600 | ERF |
| 2478. | Glyma.13G236800.1.p | Glyma.13G236800 | C3H |
| 2479. | Glyma.13G237400.1.p | Glyma.13G237400 | SBP |
| 2480. | Glyma.13G237500.1.p | Glyma.13G237500 | Dof |
| 2481. | Glyma.13G237600.1.p | Glyma.13G237600 | Dof |
| 2482. | Glyma.13G238300.1.p | Glyma.13G238300 | bHLH |
| 2483. | Glyma.13G241900.1.p | Glyma.13G241900 | Dof |
| 2484. | Glyma.13G243200.1.p | Glyma.13G243200 | NAC |
| 2485. | Glyma.13G244400.1.p | Glyma.13G244400 | MYB_related |
| 2486. | Glyma.13G245900.1.p | Glyma.13G245900 | B3 |
| 2487. | Glyma.13G247200.1.p | Glyma.13G247200 | MYB |
| 2488. | Glyma.13G249800.1.p | Glyma.13G249800 | bHLH |
| 2489. | Glyma.13G250300.1.p | Glyma.13G250300 | bHLH |
| 2490. | Glyma.13G251000.1.p | Glyma.13G251000 | GeBP |
| 2491. | Glyma.13G251300.1.p | Glyma.13G251300 | bHLH |
| 2492. | Glyma.13G253500.1.p | Glyma.13G253500 | bHLH |
| 2493. | Glyma.13G255200.1.p | Glyma.13G255200 | MIKC_MADS |
| 2494. | Glyma.13G256900.1.p | Glyma.13G256900 | MIKC_MADS |
| 2495. | Glyma.13G257000.1.p | Glyma.13G257000 | MIKC_MADS |
| 2496. | Glyma.13G260300.1.p | Glyma.13G260300 | bZIP |
| 2497. | Glyma.13G265000.1.p | Glyma.13G265000 | DBB |
| 2498. | Glyma.13G266500.1.p | Glyma.13G266500 | BES1 |
| 2499. | Glyma.13G266600.1.p | Glyma.13G266600 | BES1 |
| 2500. | Glyma.13G267400.1.p | Glyma.13G267400 | WRKY |
| 2501. | Glyma.13G267500.1.p | Glyma.13G267500 | WRKY |
| 2502. | Glyma.13G267600.1.p | Glyma.13G267600 | WRKY |
| 2503. | Glyma.13G267700.1.p | Glyma.13G267700 | WRKY |
| 2504. | Glyma.13G269400.1.p | Glyma.13G269400 | C2H2 |
| 2505. | Glyma.13G271700.1.p | Glyma.13G271700 | TCP |
| 2506. | Glyma.13G272500.1.p | Glyma.13G272500 | bZIP |
| 2507. | Glyma.13G274100.1.p | Glyma.13G274100 | ERF |
| 2508. | Glyma.13G274300.1.p | Glyma.13G274300 | NAC |
| 2509. | Glyma.13G274900.1.p | Glyma.13G274900 | SBP |
| 2510. | Glyma.13G279900.1.p | Glyma.13G279900 | NAC |
| 2511. | Glyma.13G280000.1.p | Glyma.13G280000 | NAC |
| 2512. | Glyma.13G282100.1.p | Glyma.13G282100 | MYB |
| 2513. | Glyma.13G284000.1.p | Glyma.13G284000 | NF-YC |
| 2514. | Glyma.13G284400.1.p | Glyma.13G284400 | bHLH |
| 2515. | Glyma.13G285400.1.p | Glyma.13G285400 | GRAS |
| 2516. | Glyma.13G286700.1.p | Glyma.13G286700 | bHLH |
| 2517. | Glyma.13G289400.1.p | Glyma.13G289400 | WRKY |
| 2518. | Glyma.13G290100.1.p | Glyma.13G290100 | G2-like |
| 2519. | Glyma.13G290500.1.p | Glyma.13G290500 | Trihelix |
| 2520. | Glyma.13G291400.1.p | Glyma.13G291400 | bHLH |
| 2521. | Glyma.13G292500.1.p | Glyma.13G292500 | TCP |
| 2522. | Glyma.13G292800.1.p | Glyma.13G292800 | bZIP |
| 2523. | Glyma.13G293800.1.p | Glyma.13G293800 | C2H2 |
| 2524. | Glyma.13G294000.1.p | Glyma.13G294000 | NAC |
| 2525. | Glyma.13G294300.1.p | Glyma.13G294300 | G2-like |
| 2526. | Glyma.13G295400.1.p | Glyma.13G295400 | SBP |
| 2527. | Glyma.13G296100.1.p | Glyma.13G296100 | bHLH |
| 2528. | Glyma.13G298600.1.p | Glyma.13G298600 | ERF |
| 2529. | Glyma.13G302400.1.p | Glyma.13G302400 | MYB |
| 2530. | Glyma.13G303100.1.p | Glyma.13G303100 | MYB_related |
| 2531. | Glyma.13G303200.1.p | Glyma.13G303200 | MYB |
| 2532. | Glyma.13G304300.1.p | Glyma.13G304300 | ERF |
| 2533. | Glyma.13G304900.1.p | Glyma.13G304900 | GRAS |
| 2534. | Glyma.13G305500.1.p | Glyma.13G305500 | bHLH |
| 2535. | Glyma.13G307300.1.p | Glyma.13G307300 | MYB_related |
| 2536. | Glyma.13G308200.1.p | Glyma.13G308200 | HD-ZIP |
| 2537. | Glyma.13G309200.1.p | Glyma.13G309200 | MYB |
| 2538. | Glyma.13G310100.1.p | Glyma.13G310100 | WRKY |
| 2539. | Glyma.13G311200.1.p | Glyma.13G311200 | YABBY |
| 2540. | Glyma.13G312900.1.p | Glyma.13G312900 | TALE |
| 2541. | Glyma.13G313800.1.p | Glyma.13G313800 | M-type_MADS |
| 2542. | Glyma.13G314600.1.p | Glyma.13G314600 | NAC |
| 2543. | Glyma.13G315300.1.p | Glyma.13G315300 | NAC |
| 2544. | Glyma.13G316600.1.p | Glyma.13G316600 | G2-like |
| 2545. | Glyma.13G316900.1.p | Glyma.13G316900 | bZIP |
| 2546. | Glyma.13G317000.1.p | Glyma.13G317000 | bZIP |
| 2547. | Glyma.13G317200.1.p | Glyma.13G317200 | C2H2 |
| 2548. | Glyma.13G318500.1.p | Glyma.13G318500 | ERF |
| 2549. | Glyma.13G321800.1.p | Glyma.13G321800 | C2H2 |
| 2550. | Glyma.13G321900.1.p | Glyma.13G321900 | C2H2 |
| 2551. | Glyma.13G322100.1.p | Glyma.13G322100 | bHLH |
| 2552. | Glyma.13G322900.1.p | Glyma.13G322900 | MYB |
| 2553. | Glyma.13G324200.1.p | Glyma.13G324200 | TALE |
| 2554. | Glyma.13G325100.1.p | Glyma.13G325100 | GATA |
| 2555. | Glyma.13G325200.1.p | Glyma.13G325200 | ARF |
| 2556. | Glyma.13G327500.1.p | Glyma.13G327500 | C2H2 |
| 2557. | Glyma.13G327600.1.p | Glyma.13G327600 | NAC |
| 2558. | Glyma.13G328000.1.p | Glyma.13G328000 | ARF |
| 2559. | Glyma.13G328500.1.p | Glyma.13G328500 | LBD |
| 2560. | Glyma.13G329000.1.p | Glyma.13G329000 | Dof |
| 2561. | Glyma.13G329700.1.p | Glyma.13G329700 | AP2 |
| 2562. | Glyma.13G333200.1.p | Glyma.13G333200 | MYB |
| 2563. | Glyma.13G333400.1.p | Glyma.13G333400 | C2H2 |
| 2564. | Glyma.13G334900.1.p | Glyma.13G334900 | WOX |
| 2565. | Glyma.13G335200.1.p | Glyma.13G335200 | Dof |
| 2566. | Glyma.13G337100.1.p | Glyma.13G337100 | MYB_related |
| 2567. | Glyma.13G337200.1.p | Glyma.13G337200 | GRAS |
| 2568. | Glyma.13G337300.1.p | Glyma.13G337300 | GRAS |
| 2569. | Glyma.13G337400.1.p | Glyma.13G337400 | GRAS |
| 2570. | Glyma.13G337500.1.p | Glyma.13G337500 | GRAS |
| 2571. | Glyma.13G337600.1.p | Glyma.13G337600 | GRAS |
| 2572. | Glyma.13G339900.1.p | Glyma.13G339900 | MYB |
| 2573. | Glyma.13G340600.1.p | Glyma.13G340600 | Trihelix |
| 2574. | Glyma.13G341600.1.p | Glyma.13G341600 | FAR1 |
| 2575. | Glyma.13G341700.1.p | Glyma.13G341700 | bHLH |
| 2576. | Glyma.13G342500.1.p | Glyma.13G342500 | EIL |
| 2577. | Glyma.13G344100.1.p | Glyma.13G344100 | FAR1 |
| 2578. | Glyma.13G344700.1.p | Glyma.13G344700 | DBB |
| 2579. | Glyma.13G345200.1.p | Glyma.13G345200 | bZIP |
| 2580. | Glyma.13G345700.1.p | Glyma.13G345700 | GRAS |
| 2581. | Glyma.13G346300.1.p | Glyma.13G346300 | Nin-like |
| 2582. | Glyma.13G349500.1.p | Glyma.13G349500 | C2H2 |
| 2583. | Glyma.13G349700.1.p | Glyma.13G349700 | LBD |
| 2584. | Glyma.13G352000.1.p | Glyma.13G352000 | Dof |
| 2585. | Glyma.13G352600.1.p | Glyma.13G352600 | bHLH |
| 2586. | Glyma.13G354800.1.p | Glyma.13G354800 | MYB_related |
| 2587. | Glyma.13G355700.1.p | Glyma.13G355700 | ERF |
| 2588. | Glyma.13G357100.1.p | Glyma.13G357100 | HD-ZIP |
| 2589. | Glyma.13G359800.1.p | Glyma.13G359800 | Trihelix |
| 2590. | Glyma.13G363600.1.p | Glyma.13G363600 | C3H |
| 2591. | Glyma.13G368500.1.p | Glyma.13G368500 | bHLH |
| 2592. | Glyma.13G368600.1.p | Glyma.13G368600 | bHLH |
| 2593. | Glyma.13G368700.1.p | Glyma.13G368700 | bHLH |
| 2594. | Glyma.13G369400.1.p | Glyma.13G369400 | ERF |
| 2595. | Glyma.13G369500.1.p | Glyma.13G369500 | C2H2 |
| 2596. | Glyma.13G370100.1.p | Glyma.13G370100 | WRKY |

| 2597. | Glyma.14G000400.1.p | Glyma.14G000400 | FAR1 |
| --- | --- | --- | --- |
| 2598. | Glyma.14G001600.1.p | Glyma.14G001600 | Nin-like |
| 2599. | Glyma.14G001700.1.p | Glyma.14G001700 | C2H2 |
| 2600. | Glyma.14G006800.1.p | Glyma.14G006800 | WRKY |
| 2601. | Glyma.14G006900.1.p | Glyma.14G006900 | GRAS |
| 2602. | Glyma.14G009200.1.p | Glyma.14G009200 | SAP |
| 2603. | Glyma.14G010000.1.p | Glyma.14G010000 | NF-YA |
| 2604. | Glyma.14G016000.1.p | Glyma.14G016000 | GRAS |
| 2605. | Glyma.14G016200.1.p | Glyma.14G016200 | WRKY |
| 2606. | Glyma.14G016300.1.p | Glyma.14G016300 | C3H |
| 2607. | Glyma.14G020100.1.p | Glyma.14G020100 | ERF |
| 2608. | Glyma.14G021100.1.p | Glyma.14G021100 | LBD |
| 2609. | Glyma.14G027200.1.p | Glyma.14G027200 | M-type_MADS |
| 2610. | Glyma.14G028900.1.p | Glyma.14G028900 | WRKY |
| 2611. | Glyma.14G030700.1.p | Glyma.14G030700 | NAC |
| 2612. | Glyma.14G031500.1.p | Glyma.14G031500 | G2-like |
| 2613. | Glyma.14G032200.1.p | Glyma.14G032200 | bHLH |
| 2614. | Glyma.14G032700.1.p | Glyma.14G032700 | ARF |
| 2615. | Glyma.14G034300.1.p | Glyma.14G034300 | HB-other |
| 2616. | Glyma.14G034800.1.p | Glyma.14G034800 | SRS |
| 2617. | Glyma.14G036200.1.p | Glyma.14G036200 | HSF |
| 2618. | Glyma.14G038800.1.p | Glyma.14G038800 | NF-YC |
| 2619. | Glyma.14G039200.1.p | Glyma.14G039200 | MYB |
| 2620. | Glyma.14G041500.1.p | Glyma.14G041500 | EIL |
| 2621. | Glyma.14G043900.1.p | Glyma.14G043900 | FAR1 |
| 2622. | Glyma.14G047000.1.p | Glyma.14G047000 | TALE |
| 2623. | Glyma.14G050000.1.p | Glyma.14G050000 | C3H |
| 2624. | Glyma.14G050100.1.p | Glyma.14G050100 | ERF |
| 2625. | Glyma.14G055700.1.p | Glyma.14G055700 | B3 |
| 2626. | Glyma.14G056200.1.p | Glyma.14G056200 | ERF |
| 2627. | Glyma.14G056700.1.p | Glyma.14G056700 | LBD |
| 2628. | Glyma.14G057600.1.p | Glyma.14G057600 | LBD |
| 2629. | Glyma.14G057900.1.p | Glyma.14G057900 | ERF |
| 2630. | Glyma.14G058100.1.p | Glyma.14G058100 | MYB |
| 2631. | Glyma.14G058200.1.p | Glyma.14G058200 | bHLH |
| 2632. | Glyma.14G062200.1.p | Glyma.14G062200 | MYB |
| 2633. | Glyma.14G063400.1.p | Glyma.14G063400 | MYB |
| 2634. | Glyma.14G066500.1.p | Glyma.14G066500 | C2H2 |
| 2635. | Glyma.14G066900.1.p | Glyma.14G066900 | LBD |
| 2636. | Glyma.14G067100.1.p | Glyma.14G067100 | MYB_related |
| 2637. | Glyma.14G069100.1.p | Glyma.14G069100 | MYB_related |
| 2638. | Glyma.14G069700.1.p | Glyma.14G069700 | bHLH |
| 2639. | Glyma.14G070000.1.p | Glyma.14G070000 | ERF |
| 2640. | Glyma.14G070700.1.p | Glyma.14G070700 | TALE |
| 2641. | Glyma.14G071400.1.p | Glyma.14G071400 | bZIP |
| 2642. | Glyma.14G074500.1.p | Glyma.14G074500 | MYB_related |
| 2643. | Glyma.14G076900.1.p | Glyma.14G076900 | BES1 |
| 2644. | Glyma.14G079000.1.p | Glyma.14G079000 | G2-like |
| 2645. | Glyma.14G079100.1.p | Glyma.14G079100 | B3 |
| 2646. | Glyma.14G079200.1.p | Glyma.14G079200 | B3 |
| 2647. | Glyma.14G083700.1.p | Glyma.14G083700 | HSF |
| 2648. | Glyma.14G084200.1.p | Glyma.14G084200 | bHLH |
| 2649. | Glyma.14G084300.1.p | Glyma.14G084300 | NAC |
| 2650. | Glyma.14G084600.1.p | Glyma.14G084600 | WOX |
| 2651. | Glyma.14G084700.1.p | Glyma.14G084700 | ERF |
| 2652. | Glyma.14G085500.1.p | Glyma.14G085500 | WRKY |
| 2653. | Glyma.14G086500.1.p | Glyma.14G086500 | MYB |
| 2654. | Glyma.14G088200.1.p | Glyma.14G088200 | LBD |
| 2655. | Glyma.14G088300.1.p | Glyma.14G088300 | C2H2 |
| 2656. | Glyma.14G088400.1.p | Glyma.14G088400 | bHLH |
| 2657. | Glyma.14G089200.1.p | Glyma.14G089200 | AP2 |
| 2658. | Glyma.14G089600.1.p | Glyma.14G089600 | bHLH |
| 2659. | Glyma.14G090900.1.p | Glyma.14G090900 | MYB |
| 2660. | Glyma.14G091000.1.p | Glyma.14G091000 | HD-ZIP |
| 2661. | Glyma.14G091200.1.p | Glyma.14G091200 | TALE |
| 2662. | Glyma.14G091500.1.p | Glyma.14G091500 | MYB |
| 2663. | Glyma.14G093800.1.p | Glyma.14G093800 | MYB_related |
| 2664. | Glyma.14G094800.1.p | Glyma.14G094800 | GATA |
| 2665. | Glyma.14G095900.1.p | Glyma.14G095900 | C2H2 |
| 2666. | Glyma.14G096800.1.p | Glyma.14G096800 | HSF |
| 2667. | Glyma.14G100100.1.p | Glyma.14G100100 | WRKY |
| 2668. | Glyma.14G102200.1.p | Glyma.14G102200 | bHLH |
| 2669. | Glyma.14G102900.1.p | Glyma.14G102900 | WRKY |
| 2670. | Glyma.14G103100.1.p | Glyma.14G103100 | WRKY |
| 2671. | Glyma.14G106200.1.p | Glyma.14G106200 | ERF |
| 2672. | Glyma.14G110600.1.p | Glyma.14G110600 | ARR-B |
| 2673. | Glyma.14G110900.1.p | Glyma.14G110900 | C2H2 |
| 2674. | Glyma.14G111600.1.p | Glyma.14G111600 | ERF |
| 2675. | Glyma.14G112400.1.p | Glyma.14G112400 | TALE |
| 2676. | Glyma.14G119000.1.p | Glyma.14G119000 | MYB |
| 2677. | Glyma.14G119100.1.p | Glyma.14G119100 | GRAS |
| 2678. | Glyma.14G120600.1.p | Glyma.14G120600 | MYB_related |
| 2679. | Glyma.14G123900.1.p | Glyma.14G123900 | ERF |
| 2680. | Glyma.14G127400.1.p | Glyma.14G127400 | BES1 |
| 2681. | Glyma.14G135400.1.p | Glyma.14G135400 | WRKY |
| 2682. | Glyma.14G136500.1.p | Glyma.14G136500 | bHLH |
| 2683. | Glyma.14G139500.1.p | Glyma.14G139500 | G2-like |
| 2684. | Glyma.14G140100.1.p | Glyma.14G140100 | NAC |
| 2685. | Glyma.14G143400.1.p | Glyma.14G143400 | MYB |
| 2686. | Glyma.14G145700.1.p | Glyma.14G145700 | GATA |
| 2687. | Glyma.14G146100.1.p | Glyma.14G146100 | ERF |
| 2688. | Glyma.14G147500.1.p | Glyma.14G147500 | ERF |
| 2689. | Glyma.14G152700.1.p | Glyma.14G152700 | NAC |
| 2690. | Glyma.14G154400.1.p | Glyma.14G154400 | MYB |
| 2691. | Glyma.14G155100.1.p | Glyma.14G155100 | MIKC_MADS |
| 2692. | Glyma.14G160700.1.p | Glyma.14G160700 | bHLH |
| 2693. | Glyma.14G161900.1.p | Glyma.14G161900 | ERF |
| 2694. | Glyma.14G166500.1.p | Glyma.14G166500 | ARF |
| 2695. | Glyma.14G167000.1.p | Glyma.14G167000 | bZIP |
| 2696. | Glyma.14G168700.1.p | Glyma.14G168700 | MIKC_MADS |
| 2697. | Glyma.14G171500.1.p | Glyma.14G171500 | ERF |
| 2698. | Glyma.14G174900.1.p | Glyma.14G174900 | C2H2 |
| 2699. | Glyma.14G178000.1.p | Glyma.14G178000 | bHLH |
| 2700. | Glyma.14G179600.1.p | Glyma.14G179600 | ZF-HD |
| 2701. | Glyma.14G181500.1.p | Glyma.14G181500 | bHLH |
| 2702. | Glyma.14G182800.1.p | Glyma.14G182800 | GATA |
| 2703. | Glyma.14G183800.1.p | Glyma.14G183800 | MIKC_MADS |
| 2704. | Glyma.14G185100.1.p | Glyma.14G185100 | bHLH |
| 2705. | Glyma.14G185800.1.p | Glyma.14G185800 | WRKY |
| 2706. | Glyma.14G186000.1.p | Glyma.14G186000 | WRKY |
| 2707. | Glyma.14G186100.1.p | Glyma.14G186100 | WRKY |
| 2708. | Glyma.14G189300.1.p | Glyma.14G189300 | NAC |
| 2709. | Glyma.14G190400.1.p | Glyma.14G190400 | CO-like |
| 2710. | Glyma.14G191700.1.p | Glyma.14G191700 | MYB |
| 2711. | Glyma.14G192600.1.p | Glyma.14G192600 | MYB |
| 2712. | Glyma.14G194100.1.p | Glyma.14G194100 | C3H |
| 2713. | Glyma.14G197200.1.p | Glyma.14G197200 | bZIP |
| 2714. | Glyma.14G199800.1.p | Glyma.14G199800 | WRKY |
| 2715. | Glyma.14G200200.1.p | Glyma.14G200200 | WRKY |
| 2716. | Glyma.14G202300.1.p | Glyma.14G202300 | LBD |
| 2717. | Glyma.14G202600.1.p | Glyma.14G202600 | MYB_related |
| 2718. | Glyma.14G204100.1.p | Glyma.14G204100 | bZIP |
| 2719. | Glyma.14G204400.1.p | Glyma.14G204400 | B3 |
| 2720. | Glyma.14G204400.2.p | Glyma.14G204400 | B3 |
| 2721. | Glyma.14G205600.1.p | Glyma.14G205600 | ERF |
| 2722. | Glyma.14G208500.1.p | Glyma.14G208500 | ARF |
| 2723. | Glyma.14G210000.1.p | Glyma.14G210000 | NAC |
| 2724. | Glyma.14G210600.1.p | Glyma.14G210600 | MYB_related |
| 2725. | Glyma.14G211900.1.p | Glyma.14G211900 | G2-like |
| 2726. | Glyma.14G212000.1.p | Glyma.14G212000 | C2H2 |
| 2727. | Glyma.14G214500.1.p | Glyma.14G214500 | MYB |
| 2728. | Glyma.14G216400.1.p | Glyma.14G216400 | SRS |
| 2729. | Glyma.14G216500.1.p | Glyma.14G216500 | DBB |
| 2730. | Glyma.14G217200.1.p | Glyma.14G217200 | bZIP |
| 2731. | Glyma.14G217700.1.p | Glyma.14G217700 | ARF |
| 2732. | Glyma.14G218700.1.p | Glyma.14G218700 | C2H2 |

| 2733. | Glyma.15G002100.1.p | Glyma.15G002100 | FAR1 |
| --- | --- | --- | --- |
| 2734. | Glyma.15G003300.1.p | Glyma.15G003300 | WRKY |
| 2735. | Glyma.15G004100.1.p | Glyma.15G004100 | C2H2 |
| 2736. | Glyma.15G004200.1.p | Glyma.15G004200 | ERF |
| 2737. | Glyma.15G005000.1.p | Glyma.15G005000 | bHLH |
| 2738. | Glyma.15G005100.1.p | Glyma.15G005100 | bHLH |
| 2739. | Glyma.15G008600.1.p | Glyma.15G008600 | ERF |
| 2740. | Glyma.15G011100.1.p | Glyma.15G011100 | bHLH |
| 2741. | Glyma.15G014200.1.p | Glyma.15G014200 | Trihelix |
| 2742. | Glyma.15G016500.1.p | Glyma.15G016500 | HD-ZIP |
| 2743. | Glyma.15G017300.1.p | Glyma.15G017300 | bHLH |
| 2744. | Glyma.15G018400.1.p | Glyma.15G018400 | ERF |
| 2745. | Glyma.15G019400.1.p | Glyma.15G019400 | MYB_related |
| 2746. | Glyma.15G022000.1.p | Glyma.15G022000 | bHLH |
| 2747. | Glyma.15G022800.1.p | Glyma.15G022800 | Dof |
| 2748. | Glyma.15G024500.1.p | Glyma.15G024500 | C2H2 |
| 2749. | Glyma.15G025100.1.p | Glyma.15G025100 | ERF |
| 2750. | Glyma.15G025500.1.p | Glyma.15G025500 | MYB |
| 2751. | Glyma.15G027400.1.p | Glyma.15G027400 | NF-YA |
| 2752. | Glyma.15G027900.1.p | Glyma.15G027900 | Nin-like |
| 2753. | Glyma.15G028600.1.p | Glyma.15G028600 | GRAS |
| 2754. | Glyma.15G029100.1.p | Glyma.15G029100 | bZIP |
| 2755. | Glyma.15G029500.1.p | Glyma.15G029500 | DBB |
| 2756. | Glyma.15G029900.1.p | Glyma.15G029900 | FAR1 |
| 2757. | Glyma.15G030200.1.p | Glyma.15G030200 | FAR1 |
| 2758. | Glyma.15G031800.1.p | Glyma.15G031800 | EIL |
| 2759. | Glyma.15G032700.1.p | Glyma.15G032700 | bHLH |
| 2760. | Glyma.15G032800.1.p | Glyma.15G032800 | FAR1 |
| 2761. | Glyma.15G033600.1.p | Glyma.15G033600 | C2H2 |
| 2762. | Glyma.15G033800.1.p | Glyma.15G033800 | Trihelix |
| 2763. | Glyma.15G034500.1.p | Glyma.15G034500 | MYB |
| 2764. | Glyma.15G036900.1.p | Glyma.15G036900 | GRAS |
| 2765. | Glyma.15G037000.1.p | Glyma.15G037000 | GRAS |
| 2766. | Glyma.15G037100.1.p | Glyma.15G037100 | GRAS |
| 2767. | Glyma.15G037200.1.p | Glyma.15G037200 | GRAS |
| 2768. | Glyma.15G037300.1.p | Glyma.15G037300 | MYB_related |
| 2769. | Glyma.15G039300.1.p | Glyma.15G039300 | Dof |
| 2770. | Glyma.15G039600.1.p | Glyma.15G039600 | WOX |
| 2771. | Glyma.15G040700.1.p | Glyma.15G040700 | C2H2 |
| 2772. | Glyma.15G041100.1.p | Glyma.15G041100 | MYB |
| 2773. | Glyma.15G044400.1.p | Glyma.15G044400 | AP2 |
| 2774. | Glyma.15G044800.1.p | Glyma.15G044800 | Dof |
| 2775. | Glyma.15G045200.1.p | Glyma.15G045200 | LBD |
| 2776. | Glyma.15G045700.1.p | Glyma.15G045700 | GATA |
| 2777. | Glyma.15G049000.1.p | Glyma.15G049000 | bZIP |
| 2778. | Glyma.15G051200.1.p | Glyma.15G051200 | NAC |
| 2779. | Glyma.15G053600.1.p | Glyma.15G053600 | CAMTA |
| 2780. | Glyma.15G054200.1.p | Glyma.15G054200 | C2H2 |
| 2781. | Glyma.15G057900.1.p | Glyma.15G057900 | MIKC_MADS |
| 2782. | Glyma.15G058000.1.p | Glyma.15G058000 | MIKC_MADS |
| 2783. | Glyma.15G059600.1.p | Glyma.15G059600 | MIKC_MADS |
| 2784. | Glyma.15G061400.1.p | Glyma.15G061400 | bHLH |
| 2785. | Glyma.15G062600.1.p | Glyma.15G062600 | bHLH |
| 2786. | Glyma.15G063000.1.p | Glyma.15G063000 | bHLH |
| 2787. | Glyma.15G063300.1.p | Glyma.15G063300 | GeBP |
| 2788. | Glyma.15G063900.1.p | Glyma.15G063900 | bHLH |
| 2789. | Glyma.15G064000.1.p | Glyma.15G064000 | bHLH |
| 2790. | Glyma.15G064500.1.p | Glyma.15G064500 | bHLH |
| 2791. | Glyma.15G066800.1.p | Glyma.15G066800 | MYB |
| 2792. | Glyma.15G067900.1.p | Glyma.15G067900 | B3 |
| 2793. | Glyma.15G069300.1.p | Glyma.15G069300 | MYB_related |
| 2794. | Glyma.15G070300.1.p | Glyma.15G070300 | NAC |
| 2795. | Glyma.15G071400.1.p | Glyma.15G071400 | Dof |
| 2796. | Glyma.15G075000.1.p | Glyma.15G075000 | bHLH |
| 2797. | Glyma.15G075800.1.p | Glyma.15G075800 | Dof |
| 2798. | Glyma.15G076000.1.p | Glyma.15G076000 | Dof |
| 2799. | Glyma.15G076200.1.p | Glyma.15G076200 | SBP |
| 2800. | Glyma.15G076800.1.p | Glyma.15G076800 | C3H |
| 2801. | Glyma.15G077000.1.p | Glyma.15G077000 | ERF |
| 2802. | Glyma.15G077100.1.p | Glyma.15G077100 | ERF |
| 2803. | Glyma.15G078300.1.p | Glyma.15G078300 | NAC |
| 2804. | Glyma.15G078800.1.p | Glyma.15G078800 | ARF |
| 2805. | Glyma.15G079100.1.p | Glyma.15G079100 | ERF |
| 2806. | Glyma.15G079200.1.p | Glyma.15G079200 | ERF |
| 2807. | Glyma.15G082400.1.p | Glyma.15G082400 | Dof |
| 2808. | Glyma.15G082900.1.p | Glyma.15G082900 | Trihelix |
| 2809. | Glyma.15G083600.1.p | Glyma.15G083600 | G2-like |
| 2810. | Glyma.15G085400.1.p | Glyma.15G085400 | ERF |
| 2811. | Glyma.15G086400.1.p | Glyma.15G086400 | HSF |
| 2812. | Glyma.15G088600.1.p | Glyma.15G088600 | MIKC_MADS |
| 2813. | Glyma.15G091000.1.p | Glyma.15G091000 | ARF |
| 2814. | Glyma.15G092500.1.p | Glyma.15G092500 | TCP |
| 2815. | Glyma.15G095500.1.p | Glyma.15G095500 | C3H |
| 2816. | Glyma.15G095600.1.p | Glyma.15G095600 | ERF |
| 2817. | Glyma.15G099400.1.p | Glyma.15G099400 | SBP |
| 2818. | Glyma.15G101500.1.p | Glyma.15G101500 | FAR1 |
| 2819. | Glyma.15G104600.1.p | Glyma.15G104600 | bHLH |
| 2820. | Glyma.15G109100.1.p | Glyma.15G109100 | G2-like |
| 2821. | Glyma.15G110300.1.p | Glyma.15G110300 | WRKY |
| 2822. | Glyma.15G111900.1.p | Glyma.15G111900 | TALE |
| 2823. | Glyma.15G116300.1.p | Glyma.15G116300 | GRAS |
| 2824. | Glyma.15G118800.1.p | Glyma.15G118800 | NF-YB |
| 2825. | Glyma.15G119000.1.p | Glyma.15G119000 | Trihelix |
| 2826. | Glyma.15G123000.1.p | Glyma.15G123000 | G2-like |
| 2827. | Glyma.15G123100.1.p | Glyma.15G123100 | G2-like |
| 2828. | Glyma.15G125400.1.p | Glyma.15G125400 | FAR1 |
| 2829. | Glyma.15G125500.1.p | Glyma.15G125500 | FAR1 |
| 2830. | Glyma.15G127900.1.p | Glyma.15G127900 | LBD |
| 2831. | Glyma.15G129700.1.p | Glyma.15G129700 | HD-ZIP |
| 2832. | Glyma.15G129900.1.p | Glyma.15G129900 | NF-YA |
| 2833. | Glyma.15G132000.1.p | Glyma.15G132000 | HD-ZIP |
| 2834. | Glyma.15G134100.1.p | Glyma.15G134100 | MYB |
| 2835. | Glyma.15G135100.1.p | Glyma.15G135100 | MYB_related |
| 2836. | Glyma.15G135600.1.p | Glyma.15G135600 | WRKY |
| 2837. | Glyma.15G139000.1.p | Glyma.15G139000 | WRKY |
| 2838. | Glyma.15G141300.1.p | Glyma.15G141300 | GRAS |
| 2839. | Glyma.15G141400.1.p | Glyma.15G141400 | GRAS |
| 2840. | Glyma.15G143400.1.p | Glyma.15G143400 | CAMTA |
| 2841. | Glyma.15G144000.1.p | Glyma.15G144000 | MYB |
| 2842. | Glyma.15G144500.1.p | Glyma.15G144500 | FAR1 |
| 2843. | Glyma.15G145200.1.p | Glyma.15G145200 | ARR-B |
| 2844. | Glyma.15G152000.1.p | Glyma.15G152000 | ERF |
| 2845. | Glyma.15G153900.1.p | Glyma.15G153900 | NF-YB |
| 2846. | Glyma.15G159100.1.p | Glyma.15G159100 | ERF |
| 2847. | Glyma.15G159200.1.p | Glyma.15G159200 | ERF |
| 2848. | Glyma.15G166200.1.p | Glyma.15G166200 | HB-PHD |
| 2849. | Glyma.15G166800.1.p | Glyma.15G166800 | bHLH |
| 2850. | Glyma.15G168200.1.p | Glyma.15G168200 | WRKY |
| 2851. | Glyma.15G168800.1.p | Glyma.15G168800 | HD-ZIP |
| 2852. | Glyma.15G169300.1.p | Glyma.15G169300 | GATA |
| 2853. | Glyma.15G170500.1.p | Glyma.15G170500 | bHLH |
| 2854. | Glyma.15G173300.1.p | Glyma.15G173300 | NF-YA |
| 2855. | Glyma.15G176000.1.p | Glyma.15G176000 | MYB |
| 2856. | Glyma.15G176500.1.p | Glyma.15G176500 | GRF |
| 2857. | Glyma.15G180000.1.p | Glyma.15G180000 | ERF |
| 2858. | Glyma.15G181000.1.p | Glyma.15G181000 | ARF |
| 2859. | Glyma.15G184200.1.p | Glyma.15G184200 | FAR1 |
| 2860. | Glyma.15G186300.1.p | Glyma.15G186300 | WRKY |
| 2861. | Glyma.15G190200.1.p | Glyma.15G190200 | SBP |
| 2862. | Glyma.15G192000.1.p | Glyma.15G192000 | LSD |
| 2863. | Glyma.15G202200.1.p | Glyma.15G202200 | TALE |
| 2864. | Glyma.15G203900.1.p | Glyma.15G203900 | bHLH |
| 2865. | Glyma.15G206200.1.p | Glyma.15G206200 | ARR-B |
| 2866. | Glyma.15G211900.1.p | Glyma.15G211900 | STAT |
| 2867. | Glyma.15G213300.1.p | Glyma.15G213300 | GRAS |
| 2868. | Glyma.15G215000.1.p | Glyma.15G215000 | G2-like |
| 2869. | Glyma.15G215100.1.p | Glyma.15G215100 | bHLH |
| 2870. | Glyma.15G215500.1.p | Glyma.15G215500 | Dof |
| 2871. | Glyma.15G217400.1.p | Glyma.15G217400 | C3H |
| 2872. | Glyma.15G221600.1.p | Glyma.15G221600 | AP2 |
| 2873. | Glyma.15G222100.1.p | Glyma.15G222100 | FAR1 |
| 2874. | Glyma.15G222400.1.p | Glyma.15G222400 | bZIP |
| 2875. | Glyma.15G225300.1.p | Glyma.15G225300 | MYB |
| 2876. | Glyma.15G227300.1.p | Glyma.15G227300 | NF-YC |
| 2877. | Glyma.15G228100.1.p | Glyma.15G228100 | LBD |
| 2878. | Glyma.15G232000.1.p | Glyma.15G232000 | bZIP |
| 2879. | Glyma.15G234100.1.p | Glyma.15G234100 | Trihelix |
| 2880. | Glyma.15G235000.1.p | Glyma.15G235000 | SRS |
| 2881. | Glyma.15G236400.1.p | Glyma.15G236400 | MYB_related |
| 2882. | Glyma.15G254000.1.p | Glyma.15G254000 | NAC |
| 2883. | Glyma.15G257700.1.p | Glyma.15G257700 | NAC |
| 2884. | Glyma.15G258100.1.p | Glyma.15G258100 | LBD |
| 2885. | Glyma.15G259400.1.p | Glyma.15G259400 | MYB |
| 2886. | Glyma.15G260500.1.p | Glyma.15G260500 | bHLH |
| 2887. | Glyma.15G261300.1.p | Glyma.15G261300 | NF-YC |
| 2888. | Glyma.15G263700.1.p | Glyma.15G263700 | G2-like |
| 2889. | Glyma.15G264000.1.p | Glyma.15G264000 | MYB |
| 2890. | Glyma.15G264100.1.p | Glyma.15G264100 | NAC |
| 2891. | Glyma.15G266500.1.p | Glyma.15G266500 | NAC |
| 2892. | Glyma.15G269100.1.p | Glyma.15G269100 | HD-ZIP |
| 2893. | Glyma.15G269600.1.p | Glyma.15G269600 | MYB_related |
| 2894. | Glyma.15G271900.1.p | Glyma.15G271900 | bHLH |
| 2895. | Glyma.15G272400.1.p | Glyma.15G272400 | C3H |
| 2896. | Glyma.15G273000.1.p | Glyma.15G273000 | C3H |
| 2897. | Glyma.15G273400.1.p | Glyma.15G273400 | C2H2 |
| 2898. | Glyma.15G274800.1.p | Glyma.15G274800 | C3H |

| 2899. | Glyma.16G000300.1.p | Glyma.16G000300 | ARF |
| --- | --- | --- | --- |
| 2900. | Glyma.16G004300.1.p | Glyma.16G004300 | TCP |
| 2901. | Glyma.16G005500.1.p | Glyma.16G005500 | NF-YA |
| 2902. | Glyma.16G006500.1.p | Glyma.16G006500 | MYB_related |
| 2903. | Glyma.16G007100.1.p | Glyma.16G007100 | MYB |
| 2904. | Glyma.16G007200.1.p | Glyma.16G007200 | MYB |
| 2905. | Glyma.16G007400.1.p | Glyma.16G007400 | AP2 |
| 2906. | Glyma.16G007600.1.p | Glyma.16G007600 | GRF |
| 2907. | Glyma.16G008200.1.p | Glyma.16G008200 | GRAS |
| 2908. | Glyma.16G010500.1.p | Glyma.16G010500 | C3H |
| 2909. | Glyma.16G011200.1.p | Glyma.16G011200 | Trihelix |
| 2910. | Glyma.16G012600.1.p | Glyma.16G012600 | ERF |
| 2911. | Glyma.16G012900.1.p | Glyma.16G012900 | M-type_MADS |
| 2912. | Glyma.16G016400.1.p | Glyma.16G016400 | NAC |
| 2913. | Glyma.16G016600.1.p | Glyma.16G016600 | NAC |
| 2914. | Glyma.16G016700.1.p | Glyma.16G016700 | NAC |
| 2915. | Glyma.16G017100.1.p | Glyma.16G017100 | B3 |
| 2916. | Glyma.16G017400.1.p | Glyma.16G017400 | MYB_related |
| 2917. | Glyma.16G017700.1.p | Glyma.16G017700 | bHLH |
| 2918. | Glyma.16G019400.1.p | Glyma.16G019400 | NAC |
| 2919. | Glyma.16G020500.1.p | Glyma.16G020500 | bHLH |
| 2920. | Glyma.16G021000.1.p | Glyma.16G021000 | HD-ZIP |
| 2921. | Glyma.16G022900.1.p | Glyma.16G022900 | Dof |
| 2922. | Glyma.16G023000.1.p | Glyma.16G023000 | MYB |
| 2923. | Glyma.16G023600.1.p | Glyma.16G023600 | ARF |
| 2924. | Glyma.16G023800.1.p | Glyma.16G023800 | ERF |
| 2925. | Glyma.16G023900.1.p | Glyma.16G023900 | bHLH |
| 2926. | Glyma.16G026400.1.p | Glyma.16G026400 | WRKY |
| 2927. | Glyma.16G029000.1.p | Glyma.16G029000 | bZIP |
| 2928. | Glyma.16G031400.1.p | Glyma.16G031400 | WRKY |
| 2929. | Glyma.16G031900.1.p | Glyma.16G031900 | WRKY |
| 2930. | Glyma.16G032600.1.p | Glyma.16G032600 | MYB_related |
| 2931. | Glyma.16G040000.1.p | Glyma.16G040000 | ERF |
| 2932. | Glyma.16G042300.1.p | Glyma.16G042300 | GATA |
| 2933. | Glyma.16G042900.1.p | Glyma.16G042900 | NAC |
| 2934. | Glyma.16G043200.1.p | Glyma.16G043200 | NAC |
| 2935. | Glyma.16G046300.1.p | Glyma.16G046300 | ERF |
| 2936. | Glyma.16G046700.1.p | Glyma.16G046700 | B3 |
| 2937. | Glyma.16G047600.1.p | Glyma.16G047600 | ERF |
| 2938. | Glyma.16G049400.1.p | Glyma.16G049400 | bHLH |
| 2939. | Glyma.16G050300.1.p | Glyma.16G050300 | B3 |
| 2940. | Glyma.16G050900.1.p | Glyma.16G050900 | CO-like |
| 2941. | Glyma.16G051800.1.p | Glyma.16G051800 | NAC |
| 2942. | Glyma.16G053000.1.p | Glyma.16G053000 | GRAS |
| 2943. | Glyma.16G053900.1.p | Glyma.16G053900 | TCP |
| 2944. | Glyma.16G054400.1.p | Glyma.16G054400 | WRKY |
| 2945. | Glyma.16G054500.1.p | Glyma.16G054500 | SBP |
| 2946. | Glyma.16G056000.1.p | Glyma.16G056000 | C2H2 |
| 2947. | Glyma.16G063700.1.p | Glyma.16G063700 | MYB |
| 2948. | Glyma.16G069300.1.p | Glyma.16G069300 | NAC |
| 2949. | Glyma.16G071700.1.p | Glyma.16G071700 | LBD |
| 2950. | Glyma.16G072700.1.p | Glyma.16G072700 | MYB |
| 2951. | Glyma.16G073000.1.p | Glyma.16G073000 | MYB |
| 2952. | Glyma.16G079600.1.p | Glyma.16G079600 | ERF |
| 2953. | Glyma.16G084000.1.p | Glyma.16G084000 | LBD |
| 2954. | Glyma.16G087300.1.p | Glyma.16G087300 | bHLH |
| 2955. | Glyma.16G090900.1.p | Glyma.16G090900 | bHLH |
| 2956. | Glyma.16G091300.1.p | Glyma.16G091300 | MIKC_MADS |
| 2957. | Glyma.16G091800.1.p | Glyma.16G091800 | HSF |
| 2958. | Glyma.16G092100.1.p | Glyma.16G092100 | MYB |
| 2959. | Glyma.16G092700.1.p | Glyma.16G092700 | bZIP |
| 2960. | Glyma.16G093600.1.p | Glyma.16G093600 | Trihelix |
| 2961. | Glyma.16G095200.1.p | Glyma.16G095200 | bZIP |
| 2962. | Glyma.16G097400.1.p | Glyma.16G097400 | SBP |
| 2963. | Glyma.16G101400.1.p | Glyma.16G101400 | ZF-HD |
| 2964. | Glyma.16G102000.1.p | Glyma.16G102000 | C2H2 |
| 2965. | Glyma.16G105600.1.p | Glyma.16G105600 | MIKC_MADS |
| 2966. | Glyma.16G118200.1.p | Glyma.16G118200 | FAR1 |
| 2967. | Glyma.16G128300.1.p | Glyma.16G128300 | MYB_related |
| 2968. | Glyma.16G130200.1.p | Glyma.16G130200 | NAC |
| 2969. | Glyma.16G132100.1.p | Glyma.16G132100 | SRS |
| 2970. | Glyma.16G133200.1.p | Glyma.16G133200 | bHLH |
| 2971. | Glyma.16G138400.1.p | Glyma.16G138400 | MYB_related |
| 2972. | Glyma.16G138500.1.p | Glyma.16G138500 | MYB_related |
| 2973. | Glyma.16G138700.1.p | Glyma.16G138700 | ZF-HD |
| 2974. | Glyma.16G139400.1.p | Glyma.16G139400 | GATA |
| 2975. | Glyma.16G141100.1.p | Glyma.16G141100 | C2H2 |
| 2976. | Glyma.16G141300.1.p | Glyma.16G141300 | GRAS |
| 2977. | Glyma.16G141500.1.p | Glyma.16G141500 | bZIP |
| 2978. | Glyma.16G142800.1.p | Glyma.16G142800 | TALE |
| 2979. | Glyma.16G145000.1.p | Glyma.16G145000 | Dof |
| 2980. | Glyma.16G147200.1.p | Glyma.16G147200 | bHLH |
| 2981. | Glyma.16G147500.1.p | Glyma.16G147500 | ERF |
| 2982. | Glyma.16G148700.1.p | Glyma.16G148700 | ERF |
| 2983. | Glyma.16G151500.1.p | Glyma.16G151500 | NAC |
| 2984. | Glyma.16G152100.1.p | Glyma.16G152100 | NAC |
| 2985. | Glyma.16G152200.1.p | Glyma.16G152200 | G2-like |
| 2986. | Glyma.16G152700.1.p | Glyma.16G152700 | GATA |
| 2987. | Glyma.16G154100.1.p | Glyma.16G154100 | ERF |
| 2988. | Glyma.16G155300.1.p | Glyma.16G155300 | GATA |
| 2989. | Glyma.16G156400.1.p | Glyma.16G156400 | C2H2 |
| 2990. | Glyma.16G156700.1.p | Glyma.16G156700 | GRAS |
| 2991. | Glyma.16G164800.1.p | Glyma.16G164800 | ERF |
| 2992. | Glyma.16G167500.1.p | Glyma.16G167500 | Trihelix |
| 2993. | Glyma.16G167600.1.p | Glyma.16G167600 | Trihelix |
| 2994. | Glyma.16G167700.1.p | Glyma.16G167700 | Trihelix |
| 2995. | Glyma.16G168400.1.p | Glyma.16G168400 | bZIP |
| 2996. | Glyma.16G176700.1.p | Glyma.16G176700 | WRKY |
| 2997. | Glyma.16G177000.1.p | Glyma.16G177000 | WRKY |
| 2998. | Glyma.16G178700.1.p | Glyma.16G178700 | MYB_related |
| 2999. | Glyma.16G179900.1.p | Glyma.16G179900 | GRAS |
| 3000. | Glyma.16G181700.1.p | Glyma.16G181700 | C2H2 |
| 3001. | Glyma.16G182400.1.p | Glyma.16G182400 | Nin-like |
| 3002. | Glyma.16G189400.1.p | Glyma.16G189400 | MYB |
| 3003. | Glyma.16G196200.1.p | Glyma.16G196200 | HSF |
| 3004. | Glyma.16G196700.1.p | Glyma.16G196700 | HD-ZIP |
| 3005. | Glyma.16G198800.1.p | Glyma.16G198800 | ARR-B |
| 3006. | Glyma.16G199000.1.p | Glyma.16G199000 | ERF |
| 3007. | Glyma.16G199300.1.p | Glyma.16G199300 | C3H |
| 3008. | Glyma.16G200700.1.p | Glyma.16G200700 | MIKC_MADS |
| 3009. | Glyma.16G201100.1.p | Glyma.16G201100 | Trihelix |
| 3010. | Glyma.16G201300.1.p | Glyma.16G201300 | bHLH |
| 3011. | Glyma.16G201400.1.p | Glyma.16G201400 | bHLH |
| 3012. | Glyma.16G217400.1.p | Glyma.16G217400 | NAC |
| 3013. | Glyma.16G217700.1.p | Glyma.16G217700 | MYB_related |
| 3014. | Glyma.16G217800.1.p | Glyma.16G217800 | HD-ZIP |
| 3015. | Glyma.16G218900.1.p | Glyma.16G218900 | MYB |
| 3016. | Glyma.16G219800.1.p | Glyma.16G219800 | WRKY |

| 3017. | Glyma.17G002200.1.p | Glyma.17G002200 | ZF-HD |
| --- | --- | --- | --- |
| 3018. | Glyma.17G002800.1.p | Glyma.17G002800 | NAC |
| 3019. | Glyma.17G005600.1.p | Glyma.17G005600 | NF-YB |
| 3020. | Glyma.17G007600.1.p | Glyma.17G007600 | GRAS |
| 3021. | Glyma.17G010200.1.p | Glyma.17G010200 | TALE |
| 3022. | Glyma.17G011400.1.p | Glyma.17G011400 | WRKY |
| 3023. | Glyma.17G014300.1.p | Glyma.17G014300 | M-type_MADS |
| 3024. | Glyma.17G023200.1.p | Glyma.17G023200 | bHLH |
| 3025. | Glyma.17G024300.1.p | Glyma.17G024300 | ERF |
| 3026. | Glyma.17G024400.1.p | Glyma.17G024400 | C3H |
| 3027. | Glyma.17G024600.1.p | Glyma.17G024600 | MYB_related |
| 3028. | Glyma.17G025300.1.p | Glyma.17G025300 | NF-YB |
| 3029. | Glyma.17G030600.1.p | Glyma.17G030600 | ARR-B |
| 3030. | Glyma.17G030900.1.p | Glyma.17G030900 | GATA |
| 3031. | Glyma.17G031400.1.p | Glyma.17G031400 | C2H2 |
| 3032. | Glyma.17G031600.1.p | Glyma.17G031600 | MYB |
| 3033. | Glyma.17G031900.1.p | Glyma.17G031900 | CAMTA |
| 3034. | Glyma.17G035400.1.p | Glyma.17G035400 | WRKY |
| 3035. | Glyma.17G037500.1.p | Glyma.17G037500 | MYB |
| 3036. | Glyma.17G038800.1.p | Glyma.17G038800 | CAMTA |
| 3037. | Glyma.17G039600.1.p | Glyma.17G039600 | SBP |
| 3038. | Glyma.17G042300.1.p | Glyma.17G042300 | WRKY |
| 3039. | Glyma.17G047100.1.p | Glyma.17G047100 | ARF |
| 3040. | Glyma.17G047300.1.p | Glyma.17G047300 | ERF |
| 3041. | Glyma.17G050200.1.p | Glyma.17G050200 | GRF |
| 3042. | Glyma.17G050500.1.p | Glyma.17G050500 | MYB |
| 3043. | Glyma.17G051400.1.p | Glyma.17G051400 | NF-YA |
| 3044. | Glyma.17G053700.1.p | Glyma.17G053700 | HSF |
| 3045. | Glyma.17G054000.1.p | Glyma.17G054000 | bHLH |
| 3046. | Glyma.17G054700.1.p | Glyma.17G054700 | C3H |
| 3047. | Glyma.17G055200.1.p | Glyma.17G055200 | GATA |
| 3048. | Glyma.17G056300.1.p | Glyma.17G056300 | HD-ZIP |
| 3049. | Glyma.17G057100.1.p | Glyma.17G057100 | WRKY |
| 3050. | Glyma.17G058600.1.p | Glyma.17G058600 | bHLH |
| 3051. | Glyma.17G059400.1.p | Glyma.17G059400 | HB-PHD |
| 3052. | Glyma.17G062600.1.p | Glyma.17G062600 | AP2 |
| 3053. | Glyma.17G064700.1.p | Glyma.17G064700 | C2H2 |
| 3054. | Glyma.17G065800.1.p | Glyma.17G065800 | MYB |
| 3055. | Glyma.17G066600.1.p | Glyma.17G066600 | CO-like |
| 3056. | Glyma.17G070800.1.p | Glyma.17G070800 | AP2 |
| 3057. | Glyma.17G074000.1.p | Glyma.17G074000 | WRKY |
| 3058. | Glyma.17G075200.1.p | Glyma.17G075200 | bHLH |
| 3059. | Glyma.17G076000.1.p | Glyma.17G076000 | ARR-B |
| 3060. | Glyma.17G076900.1.p | Glyma.17G076900 | C2H2 |
| 3061. | Glyma.17G079900.1.p | Glyma.17G079900 | TCP |
| 3062. | Glyma.17G080700.1.p | Glyma.17G080700 | SBP |
| 3063. | Glyma.17G080900.1.p | Glyma.17G080900 | MIKC_MADS |
| 3064. | Glyma.17G081200.1.p | Glyma.17G081200 | MIKC_MADS |
| 3065. | Glyma.17G081800.1.p | Glyma.17G081800 | Dof |
| 3066. | Glyma.17G082100.1.p | Glyma.17G082100 | bHLH |
| 3067. | Glyma.17G085600.1.p | Glyma.17G085600 | MYB_related |
| 3068. | Glyma.17G086300.1.p | Glyma.17G086300 | LBD |
| 3069. | Glyma.17G088600.1.p | Glyma.17G088600 | MYB |
| 3070. | Glyma.17G089300.1.p | Glyma.17G089300 | Dof |
| 3071. | Glyma.17G090500.1.p | Glyma.17G090500 | bHLH |
| 3072. | Glyma.17G093600.1.p | Glyma.17G093600 | E2F/DP |
| 3073. | Glyma.17G094400.1.p | Glyma.17G094400 | MYB_related |
| 3074. | Glyma.17G095000.1.p | Glyma.17G095000 | bHLH |
| 3075. | Glyma.17G096700.1.p | Glyma.17G096700 | HD-ZIP |
| 3076. | Glyma.17G097900.1.p | Glyma.17G097900 | WRKY |
| 3077. | Glyma.17G099100.1.p | Glyma.17G099100 | TCP |
| 3078. | Glyma.17G099800.1.p | Glyma.17G099800 | MYB |
| 3079. | Glyma.17G100000.1.p | Glyma.17G100000 | ZF-HD |
| 3080. | Glyma.17G100400.1.p | Glyma.17G100400 | Nin-like |
| 3081. | Glyma.17G101000.1.p | Glyma.17G101000 | Dof |
| 3082. | Glyma.17G101300.1.p | Glyma.17G101300 | M-type_MADS |
| 3083. | Glyma.17G101500.1.p | Glyma.17G101500 | NAC |
| 3084. | Glyma.17G104800.1.p | Glyma.17G104800 | TALE |
| 3085. | Glyma.17G113400.1.p | Glyma.17G113400 | YABBY |
| 3086. | Glyma.17G114500.1.p | Glyma.17G114500 | ERF |
| 3087. | Glyma.17G116200.1.p | Glyma.17G116200 | C3H |
| 3088. | Glyma.17G116700.1.p | Glyma.17G116700 | C3H |
| 3089. | Glyma.17G119600.1.p | Glyma.17G119600 | G2-like |
| 3090. | Glyma.17G121000.1.p | Glyma.17G121000 | MYB |
| 3091. | Glyma.17G121500.1.p | Glyma.17G121500 | TCP |
| 3092. | Glyma.17G123600.1.p | Glyma.17G123600 | NF-YB |
| 3093. | Glyma.17G124100.1.p | Glyma.17G124100 | ERF |
| 3094. | Glyma.17G126800.1.p | Glyma.17G126800 | C3H |
| 3095. | Glyma.17G127500.1.p | Glyma.17G127500 | GRAS |
| 3096. | Glyma.17G128500.1.p | Glyma.17G128500 | Trihelix |
| 3097. | Glyma.17G131200.1.p | Glyma.17G131200 | GRAS |
| 3098. | Glyma.17G131800.1.p | Glyma.17G131800 | ERF |
| 3099. | Glyma.17G131900.1.p | Glyma.17G131900 | ERF |
| 3100. | Glyma.17G132400.1.p | Glyma.17G132400 | TCP |
| 3101. | Glyma.17G132600.1.p | Glyma.17G132600 | TALE |
| 3102. | Glyma.17G132700.1.p | Glyma.17G132700 | MIKC_MADS |
| 3103. | Glyma.17G133800.1.p | Glyma.17G133800 | MYB |
| 3104. | Glyma.17G138100.1.p | Glyma.17G138100 | NAC |
| 3105. | Glyma.17G138200.1.p | Glyma.17G138200 | YABBY |
| 3106. | Glyma.17G142300.1.p | Glyma.17G142300 | C2H2 |
| 3107. | Glyma.17G143600.1.p | Glyma.17G143600 | MYB |
| 3108. | Glyma.17G143900.1.p | Glyma.17G143900 | ERF |
| 3109. | Glyma.17G144100.1.p | Glyma.17G144100 | MYB_related |
| 3110. | Glyma.17G144700.1.p | Glyma.17G144700 | HD-ZIP |
| 3111. | Glyma.17G145300.1.p | Glyma.17G145300 | ERF |
| 3112. | Glyma.17G145400.1.p | Glyma.17G145400 | ERF |
| 3113. | Glyma.17G146600.1.p | Glyma.17G146600 | GATA |
| 3114. | Glyma.17G150800.1.p | Glyma.17G150800 | SRS |
| 3115. | Glyma.17G152800.1.p | Glyma.17G152800 | ARR-B |
| 3116. | Glyma.17G154100.1.p | Glyma.17G154100 | NAC |
| 3117. | Glyma.17G154800.1.p | Glyma.17G154800 | LSD |
| 3118. | Glyma.17G155600.1.p | Glyma.17G155600 | C2H2 |
| 3119. | Glyma.17G155900.1.p | Glyma.17G155900 | bHLH |
| 3120. | Glyma.17G156000.1.p | Glyma.17G156000 | bHLH |
| 3121. | Glyma.17G156100.1.p | Glyma.17G156100 | bHLH |
| 3122. | Glyma.17G157600.1.p | Glyma.17G157600 | HD-ZIP |
| 3123. | Glyma.17G158000.1.p | Glyma.17G158000 | MYB |
| 3124. | Glyma.17G158300.1.p | Glyma.17G158300 | AP2 |
| 3125. | Glyma.17G158900.1.p | Glyma.17G158900 | bZIP |
| 3126. | Glyma.17G160500.1.p | Glyma.17G160500 | MYB_related |
| 3127. | Glyma.17G160800.1.p | Glyma.17G160800 | GRAS |
| 3128. | Glyma.17G162000.1.p | Glyma.17G162000 | LBD |
| 3129. | Glyma.17G162100.1.p | Glyma.17G162100 | MYB |
| 3130. | Glyma.17G162800.1.p | Glyma.17G162800 | GRAS |
| 3131. | Glyma.17G167100.1.p | Glyma.17G167100 | MYB_related |
| 3132. | Glyma.17G168900.1.p | Glyma.17G168900 | WRKY |
| 3133. | Glyma.17G169800.1.p | Glyma.17G169800 | ERF |
| 3134. | Glyma.17G170100.1.p | Glyma.17G170100 | ERF |
| 3135. | Glyma.17G170300.1.p | Glyma.17G170300 | AP2 |
| 3136. | Glyma.17G172400.1.p | Glyma.17G172400 | bHLH |
| 3137. | Glyma.17G173100.1.p | Glyma.17G173100 | bHLH |
| 3138. | Glyma.17G174900.1.p | Glyma.17G174900 | HSF |
| 3139. | Glyma.17G175500.1.p | Glyma.17G175500 | B3 |
| 3140. | Glyma.17G178200.1.p | Glyma.17G178200 | LBD |
| 3141. | Glyma.17G178500.1.p | Glyma.17G178500 | G2-like |
| 3142. | Glyma.17G180600.1.p | Glyma.17G180600 | Dof |
| 3143. | Glyma.17G180800.1.p | Glyma.17G180800 | bHLH |
| 3144. | Glyma.17G185000.1.p | Glyma.17G185000 | NAC |
| 3145. | Glyma.17G188500.1.p | Glyma.17G188500 | bZIP |
| 3146. | Glyma.17G190900.1.p | Glyma.17G190900 | MYB |
| 3147. | Glyma.17G192800.1.p | Glyma.17G192800 | GATA |
| 3148. | Glyma.17G194100.1.p | Glyma.17G194100 | ERF |
| 3149. | Glyma.17G197300.1.p | Glyma.17G197300 | bZIP |
| 3150. | Glyma.17G197500.1.p | Glyma.17G197500 | WRKY |
| 3151. | Glyma.17G201800.1.p | Glyma.17G201800 | FAR1 |
| 3152. | Glyma.17G208500.1.p | Glyma.17G208500 | CPP |
| 3153. | Glyma.17G210500.1.p | Glyma.17G210500 | ERF |
| 3154. | Glyma.17G212900.1.p | Glyma.17G212900 | bHLH |
| 3155. | Glyma.17G213800.1.p | Glyma.17G213800 | MYB_related |
| 3156. | Glyma.17G213900.1.p | Glyma.17G213900 | MYB_related |
| 3157. | Glyma.17G215500.1.p | Glyma.17G215500 | TALE |
| 3158. | Glyma.17G216100.1.p | Glyma.17G216100 | ERF |
| 3159. | Glyma.17G216800.1.p | Glyma.17G216800 | C2H2 |
| 3160. | Glyma.17G217100.1.p | Glyma.17G217100 | ARR-B |
| 3161. | Glyma.17G219700.1.p | Glyma.17G219700 | ERF |
| 3162. | Glyma.17G222300.1.p | Glyma.17G222300 | WRKY |
| 3163. | Glyma.17G222500.1.p | Glyma.17G222500 | WRKY |
| 3164. | Glyma.17G223100.1.p | Glyma.17G223100 | bHLH |
| 3165. | Glyma.17G224800.1.p | Glyma.17G224800 | WRKY |
| 3166. | Glyma.17G227600.1.p | Glyma.17G227600 | HSF |
| 3167. | Glyma.17G228000.1.p | Glyma.17G228000 | C2H2 |
| 3168. | Glyma.17G228700.1.p | Glyma.17G228700 | GATA |
| 3169. | Glyma.17G229800.1.p | Glyma.17G229800 | MYB_related |
| 3170. | Glyma.17G230000.1.p | Glyma.17G230000 | TALE |
| 3171. | Glyma.17G231900.1.p | Glyma.17G231900 | MYB |
| 3172. | Glyma.17G232600.1.p | Glyma.17G232600 | GRF |
| 3173. | Glyma.17G232700.1.p | Glyma.17G232700 | GRF |
| 3174. | Glyma.17G236100.1.p | Glyma.17G236100 | bHLH |
| 3175. | Glyma.17G236200.1.p | Glyma.17G236200 | C2H2 |
| 3176. | Glyma.17G236300.1.p | Glyma.17G236300 | LBD |
| 3177. | Glyma.17G237900.1.p | Glyma.17G237900 | MYB |
| 3178. | Glyma.17G239200.1.p | Glyma.17G239200 | WRKY |
| 3179. | Glyma.17G239900.1.p | Glyma.17G239900 | C2H2 |
| 3180. | Glyma.17G240100.1.p | Glyma.17G240100 | ERF |
| 3181. | Glyma.17G240300.1.p | Glyma.17G240300 | WOX |
| 3182. | Glyma.17G240700.1.p | Glyma.17G240700 | NAC |
| 3183. | Glyma.17G241000.1.p | Glyma.17G241000 | bHLH |
| 3184. | Glyma.17G241300.1.p | Glyma.17G241300 | HSF |
| 3185. | Glyma.17G245200.1.p | Glyma.17G245200 | MYB |
| 3186. | Glyma.17G245900.1.p | Glyma.17G245900 | B3 |
| 3187. | Glyma.17G246000.1.p | Glyma.17G246000 | B3 |
| 3188. | Glyma.17G246200.1.p | Glyma.17G246200 | G2-like |
| 3189. | Glyma.17G248900.1.p | Glyma.17G248900 | BES1 |
| 3190. | Glyma.17G253200.1.p | Glyma.17G253200 | bZIP |
| 3191. | Glyma.17G254000.1.p | Glyma.17G254000 | TALE |
| 3192. | Glyma.17G254600.1.p | Glyma.17G254600 | ERF |
| 3193. | Glyma.17G255000.1.p | Glyma.17G255000 | SRS |
| 3194. | Glyma.17G255100.1.p | Glyma.17G255100 | DBB |
| 3195. | Glyma.17G255800.1.p | Glyma.17G255800 | bZIP |
| 3196. | Glyma.17G256500.1.p | Glyma.17G256500 | ARF |
| 3197. | Glyma.17G257700.1.p | Glyma.17G257700 | C2H2 |

| 3198. | Glyma.18G003600.1.p | Glyma.18G003600 | C2H2 |
| --- | --- | --- | --- |
| 3199. | Glyma.18G004700.1.p | Glyma.18G004700 | MIKC_MADS |
| 3200. | Glyma.18G005100.1.p | Glyma.18G005100 | CAMTA |
| 3201. | Glyma.18G005600.1.p | Glyma.18G005600 | SBP |
| 3202. | Glyma.18G007100.1.p | Glyma.18G007100 | NF-YC |
| 3203. | Glyma.18G010000.1.p | Glyma.18G010000 | Trihelix |
| 3204. | Glyma.18G010800.1.p | Glyma.18G010800 | ARR-B |
| 3205. | Glyma.18G014900.1.p | Glyma.18G014900 | HD-ZIP |
| 3206. | Glyma.18G016700.1.p | Glyma.18G016700 | C2H2 |
| 3207. | Glyma.18G018200.1.p | Glyma.18G018200 | ERF |
| 3208. | Glyma.18G018400.1.p | Glyma.18G018400 | EIL |
| 3209. | Glyma.18G020900.1.p | Glyma.18G020900 | bZIP |
| 3210. | Glyma.18G024400.1.p | Glyma.18G024400 | LBD |
| 3211. | Glyma.18G025600.1.p | Glyma.18G025600 | LBD |
| 3212. | Glyma.18G025800.1.p | Glyma.18G025800 | bHLH |
| 3213. | Glyma.18G027000.1.p | Glyma.18G027000 | NF-X1 |
| 3214. | Glyma.18G029700.1.p | Glyma.18G029700 | WOX |
| 3215. | Glyma.18G030300.1.p | Glyma.18G030300 | MYB |
| 3216. | Glyma.18G036000.1.p | Glyma.18G036000 | C2H2 |
| 3217. | Glyma.18G036800.1.p | Glyma.18G036800 | MYB_related |
| 3218. | Glyma.18G039200.1.p | Glyma.18G039200 | bHLH |
| 3219. | Glyma.18G040000.1.p | Glyma.18G040000 | GRAS |
| 3220. | Glyma.18G040700.1.p | Glyma.18G040700 | MYB |
| 3221. | Glyma.18G042000.1.p | Glyma.18G042000 | M-type_MADS |
| 3222. | Glyma.18G042300.1.p | Glyma.18G042300 | C2H2 |
| 3223. | Glyma.18G043000.1.p | Glyma.18G043000 | G2-like |
| 3224. | Glyma.18G043900.1.p | Glyma.18G043900 | NAC |
| 3225. | Glyma.18G044200.1.p | Glyma.18G044200 | MYB_related |
| 3226. | Glyma.18G046800.1.p | Glyma.18G046800 | ARF |
| 3227. | Glyma.18G052100.1.p | Glyma.18G052100 | B3 |
| 3228. | Glyma.18G052500.1.p | Glyma.18G052500 | bZIP |
| 3229. | Glyma.18G052700.1.p | Glyma.18G052700 | MYB_related |
| 3230. | Glyma.18G052800.1.p | Glyma.18G052800 | M-type_MADS |
| 3231. | Glyma.18G052900.1.p | Glyma.18G052900 | M-type_MADS |
| 3232. | Glyma.18G053200.1.p | Glyma.18G053200 | M-type_MADS |
| 3233. | Glyma.18G053300.1.p | Glyma.18G053300 | M-type_MADS |
| 3234. | Glyma.18G053400.1.p | Glyma.18G053400 | M-type_MADS |
| 3235. | Glyma.18G053500.1.p | Glyma.18G053500 | M-type_MADS |
| 3236. | Glyma.18G053600.1.p | Glyma.18G053600 | M-type_MADS |
| 3237. | Glyma.18G053800.1.p | Glyma.18G053800 | M-type_MADS |
| 3238. | Glyma.18G056600.1.p | Glyma.18G056600 | WRKY |
| 3239. | Glyma.18G058000.1.p | Glyma.18G058000 | LBD |
| 3240. | Glyma.18G062200.1.p | Glyma.18G062200 | C3H |
| 3241. | Glyma.18G065200.1.p | Glyma.18G065200 | MYB |
| 3242. | Glyma.18G066100.1.p | Glyma.18G066100 | C2H2 |
| 3243. | Glyma.18G066400.1.p | Glyma.18G066400 | MYB |
| 3244. | Glyma.18G071000.1.p | Glyma.18G071000 | NF-YA |
| 3245. | Glyma.18G071600.1.p | Glyma.18G071600 | MYB |
| 3246. | Glyma.18G073300.1.p | Glyma.18G073300 | MYB_related |
| 3247. | Glyma.18G075100.1.p | Glyma.18G075100 | MYB_related |
| 3248. | Glyma.18G077500.1.p | Glyma.18G077500 | NF-YB |
| 3249. | Glyma.18G079800.1.p | Glyma.18G079800 | B3 |
| 3250. | Glyma.18G081100.1.p | Glyma.18G081100 | GRAS |
| 3251. | Glyma.18G081200.1.p | Glyma.18G081200 | WRKY |
| 3252. | Glyma.18G091600.1.p | Glyma.18G091600 | ERF |
| 3253. | Glyma.18G092200.1.p | Glyma.18G092200 | WRKY |
| 3254. | Glyma.18G095600.1.p | Glyma.18G095600 | MYB |
| 3255. | Glyma.18G105800.1.p | Glyma.18G105800 | MIKC_MADS |
| 3256. | Glyma.18G110700.1.p | Glyma.18G110700 | NAC |
| 3257. | Glyma.18G113400.1.p | Glyma.18G113400 | G2-like |
| 3258. | Glyma.18G115700.1.p | Glyma.18G115700 | bHLH |
| 3259. | Glyma.18G117100.1.p | Glyma.18G117100 | bZIP |
| 3260. | Glyma.18G118400.1.p | Glyma.18G118400 | HB-other |
| 3261. | Glyma.18G119300.1.p | Glyma.18G119300 | NAC |
| 3262. | Glyma.18G121400.1.p | Glyma.18G121400 | TCP |
| 3263. | Glyma.18G123100.1.p | Glyma.18G123100 | bHLH |
| 3264. | Glyma.18G123500.1.p | Glyma.18G123500 | HD-ZIP |
| 3265. | Glyma.18G123600.1.p | Glyma.18G123600 | MYB |
| 3266. | Glyma.18G124700.1.p | Glyma.18G124700 | WRKY |
| 3267. | Glyma.18G124800.1.p | Glyma.18G124800 | Whirly |
| 3268. | Glyma.18G125200.1.p | Glyma.18G125200 | AP2 |
| 3269. | Glyma.18G126800.1.p | Glyma.18G126800 | HD-ZIP |
| 3270. | Glyma.18G129900.1.p | Glyma.18G129900 | MYB_related |
| 3271. | Glyma.18G132400.1.p | Glyma.18G132400 | FAR1 |
| 3272. | Glyma.18G133900.1.p | Glyma.18G133900 | FAR1 |
| 3273. | Glyma.18G134200.1.p | Glyma.18G134200 | MYB |
| 3274. | Glyma.18G140400.1.p | Glyma.18G140400 | YABBY |
| 3275. | Glyma.18G141200.1.p | Glyma.18G141200 | TALE |
| 3276. | Glyma.18G143800.1.p | Glyma.18G143800 | M-type_MADS |
| 3277. | Glyma.18G144700.1.p | Glyma.18G144700 | ERF |
| 3278. | Glyma.18G145400.1.p | Glyma.18G145400 | ZF-HD |
| 3279. | Glyma.18G148000.1.p | Glyma.18G148000 | AP2 |
| 3280. | Glyma.18G150800.1.p | Glyma.18G150800 | Dof |
| 3281. | Glyma.18G151400.1.p | Glyma.18G151400 | MYB |
| 3282. | Glyma.18G154100.1.p | Glyma.18G154100 | FAR1 |
| 3283. | Glyma.18G155000.1.p | Glyma.18G155000 | B3 |
| 3284. | Glyma.18G156000.1.p | Glyma.18G156000 | MYB_related |
| 3285. | Glyma.18G156900.1.p | Glyma.18G156900 | bHLH |
| 3286. | Glyma.18G159900.1.p | Glyma.18G159900 | ERF |
| 3287. | Glyma.18G176100.1.p | Glyma.18G176100 | B3 |
| 3288. | Glyma.18G176300.1.p | Glyma.18G176300 | Dof |
| 3289. | Glyma.18G178200.1.p | Glyma.18G178200 | FAR1 |
| 3290. | Glyma.18G178400.1.p | Glyma.18G178400 | FAR1 |
| 3291. | Glyma.18G178700.1.p | Glyma.18G178700 | FAR1 |
| 3292. | Glyma.18G179300.1.p | Glyma.18G179300 | WOX |
| 3293. | Glyma.18G179400.1.p | Glyma.18G179400 | FAR1 |
| 3294. | Glyma.18G181100.1.p | Glyma.18G181100 | MYB |
| 3295. | Glyma.18G181300.1.p | Glyma.18G181300 | MYB |
| 3296. | Glyma.18G182400.1.p | Glyma.18G182400 | MYB_related |
| 3297. | Glyma.18G182500.1.p | Glyma.18G182500 | GRAS |
| 3298. | Glyma.18G183100.1.p | Glyma.18G183100 | WRKY |
| 3299. | Glyma.18G184500.1.p | Glyma.18G184500 | ARF |
| 3300. | Glyma.18G185900.1.p | Glyma.18G185900 | C2H2 |
| 3301. | Glyma.18G186000.1.p | Glyma.18G186000 | C2H2 |
| 3302. | Glyma.18G189700.1.p | Glyma.18G189700 | TALE |
| 3303. | Glyma.18G191200.1.p | Glyma.18G191200 | MYB |
| 3304. | Glyma.18G197500.1.p | Glyma.18G197500 | MYB |
| 3305. | Glyma.18G201800.1.p | Glyma.18G201800 | G2-like |
| 3306. | Glyma.18G202400.1.p | Glyma.18G202400 | Trihelix |
| 3307. | Glyma.18G204700.1.p | Glyma.18G204700 | bHLH |
| 3308. | Glyma.18G204900.1.p | Glyma.18G204900 | G2-like |
| 3309. | Glyma.18G205100.1.p | Glyma.18G205100 | GRAS |
| 3310. | Glyma.18G206600.1.p | Glyma.18G206600 | ERF |
| 3311. | Glyma.18G208800.1.p | Glyma.18G208800 | WRKY |
| 3312. | Glyma.18G210700.1.p | Glyma.18G210700 | bHLH |
| 3313. | Glyma.18G213200.1.p | Glyma.18G213200 | WRKY |
| 3314. | Glyma.18G220100.1.p | Glyma.18G220100 | GRAS |
| 3315. | Glyma.18G221000.1.p | Glyma.18G221000 | LBD |
| 3316. | Glyma.18G224100.1.p | Glyma.18G224100 | CPP |
| 3317. | Glyma.18G224300.1.p | Glyma.18G224300 | MIKC_MADS |
| 3318. | Glyma.18G224500.1.p | Glyma.18G224500 | MIKC_MADS |
| 3319. | Glyma.18G225800.1.p | Glyma.18G225800 | HD-ZIP |
| 3320. | Glyma.18G230600.1.p | Glyma.18G230600 | MYB |
| 3321. | Glyma.18G232400.1.p | Glyma.18G232400 | BES1 |
| 3322. | Glyma.18G234900.1.p | Glyma.18G234900 | MYB_related |
| 3323. | Glyma.18G237700.1.p | Glyma.18G237700 | MYB_related |
| 3324. | Glyma.18G238200.1.p | Glyma.18G238200 | WRKY |
| 3325. | Glyma.18G238600.1.p | Glyma.18G238600 | WRKY |
| 3326. | Glyma.18G241300.1.p | Glyma.18G241300 | C3H |
| 3327. | Glyma.18G242000.1.p | Glyma.18G242000 | WRKY |
| 3328. | Glyma.18G244600.1.p | Glyma.18G244600 | AP2 |
| 3329. | Glyma.18G246000.1.p | Glyma.18G246000 | bHLH |
| 3330. | Glyma.18G246100.1.p | Glyma.18G246100 | bHLH |
| 3331. | Glyma.18G246200.1.p | Glyma.18G246200 | bHLH |
| 3332. | Glyma.18G252200.1.p | Glyma.18G252200 | ERF |
| 3333. | Glyma.18G252300.1.p | Glyma.18G252300 | ERF |
| 3334. | Glyma.18G252400.1.p | Glyma.18G252400 | ERF |
| 3335. | Glyma.18G253800.1.p | Glyma.18G253800 | HD-ZIP |
| 3336. | Glyma.18G256000.1.p | Glyma.18G256000 | AP2 |
| 3337. | Glyma.18G256500.1.p | Glyma.18G256500 | WRKY |
| 3338. | Glyma.18G257200.1.p | Glyma.18G257200 | ZF-HD |
| 3339. | Glyma.18G257300.1.p | Glyma.18G257300 | ZF-HD |
| 3340. | Glyma.18G258400.1.p | Glyma.18G258400 | HD-ZIP |
| 3341. | Glyma.18G258700.1.p | Glyma.18G258700 | bHLH |
| 3342. | Glyma.18G259100.1.p | Glyma.18G259100 | MYB |
| 3343. | Glyma.18G259200.1.p | Glyma.18G259200 | ZF-HD |
| 3344. | Glyma.18G259300.1.p | Glyma.18G259300 | ZF-HD |
| 3345. | Glyma.18G260500.1.p | Glyma.18G260500 | Dof |
| 3346. | Glyma.18G261300.1.p | Glyma.18G261300 | NAC |
| 3347. | Glyma.18G261400.1.p | Glyma.18G261400 | MYB |
| 3348. | Glyma.18G261700.1.p | Glyma.18G261700 | MYB |
| 3349. | Glyma.18G262000.1.p | Glyma.18G262000 | MYB |
| 3350. | Glyma.18G262800.1.p | Glyma.18G262800 | ERF |
| 3351. | Glyma.18G263400.1.p | Glyma.18G263400 | WRKY |
| 3352. | Glyma.18G268700.1.p | Glyma.18G268700 | TCP |
| 3353. | Glyma.18G273300.1.p | Glyma.18G273300 | MYB |
| 3354. | Glyma.18G273400.1.p | Glyma.18G273400 | MYB |
| 3355. | Glyma.18G273500.1.p | Glyma.18G273500 | MIKC_MADS |
| 3356. | Glyma.18G273600.1.p | Glyma.18G273600 | MIKC_MADS |
| 3357. | Glyma.18G275400.1.p | Glyma.18G275400 | bHLH |
| 3358. | Glyma.18G277100.1.p | Glyma.18G277100 | bZIP |
| 3359. | Glyma.18G278100.1.p | Glyma.18G278100 | CO-like |
| 3360. | Glyma.18G280700.1.p | Glyma.18G280700 | TCP |
| 3361. | Glyma.18G281400.1.p | Glyma.18G281400 | ERF |
| 3362. | Glyma.18G281800.1.p | Glyma.18G281800 | Trihelix |
| 3363. | Glyma.18G287200.1.p | Glyma.18G287200 | C2H2 |
| 3364. | Glyma.18G288000.1.p | Glyma.18G288000 | WOX |
| 3365. | Glyma.18G289700.1.p | Glyma.18G289700 | Dof |
| 3366. | Glyma.18G297100.1.p | Glyma.18G297100 | LBD |
| 3367. | Glyma.18G301500.1.p | Glyma.18G301500 | NAC |

| 3368. | Glyma.19G002100.1.p | Glyma.19G002100 | MYB_related |
| --- | --- | --- | --- |
| 3369. | Glyma.19G002900.1.p | Glyma.19G002900 | NAC |
| 3370. | Glyma.19G004900.1.p | Glyma.19G004900 | LBD |
| 3371. | Glyma.19G006500.1.p | Glyma.19G006500 | MYB |
| 3372. | Glyma.19G010100.1.p | Glyma.19G010100 | HD-ZIP |
| 3373. | Glyma.19G017900.1.p | Glyma.19G017900 | MYB |
| 3374. | Glyma.19G020600.1.p | Glyma.19G020600 | WRKY |
| 3375. | Glyma.19G021100.1.p | Glyma.19G021100 | HD-ZIP |
| 3376. | Glyma.19G021400.1.p | Glyma.19G021400 | bHLH |
| 3377. | Glyma.19G021900.1.p | Glyma.19G021900 | NAC |
| 3378. | Glyma.19G022100.1.p | Glyma.19G022100 | MYB |
| 3379. | Glyma.19G022200.1.p | Glyma.19G022200 | ZF-HD |
| 3380. | Glyma.19G023200.1.p | Glyma.19G023200 | Dof |
| 3381. | Glyma.19G024500.1.p | Glyma.19G024500 | NAC |
| 3382. | Glyma.19G024700.1.p | Glyma.19G024700 | MYB |
| 3383. | Glyma.19G025000.1.p | Glyma.19G025000 | MYB |
| 3384. | Glyma.19G026000.1.p | Glyma.19G026000 | ERF |
| 3385. | Glyma.19G030900.1.p | Glyma.19G030900 | TCP |
| 3386. | Glyma.19G034500.1.p | Glyma.19G034500 | MIKC_MADS |
| 3387. | Glyma.19G034600.1.p | Glyma.19G034600 | MIKC_MADS |
| 3388. | Glyma.19G035500.1.p | Glyma.19G035500 | ERF |
| 3389. | Glyma.19G037900.1.p | Glyma.19G037900 | bZIP |
| 3390. | Glyma.19G038100.1.p | Glyma.19G038100 | MYB |
| 3391. | Glyma.19G039000.1.p | Glyma.19G039000 | CO-like |
| 3392. | Glyma.19G040300.1.p | Glyma.19G040300 | G2-like |
| 3393. | Glyma.19G044400.1.p | Glyma.19G044400 | TCP |
| 3394. | Glyma.19G045900.1.p | Glyma.19G045900 | M-type_MADS |
| 3395. | Glyma.19G046300.1.p | Glyma.19G046300 | LBD |
| 3396. | Glyma.19G047500.1.p | Glyma.19G047500 | M-type_MADS |
| 3397. | Glyma.19G047800.1.p | Glyma.19G047800 | G2-like |
| 3398. | Glyma.19G047900.1.p | Glyma.19G047900 | ARR-B |
| 3399. | Glyma.19G049400.1.p | Glyma.19G049400 | ARR-B |
| 3400. | Glyma.19G052000.1.p | Glyma.19G052000 | G2-like |
| 3401. | Glyma.19G055800.1.p | Glyma.19G055800 | MYB |
| 3402. | Glyma.19G056400.1.p | Glyma.19G056400 | NAC |
| 3403. | Glyma.19G060700.1.p | Glyma.19G060700 | MYB_related |
| 3404. | Glyma.19G061300.1.p | Glyma.19G061300 | MYB |
| 3405. | Glyma.19G061600.1.p | Glyma.19G061600 | MYB |
| 3406. | Glyma.19G062000.1.p | Glyma.19G062000 | MYB |
| 3407. | Glyma.19G063600.1.p | Glyma.19G063600 | LBD |
| 3408. | Glyma.19G067900.1.p | Glyma.19G067900 | bZIP |
| 3409. | Glyma.19G083700.1.p | Glyma.19G083700 | MYB_related |
| 3410. | Glyma.19G084000.1.p | Glyma.19G084000 | MYB_related |
| 3411. | Glyma.19G085700.1.p | Glyma.19G085700 | MYB_related |
| 3412. | Glyma.19G094000.1.p | Glyma.19G094000 | SBP |
| 3413. | Glyma.19G094100.1.p | Glyma.19G094100 | WRKY |
| 3414. | Glyma.19G095300.1.p | Glyma.19G095300 | TCP |
| 3415. | Glyma.19G095600.1.p | Glyma.19G095600 | FAR1 |
| 3416. | Glyma.19G096400.1.p | Glyma.19G096400 | GRAS |
| 3417. | Glyma.19G097700.1.p | Glyma.19G097700 | NAC |
| 3418. | Glyma.19G099700.1.p | Glyma.19G099700 | CO-like |
| 3419. | Glyma.19G100900.1.p | Glyma.19G100900 | B3 |
| 3420. | Glyma.19G102000.1.p | Glyma.19G102000 | bHLH |
| 3421. | Glyma.19G104200.1.p | Glyma.19G104200 | ERF |
| 3422. | Glyma.19G105300.1.p | Glyma.19G105300 | B3 |
| 3423. | Glyma.19G108800.1.p | Glyma.19G108800 | NAC |
| 3424. | Glyma.19G109100.1.p | Glyma.19G109100 | NAC |
| 3425. | Glyma.19G110200.1.p | Glyma.19G110200 | GATA |
| 3426. | Glyma.19G113100.1.p | Glyma.19G113100 | ERF |
| 3427. | Glyma.19G118000.1.p | Glyma.19G118000 | Dof |
| 3428. | Glyma.19G118400.1.p | Glyma.19G118400 | WOX |
| 3429. | Glyma.19G118500.1.p | Glyma.19G118500 | MYB |
| 3430. | Glyma.19G119300.1.p | Glyma.19G119300 | MYB |
| 3431. | Glyma.19G122500.1.p | Glyma.19G122500 | LBD |
| 3432. | Glyma.19G122700.1.p | Glyma.19G122700 | G2-like |
| 3433. | Glyma.19G122800.1.p | Glyma.19G122800 | bZIP |
| 3434. | Glyma.19G126800.1.p | Glyma.19G126800 | bZIP |
| 3435. | Glyma.19G127000.1.p | Glyma.19G127000 | G2-like |
| 3436. | Glyma.19G128900.1.p | Glyma.19G128900 | bHLH |
| 3437. | Glyma.19G130200.1.p | Glyma.19G130200 | bZIP |
| 3438. | Glyma.19G132500.1.p | Glyma.19G132500 | bHLH |
| 3439. | Glyma.19G132600.1.p | Glyma.19G132600 | bHLH |
| 3440. | Glyma.19G137800.1.p | Glyma.19G137800 | HSF |
| 3441. | Glyma.19G138000.1.p | Glyma.19G138000 | AP2 |
| 3442. | Glyma.19G139100.1.p | Glyma.19G139100 | FAR1 |
| 3443. | Glyma.19G140400.1.p | Glyma.19G140400 | C2H2 |
| 3444. | Glyma.19G141700.1.p | Glyma.19G141700 | C3H |
| 3445. | Glyma.19G142000.1.p | Glyma.19G142000 | ERF |
| 3446. | Glyma.19G142600.1.p | Glyma.19G142600 | C2H2 |
| 3447. | Glyma.19G143900.1.p | Glyma.19G143900 | bHLH |
| 3448. | Glyma.19G144200.1.p | Glyma.19G144200 | bHLH |
| 3449. | Glyma.19G145300.1.p | Glyma.19G145300 | bZIP |
| 3450. | Glyma.19G146000.1.p | Glyma.19G146000 | SBP |
| 3451. | Glyma.19G146500.1.p | Glyma.19G146500 | G2-like |
| 3452. | Glyma.19G146600.1.p | Glyma.19G146600 | G2-like |
| 3453. | Glyma.19G147100.1.p | Glyma.19G147100 | C3H |
| 3454. | Glyma.19G149000.1.p | Glyma.19G149000 | HD-ZIP |
| 3455. | Glyma.19G155300.1.p | Glyma.19G155300 | bHLH |
| 3456. | Glyma.19G159500.1.p | Glyma.19G159500 | HSF |
| 3457. | Glyma.19G159600.1.p | Glyma.19G159600 | C2H2 |
| 3458. | Glyma.19G160900.1.p | Glyma.19G160900 | bHLH |
| 3459. | Glyma.19G162500.1.p | Glyma.19G162500 | C2H2 |
| 3460. | Glyma.19G162900.1.p | Glyma.19G162900 | LBD |
| 3461. | Glyma.19G163000.1.p | Glyma.19G163000 | LBD |
| 3462. | Glyma.19G163300.1.p | Glyma.19G163300 | bHLH |
| 3463. | Glyma.19G163700.1.p | Glyma.19G163700 | ERF |
| 3464. | Glyma.19G163800.1.p | Glyma.19G163800 | ERF |
| 3465. | Glyma.19G163900.1.p | Glyma.19G163900 | ERF |
| 3466. | Glyma.19G164100.1.p | Glyma.19G164100 | ERF |
| 3467. | Glyma.19G164600.1.p | Glyma.19G164600 | MYB |
| 3468. | Glyma.19G165600.1.p | Glyma.19G165600 | NAC |
| 3469. | Glyma.19G167500.1.p | Glyma.19G167500 | G2-like |
| 3470. | Glyma.19G174000.1.p | Glyma.19G174000 | C2H2 |
| 3471. | Glyma.19G174100.1.p | Glyma.19G174100 | C2H2 |
| 3472. | Glyma.19G174200.1.p | Glyma.19G174200 | C2H2 |
| 3473. | Glyma.19G177400.1.p | Glyma.19G177400 | WRKY |
| 3474. | Glyma.19G178000.1.p | Glyma.19G178000 | MYB_related |
| 3475. | Glyma.19G178200.1.p | Glyma.19G178200 | AP2 |
| 3476. | Glyma.19G178500.1.p | Glyma.19G178500 | NF-YB |
| 3477. | Glyma.19G179700.1.p | Glyma.19G179700 | S1Fa-like |
| 3478. | Glyma.19G180300.1.p | Glyma.19G180300 | NAC |
| 3479. | Glyma.19G180400.1.p | Glyma.19G180400 | C2H2 |
| 3480. | Glyma.19G181900.1.p | Glyma.19G181900 | ARF |
| 3481. | Glyma.19G184500.1.p | Glyma.19G184500 | MYB |
| 3482. | Glyma.19G186200.1.p | Glyma.19G186200 | C2H2 |
| 3483. | Glyma.19G189700.1.p | Glyma.19G189700 | HD-ZIP |
| 3484. | Glyma.19G190000.1.p | Glyma.19G190000 | Trihelix |
| 3485. | Glyma.19G191700.1.p | Glyma.19G191700 | HSF |
| 3486. | Glyma.19G192300.1.p | Glyma.19G192300 | Trihelix |
| 3487. | Glyma.19G192400.1.p | Glyma.19G192400 | ERF |
| 3488. | Glyma.19G192700.1.p | Glyma.19G192700 | GRF |
| 3489. | Glyma.19G193400.1.p | Glyma.19G193400 | bZIP |
| 3490. | Glyma.19G194500.1.p | Glyma.19G194500 | bZIP |
| 3491. | Glyma.19G195800.1.p | Glyma.19G195800 | NAC |
| 3492. | Glyma.19G196600.1.p | Glyma.19G196600 | B3 |
| 3493. | Glyma.19G199200.1.p | Glyma.19G199200 | Dof |
| 3494. | Glyma.19G199500.1.p | Glyma.19G199500 | TALE |
| 3495. | Glyma.19G200300.1.p | Glyma.19G200300 | Dof |
| 3496. | Glyma.19G200800.1.p | Glyma.19G200800 | NF-YA |
| 3497. | Glyma.19G206100.1.p | Glyma.19G206100 | ARF |
| 3498. | Glyma.19G207100.1.p | Glyma.19G207100 | CO-like |
| 3499. | Glyma.19G208700.1.p | Glyma.19G208700 | HB-other |
| 3500. | Glyma.19G208900.1.p | Glyma.19G208900 | C2H2 |
| 3501. | Glyma.19G213100.1.p | Glyma.19G213100 | ERF |
| 3502. | Glyma.19G214600.1.p | Glyma.19G214600 | C2H2 |
| 3503. | Glyma.19G214900.1.p | Glyma.19G214900 | MYB |
| 3504. | Glyma.19G216200.1.p | Glyma.19G216200 | bZIP |
| 3505. | Glyma.19G216700.1.p | Glyma.19G216700 | GRAS |
| 3506. | Glyma.19G217000.1.p | Glyma.19G217000 | WRKY |
| 3507. | Glyma.19G217800.1.p | Glyma.19G217800 | WRKY |
| 3508. | Glyma.19G218500.1.p | Glyma.19G218500 | MYB_related |
| 3509. | Glyma.19G218800.1.p | Glyma.19G218800 | MYB |
| 3510. | Glyma.19G219000.1.p | Glyma.19G219000 | MYB |
| 3511. | Glyma.19G221700.1.p | Glyma.19G221700 | WRKY |
| 3512. | Glyma.19G222000.1.p | Glyma.19G222000 | bHLH |
| 3513. | Glyma.19G222200.1.p | Glyma.19G222200 | MYB |
| 3514. | Glyma.19G224600.1.p | Glyma.19G224600 | MYB |
| 3515. | Glyma.19G224700.1.p | Glyma.19G224700 | bHLH |
| 3516. | Glyma.19G225400.1.p | Glyma.19G225400 | C2H2 |
| 3517. | Glyma.19G228300.1.p | Glyma.19G228300 | TALE |
| 3518. | Glyma.19G229900.1.p | Glyma.19G229900 | GATA |
| 3519. | Glyma.19G234500.1.p | Glyma.19G234500 | C2H2 |
| 3520. | Glyma.19G236400.1.p | Glyma.19G236400 | NF-YC |
| 3521. | Glyma.19G236900.1.p | Glyma.19G236900 | bHLH |
| 3522. | Glyma.19G242600.1.p | Glyma.19G242600 | GeBP |
| 3523. | Glyma.19G243500.1.p | Glyma.19G243500 | Trihelix |
| 3524. | Glyma.19G244800.1.p | Glyma.19G244800 | bZIP |
| 3525. | Glyma.19G247600.1.p | Glyma.19G247600 | G2-like |
| 3526. | Glyma.19G248100.1.p | Glyma.19G248100 | MYB |
| 3527. | Glyma.19G248900.1.p | Glyma.19G248900 | ERF |
| 3528. | Glyma.19G249500.1.p | Glyma.19G249500 | Whirly |
| 3529. | Glyma.19G252600.1.p | Glyma.19G252600 | bZIP |
| 3530. | Glyma.19G253100.1.p | Glyma.19G253100 | ERF |
| 3531. | Glyma.19G254800.1.p | Glyma.19G254800 | WRKY |
| 3532. | Glyma.19G256700.1.p | Glyma.19G256700 | bHLH |
| 3533. | Glyma.19G256800.1.p | Glyma.19G256800 | ERF |
| 3534. | Glyma.19G257300.1.p | Glyma.19G257300 | LBD |
| 3535. | Glyma.19G257400.1.p | Glyma.19G257400 | MYB |
| 3536. | Glyma.19G257500.1.p | Glyma.19G257500 | Dof |
| 3537. | Glyma.19G259500.1.p | Glyma.19G259500 | NAC |
| 3538. | Glyma.19G259700.1.p | Glyma.19G259700 | NAC |
| 3539. | Glyma.19G260900.1.p | Glyma.19G260900 | MYB_related |
| 3540. | Glyma.19G261300.1.p | Glyma.19G261300 | B3 |
| 3541. | Glyma.19G262700.1.p | Glyma.19G262700 | ERF |
| 3542. | Glyma.19G264200.1.p | Glyma.19G264200 | MYB |

| 3543. | Glyma.20G000600.1.p | Glyma.20G000600 | NF-YB |
| --- | --- | --- | --- |
| 3544. | Glyma.20G001600.1.p | Glyma.20G001600 | TCP |
| 3545. | Glyma.20G001900.1.p | Glyma.20G001900 | MIKC_MADS |
| 3546. | Glyma.20G005800.1.p | Glyma.20G005800 | C2H2 |
| 3547. | Glyma.20G005900.1.p | Glyma.20G005900 | C2H2 |
| 3548. | Glyma.20G006000.1.p | Glyma.20G006000 | G2-like |
| 3549. | Glyma.20G006400.1.p | Glyma.20G006400 | SBP |
| 3550. | Glyma.20G007700.1.p | Glyma.20G007700 | bZIP |
| 3551. | Glyma.20G008700.1.p | Glyma.20G008700 | B3 |
| 3552. | Glyma.20G009800.1.p | Glyma.20G009800 | G2-like |
| 3553. | Glyma.20G011700.1.p | Glyma.20G011700 | MYB |
| 3554. | Glyma.20G012700.1.p | Glyma.20G012700 | C2H2 |
| 3555. | Glyma.20G013000.1.p | Glyma.20G013000 | MYB |
| 3556. | Glyma.20G014400.1.p | Glyma.20G014400 | HD-ZIP |
| 3557. | Glyma.20G016400.1.p | Glyma.20G016400 | Nin-like |
| 3558. | Glyma.20G016500.1.p | Glyma.20G016500 | Nin-like |
| 3559. | Glyma.20G017600.1.p | Glyma.20G017600 | WOX |
| 3560. | Glyma.20G023400.1.p | Glyma.20G023400 | LBD |
| 3561. | Glyma.20G024700.1.p | Glyma.20G024700 | FAR1 |
| 3562. | Glyma.20G026600.1.p | Glyma.20G026600 | C2H2 |
| 3563. | Glyma.20G028000.1.p | Glyma.20G028000 | WRKY |
| 3564. | Glyma.20G030500.1.p | Glyma.20G030500 | WRKY |
| 3565. | Glyma.20G031000.1.p | Glyma.20G031000 | ERF |
| 3566. | Glyma.20G032700.1.p | Glyma.20G032700 | bZIP |
| 3567. | Glyma.20G032900.1.p | Glyma.20G032900 | MYB |
| 3568. | Glyma.20G033300.1.p | Glyma.20G033300 | NAC |
| 3569. | Glyma.20G034000.1.p | Glyma.20G034000 | M-type_MADS |
| 3570. | Glyma.20G034100.1.p | Glyma.20G034100 | MYB |
| 3571. | Glyma.20G035200.1.p | Glyma.20G035200 | Dof |
| 3572. | Glyma.20G035300.1.p | Glyma.20G035300 | G2-like |
| 3573. | Glyma.20G035700.1.p | Glyma.20G035700 | B3 |
| 3574. | Glyma.20G035800.1.p | Glyma.20G035800 | B3 |
| 3575. | Glyma.20G036500.1.p | Glyma.20G036500 | ZF-HD |
| 3576. | Glyma.20G037100.1.p | Glyma.20G037100 | SRS |
| 3577. | Glyma.20G042000.1.p | Glyma.20G042000 | HSF |
| 3578. | Glyma.20G047500.1.p | Glyma.20G047500 | C2H2 |
| 3579. | Glyma.20G047600.1.p | Glyma.20G047600 | MYB |
| 3580. | Glyma.20G049200.1.p | Glyma.20G049200 | bZIP |
| 3581. | Glyma.20G050400.1.p | Glyma.20G050400 | M-type_MADS |
| 3582. | Glyma.20G051500.1.p | Glyma.20G051500 | EIL |
| 3583. | Glyma.20G053500.1.p | Glyma.20G053500 | FAR1 |
| 3584. | Glyma.20G060400.1.p | Glyma.20G060400 | CO-like |
| 3585. | Glyma.20G062300.1.p | Glyma.20G062300 | MYB_related |
| 3586. | Glyma.20G068700.1.p | Glyma.20G068700 | MYB_related |
| 3587. | Glyma.20G068900.1.p | Glyma.20G068900 | MYB_related |
| 3588. | Glyma.20G070000.1.p | Glyma.20G070000 | ERF |
| 3589. | Glyma.20G070100.1.p | Glyma.20G070100 | ERF |
| 3590. | Glyma.20G070300.1.p | Glyma.20G070300 | ZF-HD |
| 3591. | Glyma.20G075300.1.p | Glyma.20G075300 | ZF-HD |
| 3592. | Glyma.20G075400.1.p | Glyma.20G075400 | ZF-HD |
| 3593. | Glyma.20G075600.1.p | Glyma.20G075600 | ZF-HD |
| 3594. | Glyma.20G075800.1.p | Glyma.20G075800 | ZF-HD |
| 3595. | Glyma.20G076100.1.p | Glyma.20G076100 | FAR1 |
| 3596. | Glyma.20G082300.1.p | Glyma.20G082300 | MYB |
| 3597. | Glyma.20G088600.1.p | Glyma.20G088600 | bHLH |
| 3598. | Glyma.20G090700.1.p | Glyma.20G090700 | MYB |
| 3599. | Glyma.20G091200.1.p | Glyma.20G091200 | bHLH |
| 3600. | Glyma.20G093700.1.p | Glyma.20G093700 | C3H |
| 3601. | Glyma.20G097500.1.p | Glyma.20G097500 | TALE |
| 3602. | Glyma.20G097900.1.p | Glyma.20G097900 | MYB_related |
| 3603. | Glyma.20G099100.1.p | Glyma.20G099100 | C2H2 |
| 3604. | Glyma.20G099400.1.p | Glyma.20G099400 | WOX |
| 3605. | Glyma.20G103400.1.p | Glyma.20G103400 | TCP |
| 3606. | Glyma.20G107500.1.p | Glyma.20G107500 | bHLH |
| 3607. | Glyma.20G107900.1.p | Glyma.20G107900 | B3 |
| 3608. | Glyma.20G108000.1.p | Glyma.20G108000 | B3 |
| 3609. | Glyma.20G108100.1.p | Glyma.20G108100 | B3 |
| 3610. | Glyma.20G108300.1.p | Glyma.20G108300 | B3 |
| 3611. | Glyma.20G108400.1.p | Glyma.20G108400 | B3 |
| 3612. | Glyma.20G108600.1.p | Glyma.20G108600 | G2-like |
| 3613. | Glyma.20G109400.1.p | Glyma.20G109400 | C2H2 |
| 3614. | Glyma.20G111800.1.p | Glyma.20G111800 | MYB_related |
| 3615. | Glyma.20G113600.1.p | Glyma.20G113600 | bZIP |
| 3616. | Glyma.20G115300.1.p | Glyma.20G115300 | ERF |
| 3617. | Glyma.20G115600.1.p | Glyma.20G115600 | CO-like |
| 3618. | Glyma.20G115700.1.p | Glyma.20G115700 | LBD |
| 3619. | Glyma.20G117000.1.p | Glyma.20G117000 | MYB |
| 3620. | Glyma.20G121800.1.p | Glyma.20G121800 | C3H |
| 3621. | Glyma.20G130200.1.p | Glyma.20G130200 | bHLH |
| 3622. | Glyma.20G130800.1.p | Glyma.20G130800 | HB-PHD |
| 3623. | Glyma.20G132100.1.p | Glyma.20G132100 | FAR1 |
| 3624. | Glyma.20G133000.1.p | Glyma.20G133000 | LBD |
| 3625. | Glyma.20G133600.1.p | Glyma.20G133600 | bHLH |
| 3626. | Glyma.20G133700.1.p | Glyma.20G133700 | bHLH |
| 3627. | Glyma.20G136400.1.p | Glyma.20G136400 | M-type_MADS |
| 3628. | Glyma.20G136500.1.p | Glyma.20G136500 | M-type_MADS |
| 3629. | Glyma.20G136600.1.p | Glyma.20G136600 | M-type_MADS |
| 3630. | Glyma.20G136700.1.p | Glyma.20G136700 | M-type_MADS |
| 3631. | Glyma.20G136800.1.p | Glyma.20G136800 | M-type_MADS |
| 3632. | Glyma.20G142200.1.p | Glyma.20G142200 | HD-ZIP |
| 3633. | Glyma.20G148500.1.p | Glyma.20G148500 | TCP |
| 3634. | Glyma.20G149100.1.p | Glyma.20G149100 | CPP |
| 3635. | Glyma.20G150300.1.p | Glyma.20G150300 | HSF |
| 3636. | Glyma.20G152900.1.p | Glyma.20G152900 | bHLH |
| 3637. | Glyma.20G153000.1.p | Glyma.20G153000 | bHLH |
| 3638. | Glyma.20G153700.1.p | Glyma.20G153700 | MIKC_MADS |
| 3639. | Glyma.20G154200.1.p | Glyma.20G154200 | MIKC_MADS |
| 3640. | Glyma.20G154400.1.p | Glyma.20G154400 | TCP |
| 3641. | Glyma.20G155100.1.p | Glyma.20G155100 | ERF |
| 3642. | Glyma.20G155200.1.p | Glyma.20G155200 | ERF |
| 3643. | Glyma.20G156100.1.p | Glyma.20G156100 | FAR1 |
| 3644. | Glyma.20G156500.1.p | Glyma.20G156500 | HD-ZIP |
| 3645. | Glyma.20G156800.1.p | Glyma.20G156800 | HSF |
| 3646. | Glyma.20G157900.1.p | Glyma.20G157900 | MYB |
| 3647. | Glyma.20G158100.1.p | Glyma.20G158100 | MYB |
| 3648. | Glyma.20G160200.1.p | Glyma.20G160200 | Nin-like |
| 3649. | Glyma.20G161000.1.p | Glyma.20G161000 | C2H2 |
| 3650. | Glyma.20G162100.1.p | Glyma.20G162100 | GRAS |
| 3651. | Glyma.20G162800.1.p | Glyma.20G162800 | MYB_related |
| 3652. | Glyma.20G163200.1.p | Glyma.20G163200 | WRKY |
| 3653. | Glyma.20G166500.1.p | Glyma.20G166500 | Trihelix |
| 3654. | Glyma.20G166600.1.p | Glyma.20G166600 | Trihelix |
| 3655. | Glyma.20G166800.1.p | Glyma.20G166800 | Trihelix |
| 3656. | Glyma.20G168500.1.p | Glyma.20G168500 | ERF |
| 3657. | Glyma.20G172100.1.p | Glyma.20G172100 | NAC |
| 3658. | Glyma.20G172800.1.p | Glyma.20G172800 | ERF |
| 3659. | Glyma.20G175500.1.p | Glyma.20G175500 | NAC |
| 3660. | Glyma.20G176500.1.p | Glyma.20G176500 | C2H2 |
| 3661. | Glyma.20G176800.1.p | Glyma.20G176800 | GRAS |
| 3662. | Glyma.20G177600.1.p | Glyma.20G177600 | LBD |
| 3663. | Glyma.20G178500.1.p | Glyma.20G178500 | MYB |
| 3664. | Glyma.20G180000.1.p | Glyma.20G180000 | ARF |
| 3665. | Glyma.20G180100.1.p | Glyma.20G180100 | GATA |
| 3666. | Glyma.20G181000.1.p | Glyma.20G181000 | C2H2 |
| 3667. | Glyma.20G184000.1.p | Glyma.20G184000 | C2H2 |
| 3668. | Glyma.20G184100.1.p | Glyma.20G184100 | MYB |
| 3669. | Glyma.20G184200.1.p | Glyma.20G184200 | MYB |
| 3670. | Glyma.20G184300.1.p | Glyma.20G184300 | bHLH |
| 3671. | Glyma.20G184500.1.p | Glyma.20G184500 | Trihelix |
| 3672. | Glyma.20G185200.1.p | Glyma.20G185200 | MYB_related |
| 3673. | Glyma.20G185500.1.p | Glyma.20G185500 | bHLH |
| 3674. | Glyma.20G185800.1.p | Glyma.20G185800 | NAC |
| 3675. | Glyma.20G186200.1.p | Glyma.20G186200 | RAV |
| 3676. | Glyma.20G186400.1.p | Glyma.20G186400 | C2H2 |
| 3677. | Glyma.20G186500.1.p | Glyma.20G186500 | G2-like |
| 3678. | Glyma.20G188100.1.p | Glyma.20G188100 | Trihelix |
| 3679. | Glyma.20G189000.1.p | Glyma.20G189000 | Trihelix |
| 3680. | Glyma.20G192300.1.p | Glyma.20G192300 | NAC |
| 3681. | Glyma.20G192500.1.p | Glyma.20G192500 | NAC |
| 3682. | Glyma.20G193000.1.p | Glyma.20G193000 | C2H2 |
| 3683. | Glyma.20G193600.1.p | Glyma.20G193600 | G2-like |
| 3684. | Glyma.20G195900.1.p | Glyma.20G195900 | ERF |
| 3685. | Glyma.20G196400.1.p | Glyma.20G196400 | ERF |
| 3686. | Glyma.20G197000.1.p | Glyma.20G197000 | ERF |
| 3687. | Glyma.20G198500.1.p | Glyma.20G198500 | NF-YB |
| 3688. | Glyma.20G199300.1.p | Glyma.20G199300 | MYB |
| 3689. | Glyma.20G200500.1.p | Glyma.20G200500 | GRAS |
| 3690. | Glyma.20G203500.1.p | Glyma.20G203500 | ERF |
| 3691. | Glyma.20G203600.1.p | Glyma.20G203600 | ERF |
| 3692. | Glyma.20G203700.1.p | Glyma.20G203700 | ERF |
| 3693. | Glyma.20G206000.1.p | Glyma.20G206000 | C2H2 |
| 3694. | Glyma.20G207900.1.p | Glyma.20G207900 | C2H2 |
| 3695. | Glyma.20G209700.1.p | Glyma.20G209700 | MYB |
| 3696. | Glyma.20G214300.1.p | Glyma.20G214300 | ZF-HD |
| 3697. | Glyma.20G215700.1.p | Glyma.20G215700 | ERF |
| 3698. | Glyma.20G216600.1.p | Glyma.20G216600 | Dof |
| 3699. | Glyma.20G216700.1.p | Glyma.20G216700 | bHLH |
| 3700. | Glyma.20G219100.1.p | Glyma.20G219100 | C3H |
| 3701. | Glyma.20G223300.1.p | Glyma.20G223300 | MYB |
| 3702. | Glyma.20G224000.1.p | Glyma.20G224000 | Trihelix |
| 3703. | Glyma.20G224500.1.p | Glyma.20G224500 | bZIP |
| 3704. | Glyma.20G224700.1.p | Glyma.20G224700 | bHLH |
| 3705. | Glyma.20G228300.1.p | Glyma.20G228300 | GeBP |
| 3706. | Glyma.20G228500.1.p | Glyma.20G228500 | GeBP |
| 3707. | Glyma.20G231800.1.p | Glyma.20G231800 | bHLH |
| 3708. | Glyma.20G232400.1.p | Glyma.20G232400 | NF-YC |
| 3709. | Glyma.20G234500.1.p | Glyma.20G234500 | MYB_related |
| 3710. | Glyma.20G234900.1.p | Glyma.20G234900 | NF-YB |
| 3711. | Glyma.20G235100.1.p | Glyma.20G235100 | C2H2 |
| 3712. | Glyma.20G246400.1.p | Glyma.20G246400 | bZIP |
| 3713. | Glyma.20G247300.1.p | Glyma.20G247300 | RAV |
| 3714. | Glyma.20G247500.1.p | Glyma.20G247500 | C3H |
| 3715. | Glyma.20G248100.1.p | Glyma.20G248100 | bHLH |
| 3716. | Glyma.20G249900.1.p | Glyma.20G249900 | Trihelix |

| 3717. | Glyma.U001100.1.p | Glyma.U001100 | C2H2 |
| --- | --- | --- | --- |
| 3718. | Glyma.U009200.1.p | Glyma.U009200 | TALE |
| 3719. | Glyma.U009700.1.p | Glyma.U009700 | NF-YB |
| 3720. | Glyma.U012500.1.p | Glyma.U012500 | LBD |
| 3721. | Glyma.U013000.1.p | Glyma.U013000 | M-type_MADS |
| 3722. | Glyma.U013800.1.p | Glyma.U013800 | GRAS |
| 3723. | Glyma.U014700.1.p | Glyma.U014700 | bHLH |
| 3724. | Glyma.U015500.1.p | Glyma.U015500 | bHLH |
| 3725. | Glyma.U015800.1.p | Glyma.U015800 | SBP |
| 3726. | Glyma.U018000.1.p | Glyma.U018000 | MIKC_MADS |
| 3727. | Glyma.U018600.1.p | Glyma.U018600 | bZIP |
| 3728. | Glyma.U019400.1.p | Glyma.U019400 | C2H2 |
| 3729. | Glyma.U019800.1.p | Glyma.U019800 | ARF |
| 3730. | Glyma.U020900.1.p | Glyma.U020900 | LBD |
| 3731. | Glyma.U021300.1.p | Glyma.U021300 | Dof |
| 3732. | Glyma.U022100.1.p | Glyma.U022100 | AP2 |
| 3733. | Glyma.U024400.1.p | Glyma.U024400 | M-type_MADS |
| 3734. | Glyma.U024500.1.p | Glyma.U024500 | M-type_MADS |
| 3735. | Glyma.U025800.1.p | Glyma.U025800 | SBP |
| 3736. | Glyma.U027500.1.p | Glyma.U027500 | MYB |
| 3737. | Glyma.U028100.1.p | Glyma.U028100 | MYB |
| 3738. | Glyma.U028600.1.p | Glyma.U028600 | GRF |
| 3739. | Glyma.U028700.1.p | Glyma.U028700 | GRF |
| 3740. | Glyma.U029500.1.p | Glyma.U029500 | bZIP |
| 3741. | Glyma.U035800.1.p | Glyma.U035800 | bHLH |
| 3742. | Glyma.U037700.1.p | Glyma.U037700 | AP2 |
| 3743. | Glyma.U037800.1.p | Glyma.U037800 | bHLH |
| 3744. | Glyma.U038900.1.p | Glyma.U038900 | MYB |
| 3745. | Glyma.U039100.1.p | Glyma.U039100 | HD-ZIP |
| 3746. | Glyma.U039200.1.p | Glyma.U039200 | TALE |
| 3747. | Glyma.U040700.1.p | Glyma.U040700 | M-type_MADS |
